# Diencephalic and Neuropeptidergic Dysfunction in Zebrafish with Autism Risk Mutations

**DOI:** 10.1101/2024.01.18.576309

**Authors:** Mary E.S. Capps, Anna J. Moyer, Claire L. Conklin, Verdion Martina, Emma G. Torija-Olson, Morgan C. Klein, William C. Gannaway, Caleb C.S. Calhoun, Michael D. Vivian, Summer B. Thyme

**Affiliations:** Department of Neurobiology, The University of Alabama at Birmingham Heersink School of Medicine, Birmingham, AL, USA; Department of Biochemistry and Molecular Biotechnology, UMass Chan Medical School, Worcester, MA, USA

## Abstract

Hundreds of human mutations are linked to autism and related disorders, yet the functions of many of these mutated genes during vertebrate neurodevelopment are unclear. We generated 27 zebrafish mutants with presumptive protein-truncating mutations or specific missense variants corresponding to autism-risk alleles in 17 human genes. We observed baseline and stimulus-driven behavioral changes at larval stages, as well as social behavior differences in lines tested as juveniles. Imaging whole-brain activity revealed a near identical activity map for mutations in the unrelated genes *kmt5b* and *hdlbpa*, defined by increased activity mainly in the diencephalon. Mutating 7 of the 17 risk genes resulted in substantial brain size differences. Using RNA sequencing, we further defined molecular drivers of the observed phenotypes, identifying targetable disruptions in neuropeptide signaling, neuronal maturation, and cell proliferation. This multi-modal screen nominated brain regions, cell types, and molecular pathways that may contribute to autism susceptibility.

**Teaser:** Zebrafish screen uncovers diencephalon, social interaction, and neuropeptidergic signaling phenotypes in ASD risk mutants.

## Introduction

Recent exome sequencing studies have identified dozens of inherited and *de novo* protein-coding variants that significantly increase risk for autism spectrum disorders (ASDs) (*1–4*). Many of these genes with large effect *de novo* mutations cause rare neurodevelopmental disorders such as Coffin-Siris syndrome (*5–10*). Although autism is a feature of such disorders, the wide phenotypic spectrum resulting from mutating these associated genes raises the question to what extent molecular convergence underlying ASD exists. Even though there is convergence to general categories such as gene expression regulation and synaptic physiology (*1*, *3*), new functions of these genes are still being discovered (*11*). Understanding the mechanisms bridging genes and neural phenotypes is critical for developing new therapies, whether they act on shared pathways or require more precise approaches.

The zebrafish model offers unique features that facilitate much-needed large-scale characterization of the neurodevelopmental roles of ASD risk genes. Multiple groups have uncovered circuit, molecular, and behavioral functions of such genes using zebrafish. Previous findings included changes to brain structure and activity, altered sensory responsivity, reduced social interaction, and a role for microglia (*12–19*). External early brain development, optical transparency, small size, and low cost make larval zebrafish an ideal choice for genetic and pharmacological screens (*20*), and more complex medium-throughput behavioral studies such as social interaction are possible at juvenile stages (*21*). The technological advance of whole-brain activity mapping in zebrafish by measuring phosphorylated-Erk (pErk) levels, which respond rapidly to calcium influx following neuronal activity, offers a combination of throughput, sensitivity, and resolution that was not previously available (*22*). This method enabled detection of precise differences in the function of specific brain areas for genetic mutants, behavioral paradigms, and drug molecules (*17*, *23–28*).

Here, we leveraged the advantages of zebrafish to define the neural functions of 17 genes often mutated in ASD. After generating mutants with protein-truncating or missense mutations in zebrafish orthologs, we conducted a two-tiered phenotypic screen (**Fig. 1A**). All mutant lines were subjected to pErk brain activity mapping and a larval behavioral pipeline, including both baseline movement and responses to sensory stimulation. Most mutants had strong phenotypes in one or both assays. A subset of mutants underwent a juvenile social interaction assay, which has previously revealed phenotypes for similar mutants (*15*, *16*) and has validity for modeling core symptoms of ASD. To uncover molecular disruptions leading to observed phenotypes, we collected mRNA expression data for several mutants. Cellular differences in these lines were predicted using Gene Set Enrichment Analysis (GSEA) with published single-cell sequencing data (scRNA-seq) from a similar age and tissue (*29*). Brain activity maps and transcriptional changes nominated the diencephalon as a vulnerable brain area in ASD risk.

**Fig. 1.**
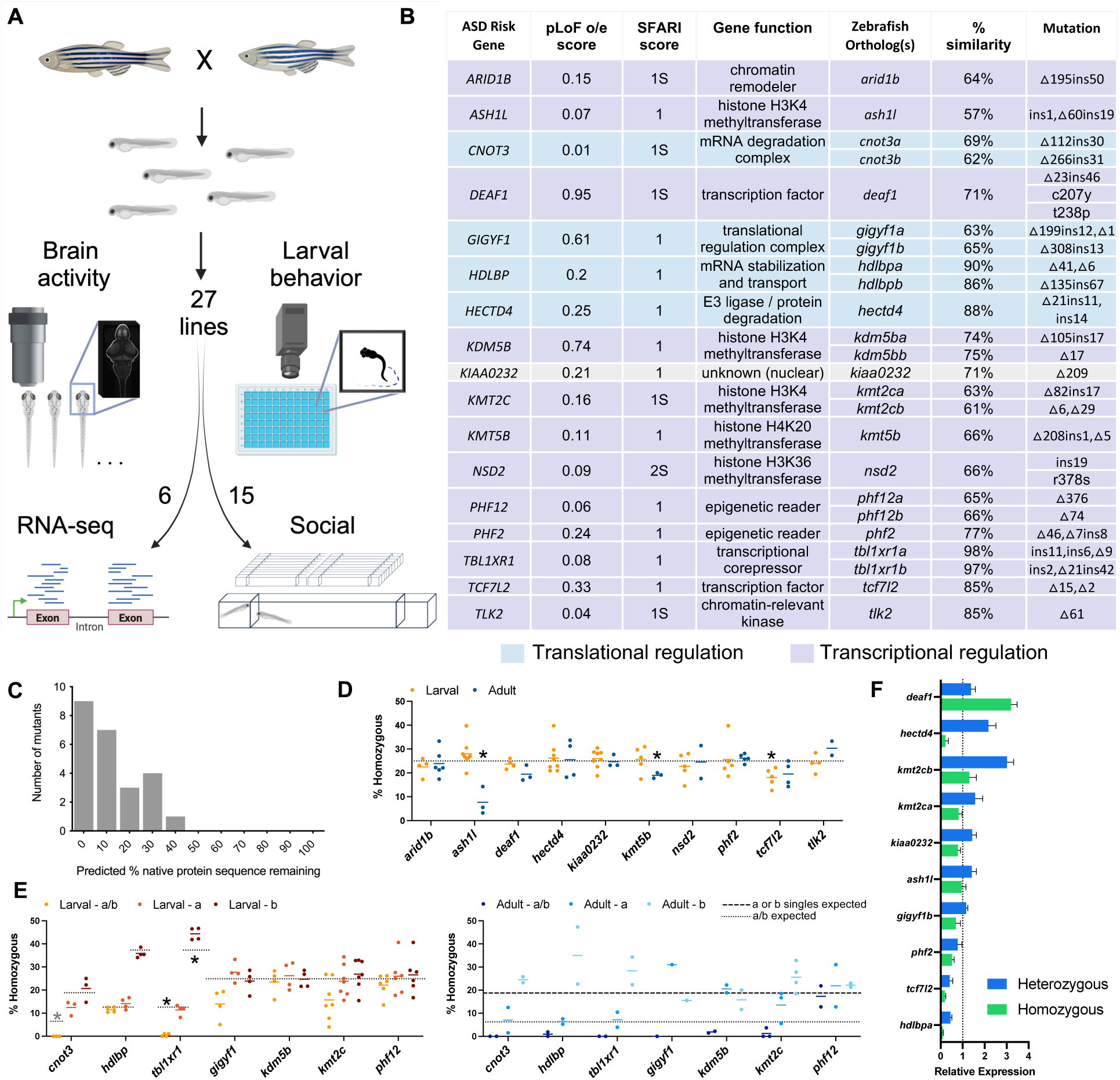
Generation and analysis of mutants for genes that increase risk for ASD. (**A**) Pipeline of the screen. Mutant lines generated with CRISPR/Cas9 were crossed together, and sibling larvae were assessed for changes to brain morphology, brain activity, and behavioral profiling in 96-well plates. Subsets of lines were further characterized with bulk mRNA-sequencing of dissected heads or social behavior. Created with BioRender.com. (**B**) Selected genes that increase risk for ASD and corresponding zebrafish CRISPR/Cas9 alleles. (**C**) Protein sequence predicted to remain for the lines with predicted truncating mutations, based on sequence alignment identity. (**D**) Survival of lines for single-gene alleles, where 25% is the expected Mendelian outcome for homozygous larvae derived from crosses of heterozygous parents. Significance was determined using a one sample t test compared to 25%. (**E**) Survival of double mutant lines, as larvae and adult. The dashed lines indicate the survival expectation for each genotype, as different parental combinations were used to produce the counted animals. The gray asterisk marks an incalculable p-value because the survival was always zero. (**F**) The relative change to mRNA levels using quantitative PCR for selected zebrafish mutant lines. The heterozygous and homozygous samples are normalized relative to the wild-type level, indicated by the dashed line at 1.

## Results

### Generating zebrafish mutants for ASD-risk genes

From the dozens of genes that increase risk for ASD, we selected those to include in our screen based on their known functions and relationships. We focused on two categories of genes (**Fig. 1B**): transcriptional regulators and a second, smaller category we refer to as translational regulators, which function post-transcriptionally through degradation or stabilization of protein or mRNA. We were particularly interested in those proteins known to interact with other ASD risk factors, directly or indirectly. For example, HECTD4 was included because it binds to GRIA3 (*30*) (**fig. S1**), identified through public databases (*31–33*). Both TBL1XR1 and TCF7L2 are effectors of Wnt signaling during neurodevelopment (*34*, *35*). Several of the selected genes also have similar functions, such as those involved in the deposition of methylation marks on histone tails (**Fig. 1B**). By choosing genes that could impact shared mechanisms, we expected to increase the chance of uncovering convergent phenotypes.

For orthologs of 17 human risk genes, we generated CRISPR/Cas9 mutations in coding regions predicted to yield premature stop codons and protein truncation (**Fig. 1B**, **Fig. 1C**, **Table S1**). Although not intentional, most selected genes were highly resistant to loss-of-function mutations in humans (o/e score < 0.35) (*36*) (**Fig. 1B**). When two zebrafish orthologs were present, both were mutated, and the a/b double mutant was used in all experiments unless only one ortholog was determined to confer the phenotype. For the *DEAF1* and *NSD2* genes, the reported ASD risk alleles are often missense instead of truncating (*1*, *37*, *38*). Thus, we generated point mutations corresponding to the human C207Y and T238P *DEAF1* and R378S *NSD2* variants. These lines were made by co-injecting an oligonucleotide with the desired changes (*39*) as well as additional silent nucleotide substitutions to introduce a restriction enzyme site for genotyping.

Mutating these genes impacted the survival of homozygous animals. For the single-gene alleles, crosses of heterozygous parents yielded near the expected 25% homozygous for all lines at the larval stages (**Fig. 1D**). Although a slightly lower number of *tcf7l2* larvae were observed (p = 0.0153), the difference was not significant in adulthood. Adult survival of the homozygous zebrafish was below expectation for the *kmt5b* and *ash1l* lines (p = 0.0113, 0.0359, respectively). For lines with mutations in two orthologs, the yield of healthy double homozygous larvae was significantly below expectation for *tbl1xr1*-ab (p < 0.001) and *cnot3*-ab (a/b survival was always 0) (**Fig. 1E**). Two lines had reduced survival as larvae that neared significance: *cnot3a* (larval p = 0.0736) and *gigyf1-*ab (larval p = 0.0565). Significance was not reached for any of the adult double-mutant data, as we did not process sufficient batches of adult animals during the duration of the screen. Comparing the survival of animals homozygous for individual orthologs can give insight into which ortholog contributes more to phenotypes in the double mutant. The adult survival outcomes, although insignificant, were lower than the Mendelian ratio for six of the seven double homozygous lines. Since the *cnot3a*, *hdlbpa* (p = 0.0587), and *tbl1xr1a* more strongly impacted survival than the “b” orthologs, the “a” ortholog was separately assessed for some neural phenotyping experiments.

Predicted protein-truncating mutations had a varied impact on mRNA levels (**Fig. 1F**, **fig. S2**). Most did not follow the expected pattern of a partial reduction in heterozygotes and further in homozygotes (e.g., *tcf7l2*). Many mutations did not yield significant changes between wild-type and homozygous siblings (e.g., *ash1l*). In several cases, the heterozygous samples showed a robust increase of the mRNA (e.g., *hectd4*), suggesting that genetic compensation stimulated by nonsense-mediated decay (*40*) upregulated the remaining wild-type allele. This full-length, wild-type mRNA would not be subject to nonsense-mediated decay, whereas both copies would undergo degradation in the homozygotes. Although mutants tested here have diverse expression outcomes, there is no clear relationship between the mRNA expression in the homozygotes and their neural phenotypes.

### ASD-risk mutations alter behavioral and sensory outputs

To uncover baseline and stimulus-driven behavioral abnormalities, we subjected homozygous mutant and sibling control larvae to monitoring and perturbation in 96-well plates from 4-7 dpf (**Fig. 2A**). We measured sleep (*41*), response to light and dark flashes (*42*, *43*), acoustic startle threshold, sensorimotor gating (*44*), and acoustic startle habituation (*43*). As previously (*17*), we calculated baseline parameter differences across sections of the multi-day experiment. Findings from these time windows and multiple related measures for both baseline (**Fig. 2B**) and stimulus-driven (**Fig. 2C**) results were integrated to summarize and visualize the outcomes.

**Fig. 2.**
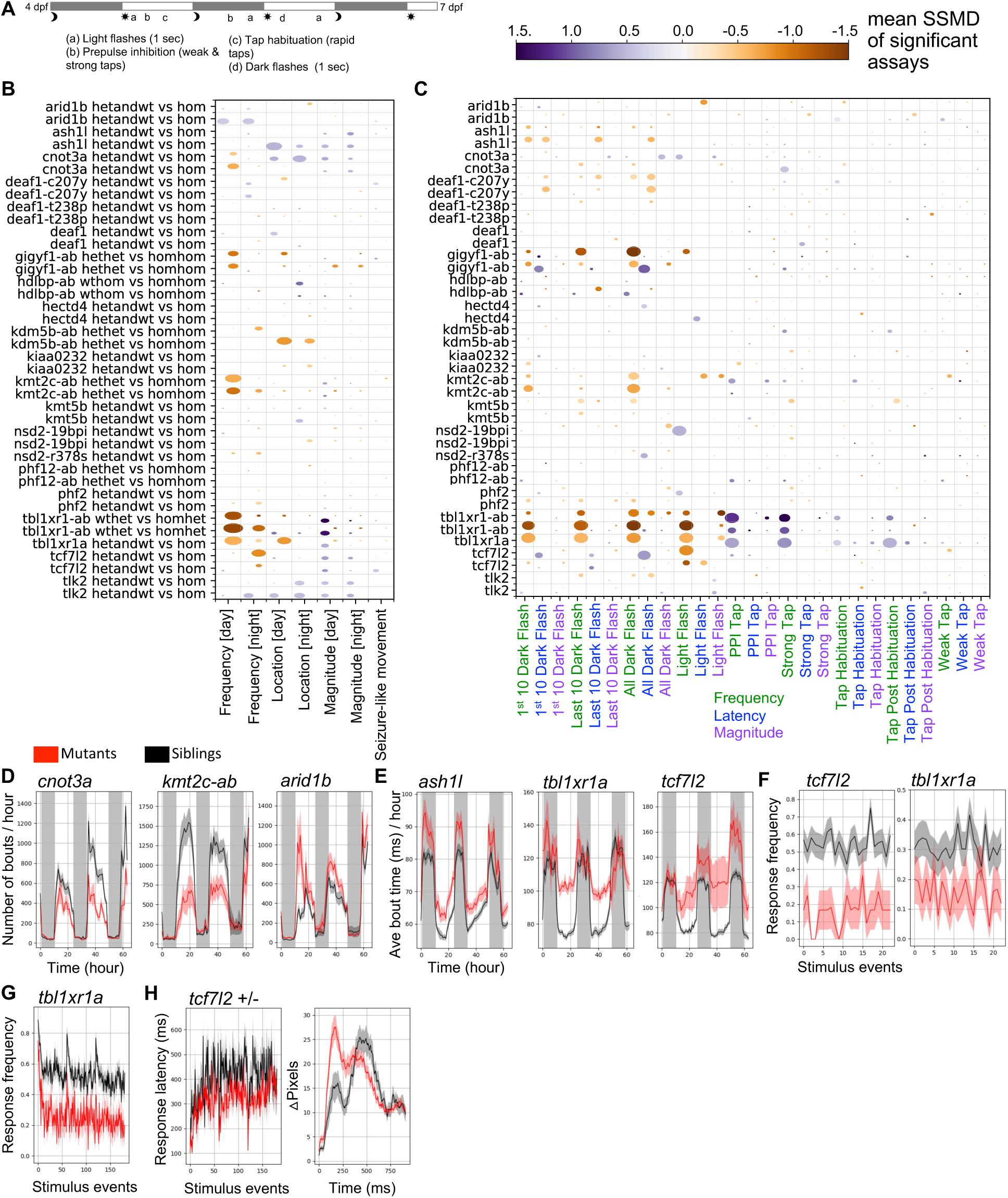
Behavioral phenotypes in zebrafish mutants. (**A)** Behavioral assay protocol. Frequency of movement, magnitude of movement, and location preferences were calculated for the baseline data. Stimulus responses were quantified from high-speed (285 frames-per-second) 1-second-long movies. (**B**) Merged summary of baseline behavioral phenotypes. The size of the bubble represents the percent of significant measurements in the summarized category, and the color represents the mean of the strictly standardized mean difference (SSMD) of the significant assays in that category. (**C**) Merged summary of stimulus-driven behavioral phenotypes. The bubble size and color are calculated the same as in panel B. (**D**) Examples of altered frequency of movement for the duration of the experiment. (**E**) Examples of increased movement magnitude (bout time) for the duration of the experiment. (**F**) Example of a reduced response frequency to one second light flashes (6 dpf evening light flashes). P-value = 0.002 (*tbl1xr1a*). P-value = <0.0001 (*tcf7l2*). (**G**) Reduced response frequency to one second dark flashes (all dark flash blocks, 6 dpf). P-value <0.0001. (**H**) Decreased response latency (left) for the *tcf7l2* heterozygous larvae compared to wild-type siblings. P-value = 0.0009. The response graph (right) is an average of larvae in the mutant and control groups for events where a response was observed. Plots of mutant compared to control groups in panels D, E, F, G, and H represent mean ± s.e.m.. The N for all experiments is available in Table S1. P-values are Kruskal-Wallis ANOVA.

Repeatable baseline movement differences included changes to the frequency of movement bouts as well as the magnitude of these movements (**Fig. 2B**, **fig. S3**, **fig. S4**, **fig. S5**, **fig. S6**). Reduced movement was more common than increased, as it was observed in four lines (*cnot3a*, *gigyf1*-ab, *kmt2c-*ab, *tbl1xr1*-ab), whereas only one line (*arid1b*) displayed increased daytime movement (**Fig. 2B**, **Fig. 2D**). Although the lines with reduced movement frequency were healthy when tested as larvae, as animals without swim bladders were not included, fewer survive to adulthood (**Fig. 1E**), indicating that reduced larval movement frequency could predict poor adult health for some genes. The most common shared behavioral difference was increased movement magnitude (**Fig. 2B**, **fig. S6**). The magnitude assessment includes measures such as the movement displacement and time elapsed during the movement. The main driver of this differences was the “bout time” measure, which was increased in multiple mutants (**Fig. 2E**, **fig. S6**), including those that were healthy in adulthood (e.g., *tcf7l2*).

Analysis of high-speed responses to visual and acoustic perturbations revealed that visual stimulation was more often affected in this mutant set (**Fig. 2C**, **fig. S7**, **fig. S8**, **fig. S9**, **fig. S10**). Animals from the *tbl1xr1a*, *gigyf1*-ab, and *kmt2c*-ab lines showed a robust reduction in their frequency of response to dark flashes, known to elicit the O-bend escape behavior (*43*). The *ash1l*, *kmt5b*, and *phf2* lines also exhibited a milder reduction in the dark flash response (**fig. S10**). The line with the most stimulus-driven phenotypes was *tbl1xr1a*, as light flash (**Fig. 2F**), dark flash (**Fig. 2G**), and acoustic stimuli responses (**Fig. 2E**, **fig. S10**) were affected. The light flash response frequency was reduced in the *tcf7l2* homozygous mutants (**Fig. 2F**). While most responses were decreased, the *deaf1*-c207y homozygous and *tcf7l2* heterozygous lines displayed a shorter response latency to dark flashes (**fig. S10**, **Fig. 2H**), indicating a higher stimulus sensitivity. Among those lines tested alongside wild-type sibling controls, the *tcf7l2* line was the only one with strong, repeatable phenotypes in the heterozygotes. The preponderance of visual phenotypes over acoustic in these mutants for ASD risk genes was not observed in a previous larger screen of zebrafish lines for schizophrenia-associated genes (*17*). It is possible that the distinct behavioral patterns of genes in different mutant groups reflect meaningful pathophysiology, as problems with the visual system have been reported in up to 50% of people with ASD (*45*, *46*).

### ASD-risk mutations alter brain activity and morphology

Using pErk brain activity mapping (*17*, *23–28*), we determined whole-brain activity and morphology differences between mutant larvae and sibling control populations. In this method, confocal stacks from animals stained with pErk and total-Erk (tErk) are registered to a common 6 dpf reference brain. The pErk / tErk ratio is a proxy of neuronal activity, and the deformation matrix generated by the registration indicates areas of increased or decreased brain size. The larvae were unstimulated for at least thirty minutes prior to rapid fixation. After determining the statistical differences between groups, the positive and negative signals in selected major brain regions (**Fig. 3A**) were quantified for activity (**Fig. 3B**) and structure (**Fig. 3C**).

**Fig. 3.**
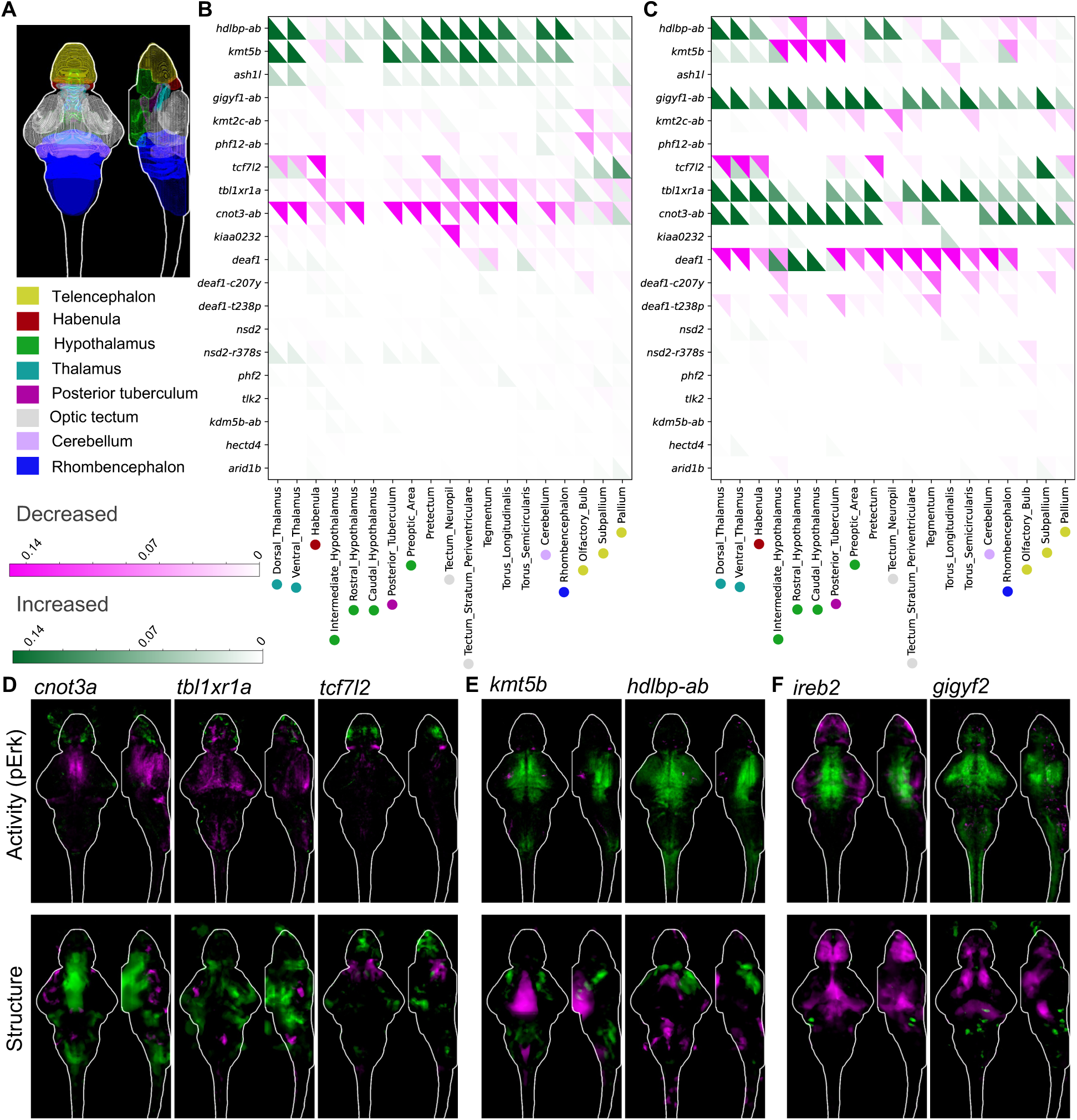
Whole-brain morphology phenotypes in zebrafish mutants. (**A**) Location of selected major regions in the zebrafish brain (*22*). (**B**) Summary of pErk comparisons between groups, where magenta represents decreased activity and green represents increased. The signal in each region was summed and divided by the total size of that region. Displayed data are the following comparisons from the run1 set: -/- versus +/+ for single gene mutants and -/-;-/- versus +/+;+/+ for all double mutants except *cnot3*-ab (-/-;+/- versus +/+;+/+). The N for all experiments is available in Table S1. (**C**) Summary of structure comparisons between groups, where magenta represents decreased size and green represents increased. The structural measure was calculated using deformation-based morphometry, as previously described (*17*). (**D**) Examples of activity and structural differences in mutants displayed as sum-of-slices projections (Z- and X-axes). Brain images represent the significant differences in signal between two groups. (**E**) Examples of activity and structural differences in mutants displayed as sum-of-slices projections (Z- and X-axes). Brain images represent the significant differences in signal between two groups. (**F**) Brain activity and structural maps from data in our previous screen of 132 schizophrenia-associated genes (*17*).

Approximately half of the mutants in ASD risk genes had substantial, repeatable changes to brain activity (**Fig. 3B**, **fig. S11**, **fig. S12**) or structure (**Fig. 3C**, **fig. S13**, **fig. S14**). Compared to a similar published screen of 132 genes associated with schizophrenia (*17*), more mutants had large brain structural changes (41% versus 12%). Both increased and decreased activity and size were observed in these lines. Mutated *tbl1xr1a* and *cnot3a* resulted in substantially decreased activity throughout the brain, particularly in the diencephalon and mesencephalon, and increased brain size in adjacent areas (**Fig. 3D**). The *tcf7l2* mutant line also had reduced activity in the diencephalon, mainly in the thalamus and habenula, but, unlike *tbl1xr1a* and *cnot3a*, the brain size was also reduced in these areas (**Fig. 3D**). This transcription factor was previously reported to regulate thalamic and habenular differentiation in mice and zebrafish (*47*, *48*). The reduced response to light stimulation observed in this line (**Fig. 2F**) is also compatible with disrupted habenula function (*49*).

The most similar brain activity maps were in the *kmt5b* and *hdlbp*-ab or *hdlbpa* lines (**Fig. 3B**, **Fig. 3E**). Both had increased activity throughout the brain, with an exceptionally strong signal in the thalamus and posterior tuberculum of the diencephalon. Mutants with shared brain activity phenotypes are relatively uncommon, as only a small number were discovered among the 132 genes associated with schizophrenia (*17*). In that screen, the most commonly overlapping areas were in the telencephalon and tectum. Thus, altered activity and structure of the diencephalon is more representative of the current mutant set. Among the set of 132 mutants, we identified two with a similar phenotype to *kmt5b* and *hdlbp*-ab: *ireb2* and *gigyf2* (**Fig. 3F**). Gigyf2 is itself an ASD risk factor that interacts with Gigyf1 and Cnot3 (fig. S1) (*50*, *51*), reinforcing the indication that the areas in this shared activity pattern are relevant to ASD. Although all four of these lines have an overlapping brain activity pattern, all have differing morphology maps (**Fig. 3E**, **Fig. 3F**). A lack of correlation between activity and structure is also present in the overall data (**Fig. 3B**, **Fig. 3C**), as several genes have mild brain activity phenotypes but substantial brain size differences (e.g., *deaf1*, *gigyf1*-ab). Taken together, we observed both shared and divergent phenotypes in brain size and structure.

### Differential gene expression in the *deaf1* allelic series

Next, to define the molecular underpinnings of the observed neural phenotypes, we selected mutant lines for further characterization with RNA sequencing (RNA-seq). To address how the missense mutations in *deaf1* related to truncating alleles, we compared sequencing from dissected heads of homozygous mutants and wild-type siblings. The missense mutations have similar, albeit non-identical, impacts on brain activity (**fig. S11**, **fig. S12**), brain structure (**fig. S13**, **fig. S14**), and behavior (**Fig. 2B**, **Fig. 2C**, **fig. S6**, **fig. S10**), and it is not clear how underlying gene expression differences yield phenotypic differences. We selected the *deaf1* allelic series for follow-up studies over the *nsd2* series, as the *deaf1* mutants had more divergent phenotypes; the *deaf1*-23d46i mutation caused more structural changes than the missense variants (**Fig. 3C**, **fig. S13**, **fig. S14**) and the *deaf1*-c207y mutation resulted in slightly increased night movement frequency (**fig. S6**) and decreased latency of dark flash responses (**fig. S10**). As Deaf1 is a transcription factor, it is expected to produce phenotypes by directly modulating expression of its target genes. The missense mutations are predicted to impact DNA-binding or transcriptional activity but could result in more complex outcomes than truncating mutations through retained interactions with binding partners (*6*, *52*).

We compared the gene expression outcomes across *deaf1* mutations and developmental time for the truncating (23d46i) line. Several core genes are robustly dysregulated in the same direction across multiple datasets (**Fig. 4A**, **fig. S15**). Comparing data from three independently generated lines allowed us to identify the changes to gene expression most important to the mutant phenotypes, as allele-specific expression of genes on the same chromosome in linkage disequilibrium with CRISPR/Cas9 mutation is common in outbred zebrafish lines (*53*) (**fig. S16**). Rather than removing all chromosome 25 genes from the analysis, the convergence of the three datasets allowed us to include those that are likely biologically meaningful (**fig. S16**). Observed gene expression differences from the three lines matched published literature results for orthologs of *DEAF1*, *EIF4G3*, *UBE2M*, and *RAC3* (*6*, *52*) (**Fig. 4A**). The *deaf1* transcription factor is known to be autoregulatory, as was observed in the *deaf-c207y* and *deaf1-t238p* differential expression. The RT-qPCR results (**Fig. 1F**, **fig. S2**) indicate our *deaf1* truncating allele is also upregulated; we expect that the different outcome in RNA-seq for this allele was due to the reliance on poly-adenylated transcripts in the preparation, which are degraded from the 3′ end by nonsense-mediated decay. Mutation of *RAC3* orthologs, consistently downregulated in mutants (**Fig. 4B**), causes a neurodevelopmental syndrome similar to mutations in *DEAF1* (*6*, *54*). This gene is likely involved in neuronal maturation and migration (*55*). Although the brain size of *deaf1* mutants is reduced (**Fig. 3C**, **fig. S13**, **fig. S14**), genes related to proliferation were not among those consistently dysregulated, except for *cdkn1bb* and *cdc27* (**Fig. 4A**, **Fig. 4C**).

**Fig. 4.**
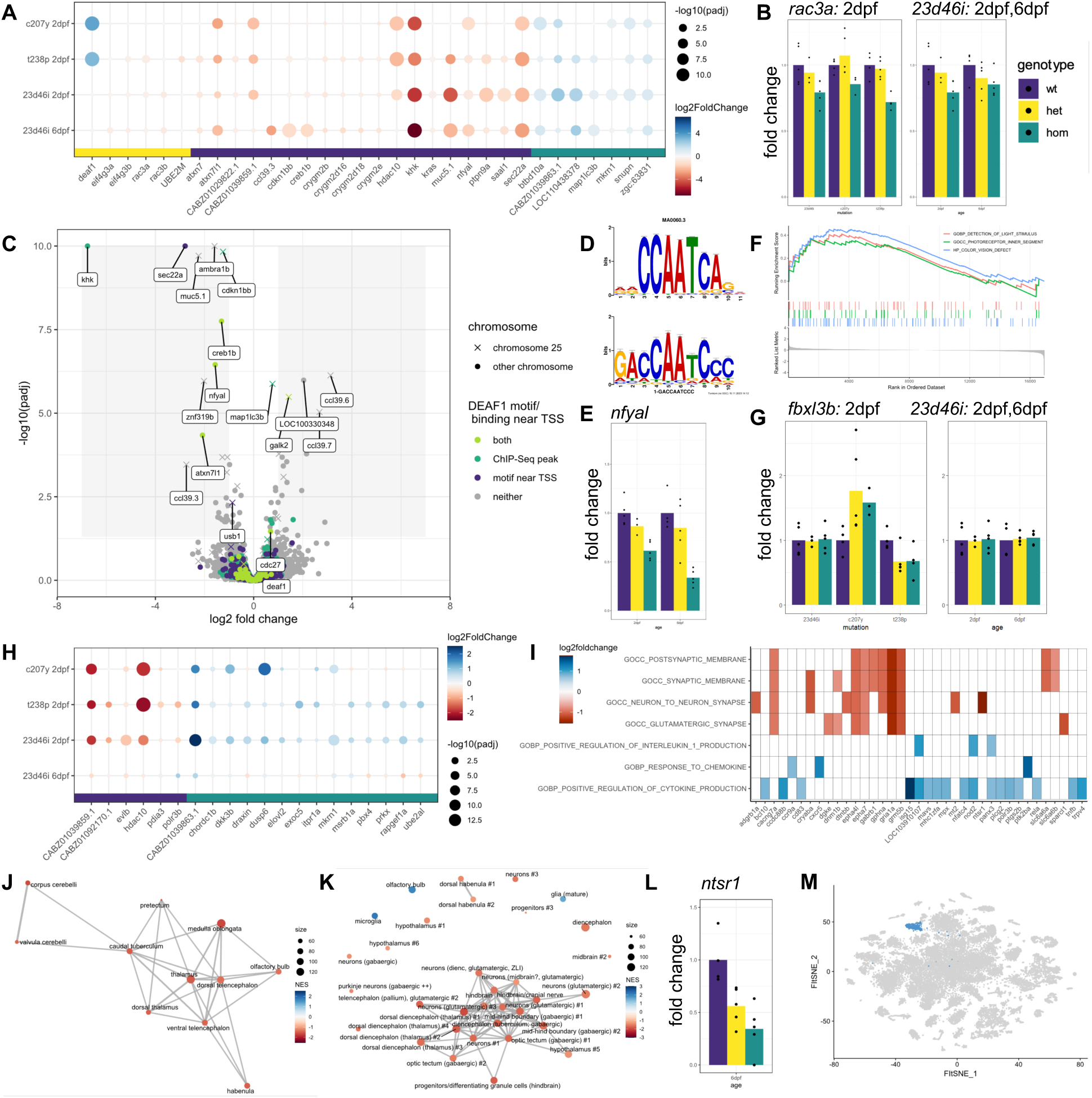
Differential gene expression analysis of *deaf1* mutant lines. (**A**) Genes shared across at least two of the four datasets. (**B**) Example expression of *rac3a*, one of the top downregulated genes the mouse hippocampus when *Deaf1* is downregulated (*52*). The graph on the left compares the 23d46i, c207y, and t238p lines at 2 dpf, while the graph on the right compares the 23d46i line at 2 dpf and 6 dpf. P-values for all plots of fold change are available in the tables in Data S1 (**C**) Volcano plot identifying putative direct targets of Deaf1 and other differentially expressed genes at 6 dpf in the 23d46i line. Genes located on the same chromosome as *deaf1* (chr25) are annotated. (**D**) *NFYA1* motif result from Tomtom (top) and query (bottom) from STREME. (**E**) Reduced expression of *nfyal* in the 23d46i lines. (**F**) GSEA of visual terms in the *deaf1*-c207y line. (**G**) Upregulation of *fbxl3b* in the c207y line compared to the other mutant alleles. (**H**) Genes that are most different between the 6 dpf sample and other three samples. (**I**) Selected GSEA C5 molecular signatures for the *deaf1*-23d46i 6 dpf data. (**J**) Network Enrichment Map of the ZFA enrichment results. Nodes represent enriched ZFA terms and edges join those terms with overlapping gene expression. (**K**) GSEA network plot with single-cell types from the same age. (**L**) Downregulation of *ntsr1*, marker of 34 cluster, in the 6 dpf data. (**M**) Cluster 34 in the published scRNA-seq data from 5 dpf heads (*29*).

Since Deaf1 is a transcription factor, some differentially expressed genes in the mutant lines are expected to be direct targets. DEAF1 is known to activate and repress its targets (*6*, *52*), and our mutations resulted in an approximately equal distribution of up- and down-regulated differentially-expressed genes (**Fig. 4C**, **fig. S17**). To discover putative direct targets, we used existing ENCODE DEAF1 ChIP-seq data (*56*) and profiled promoter regions for the known DEAF1 binding motif. DEAF1 ChIP-seq peaks are enriched at promoters (**fig. S18**). Searching the promoters of differentially expressed genes shared across the three lines identified DNA sequence motifs (**fig. S18**), including for the zebrafish ortholog of the DEAF1 target gene *NFYA* (**Fig. 4C**, **Fig. 4D**, **Fig. 4E**, **fig. S17**). It was previously noted that DEAF1 and NFYA have overlapping ChIP-seq peaks (*57*), as we also observe for genes with differential expression in our mutant lines (**fig. S18**). Knockdown of *NFYA* in cultured neurons results in downregulation of *KHK* and *ATXN71L* (*58*), the orthologs of which are downregulated in our data (**Fig. 4C**). Thus, some genes are differentially expressed in the mutant lines because of the downstream effect of dysregulating *nfyal* in conjunction with *deaf1*.

Comparing the differentially expressed genes from the three lines uncovered divergent molecular changes that may underlie phenotypic differences. The *deaf1*-c207y line displayed differences in visual behaviors, which were not present in the other two, yet a milder brain structural phenotype (**fig. S13**, **fig. S14**). Searching genes specifically misexpressed in this line uncovered an enrichment for genes involved in visual function (**Fig. 4F**, **fig. S19**), such as upregulation of the ortholog of recoverin (*rcvrnb*), which may enhance visual sensitivity (*59*). The night-specific movement frequency differences in this line could be due to changes to the *fbxl3b* gene (**Fig. 4G**), either directly or as a downstream effect of changes to vision, as Fbxl3 is involved in circadian rhythm maintenance in mammals (*60*). Several other genes show stronger dysregulation in the truncating and t238p lines than in *deaf1*-c207y (e.g., *draxin*, *evlb, pbx4, polr3b*) (**Fig. 4H**), which may reflect size differences of the 6 dpf brain. Early transcriptional differences specific to the 2 dpf datasets, such as *hdac10*, may be involved in initiating the pathways driving formation of the 6 dpf brain. Taken together, comparisons of the molecular changes in this allelic series highlighted shared and distinct disruptions underlying mutant neural and behavioral phenotypes.

Disrupted pathways and cell types that define the *deaf1* 6 dpf phenotype were predicted using Gene Set Enrichment Analysis (GSEA). Terms related to the formation of mature neuronal circuits and synapses were among the top differential terms from the C5 set of molecular signatures (**Fig. 4I**). Changes to immune system pathways may be related to the infection susceptibility in humans with *DEAF1* mutations (*6*). Using the Zebrafish Anatomy Ontology (ZFA) with CNS terms highlighted differential gene expression in areas that overlap with the observed diencephalic structural changes (**fig. S13**, **fig. S14**) such as thalamus (**Fig. 4J**, **fig. S20**). Refining this approach to the single-cell level with published 5 dpf brain scRNA-seq (*29*) uncovered reduced expression in the markers of mainly diencephalic cell types (**Fig. 5K**, **fig. S20**). Some differentially expressed genes were specific to small clusters in the scRNA-seq, such as *ntsr1,* a top marker of “hypothalamus #1” (**Fig. 4K**, **Fig. 4L**, **Fig. 4M**, **fig. S20**). Therefore, using GSEA to deconvolve bulk RNA-seq data can nominate potentially affected cell types if a corresponding scRNA-seq dataset exists.

**Fig. 5.**
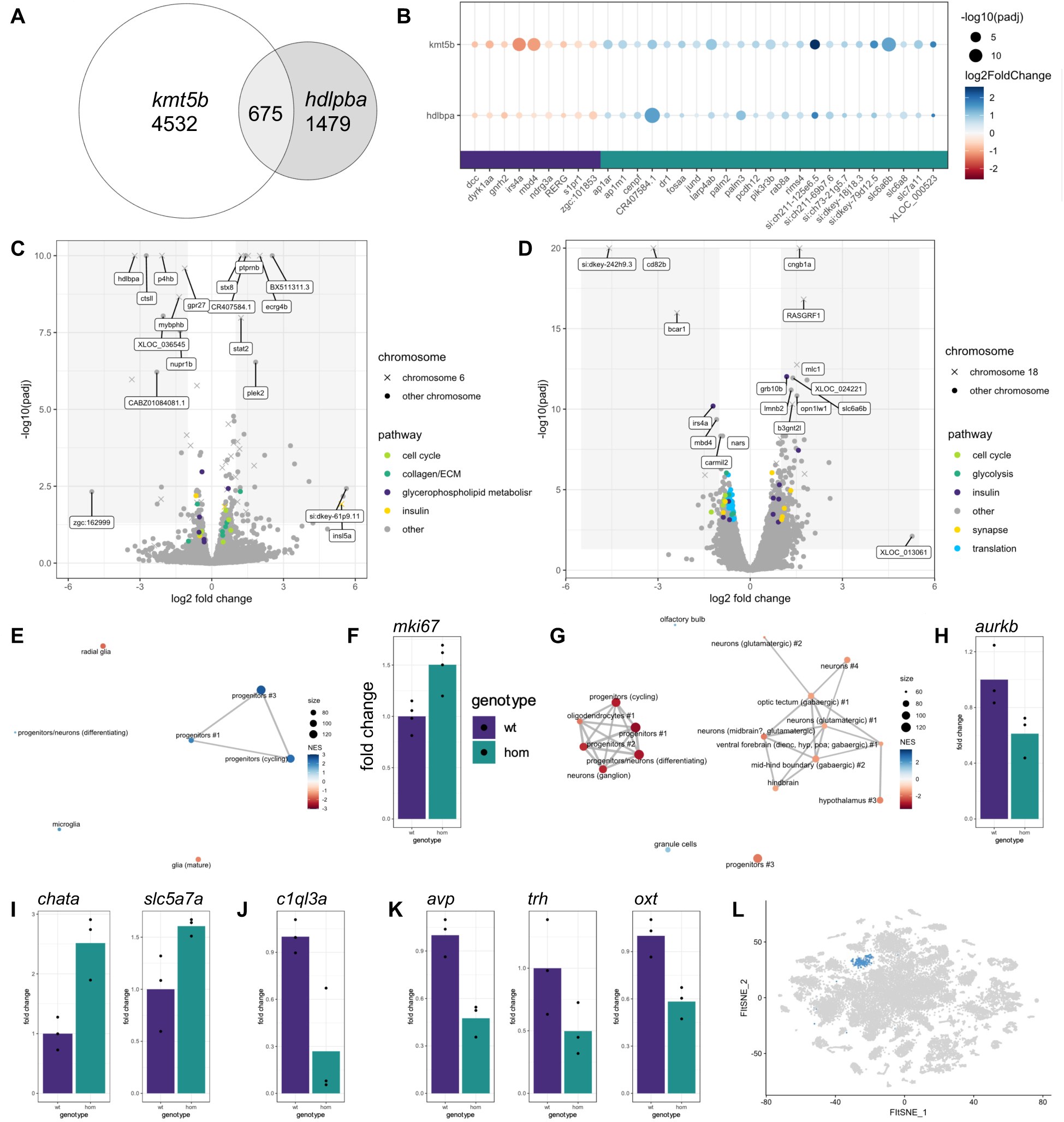
Differential gene expression analysis of *kmt5b* and *hdlbpa* mutant lines. (**A**) Euler diagram of the intersection of *kmt5b* and *hdlbpa* differentially expressed genes (p-value < 0.05). (**B**) Differentially expressed genes shared between *kmt5b* and *hdlbpa*. (**C**) Volcano plot of the *hdlbpa* RNA sequencing data with genes in selected pathways highlighted. Genes located on the same chromosome as *hdlbpa* (chr6) are annotated. (**D**) Volcano plot of the *kmt5b* RNA sequencing data with genes in selected pathways highlighted. Genes located on the same chromosome as *kmt5b* (chr18) are annotated. (**E**) GSEA network plot with single-cell types from the same age for *hdlbpa*. (**F**) Upregulation of *mki67*, a marker of proliferation, in *hdlbpa* mutants. (**G**) GSEA network plot with single-cell types from the same age for *kmt5b*. (**H**) Downregulation of *aurkb*, a marker of proliferation, in *kmt5b* mutants. (**I**) Upregulation of markers of cholinergic neurons in *kmt5b* mutants. (**J**) Downregulation of defined neuron type marker in *kmt5b* mutants. (**K**) Downregulation of markers of neuroendocrine cells, cluster 37, in *kmt5b* mutants. (**L**) Cluster 37 in the published scRNA-seq data from 5 dpf heads (*29*).

### Differential gene expression in mutants with shared brain activity patterns

Using the same RNA-seq and GSEA approaches as for *deaf1* (above), we next investigated the cellular and molecular basis underlying the shared brain activity patterns in the *kmt5b* and *hdlbpa* mutants. As with *deaf1*, we excluded genes located on the same chromosome prior to analysis (**fig. S21**). The *kmt5b* mutants yielded more differentially expressed genes than *hdlbpa*, and only a small proportion of the genes were shared between the two lines (**Fig. 5A**). Given that *kmt5b* modifies the epigenome (*61*) and *hdlbpa* regulates translation of secreted proteins (*62*), this difference is not unexpected. Among the shared genes (**Fig. 5B**), we noticed upregulation of *fosaa* and *jund*, immediate early genes that likely reflect the increased neuronal activity in these lines (**Fig. 3E**). Both lines also shared disrupted insulin signaling, identified by changes to several genes, including some shared, such as *irs4a* and *pik3r3b* (**Fig. 5B**), and some divergent, such as *insl5a* (*hdlbpa*) and *grb10b* (*kmt5b*) (**Fig. 5C**, **Fig. 5D**). A relationship between autism and insulin signaling has been hypothesized to occur via contributions of the PI3K/Tor pathway activation (*63*). While compelling, we remain cautious about this molecular overlap since two genes involved in the pathway are on the same chromosome as *hdlbpa* (**Fig. 5C**).

Mutation of either *kmt5b* or *hdlbpa* impacted progenitors in 6 dpf brains but in opposite directions. Cell cycle genes are upregulated in *hdlbpa* (**Fig. 5C**), as are progenitors using GSEA of the scRNA-seq clusters (**Fig. 5E**, **fig. S22**) and molecular signatures related to cell cycle and chromatin (**fig. S22**). Accordingly, the canonical proliferation marker *mki67* is upregulated (**Fig. 5F**). Given the mild brain structural differences (both increased and decreased), the upregulated transcription of proliferative markers in *hdlbpa* mutants is surprising, and its effect may be counteracted by currently unclear mechanisms. A reduction in cell cycle genes (**Fig. 5D**), progenitor markers (**Fig. 5G**, **fig. S23**), molecular signatures (**fig. S23**), and classical cell cycle markers (**Fig. 5H**) in the *kmt5b* mutants could explain their substantially smaller diencephalon (**Fig. 3E**).

The differentially expressed genes in the *kmt5b* dataset pointed to specifically disrupted, possibly missing, mature neuronal cell types in this mutant. We observed upregulation of *chata* (choline O-acetyltransferase) and *slc5a7a*, which transports choline into cholinergic neurons (**Fig. 5I**). As no cholinergic neuron cluster is annotated in the 5 dpf scRNA-seq data (*29*), and thus other marker genes are not known, it is unclear whether these transcriptional changes reflect decreased cholinergic optic tectum or hindbrain neurons (*64*) or reduced transcript expression. One of the most significantly reduced genes that mark specific neuronal cell types from the scRNA-seq is *c1ql3a* (**Fig. 5J**, **fig. S23**). The impacted *c1ql3a*-expressing glutamatergic types have an unclear spatial location but share some markers with other clusters thought to be at the midbrain-hindbrain boundary (**fig. S23**); the C1ql3 ortholog is expressed broadly throughout the mouse brain, although in discrete excitatory neuron subsets (*65*). While most implicated neuronal cell types were annotated as generally in the diencephalon or in unclear locations (*29*), the “hypothalamus #3” type stood out (**Fig. 5G**, **fig. S23**). The marker genes of this specific hypothalamic neuron type were *avp*, *oxt*, and *trh*, which were all downregulated (**Fig. 5K**). The reduction in all three marker genes indicates that this neuron type (**Fig. 5L**, **fig. S24**), known as neuroendocrine cells and specified by the Orthopedia transcription factor (*66*), is likely decreased in *kmt5b* mutants.

### ASD-risk mutations impair juvenile social behavior

Extending our mutant analysis beyond the larval stage, we characterized juvenile (21 dpf) social behavior. The behavioral test, which assesses a juvenile’s preference to interact with a conspecific over an empty control well (*67*), has uncovered reduced interaction in zebrafish mutants in ASD risk genes (*15*, *16*). We selected mutants to test that were healthy in adulthood (**Fig. 1D**, **Fig. 1E**) for the assay, excluding those that were lethal or had lower survival as adults (*cnot3a, hdlbpa, tbl1xr1a, kmt2c*-ab, *gigyf1*-ab, and *ash1l* homozygous). Following removal of a white comb that blocks visual access to the stimulus fish, an increase in time spent near the stimulus compared to the control well was observed (**Fig. 6A**, **fig. S25**). While some mutants interacted similarly to control siblings, *tcf7l2* and *kmt5b* had substantially reduced interaction with the stimulus fish (**Fig. 6A**). We also noticed that the *arid1b* line displayed a mild, repeatable reduction in interaction that was mainly at the beginning of the experiment (**Fig. 6B**). Thus, generating a single value per fish to represent the social preference is not representative of interaction over the entirety of the collected data. While the social preference per fish over the entire experiment was not significant for individual *arid1b* experiments, minutes 20-70 for sets 1 and 2 achieved significance, as did the combined data from three runs (**Fig. 6C**). Most lines did not display differences in social behavior (**Fig 6D**, f**ig. S25**).

**Fig. 6.**
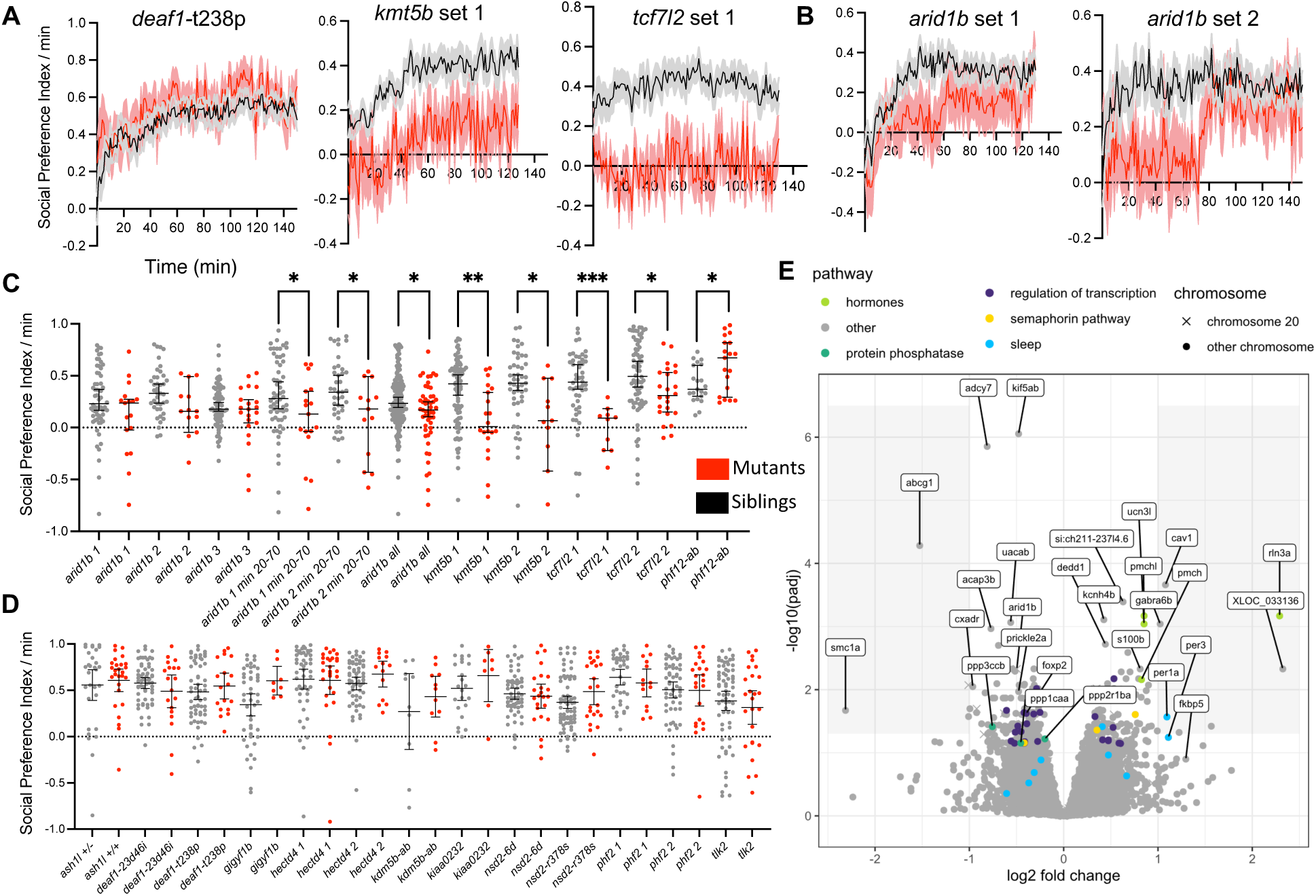
Juvenile social behavioral and transcriptomic differences. (**A**) Social Preference Index over the duration of the experiment, following removal of a white comb that blocks the view of the stimulus fish. The Social Preference Index per minute is defined as the number of frames in the zone near the stimulus fish (25% of the arena) minus the frames near the empty control well (25%) per minute. Plots of mutant compared to control groups represent mean ± s.e.m. The N for all social behavior experiments is in Table S1. (**B**) Social Preference Index for two *arid1b* datasets that show differences in interaction level over the duration of the experiment. Plots of mutant compared to control groups represent mean ± s.e.m. (**C**) Lines with social interaction phenotypes. The Social Preference Index per minute was averaged for each fish across the entire experiment (every point represents one animal). Significance was determined with the two-tailed Mann-Whitney test (*arid1b* 1 min 20-70: 0.0387, *arid1b* 2 min 20-70: 0.0407, *arid1b* all: 0.0228, *kmt5b* 1: 0.001, *kmt5b* 2: 0.0117, *tcf7l2* 1: 0.0001, *tcf7l2* 2: 0.0147, *phf12-ab*: 0.0286). Error bars represent the mean and 95% confidence interval. (**D**) Lines without social interaction phenotypes. (**E**) Volcano plot of differentially expressed genes in *arid1b* adult brain samples. N = 7 homozygous, 8 heterozygous.

With the discovery of this new juvenile behavioral phenotype, we selected the *arid1b* line for further characterization. Given the minimal neural and behavioral phenotypes in *arid1b* at larval stages (**Fig. 2**, **Fig. 3**), we had previously not chosen to explore its molecular differences. RNA-sequencing of pooled, dissected whole adult *arid1b* homozygous and heterozygous sibling adult brains revealed multiple disrupted pathways (**Fig. 6E**, **fig. S26**). Several neuropeptides were upregulated, including orthologs of *Rlxn3* and *Ucn3* (**Fig. 6E**), implicated in social behavior in rodents (*68*, *69*). Increased activity of the relaxin-3 receptor promotes social avoidance in rats and, thus, the significantly increased *rlxn3a* levels in the *arid1b* adult brain could contribute to the mutant’s behavioral phenotype.

## Discussion

Our work combines zebrafish behavioral analysis, brain activity and structural mapping, and RNA sequencing to investigate the neural roles of genes that increase risk for ASD. Many of our findings relate to previous literature and hypotheses regarding ASD, highlighting the relevance of the zebrafish model for rapid characterization of many genes simultaneously. Classic zebrafish genetic screens typically uncovered differences in fewer than 10% of mutants (*70–73*) compared to half of the mutants tested here, reinforcing the power of these methods for sensitive detection of neural phenotypes at a large scale. The high protein conservation of zebrafish also allowed us to test specific missense mutations. Although more challenging to generate than truncating mutations, these substitutions resulted in unique phenotypes that only partially overlapped with typical truncating loss-of-function alleles, underscoring the importance of precision modeling.

Comparing data across mutant lines identified points of convergence between genes with unrelated functions. At the anatomical level, *hdlbpa* and *kmt5b* shared increased brain activity in the diencephalon (**Fig. 3E**), including the posterior tuberculum and thalamus. At the molecular level, marker genes related to neuropeptide signaling (neurotensin, oxytocin, arginine-vasopressin, and relaxin-3) were dysregulated based on RNA-sequencing results. We found that GSEA using existing scRNA-seq data is a powerful and sensitive approach for deconvolving bulk sequencing, as diencephalic neuronal populations with small numbers relative to the whole brain were implicated by this method (**Fig. 4M**, **Fig. 5L**). While GSEA has been previously used to compare zebrafish bulk expression results with other species (*18*), leveraging zebrafish single-cell sequencing data from a similar age and tissue yielded testable hypotheses for these mutant lines. Together, our work highlights brain regions potentially vulnerable in ASD and provides insights into the molecular basis for these phenotypes.

Several of our main results align with previously proposed causes of ASD. Although it is a complex topic (*74*), an imbalance of excitation and inhibition (E-I) in ASD is an accepted model in the field (*75*, *76*). Increased activity throughout the brain, as we found shared in several mutants (**Fig. 3E**, **Fig. 3F**), would be expected from an increased E-I ratio. The classification of increased pErk as neuronal activity is further corroborated by the upregulation of the immediate-early genes *fosaa* and *jund* in mutant lines (**Fig. 5B**). Second, reduced neuropeptide signaling, particularly oxytocin, is another accepted model of ASD (*77*, *78*). Both oxytocin (*oxt*) and arginine-vasopressin (*avp*), downregulated in the *kmt5b* mutants (**Fig. 4K**), have been implicated in social behavior and ASD (*79*). Although relaxin-3 signaling, dysregulated in *aridb1* mutants with social behavior deficits, has not been convincingly implicated in ASD, it is known to impact social behavior in rodents (*69*). Changes to neuropeptidergic signaling pathways, such as the likely missing neuroendocrine cells in *kmt5b* mutants, may play a role in reduced social behavior in these zebrafish lines (**Fig. 6A**).

The diencephalon’s involvement in ASD and social behavior is also supported by previous literature. The hypothalamus (*80*) and thalamus (*81*) have both been implicated in ASD in studies of the human brain. The hypothalamus is a well-established hub of social behavior across species (*82*). In a recent study of zebrafish mutants in other ASD risk genes, the thalamus and posterior tuberculum also stood out among the brain regions with aberrant neural activity (*18*). This convergence is exciting in light of a recent study that identified the dorsal thalamus as the center of a tectothalamic circuit driving social affiliation (*83*). We report strongly reduced social interaction in the *tcf7l2* mutant (**Fig. 6A**), which has a substantially smaller thalamus (**Fig. 3D**), corroborating this connection.

Our study nominates directions for therapeutic development and highlights challenges. Normalization of E-I balance, the number of mature neurons in diencephalic clusters, and neuropeptide signaling are possible avenues of future intervention. Our data reinforced known molecular mechanisms (e.g., oxytocin) and suggested compelling new ones (e.g., relaxin-3). Gene expression analysis identified putative druggable targets in these lines, such as relaxin-3 receptors in *arid1b* and Rac3 in *deaf1*. However, mutants with shared brain activity patterns do not necessarily share transcriptomic profiles or anatomical changes (**Fig. 3**, **Fig. 5**). More work is needed to understand how these activity patterns arise, as many risk genes, including *hdlbpa* (*62*), impact physiology through changes to protein expression or synaptic physiology rather than mRNA dysregulation. Although less traditional than targeting molecular differences, it may be necessary to consider the correction of emergent phenotypes, such as increased brain activity, in developing a common treatment. Heterogeneity among people with ASD may also limit treatment scope. For example, intranasal oxytocin has yielded mixed outcomes as a treatment for reduced social abilities in ASD (*84*, *85*). Divergent mutant phenotypes may reflect a heterogeneity in people with ASD, as decreased oxytocin was observed only in the *kmt5b* mutants. While it remains unclear whether it will be possible to target shared pathways or whether precision approaches will be required, these mutant lines and their described phenotypes are a valuable starting point for translating genetic risk mutations into druggable mechanisms.

## Materials and Methods

### Zebrafish husbandry

Zebrafish experiments were approved by the UAB Institutional Animal Care and Use Committee (IACUC protocols 22155 and 21744). All animals, adults and experimental larvae, were maintained on a 14 h/10 h light/dark cycle at 28°C. Mutants were generated as in our previous work (*17*), in an Ekkwill-based strain. Mutant sequences, guide RNA targets, and genotyping primers are available in Table S1. Clean heterozygous mutant lines – at least F2s – were generated prior to experimentation by outcrosses to wild type. Larvae used for experimentation were grown in fish water with methylene blue in 150 mm petri dishes at a density of less than 160 per dish. Debris was removed twice prior to experimental tests beginning at 4 dpf for behavior and 6 dpf for pErk. Only healthy larvae with inflated swim bladders were used in all experiments. Control animals were always siblings from the same clutch, derived from a single parental pair, and all larvae were genotyped after experimentation.

### Brain activity and morphology

Phosphorylated-ERK (pErk) antibody staining was conducted as previously described (*17*). The following minor modifications were made to the standard pErk mapping protocol: paraformaldehyde (PFA) (Polysciences) was diluted to 4% with 1X Phosphate Buffered Saline without 0.25% Triton, the larvae were not fed on 5 dpf, the antigen retrieval step of incubation at 70°C for 20 minutes in 150 mM Tris-HCl pH 9.0 was skipped, the tissue permeabilization with 0.05% Trypsin-EDTA on ice was reduced to 30 minutes from 45, and the total ERK antibody (Cell Signaling, #4696) was used at 3:1000 instead of 1:500. Typically, the primary antibody exposure was for 2-3 days instead of one day. Images were collected using a Zeiss LSM 900 upright confocal microscope with a 20X/1.0 NA water-dipping objective. Larvae were removed from the agarose and genotyped after imaging. As previously (*17*, *22*), confocal stacks of the larval brains were registered to a standard zebrafish reference brain using Computational Morphometry Toolkit (CMTK). Using MapMAPPING (*17*, *22*), the significance threshold was set based on a false discovery rate (FDR) where 0.05% of control pixels would be called as significant for both the brain activity and structural measurements. The N for each assay is available in Table S1.

### Larval behavior

Larval behavior assays were conducted and analyzed as described (*17*, *86*) using custom-built behavioral systems. Briefly, the pipeline included the following on day 5: light flashes (9:11-9:25), mixed acoustic stimuli (prepulse, strong, and weak, from 9:38-2:59), three blocks of acoustic habituation (3:35-6:35); on the night following day 5: mixed acoustic stimuli (1:02-5:00), light flashes (6:01-6:20); on day 6: three blocks of dark flashes with an hour between them (10:00-3:00), mixed acoustic stimuli and light flashes (4:02-6:00). All code for behavior analysis is available at https://github.com/thymelab/ZebrafishBehavior, as are new scripts for merging individual measures and generating bubble plot visualizations (DownstreamAnalysis repository). The data are merged and shown in bubble plots with the possibility of two bubbles in each box because it is possible for some measures in the merged group to be increased (purple) and some decreased (orange). This is particularly true if there is a change in the phenotype across the duration of the experiment (e.g., different movement on night of 4 dpf versus 6 dpf). The N for each assay is available in Table S1.

### Juvenile social behavior

The social behavior arena was adapted from the published Fishbook assay (*21*), with minor modifications to the well sizes to maximize the number of chambers possible in our behavioral systems (20 test wells). The components of the arena and a modified version of our larval behavioral analysis code are available at https://github.com/thymelab/ZebrafishSocial. As in the Fishbook assay, 21 dpf larvae were placed in the test and stimulus chambers with a transfer pipette, cut to achieve a wider opening. The positions of the test fish were tracked for a minimum of an hour (15 frames-per-second) after removal of the white comb to expose them to the stimulus fish, using our previously published LabVIEW-based code (*86*). The health of the test and stimulus fish were checked in movies that correspond to the tracking and with activity measures. Unhealthy fish, likely damaged in the loading procedure and identified blind to genotype, were not included. To achieve a higher N, simultaneous tests were completed with additional animals from the same genetic cross. The modified code enables data from multiple behavioral systems to be combined. The caveat with combining the data is that each system could have subtle differences in light and temperature. Further, the code expects animals in different systems to initiate the experiment under the same conditions, and there were slight variations in the time between removal of the white comb and collection of position data. The N for each assay is available in Table S1.

### Real-Time Quantitative Reverse Transcription PCR (RT-qPCR)

The heads of anesthetized larval zebrafish (Syncaine MS 222 Fish Anesthetic) were dissected, frozen on dry ice, and stored at -80°C. The remaining bodies were collected and genotyped. Typically, four heads were combined and homogenized for each biological replicate, and 3-5 biological replicates were collected for each experimental group. RNA was extracted from the samples using the E.Z.N.A. MicroElute Total RNA Kit (Omega Bio-Tek R6834-02), including an incubation at room temperature for 15 min with E.Z.N.A. RNase-Free DNase I Set (Omega Bio-Tek E1091). RNA concentration and purity were determined using the ThermoScientific Nanodrop One Spectrophotometer. Complementary DNA (cDNA) synthesis was performed with iScript Reverse Transcription Supermix for RT-qPCR (Bio-Rad #1708840) using 200 ng of RNA template and manufacturer protocol. RT-qPCR was performed with SsoAdvanced Universal SYBR Green Supermix (Bio-Rad #1725271) using the manufacturer protocol and a Bio-Rad CFX96 Touch Real-Time PCR Detection System Module on a C1000 Touch Thermal Cycler. Each reaction contained 4 ng of cDNA input. Primers for qPCR (Table S1) were designed to span two exons downstream of the target gene mutation and created with all considerations recommended by the manufacturer’s manual. The *rpl13a* gene was used as the reference with published primers (*87*). Raw data from RT-qPCR assays was collected using CFX Maestro Software (Bio-Rad), and analysis of gene expression was performed using the comparative method delta-Ct (dCT)*.

### RNA sequencing

Bulk RNA-Sequencing was performed based on the SMART-Seq2 protocol (*88*) with modifications. Zebrafish head collection and RNA extraction were performed as specified for the above RT-qPCR method. For reverse transcription, 6 µL RNA was combined with the following: 0.3 µL 10 µM RT Oligo (5′-AGACGTGTGCTCTTCCGATCT(30)VN-3′)

3 µL 10 mM dNTP mix (Thermo-Fischer, R0192)

0.3 µL RNase Inhibitor 40 U/µL (Life Technologies, AM2694)

2.5 µL 1 M Trehalose (Life Sciences, TS1M-100).

Samples were incubated at 72°C for 3 minutes and immediately placed on ice following incubation. The following mix was then added to each sample:

6 µL 5X Maxima RT Buffer (Thermo-Fischer, EP0751)

0.3 µL Maxima RNase H-Minus RT 200 U/µL (Thermo-Fischer, EP0751)

10.35 µL 1 M Trehalose (Life Sciences TS1M-100)

0.3 µL 1 M MgCl2 (Invitrogen, AM9530G)

0.3 µL 10 µM TSO (5′-AGACGTGTGCTCTTCCGATCTNNNNNrGrGrG-3′)

0.75 µL RNase Inhibitor 40 U/µL (Life Technologies, AM2694)

Samples were mixed gently via pipette and were incubated in a thermal cycler at 50°C for 90 minutes, followed by a 5-minute inactivation period at 85°C. Seven µL of the reverse transcription reaction was added to the following master mix, and PCR was performed:

5 µL Nuclease Free Water

0.5 µL 10 µM PCR Oligo (5′-AGACGTGTGCTCTTCCGATCT-3′)

12.5 Kapa HiFi HotStart PCR ReadyMix (KAPA Biosystems, KK2601)

Whole Transcriptome Amplification was performed for 14 cycles (67°C annealing for 20 seconds and 6-minute extension at 72°C). PCR products were purified with a 0.8X AMPure XP SPRI (Beckman-Coulter, A63881) clean-up according to the manufacturer instructions, and DNA was eluted from beads with 10 µL Elution Buffer. Sample concentration was determined with the Promega QuantiFluor ONE dsDNA System (Promega, E4870) and Quantus Fluorometer (Promega, E6150), and then diluted to 0.2 ng/µL. Nextera XT DNA Library Preparation (Illumina, FC-131-1096) was carried out according to the manufacturer protocol, except for using a final volume of 25 µL. The DNA libraries were normalized by the Heflin Genomics Core using qPCR and sequenced with either a NovaSeq 6000 or NextSeq 550. The typical read depth was 30 million per sample, and any samples with abnormally low reads (<5 million) were discarded.

### RNA sequencing analysis

Single-end sequencing reads were aligned to GRCz11 release 104 using the Lawson Lab Zebrafish Transcriptome Annotation version 4.3.2 (*89*) using the STAR aligner (2.7.3a-GCC-6.4.0-2.28) (*90*). The resulting raw counts files were normalized using rlog counts method in DESeq2 (*91*). Prior to normalization, transcripts with zero counts in fewer than four samples were removed (8-15 samples were typically normalized together). An example DESeq2 RScript and input files are provided at https://github.com/thymelab/BulkRNASeq. Two sets of *arid1b* adult brain samples were collected and batch effects were removed with *ComBat-seq* (*92*) prior to analysis (script also on GitHub). Fold-change tables and analysis code for all experiments are in Data S1. Normalized counts are available from Zenodo. Raw counts and sequencing files are available from GEO (GSE253405).

To identify chromosome 25 genes dysregulated across the 2 dpf *deaf1* mutants, chromosome 25 genes were first filtered to include those with p-values < 0.05 in any of the three 2 dpf homozygous versus wildtype comparisons, then further filtered to include genes with log2 fold changes > 0.3 or < -0.3 across two of the three comparisons and the same sign in the third comparison. Based on these criteria, 487 chromosome 25 genes not shared between mutants were excluded from further analysis. To identify genes dysregulated across *deaf1* mutant conditions, genes were filtered to include those with p-values < 0.05 in two or more comparisons, resulting in 187 upregulated genes and 139 downregulated genes. For the *kmt5b* and *hdlbpa* datasets, all genes on the same chromosome as the mutation were removed.

### DEAF1 ChIP-Seq and motif annotation

DEAF1 ChIP-seq peak data from human K562 cells were downloaded from ENCODE (ENCFF813TRY) (*56*), and peaks were annotated using the annotatePeak function from the ChIPseeker_1.38.0 package (*93*) with TSS region spanning -2000 to +2000 of the annotated TSS and flanking gene distance of 250. Plots of binding distance from TSS and feature distribution based on chromosomal position were created using the plotAvgProf and plotAnnoBar functions. Human genes with DEAF1 peaks annotated at promoters were converted to zebrafish gene names using the orthogene_1.8.0 package with “keep both species” mapping option. Zebrafish orthologs of *ATXN7L1* and *EIF4G3* were manually annotated. Zebrafish genes with the DEAF1 binding motif (TCG(N5-N11)TCG) within -100 and -1 of the TSS were annotated using RSAT - genome-scale dna-pattern (https://rsat.france-bioinformatique.fr/metazoa/genome-scale-dna-pattern_form.cgi). Human DEAF1 (ENCSR387SYS) and NFYA (ENCSR163VTS) peaks were visualized on custom tracks using the UCSC Genome Browser (*94*). To identify motifs in the promoter regions of differentially expressed genes, the upstream sequences (-300 to -1) of the 326 differentially expressed genes shared between the *deaf1* mutants were retrieved from Ensembl Release 110 (http://rsat.sb-roscoff.fr/retrieve-ensembl-seq_form.cgi) and submitted to STREME (*95*) using right alignment. Motifs identified by STREME were compared to known vertebrate motifs using Tomtom Motif Comparison Tool (*96*).

### Gene Set Enrichment Analysis

Gene Set Enrichment Analysis (GSEA) (*97*) was performed using the GSEA function from the clusterProfiler_4.10.0 R package. For enrichment of genes based on chromosomal position, a custom gene set was created using RefSeq Genes and Gene Predictions from the GRCz11/danRer11 assembly, and genes were sorted by absolute value of log2 fold change before performing GSEA. For GSEA using molecular signatures, the zebrafish C5 ontology gene set (containing GO:BP, GO:CC, GO:MF, and HPO terms) was obtained using the msigdbr function from the msigdbr_7.5.1 package (*98*), and genes were sorted by log2 fold change. To perform GSEA using the zebrafish anatomy and development ontology (ZFA), the ZFA OBO file (zebrafish_anatomy.obo) and expression data for wildtype fish (wildtype-expression_fish.txt) were downloaded from zfin.org (*99*, *100*). Expression of genes reported for each anatomy term was associated with all ancestors of the term, and gene sets were created for all descendants of head terms and for all descendants of CNS terms with at least 80 genes per anatomy term. To perform GSEA using 5 dpf single-cell data (*29*), the single-cell object GSE158142_zf5dpf_cc_filt.cluster.rds was downloaded from NCBI GEO, and markers for all annotated clusters were identified using the FindAllMarkers function from the Seurat_5.0.0 package (*101*). The top 250 markers for each cluster were used to create custom gene sets for all clusters and for all CNS clusters (excluding eye). The enrichplot_1.22.0 package was used to create enrichment plots (gseaplot2 function), heatmaps (heatplot function), and network plots (emapplot function). The complete results of all GSEA analyses are available in Data S1.

## Supporting information

Data S1 - RNA-seq analysis files

## Acknowledgments

We thank the Heflin Genomics Institute at UAB, UAB fish facility staff, the Research Computing team at UAB, and the UAB Department of Neurobiology for supporting this study. The UAB Cheaha supercomputer was used for all analyses in this work. We also thank Justine Pinskey for helpful comments on the manuscripts and the following undergraduate students and technicians for experimental assistance supporting this study: Lynne Zhou, Mandy Chen, Katlyn Foy, Lalit Pusapati, Jaqueline Martinez, Brandie Cline, Vaishnavi Balaji, Gretchen Kioschos, and Elijah Quiñones.

## Funding

This research was funded by the following sources:

Klingenstein-Simons Fellowship Award in Neuroscience (SBT)

NARSAD New Investigator Award from the Brain and Behavior Research Foundation (SBT)

Undiagnosed Disease Network Gene Function Study (SBT)

Simons Foundation SFARI Pilot Award (SBT)

Jerome Lejeune Foundation Fellowship (AJM)

## Author contributions

SBT conceived of the study, analyzed imaging and behavioral data, and wrote the manuscript with contributions from MSC and AJM. MSC collected all RNA-sequencing and RT-qPCR data, as well as contributed to brain activity imaging, social behavior, and larval behavior studies. AJM analyzed the RNA-sequencing data. CLC established the social behavior assay, using previous code developed by MDV, as well as collected most of the brain activity and structural imaging data. VM generated all mutant lines and collected some larval behavioral data. EGT, MCK, WCG, and CCSC contributed to the experimental data in this work by genotyping and propagating zebrafish lines, running larval behavior experiments, and collecting brain imaging data. EGT also contributed to the initial RNA-sequencing analysis pipeline.

## Data and materials availability

All data are available in the main text, the Supplementary Materials, or appropriate databases. All mutants described in the paper are available from ZIRC. Code is available from https://github.com/thymelab (several repositories, listed throughout the Materials and Methods). Processed behavioral and imaging data is available from Zenodo under DOI 10.5281/zenodo.10530481, and raw behavioral and imaging files that are too large for Zenodo are available upon request. RNA-sequencing data, raw counts and fastq files, is available from GEO under access number GSE253405.

