## Supplementary Materials for "Diencephalic and Neuropeptidergic Dysfunction in Zebrafish with Autism Risk Mutations"

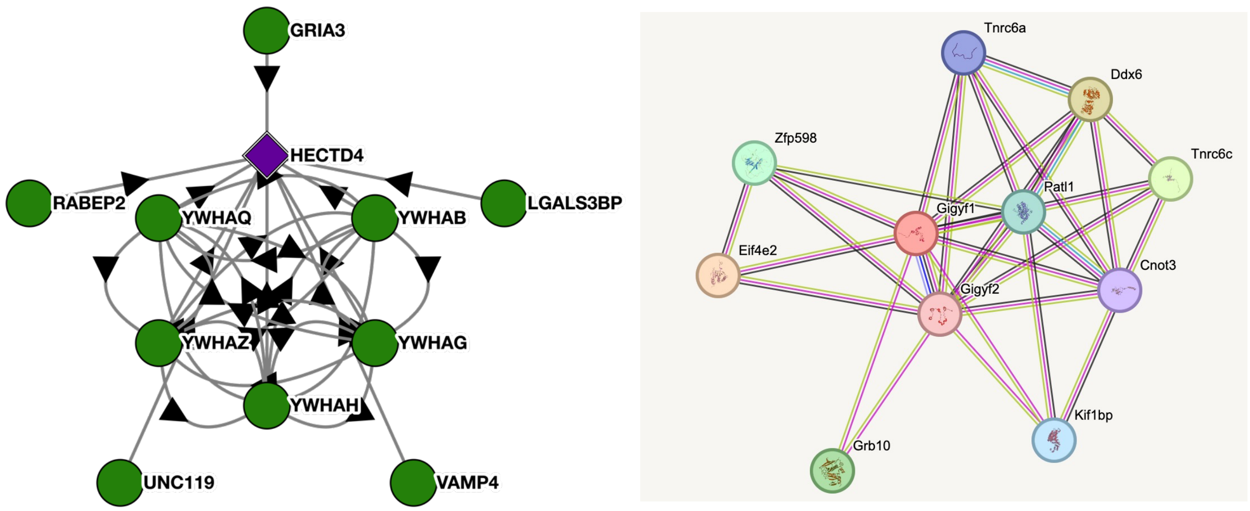


**Fig. S1. Protein-protein interaction networks from public databases.**
Protein-protein interaction network for HECTD4 (Bioplex 3.0) and Cnot3/Gigyf1/Gigyf2 (STRING 12.0).

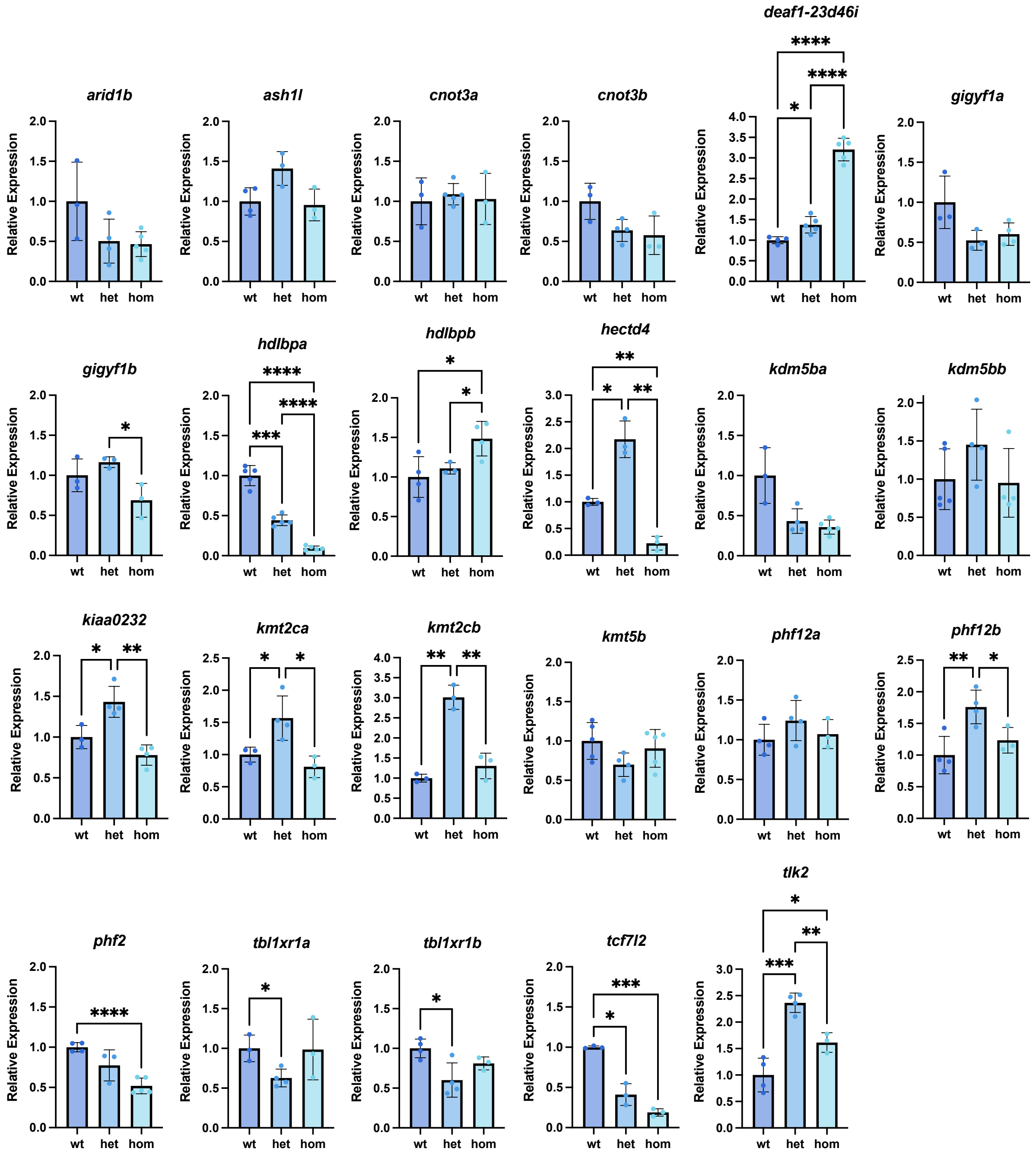


**Fig. S2. Quantitative RT-PCR for truncating mutants.**

Each biological sample consisted of 4 dissected heads, and 3-5 biological samples were included for each genotype. Duplicated genes were tested separately with the other ohnolog as wild type. The expression of each gene was normalized to *rpl13a*. Statistical significance was calculated with the Brown-Forsythe and Welch ANOVA.

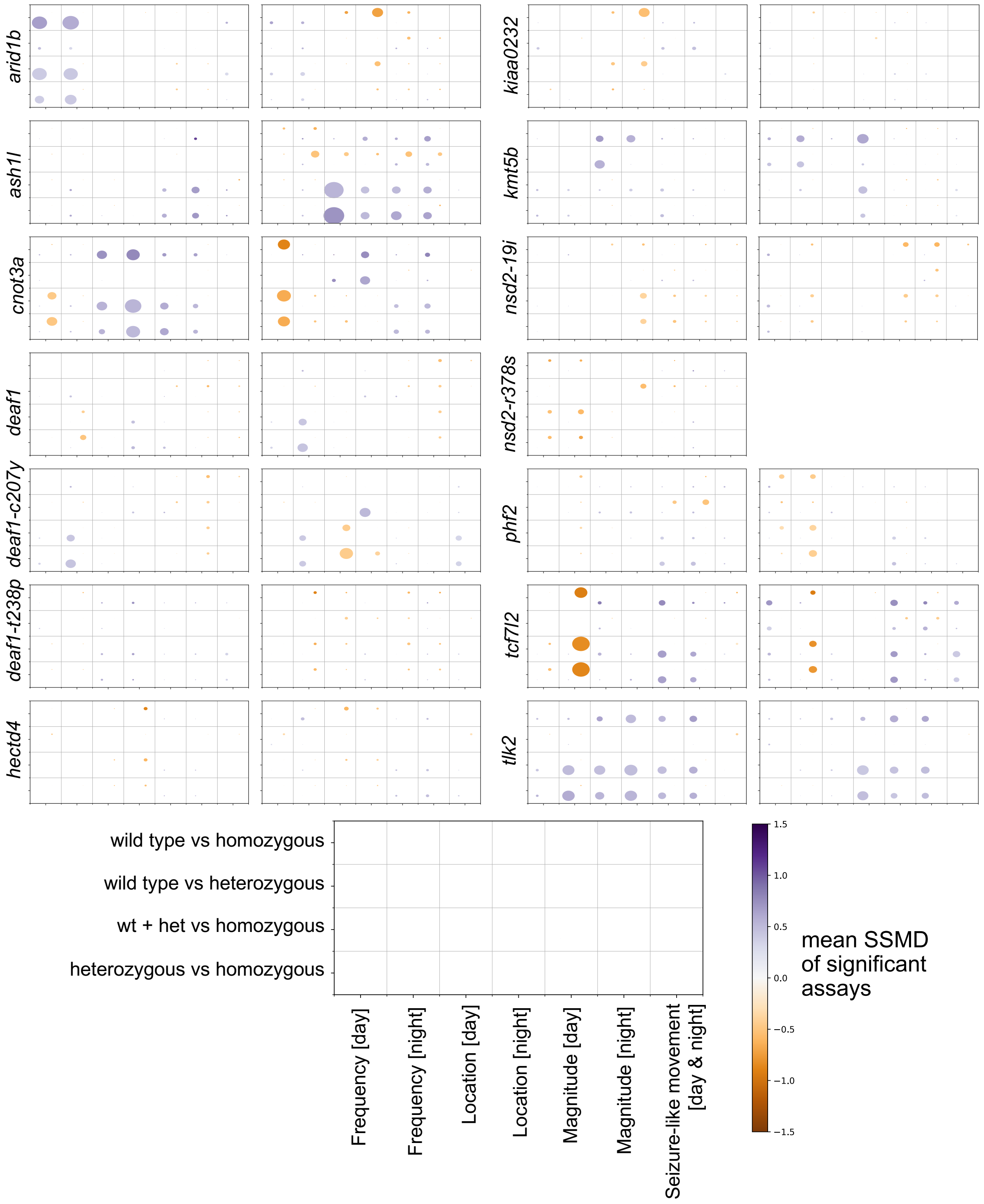


**Fig. S3. Baseline behavior summary data for single-gene mutants.**

Baseline behavior dot plots for all sibling comparisons from a heterozygous parental in-cross for zebrafish mutants representing ASD-risk genes with a single ortholog. Dot size corresponds to the percent of significant assays in the category (e.g., Magnitude). Replicate experiments using different parental pairs are shown side-by-side for all mutants except *nsd2*-r378s. All N are in Table S1.

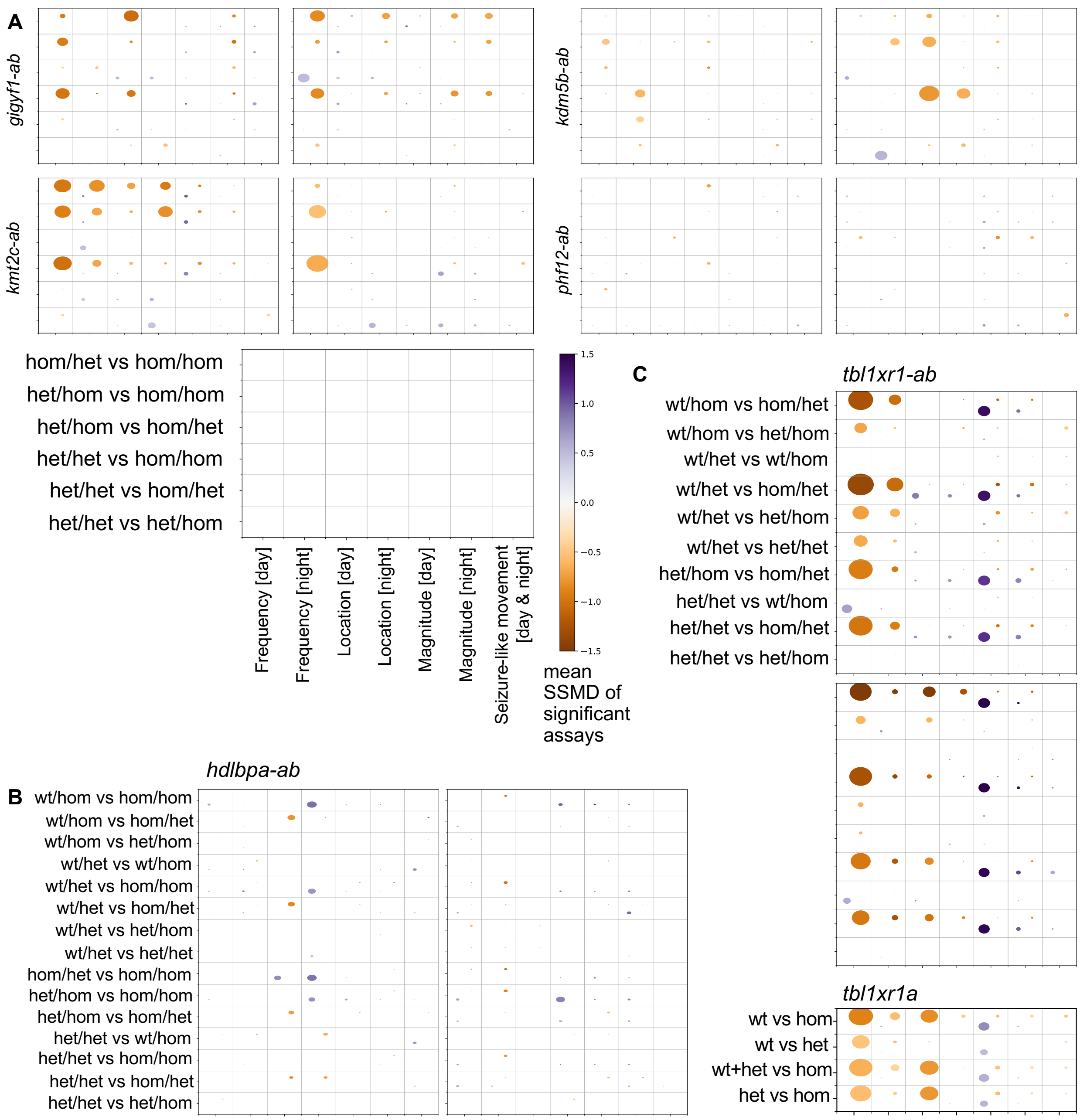


**Fig. S4. Baseline behavior summary data for double-gene mutants.**

Baseline behavior dot plots for all sibling comparisons from a parental cross for zebrafish mutants representing ASD-risk genes with two orthologs. Dot size corresponds to the percent of significant assays in the category (e.g., Magnitude). Two experiments using different parental pairs are shown side-by-side or above-below (*tbl1xr1-ab* only). (**A**) Parental cross: heterozygous/homozygous with homozygous/heterozygous. (**B**) Parental cross: heterozygous/heterozygous with heterozygous/homozygous. (**C**) Parental cross for *tbl1xr1-ab*: heterozygous/heterozygous with heterozygous/homozygous with homozygous-homozygous larvae dying before the experiment starting at 4 dpf. Parental cross for *tbl1xr1a*: standard single-mutant heterozygous with heterozygous. All N are in Table S1.

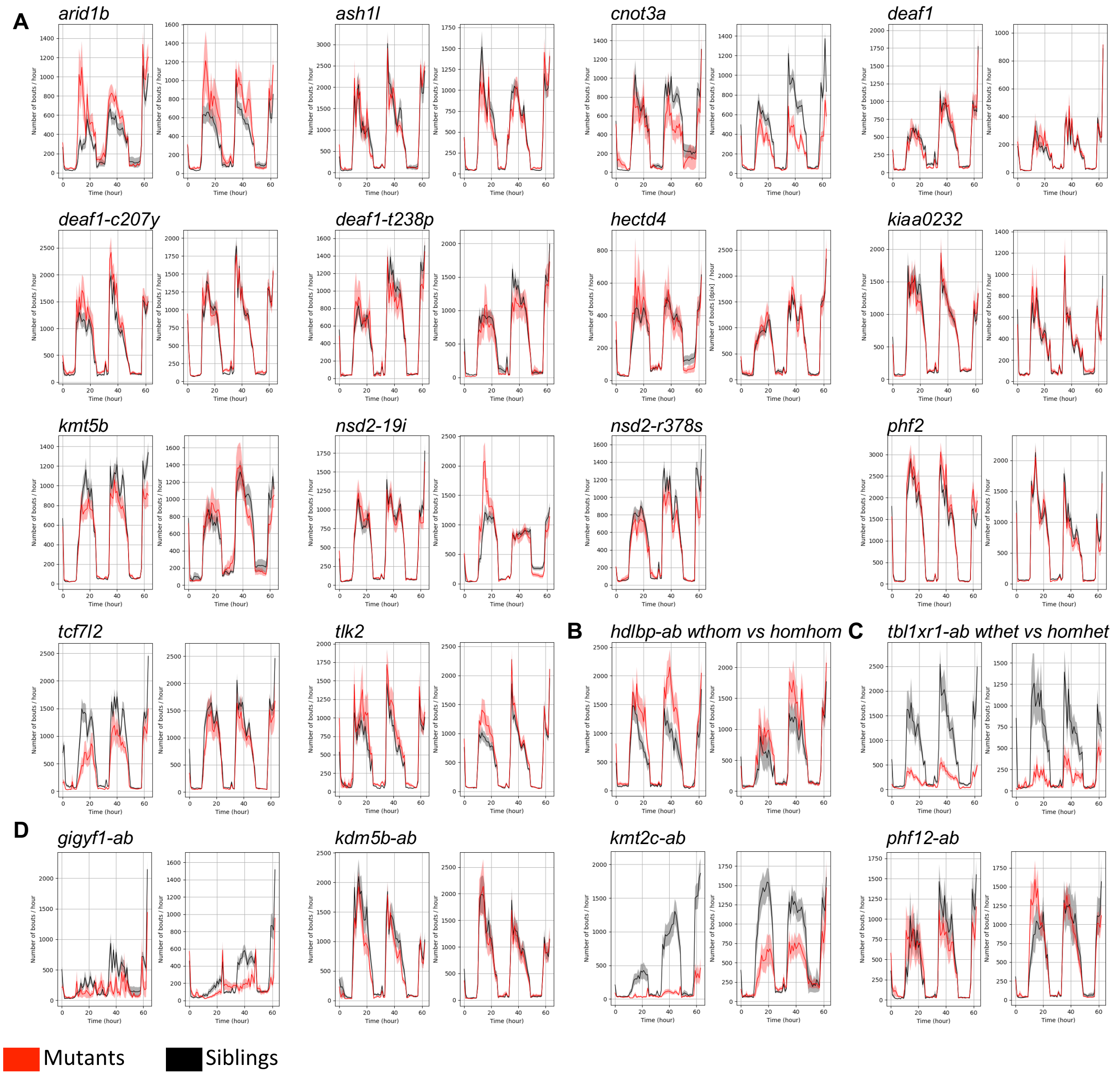


**Fig. S5. Frequency of motion example plots for each mutant.**

Frequency of motion measured as number of bouts/hour over the entire duration of the experiment. The biological replicates are shown side-by-side. All N are in Table S1. (**A**) All single-gene mutants with heterozygous parental in-crosses. The heterozygous and wild-type siblings (black) are compared to the homozygous (red). (**B**) The *hdlbp-ab* parental cross was heterozygous/heterozygous with heterozygous/homozygous, and this sibling comparison is wild-type/homozygous (black) versus homozygous/homozygous (red). (**C**) The *tbl1xr1-ab* parental cross was heterozygous/heterozygous with heterozygous/homozygous, and this sibling comparison is wild-type/heterozygous (black) versus homozygous/heterozygous (red), as all homozygous/homozygous larvae die before 4 dpf. (**D**) Double-gene mutants with parental crosses of heterozygous/homozygous with homozygous/heterozygous. This sibling comparison is heterozygous/heterozygous (black) versus homozygous/homozygous (red).

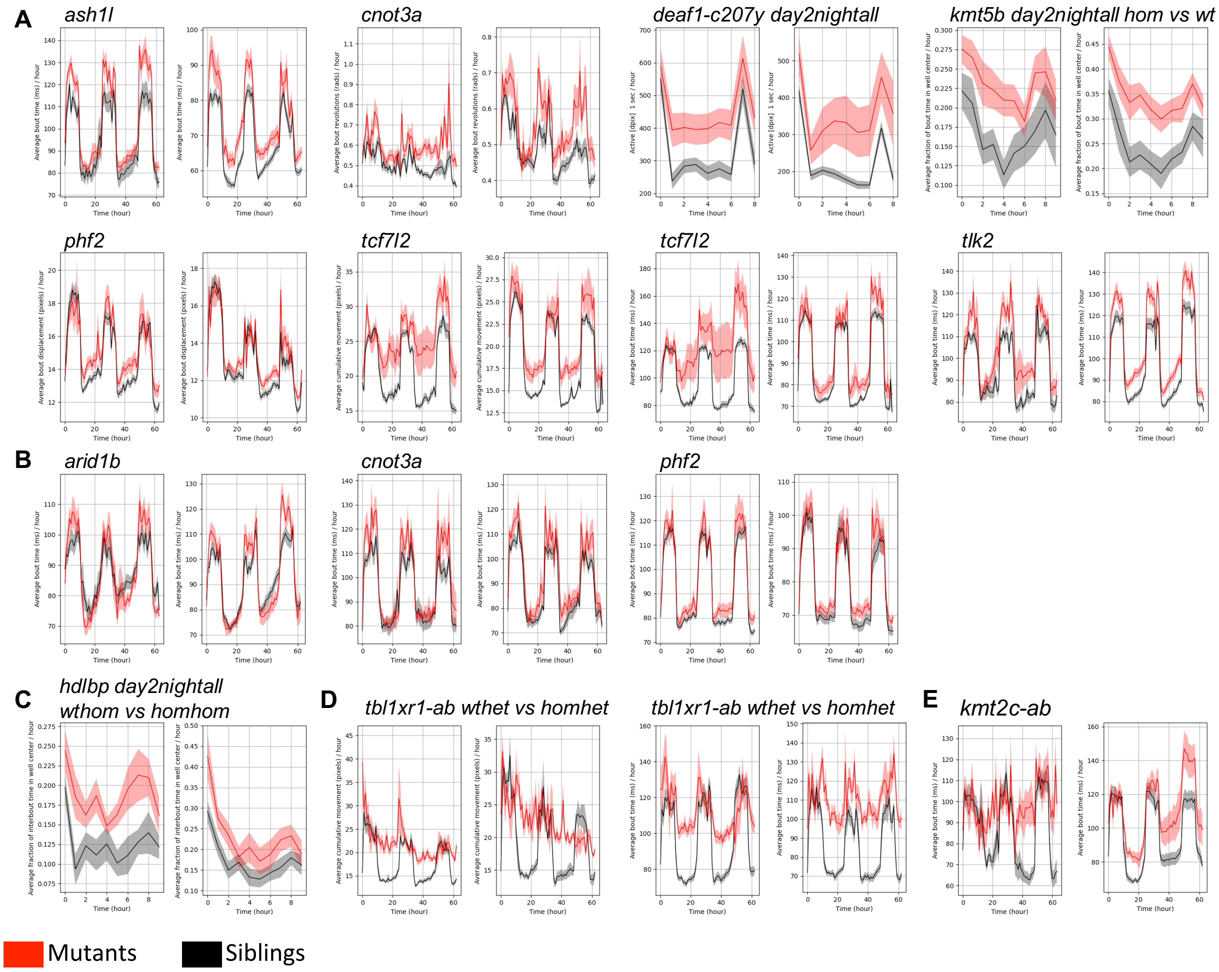


**Fig. S6. Baseline motion phenotypes.**

Raw measurements plots corresponding to significant, repeatable findings. When the entire experiment is shown (all examples where there is no label of a shorter time window) significance is not calculated (individual smaller time windows within the experiment are significant, p-values in Zenodo files). Replicate experiments from different parental crosses are displayed side-by-side. All N are in Table S1. (**A**) Single-gene mutants from heterozygous parental in-crosses. The heterozygous and wild-type siblings (black) are compared to the homozygous (red) except when otherwise stated (*kmt5b* only). P-value for *deaf1*-c207y day2nightall = 0.007 (left), 0.02 (right) and p-value for *kmt5b* day2nightall = 0.01 (left), 0.004 (right). (**B**) Additional mutants with a weaker (still repeatable) increased "bout time" phenotype that were not included in panel A. (**C**) Location phenotype for *hdlbp-ab* mutants; p-value = 0.004 (left), 0.07 (right) or 0.005 with the linear-mixed model significance method instead of the Kruskal-Wallis ANOVA (see Thyme, *et al.*, Cell 2019). (**D**) Two Magnitude measures showing phenotypes for *tbl1xr1-ab* homozygous/heterozygous mutants (red) compared to wild-type/heterozygous siblings (black). (**E**) Increased bout time for the heterozygous/heterozygous siblings (black) versus homozygous/homozygous mutants (red) for the *kmt2c-ab* double mutant.

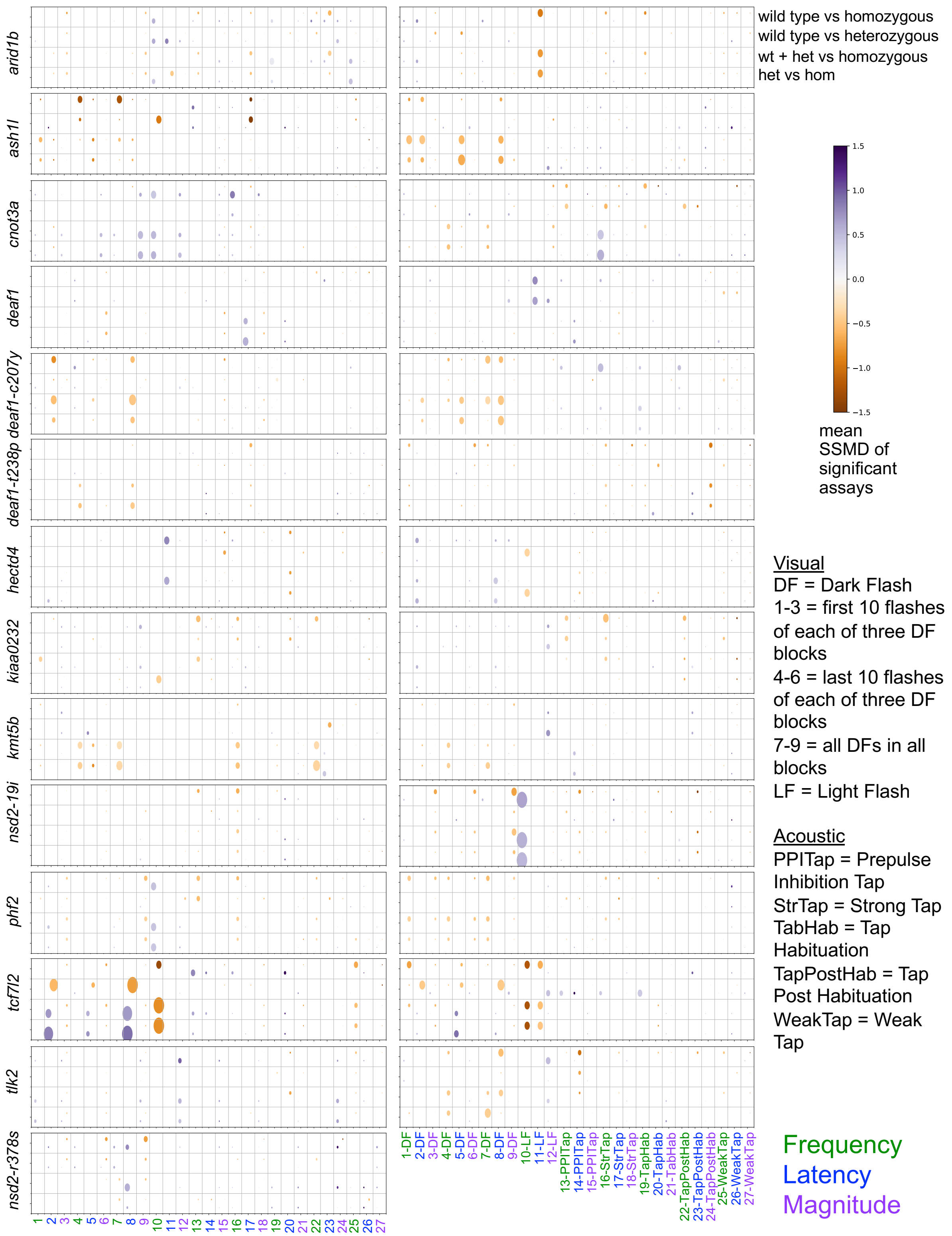


**Fig. S7. Stimulus-driven behavior summary data for single-gene mutants.**

Stimulus-driven behavior dot plots for all sibling comparisons from a heterozygous parental in-cross for single-gene mutants. Dot size corresponds to the percent of significant assays in the category (e.g., 27-Weak Tap Magnitude). Replicates are shown side-by-side, all N in Table S1.

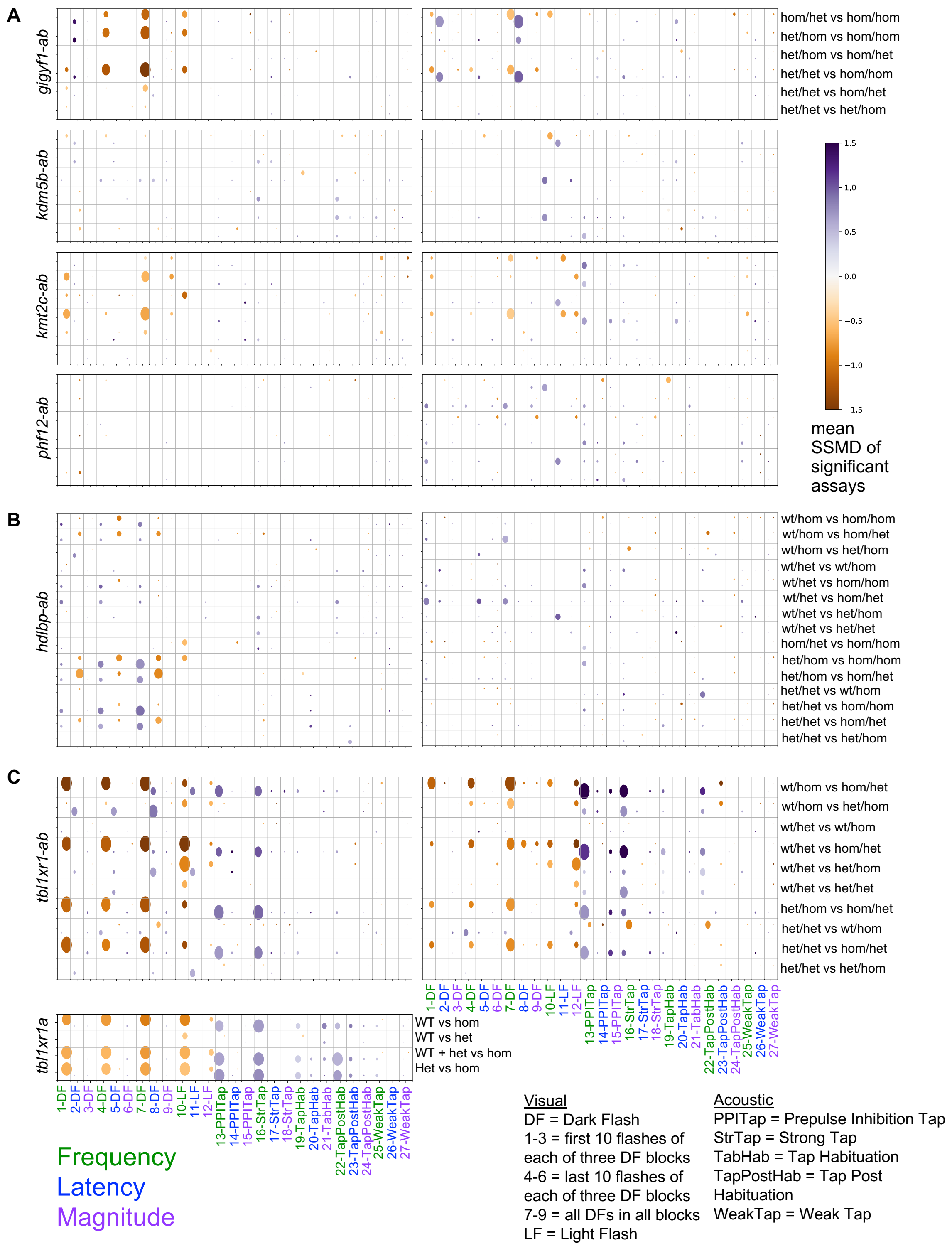


**Fig. S8. Stimulus-driven behavior summary data for double-gene mutants.**
Dot size is the percent of significant assays in the category. Replicates are shown side-by-side. (**A**) Parental cross: heterozygous/homozygous with homozygous/heterozygous. (**B**) Parental cross: heterozygous/heterozygous with heterozygous/homozygous. (**C**) Parental cross for *tbl1xr1-ab*: heterozygous/heterozygous with heterozygous/homozygous Parental cross for *tbl1xr1a*: standard single-mutant heterozygous with heterozygous. All N are in Table S1.

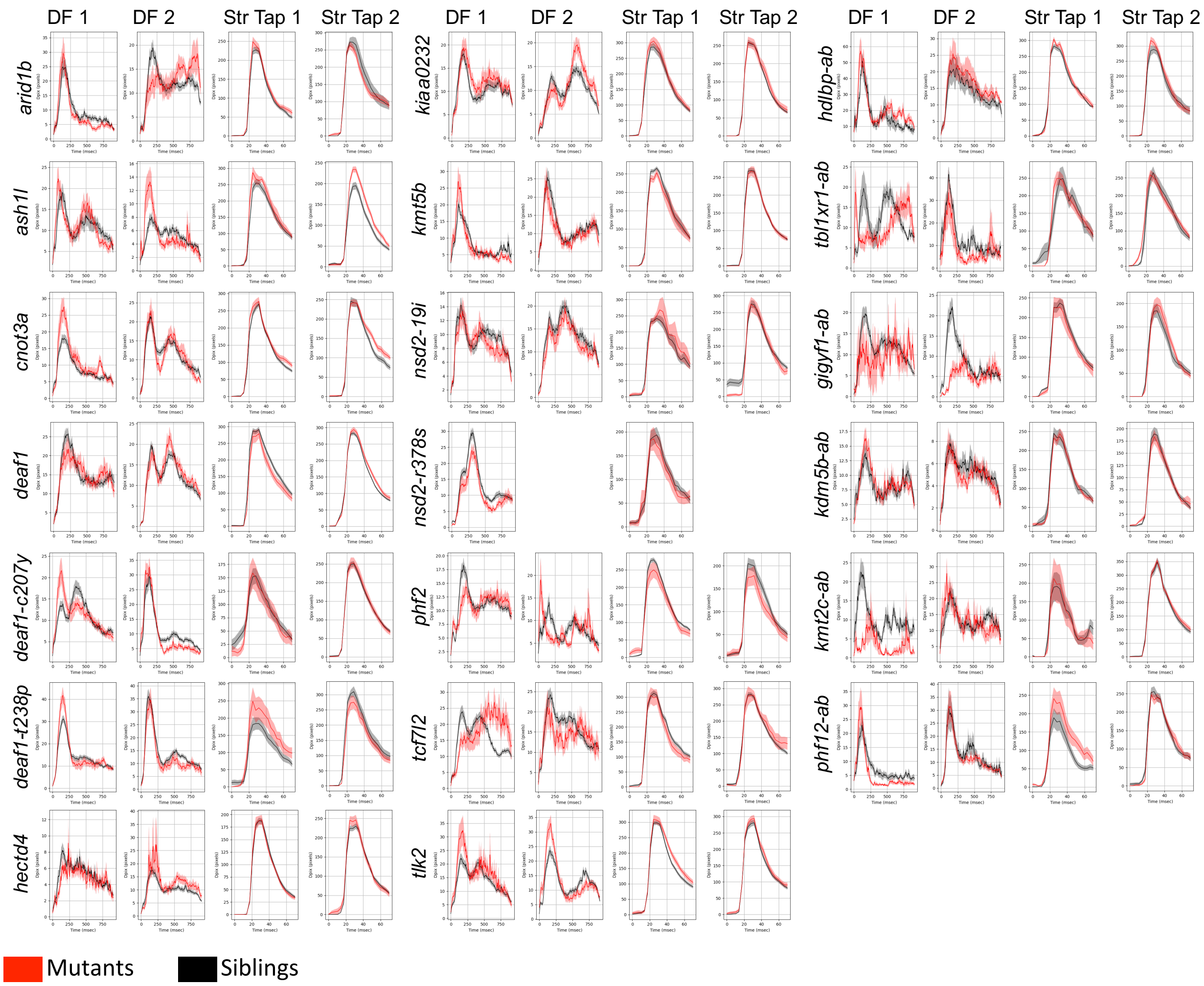


**Fig. S9. Example stimulus-driven behavioral response plot for all mutants.**

Examples (DF1) and replicates (DF2) for the dark flash responses (all dark flashes), as well as for a strong tap. The strong tap event selected was from the 5 dpf prepulse inhibition block, where there is no prepulse (a0f1000d5pD300a1f1000d5p). The response graphs an average of larvae in the mutant and control groups for events where a response was observed. All N are in Table S1.

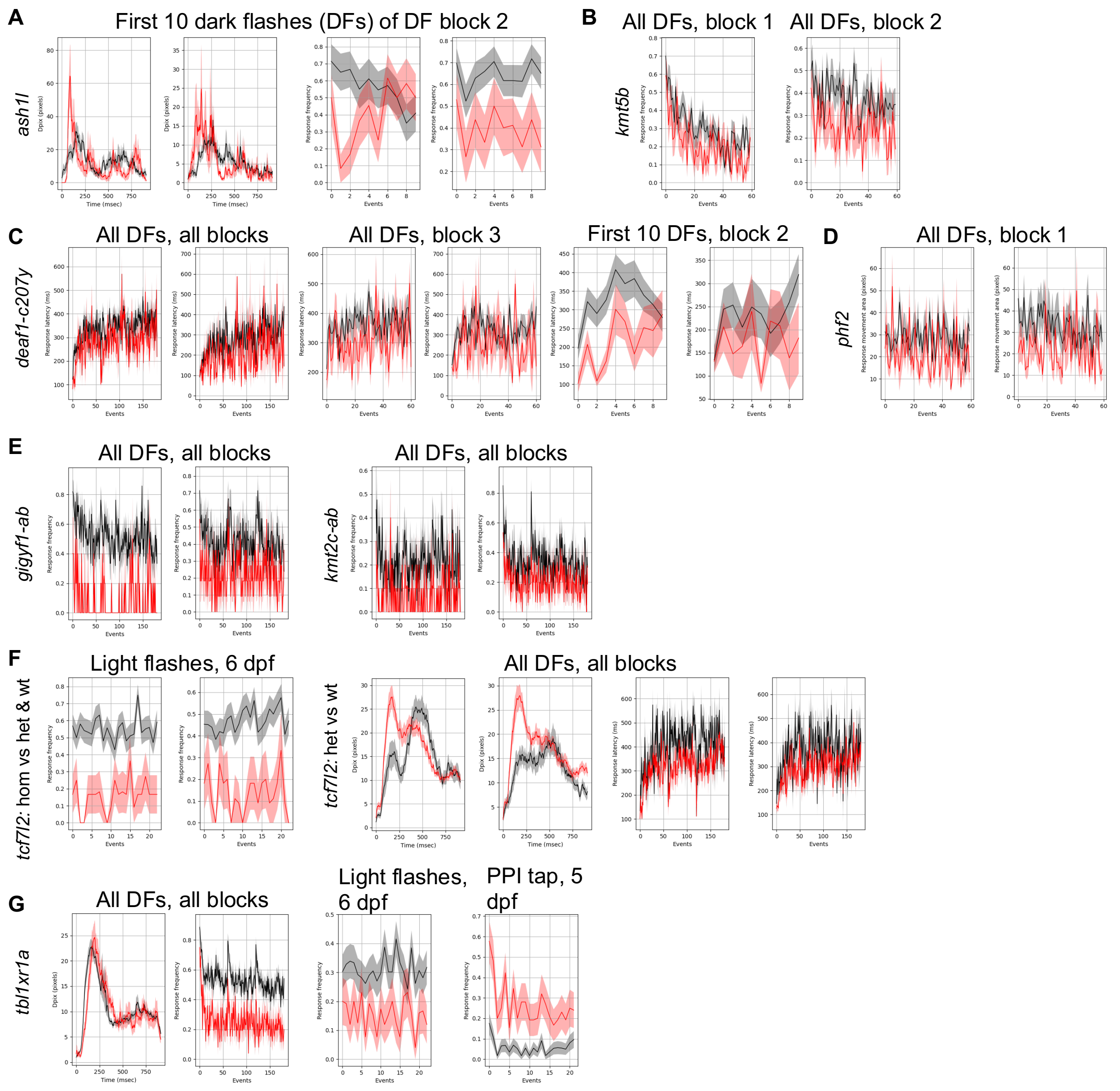


**Fig. S10. Stimulus-driven responses for repeatable findings.**

All N are in Table S1. P-values listed are from Kruskal-Wallis ANOVA. Biological replicates (different clutches) are shown side-by-side. (**A**) First 10 dark flashes of block 2 (day6dpfdf2a) response frequency p-value = 0.033, 0.0042. The corresponding response graphs are shown on the left (no p-value is calculated for them). (**B**) *kmt5b* frequency for all dark flashes block 1 p-value = 0.028 and block 2 for replicate = 0.024 (block 1 replicate = 0.1). *deaf1*-c207y response latency for all dark flashes = 0.0045, 0.007, block 3 = 0.022, 0.006, block 2, first 10 = 0.0008, 0.027. (**C**) *phf2* response area (day6dpfdf1) for all dark flashes of block 1 = 0.0006, 0.0013. (**D**) Response frequency for two double genes. *gigyf1-ab* response frequency for all dark flashes = 0.00094, 0.028. *kmt2c-ab* response frequency for all dark flashes = 0.0028, 0.025. (**F**) *tcf7l2* light flash (day6dpfmslf) response frequency = 0.00003, 0. <0.0001. *tcf7l2* het vs wt response latency for all dark flashes, all blocks = 0.0009, 0.037. Response graphs are shown on the left (no p-value). (**G**) *tbl1xr1a* dark flashes = <0.0001, light flashes = 0.002, and ppi tap = <0.0001.

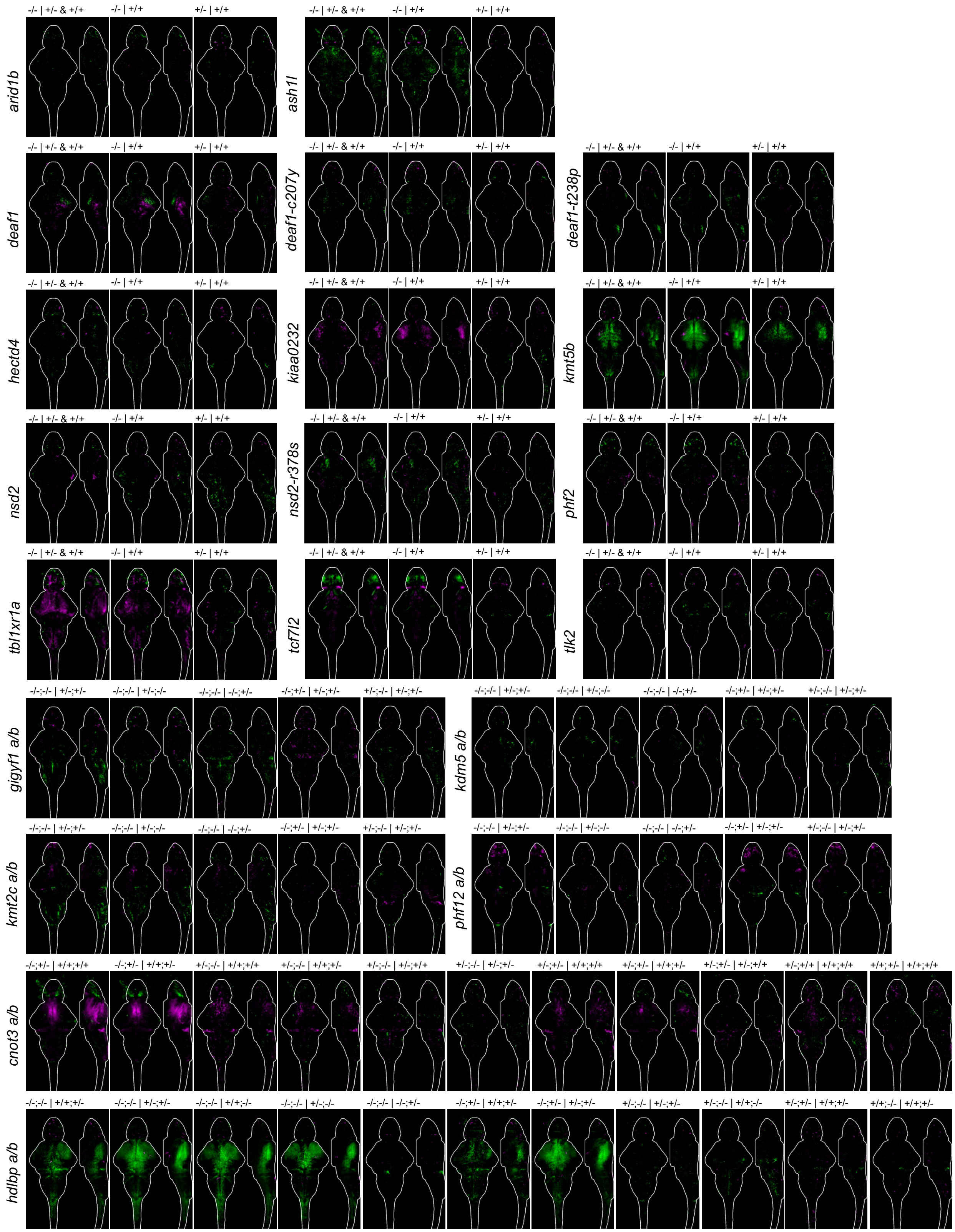


**Fig. S11. Brain activity maps for multiple genetic comparisons.**

Comparisons are shown above the sum-of-slices intensity projection with a 6 dpf brain outline, where the genotype before the | is being compared to the one after. All N are in Table S1.

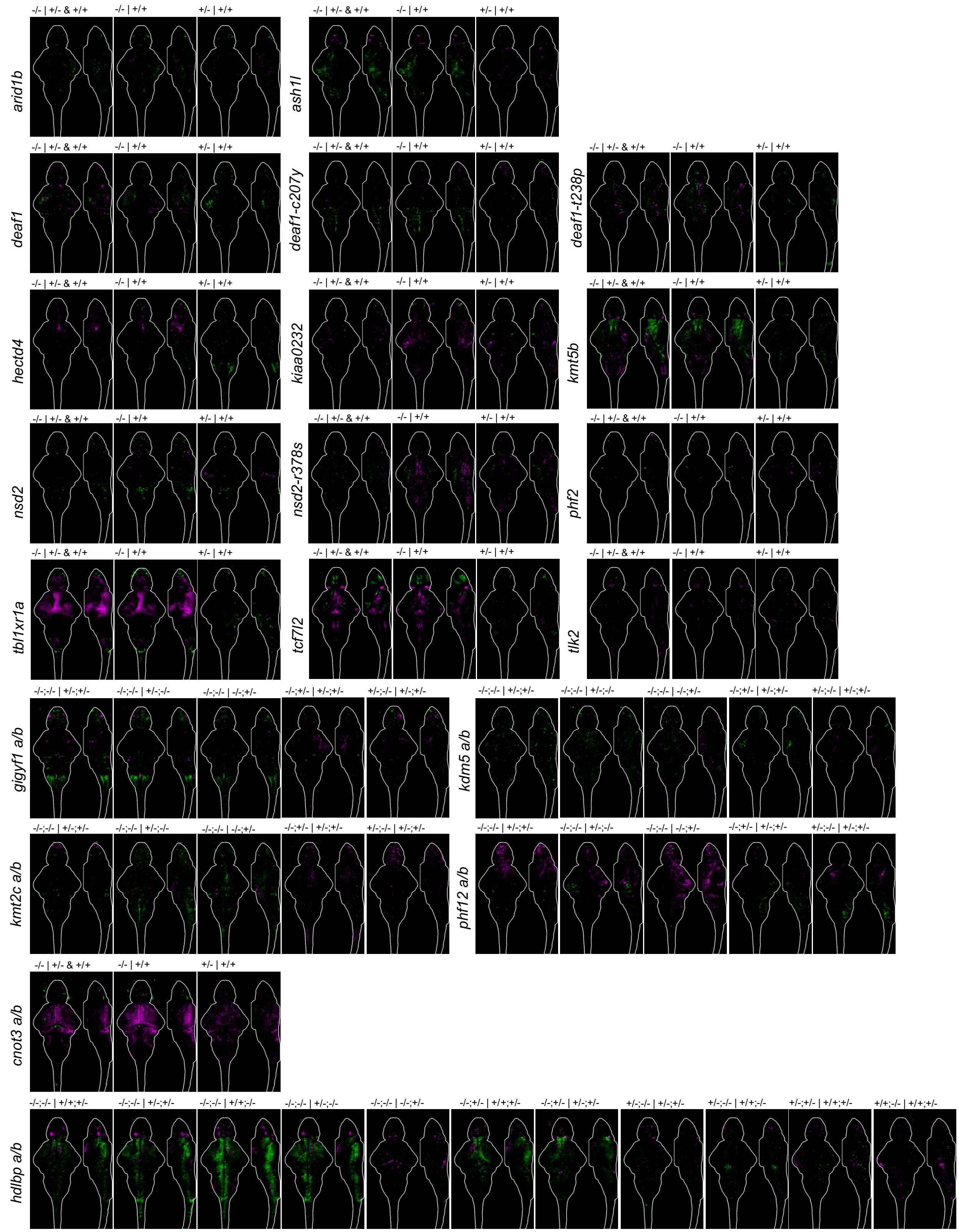


**Fig. S12. Second set of brain activity maps for multiple genetic comparisons.** Comparisons are shown above the sum-of-slices intensity projection with a 6 dpf brain outline, where the genotype before the | is being compared to the one after. All N are in Table S1.

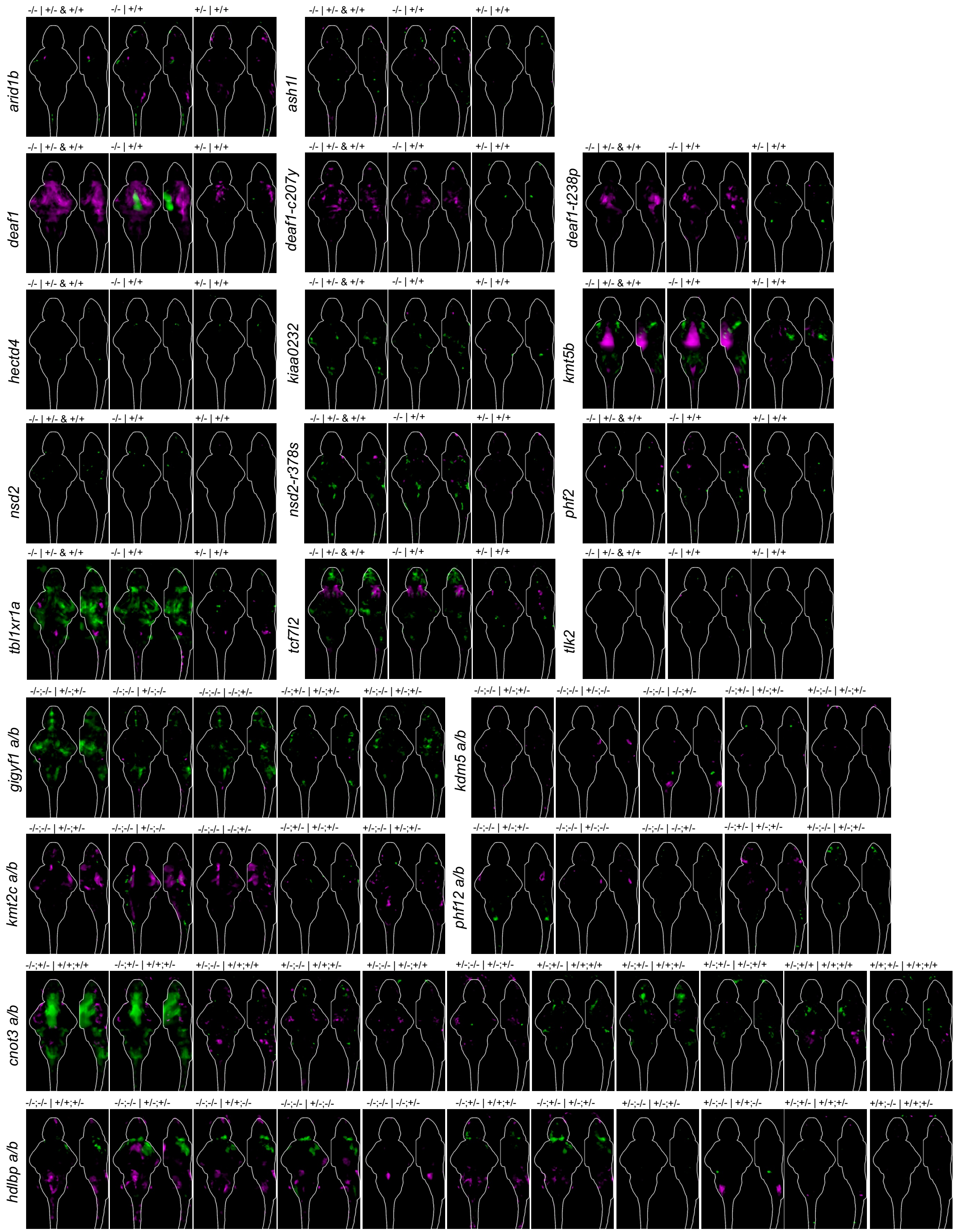


**Fig. S13. Brain structural maps for multiple genetic comparisons.**

Comparisons are shown above the sum-of-slices intensity projection with a 6 dpf brain outline, where the genotype before the | is being compared to the one after. All N are in Table S1.

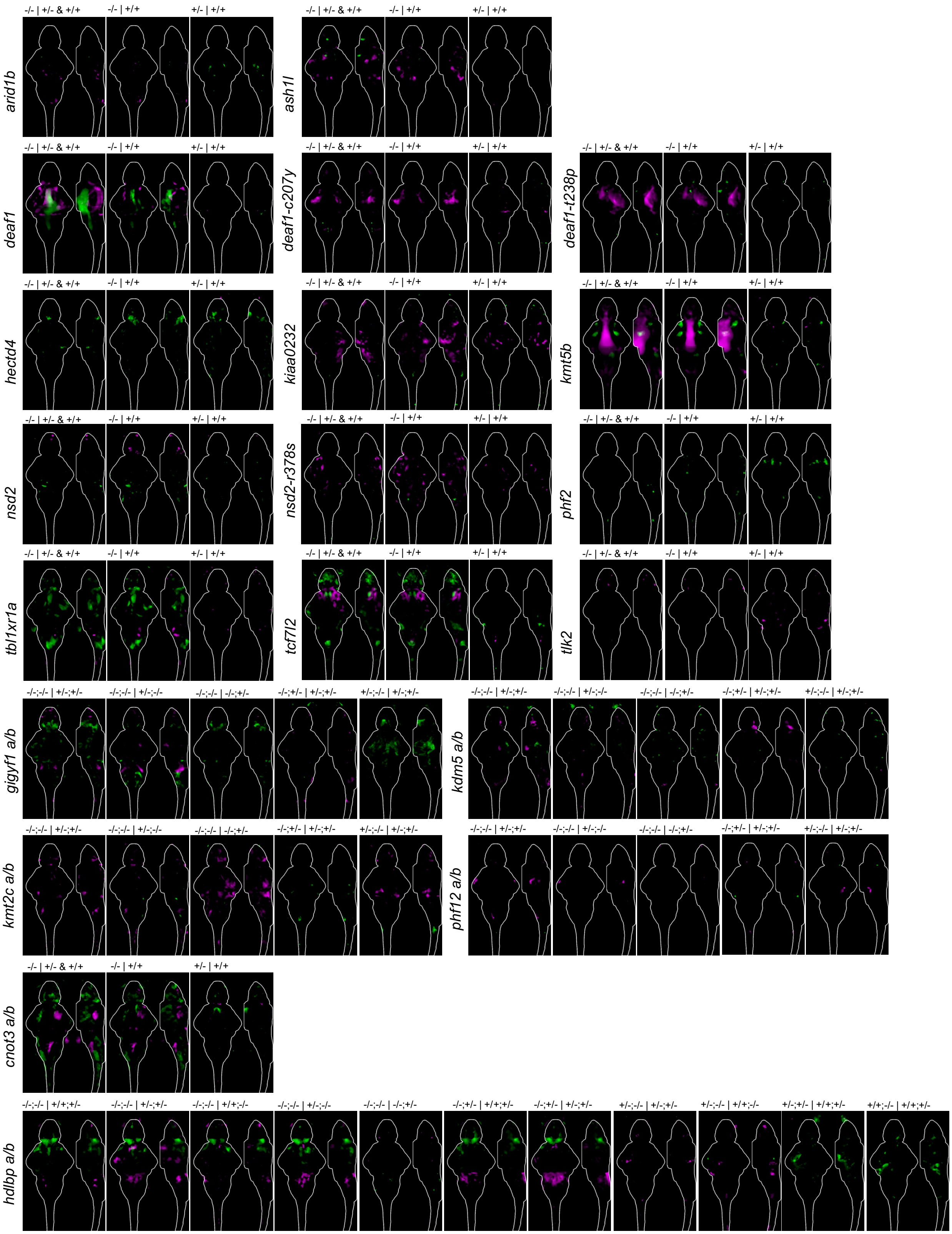


**Fig. S14. Second set of brain structural maps for multiple genetic comparisons.** Comparisons are shown above the sum-of-slices intensity projection with a 6 dpf brain outline, where the genotype before the | is being compared to the one after. All N are in Table S1.

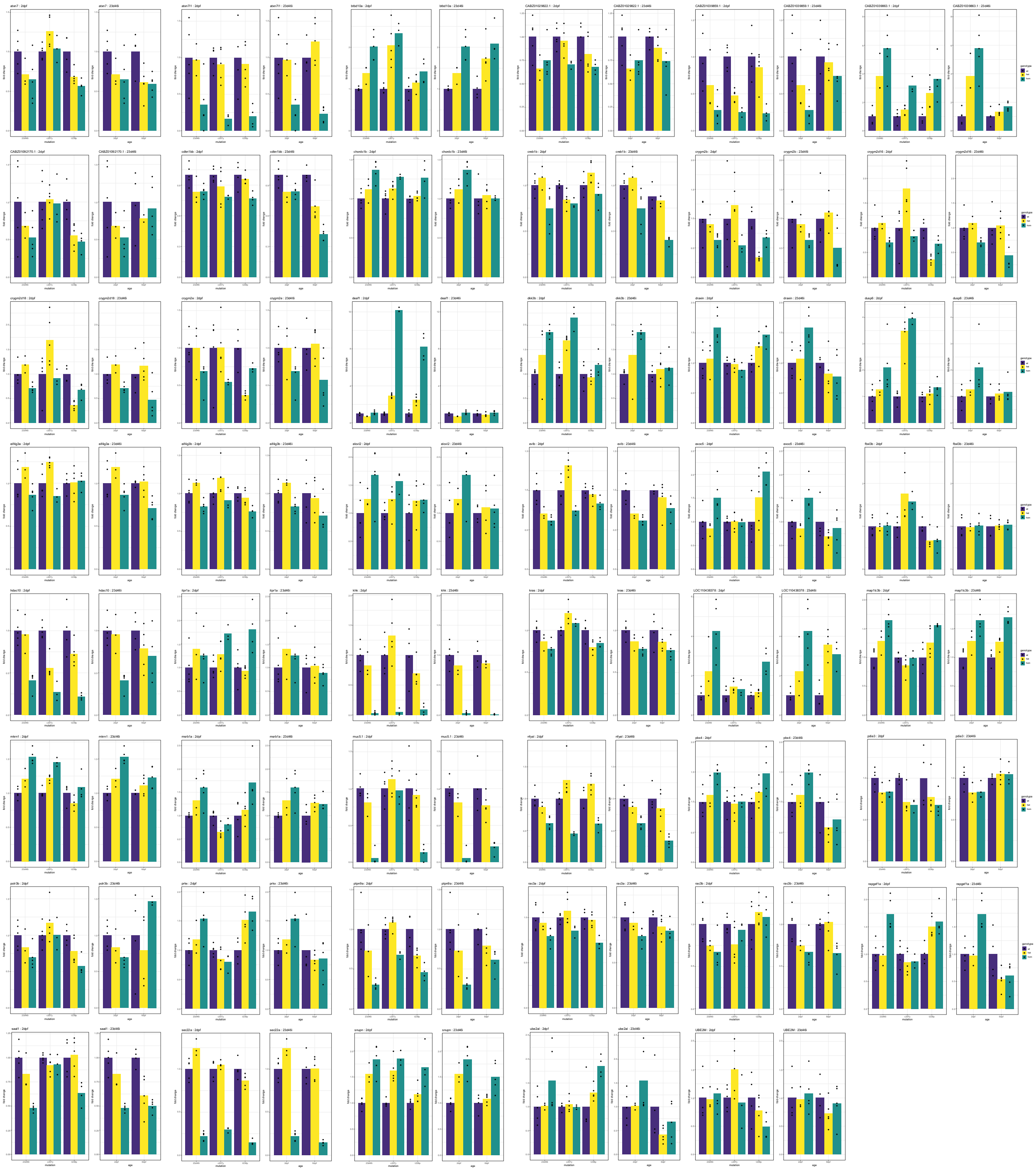

**Fig. S15. Individual gene expression plots for *deaf1* data.**

Individual bar plots showing differential expression of genes of interest between wild-type, heterozygous, and homozygous samples for all four datasets. The *deaf1*-23d46i 2 dpf data is shown both as a comparison with the other 2 dpf data and with the *deaf1*-23d46i 6 dpf data. P-values can be found in the RNA-seq analysis tables in Data S1.

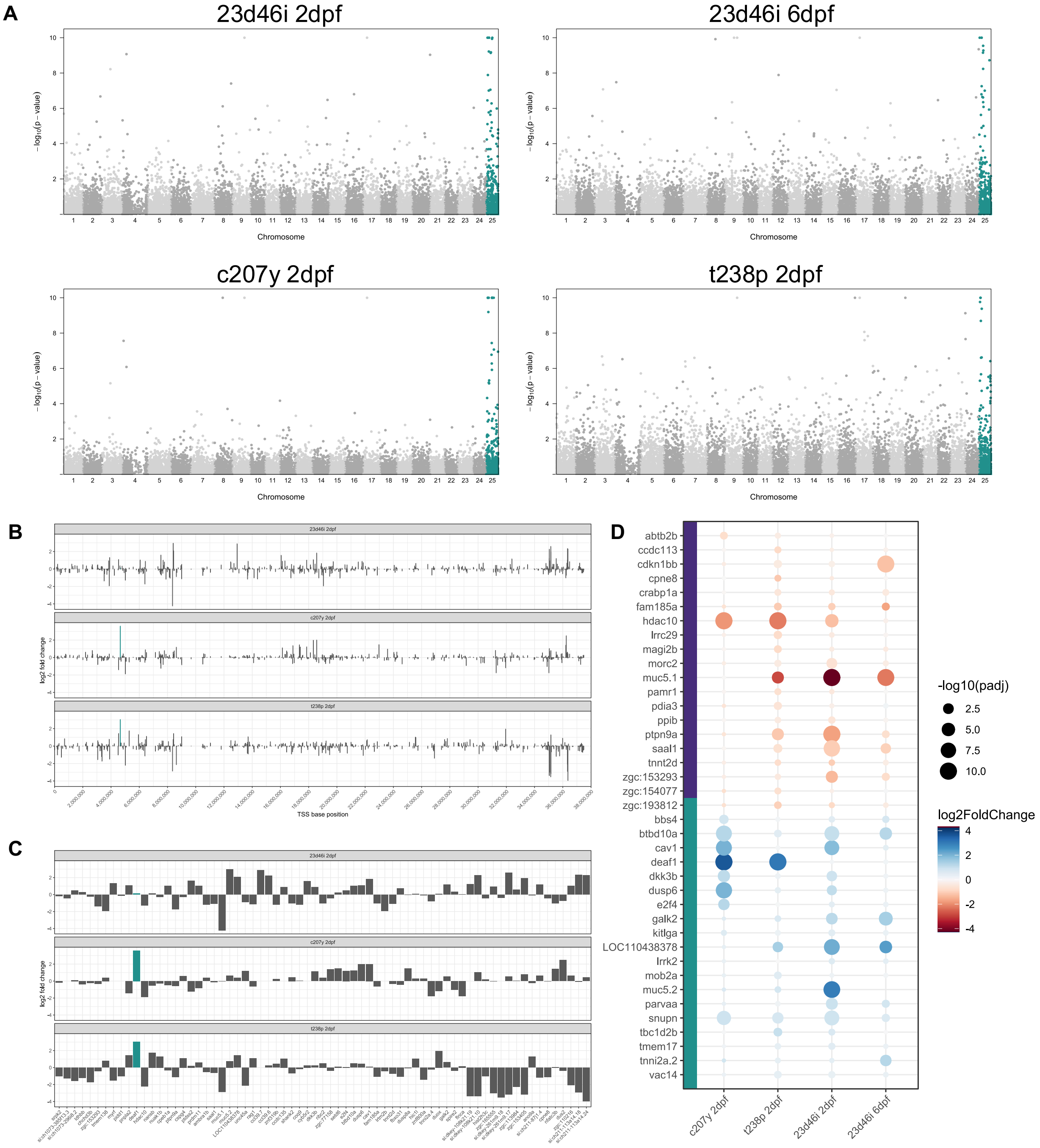


**Fig. S16. Enrichment of DEGs on the same chromosome as mutant alleles.**

(**A**) Manhattan plots for *deaf1* homozygous RNA-seq compared to the wild-type siblings, showing an enrichment for genes on chromosome 25 (chr25) in all four mutant lines. (**B**) Location of DEGs in the 2 dpf datasets. The *deaf1* location is marked with a teal bar. (**C**) Chr25 genes with a log2FoldChange > 1, in order of chromosome location. (**D**) Genes with consistent dysregulation (log2FoldChange > 0.3 or <-0.3 in at least two of the three 2 dpf datasets). These genes were included in subsequent analysis, while other chr25 genes were not.

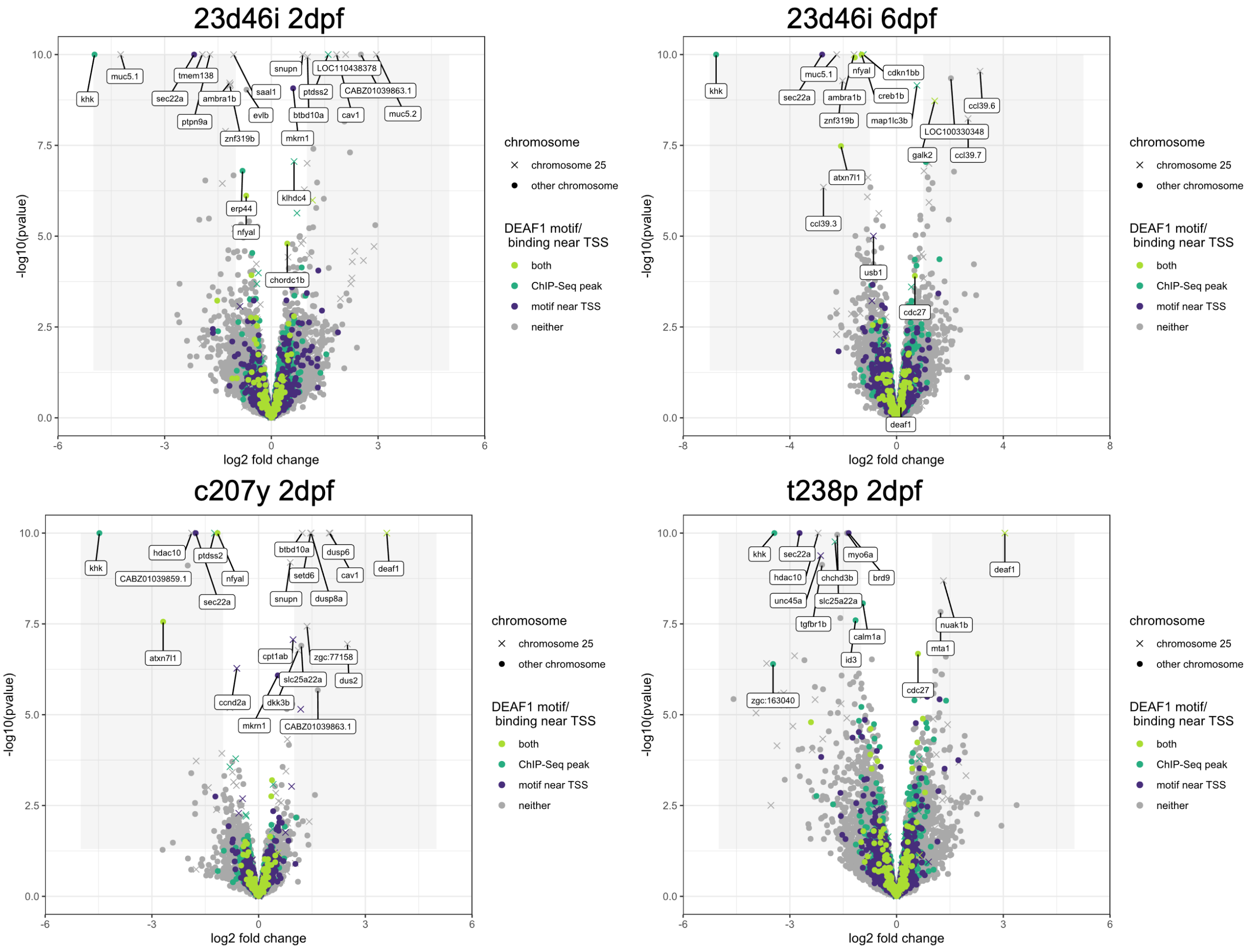


**Fig. S17. Differential gene expression in four *deaf1* mutant datasets.**

Volcano plots with labels for genes on the same chromosome (chr25) as *deaf1* and putative direct targets of Deaf1 that have ChIP-seq peaks or DEAF1 binding motifs. Genes are both upregulated and downregulated approximately equally in all datasets, and many differentially expressed genes are shared.

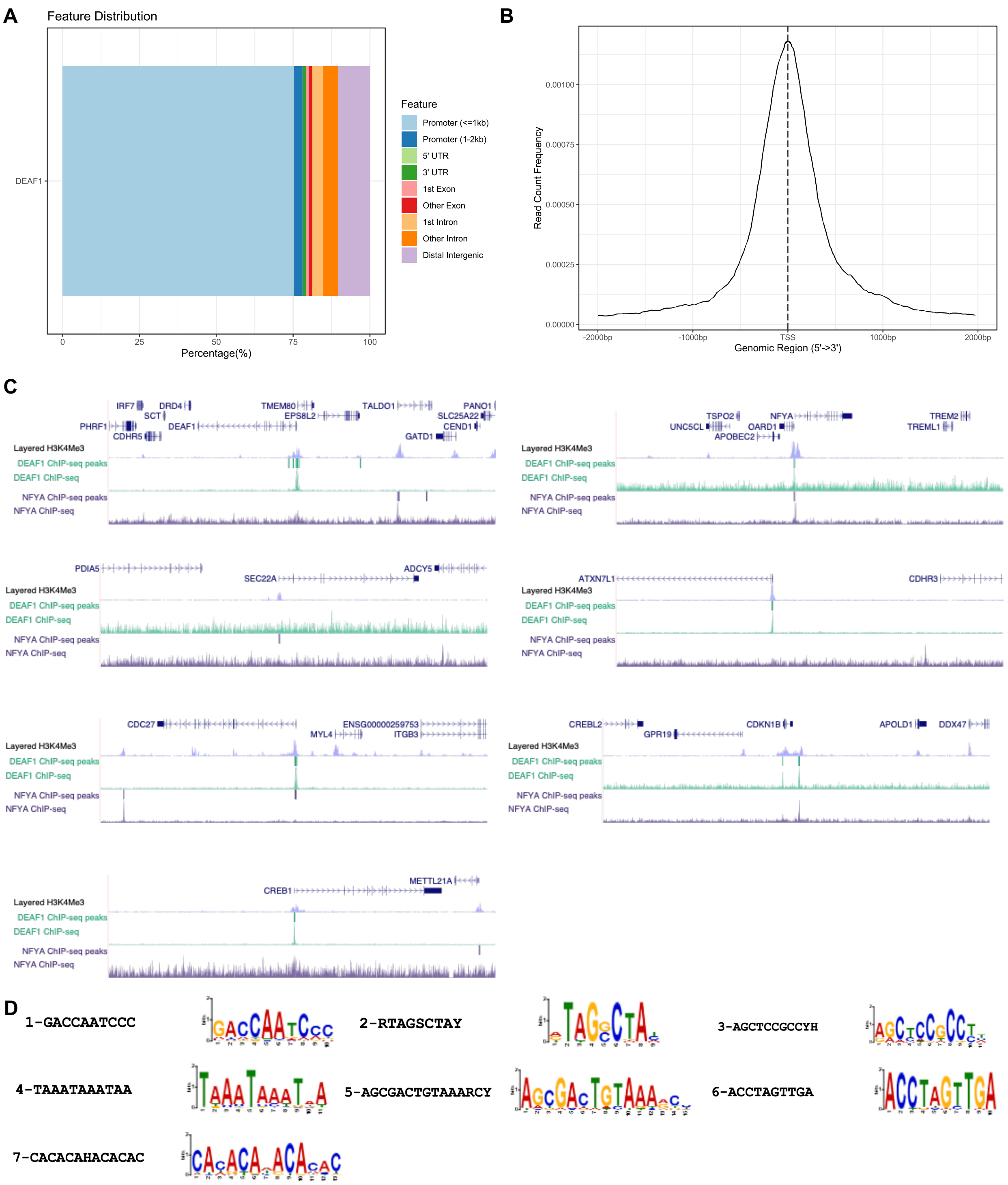


**Fig. S18. Transcription factor binding sites.**

(**A**) DEAF1 binding is enriched at promoters in ChIP-seq data from ENCODE (ENCFF813TRY). (**B**) DEAF1 binding is centered close to the transcription start site (TSS). (**C**) Orthologs of differentially expressed genes (DEGs) in the zebrafish *deaf1* mutant data have DEAF1 (ENCSR387SYS) and/or NFYA (ENCSR163VTS) ChIP-seq peaks at their TSSs. (**D**) Motifs discovered with STREME in the promoter regions of zebrafish DEGs.

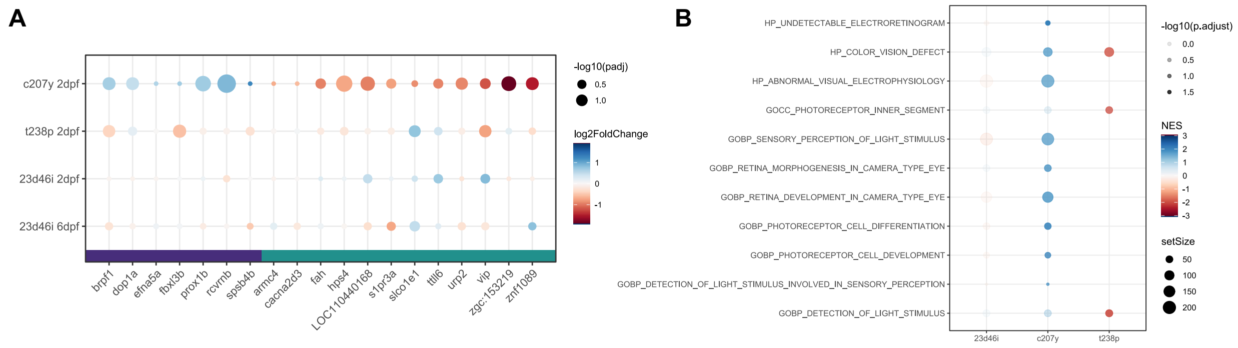


**Fig. S19. Divergent expression results in *deaf1*-c207y mutants.**

(**A**) Genes that are differentially expressed, based on a p-value of 0.01, in the *deaf1*-c207y samples, but not in the other *deaf1* mutant data. Selected genes of interest are shown from 49 identified. (**B**) C5 ontology terms identified with GSEA that are uniquely impacted in the *deaf1*-c207y data and related to the visual system.

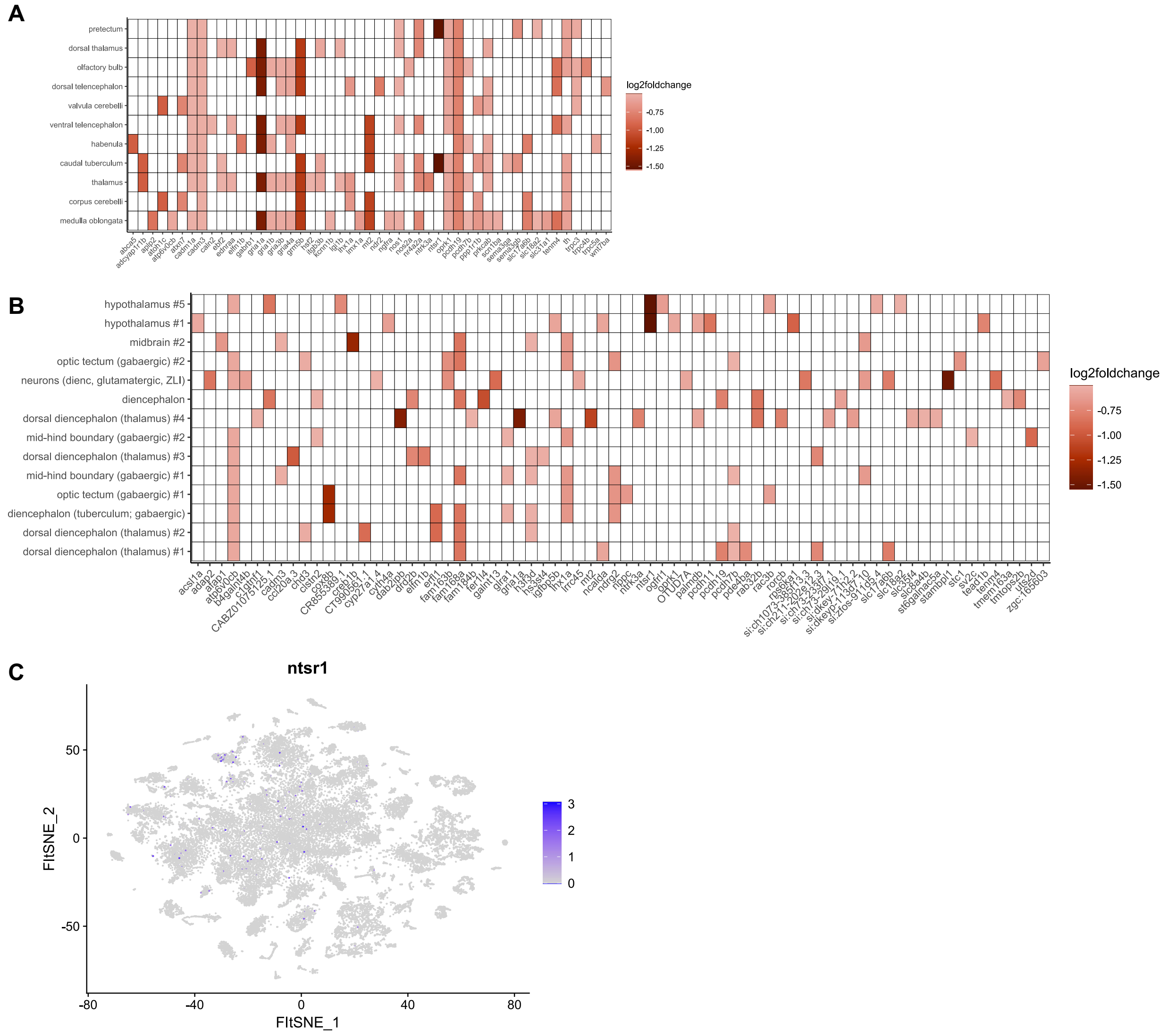


**Fig. S20. Genes contributing to GSEA in *deaf1*-23d46i 6 dpf mutants.**

(**A**) Genes contributing to ZFA terms corresponding to anatomical regions with differential gene expression. (**B**) Genes contributing to cell types from scRNA-seq. (**C**) Expression of *ntsr1*, marking cluster 34, in the scRNA-seq data from the 5 dpf head samples in Raj et al., 2020 (*29*).

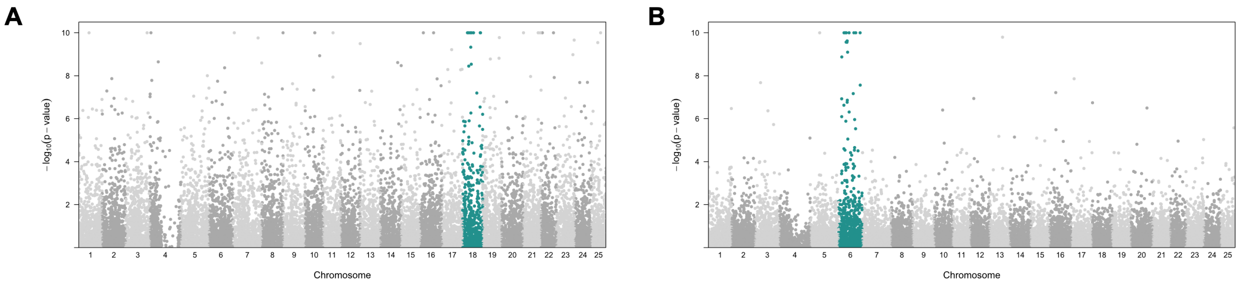


**Fig. S21. Manhattan plots for *kmt5b* and *hdlbpa* gene expression data.**

Both datasets show an enrichment of genes on the same chromosome as the mutant allele (chr18 for *kmt5b*, chr6 for *hdlbpa*). Differentially expressed genes on the same chromosome were not included in GSEA.

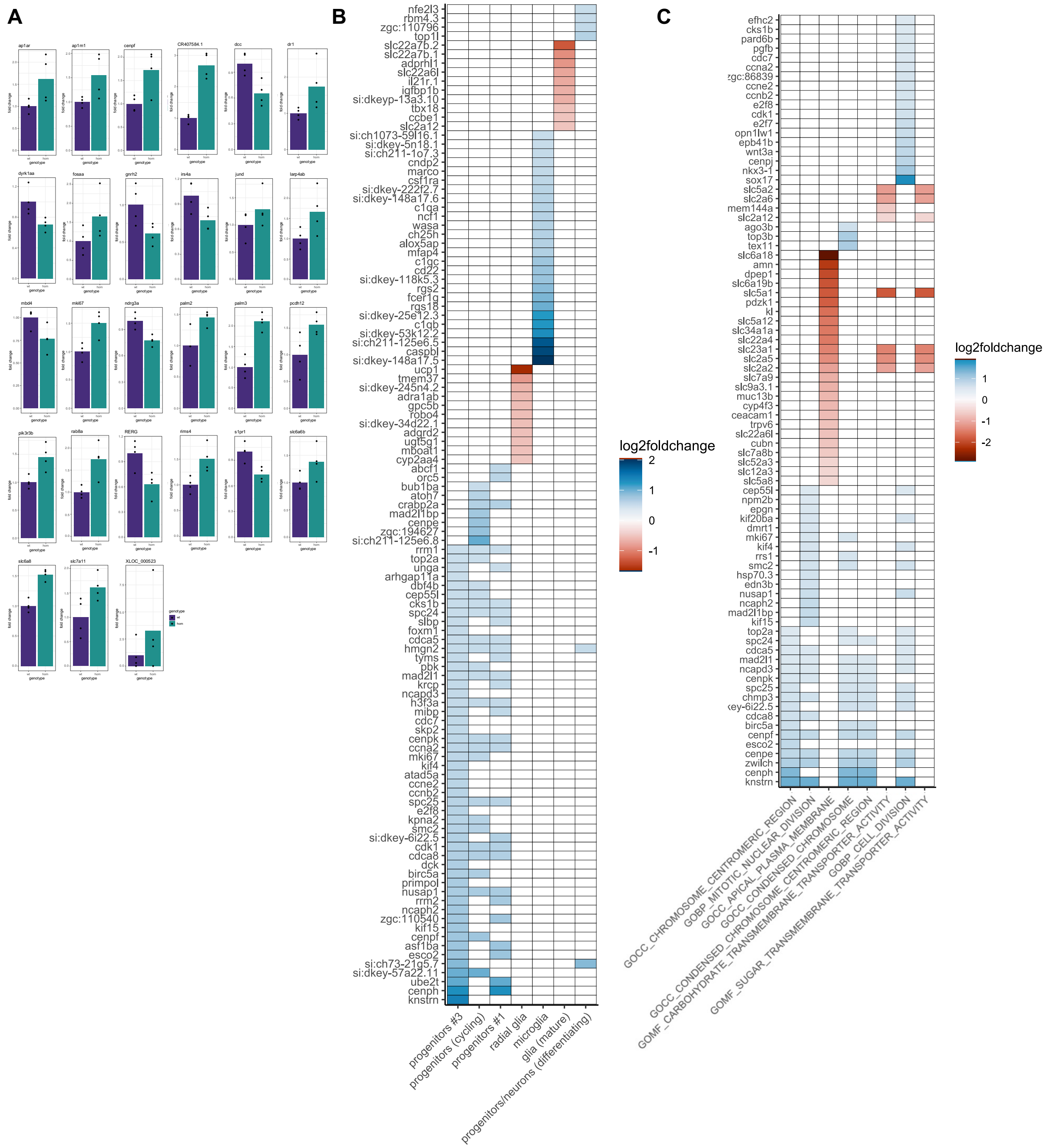


**Fig. S22. Expression analysis of *hdlbpa* mutants.**

(**A**) Individual bar plots showing differential expression of genes of interest between wild-type and homozygous samples. (**B**) Genes contributing to cell types from scRNA-seq identified with GSEA. (**C**) Genes contributing to C5 ontology terms identified with GSEA.

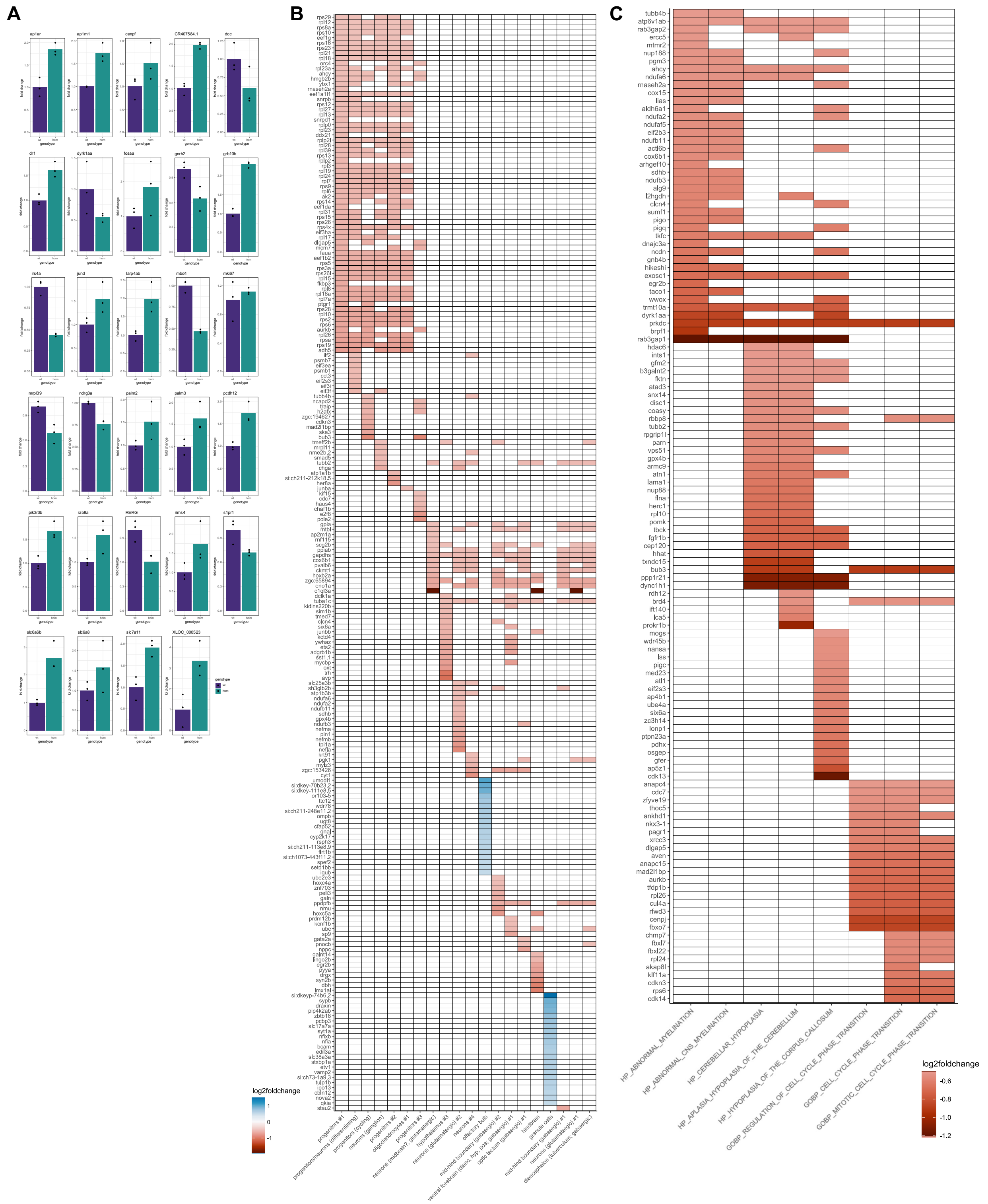


**Fig. S23. Expression analysis of *kmt5b* mutants.**

(**A**) Individual bar plots showing differential expression of genes of interest between wild-type and homozygous samples. (**B**) Genes contributing to cell types from scRNA-seq identified with GSEA. (**C**) Genes contributing to C5 ontology terms identified with GSEA.

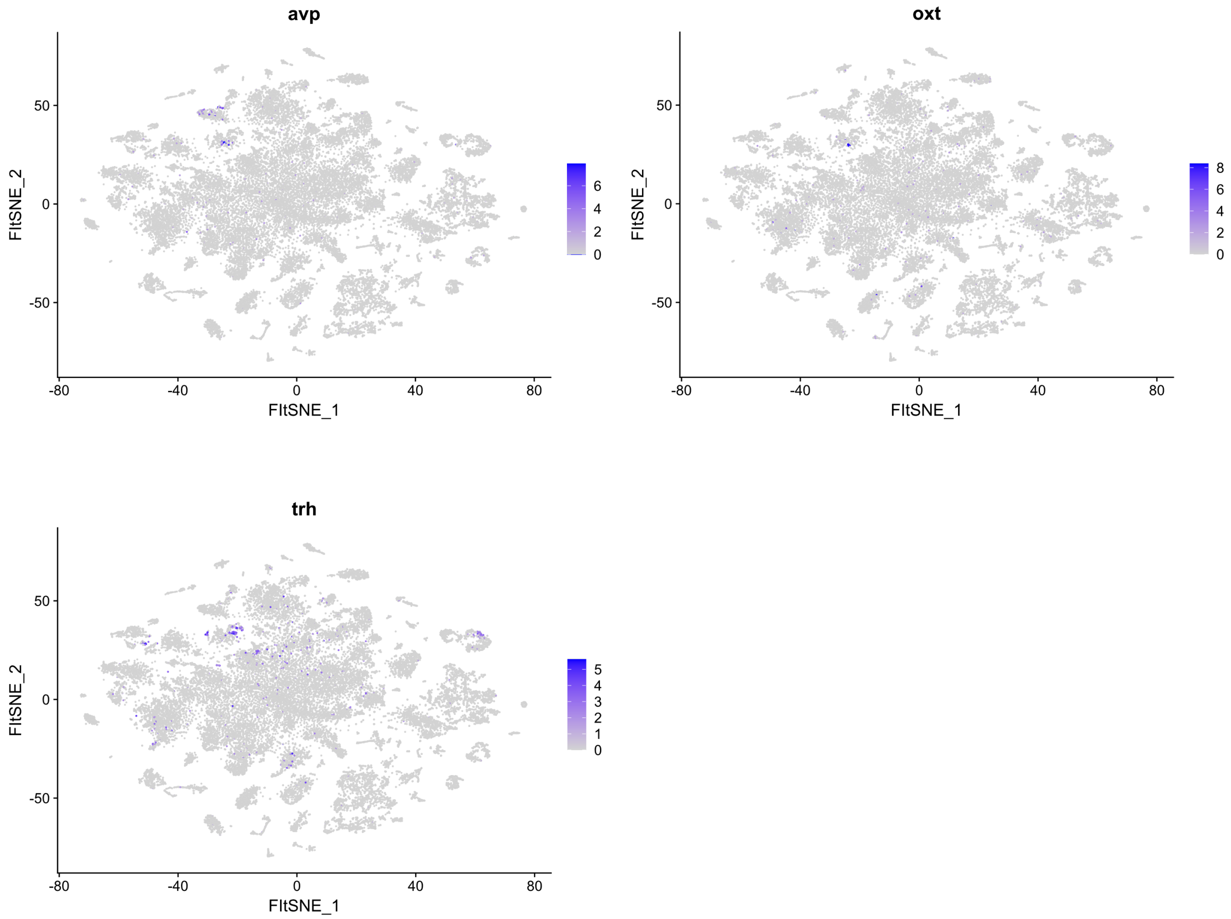


**Fig. S24. Expression of neuroendocrine cell markers in 5 dpf scRNA-seq data.**
These genes are markers of cluster 37 in the scRNA-seq data from the 5 dpf head samples in Raj et al., 2020 (*29*). Although they mark the same cluster and are known to have shared developmental origins (*66*), they are not expressed by the same neurons.

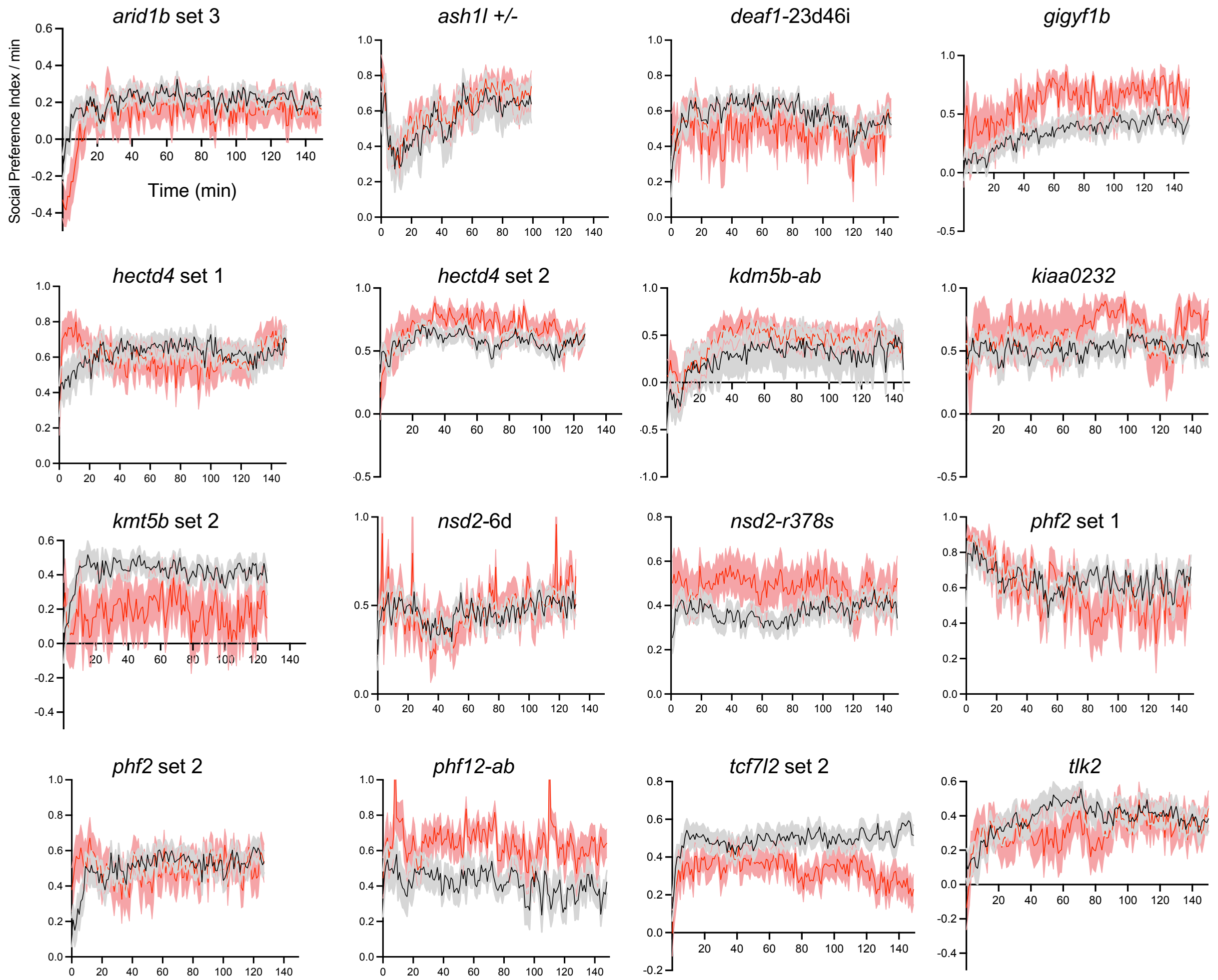


**Fig. S25. Additional social interaction data.**

Plots of mutant compared to control groups represent mean ± s.e.m. The N for all social behavior experiments is available in Table S1.

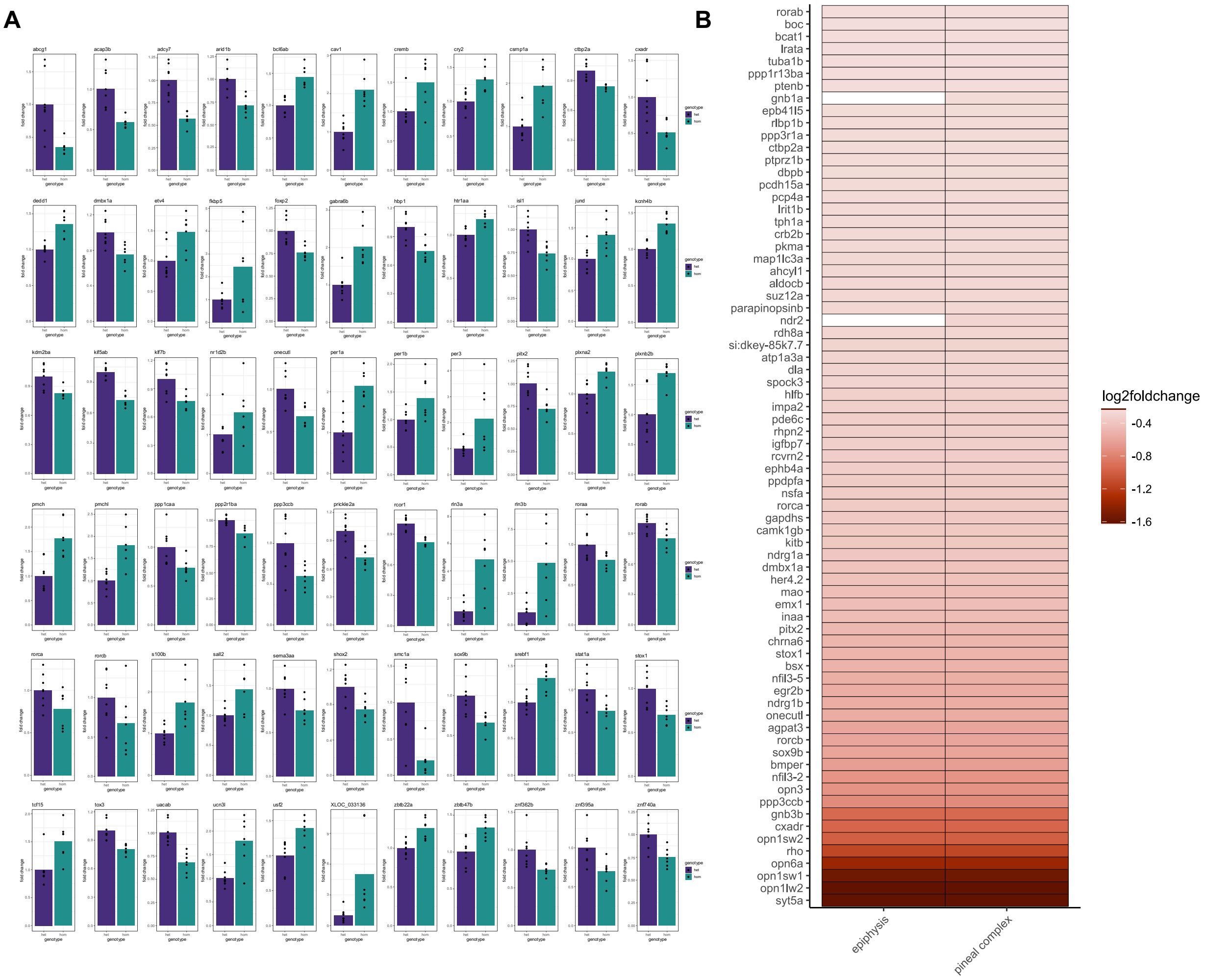


**Fig. S26. Expression analysis of *arid1b* mutants.**

(**A**) Individual bar plots showing differential expression of genes of interest between homozygous and heterozygous brains. (**B**) Genes contributing to ZFA terms identified with GSEA. Although it is possible that the pineal complex is impacted in these mutants, these genes are also shared with the eye and retina. There is far less adult expression data that contributes to the ZFA terms, making the outcome less reliable than for the 6 dpf findings.

**Table S1. Mutants generated and corresponding genotyping and experimental information.**

**Dataset S1. RNA-sequencing analysis files.**
