## Supplementary material for "Diencephalic and Neuropeptidergic Dysfunction in Zebrafish with Autism Risk Mutations": Data S1 - RNA-seq analysis files: arid1b_rnaseq.html

Arid1b Adult RNAseq Analysis


Code 

- Show All Code
- Hide All Code

### Arid1b Adult RNAseq Analysis

###### Anna Moyer

#### *2023-11-29*

Compare arid1b hom and het adult samples.

### 1 Setup

#### 1.1 Set seed and working directory

```
#set random seed
set.seed(123)

#set working directory
setwd("C:/Users/annam/Desktop/lab work/autismrnaseq/arid1b_updatedrnaseq")
```

#### 1.2 Load packages

```
library("tidyverse") 
library("DESeq2") #finding differentially expressed genes 
library("cowplot") #arranging plots into grids
library("viridis") #viridis color schemes
library("scales") #use to get color schemes for viridis
library("ggrepel") #repels text labels on plots
library("RColorBrewer") #pick colors
library("DT") #interactive and searchable tables of our GSEA results
library("GSEABase") #functions and methods for Gene Set Enrichment Analysis
library("Biobase") #base functions for bioconductor; required by GSEABase
library("GSVA") #Gene Set Variation Analysis, a non-parametric and unsupervised method for estimating variation of gene set enrichment across samples.
library("gprofiler2") #tools for accessing the GO enrichment results using g:Profiler web resources
library("clusterProfiler") # provides a suite of tools for functional enrichment analysis
library("msigdbr") # access to msigdb collections directly within R
library("enrichplot") # great for making the standard GSEA enrichment plots
library("ontologyIndex") #for parsing obo files
library("BaseSet") #for importing gaf file
library("plotly") #make interactive plots
library("orthogene") #convert gene names between species
library("ChIPseeker") #for annotating and browsing chip-seq data
library("TxDb.Hsapiens.UCSC.hg38.knownGene", pos = .Machine$integer.max) #human gene annotation for chip-seq data
library("EnsDb.Hsapiens.v75", pos = .Machine$integer.max) #human gene annotation for chip-seq data
library("org.Dr.eg.db") #convert ensembl gene names to gene id
library("lattice") #used for making manhattan plot
library("ggpubr") #calculating correlation coefficient 
library("colorspace") #colors for heatmap
library("ggraph") #library for making network graphs
library("svglite") #export svgs in loop
library("eulerr") #make venn diagrams
```

#### 1.3 Import files

```
#import  deseq output for arid1b data
arid1b <- read.csv("arid1badbrain-allhomvshetNOb1hom3COMBAT_allresults_wt_hom-with-normalized.csv")[,2:9]

#import  normalized counts for arid1b data
arid1b_genecounts <- read.csv("arid1badbrain-allhomvshetNOb1hom3COMBAT_normalized_reads_gene_list.csv")[,2:19]

#add some information 
#limit padj to 1e-10
arid1b <- arid1b %>% mutate(padjbound = ifelse(arid1b$padj < 1e-10, 1e-10, arid1b$padj))

#limit pvalue to 1e-10
arid1b <- arid1b %>% mutate(pvaluebound = ifelse(arid1b$pvalue < 1e-10, 1e-10, arid1b$pvalue))

#import annotations for chromosome and TSS
chromosomeinfo <- readr::read_tsv(file="ncbi_refseqgenes")

#keep chromosome and gene name
chromosomeinfo <- chromosomeinfo %>% dplyr::select(chrom, name2, txStart)

arid1b$chrom <- NA
arid1b$txStart <- NA

#add chromosome and txStart information
for (x in 1:length(arid1b$LLgeneAbbrev)) {
  gene <- arid1b$LLgeneAbbrev[x]
  chromosomeinfosub <- chromosomeinfo %>% dplyr::filter(name2 == gene)
  chr <- chromosomeinfosub$chrom[1]
  txStart <- chromosomeinfosub$txStart[1]
  arid1b$chrom[x] <- chr
  arid1b$txStart[x] <- txStart
}


#add a column with chromosome number
arid1b <- arid1b %>% mutate(chromosomenumber = str_replace_all(chrom, c("chr1"="1", "chr2"="2", "chr3"="3", "chr4"="4", "chr5"="5", "chr6"="6", "chr7"="7", "chr8"="8", "chr9"="9", "chr10"="10", "chr11"="11", "chr12"="12", "chr13"="13", "chr14"="14", "chr15"="15", "chr16"="16", "chr17"="17", "chr18"="18", "chr19"="19", "chr20"="20", "chr21"="21", "chr22"="22", "chr23"="23", "chr24"="24", "chr25"="25")))

arid1b <- distinct(arid1b)

#export csv
write.csv(arid1b, file = "arid1b_annotated.csv")
```

### 2 Plotting DEGs by chromosome

Are differentially expressed genes located at a particular place in
the genome?

#### 2.1 Make Manhattan plot for arid1b data

This function is from: https://genome.sph.umich.edu/wiki/Code\_Sample:\_Generating\_Manhattan\_Plots\_in\_R

```
#import  comparisons (to skip running previous steps)
arid1b <- read.csv("arid1b_annotated.csv")[,2:14]

#function for making manhattan plots
manhattan.plot<-function(chr, pos, pvalue, 
    sig.level=NA, annotate=NULL, ann.default=list(),
    should.thin=T, thin.pos.places=2, thin.logp.places=2, 
    xlab="Chromosome", ylab=expression(-log[10](p-value)),
    col=c("gray","darkgray"), panel.extra=NULL, pch=20, cex=0.8,...) {

    if (length(chr)==0) stop("chromosome vector is empty")
    if (length(pos)==0) stop("position vector is empty")
    if (length(pvalue)==0) stop("pvalue vector is empty")

    #make sure we have an ordered factor
    if(!is.ordered(chr)) {
        chr <- ordered(chr)
    } else {
        chr <- chr[,drop=T]
    }

    #make sure positions are in kbp
    if (any(pos>1e6)) pos<-pos/1e6;

    #calculate absolute genomic position
    #from relative chromosomal positions
    posmin <- tapply(pos,chr, min);
    posmax <- tapply(pos,chr, max);
    posshift <- head(c(0,cumsum(posmax)),-1);
    names(posshift) <- levels(chr)
    genpos <- pos + posshift[chr];
    getGenPos<-function(cchr, cpos) {
        p<-posshift[as.character(cchr)]+cpos
        return(p)
    }

    #parse annotations
    grp <- NULL
    ann.settings <- list()
    label.default<-list(x="peak",y="peak",adj=NULL, pos=3, offset=0.5, 
        col=NULL, fontface=NULL, fontsize=NULL, show=F)
    parse.label<-function(rawval, groupname) {
        r<-list(text=groupname)
        if(is.logical(rawval)) {
            if(!rawval) {r$show <- F}
        } else if (is.character(rawval) || is.expression(rawval)) {
            if(nchar(rawval)>=1) {
                r$text <- rawval
            }
        } else if (is.list(rawval)) {
            r <- modifyList(r, rawval)
        }
        return(r)
    }

    if(!is.null(annotate)) {
        if (is.list(annotate)) {
            grp <- annotate[[1]]
        } else {
            grp <- annotate
        } 
        if (!is.factor(grp)) {
            grp <- factor(grp)
        }
    } else {
        grp <- factor(rep(1, times=length(pvalue)))
    }
  
    ann.settings<-vector("list", length(levels(grp)))
    ann.settings[[1]]<-list(pch=pch, col=col, cex=cex, fill=col, label=label.default)

    if (length(ann.settings)>1) { 
        lcols<-trellis.par.get("superpose.symbol")$col 
        lfills<-trellis.par.get("superpose.symbol")$fill
        for(i in 2:length(levels(grp))) {
            ann.settings[[i]]<-list(pch=pch, 
                col=lcols[(i-2) %% length(lcols) +1 ], 
                fill=lfills[(i-2) %% length(lfills) +1 ], 
                cex=cex, label=label.default);
            ann.settings[[i]]$label$show <- T
        }
        names(ann.settings)<-levels(grp)
    }
    for(i in 1:length(ann.settings)) {
        if (i>1) {ann.settings[[i]] <- modifyList(ann.settings[[i]], ann.default)}
        ann.settings[[i]]$label <- modifyList(ann.settings[[i]]$label, 
            parse.label(ann.settings[[i]]$label, levels(grp)[i]))
    }
    if(is.list(annotate) && length(annotate)>1) {
        user.cols <- 2:length(annotate)
        ann.cols <- c()
        if(!is.null(names(annotate[-1])) && all(names(annotate[-1])!="")) {
            ann.cols<-match(names(annotate)[-1], names(ann.settings))
        } else {
            ann.cols<-user.cols-1
        }
        for(i in seq_along(user.cols)) {
            if(!is.null(annotate[[user.cols[i]]]$label)) {
                annotate[[user.cols[i]]]$label<-parse.label(annotate[[user.cols[i]]]$label, 
                    levels(grp)[ann.cols[i]])
            }
            ann.settings[[ann.cols[i]]]<-modifyList(ann.settings[[ann.cols[i]]], 
                annotate[[user.cols[i]]])
        }
    }
    rm(annotate)

    #reduce number of points plotted
    if(should.thin) {
        thinned <- unique(data.frame(
            logp=round(-log10(pvalue),thin.logp.places), 
            pos=round(genpos,thin.pos.places), 
            chr=chr,
            grp=grp)
        )
        logp <- thinned$logp
        genpos <- thinned$pos
        chr <- thinned$chr
        grp <- thinned$grp
        rm(thinned)
    } else {
        logp <- -log10(pvalue)
    }
    rm(pos, pvalue)
    gc()

    #custom axis to print chromosome names
    axis.chr <- function(side,...) {
        if(side=="bottom") {
            panel.axis(side=side, outside=T,
                at=((posmax+posmin)/2+posshift),
                labels=levels(chr), 
                ticks=F, rot=0,
                check.overlap=F
            )
        } else if (side=="top" || side=="right") {
            panel.axis(side=side, draw.labels=F, ticks=F);
        }
        else {
            axis.default(side=side,...);
        }
     }

    #make sure the y-lim covers the range (plus a bit more to look nice)
    prepanel.chr<-function(x,y,...) { 
        A<-list();
        maxy<-ceiling(max(y, ifelse(!is.na(sig.level), -log10(sig.level), 0)))+.5;
        A$ylim=c(0,maxy);
        A;
    }

    xyplot(logp~genpos, chr=chr, groups=grp,
        axis=axis.chr, ann.settings=ann.settings, 
        prepanel=prepanel.chr, scales=list(axs="i"),
        panel=function(x, y, ..., getgenpos) {
            if(!is.na(sig.level)) {
                #add significance line (if requested)
                panel.abline(h=-log10(sig.level), lty=2);
            }
            panel.superpose(x, y, ..., getgenpos=getgenpos);
            if(!is.null(panel.extra)) {
                panel.extra(x,y, getgenpos, ...)
            }
        },
        panel.groups = function(x,y,..., subscripts, group.number) {
            A<-list(...)
            #allow for different annotation settings
            gs <- ann.settings[[group.number]]
            A$col.symbol <- gs$col[(as.numeric(chr[subscripts])-1) %% length(gs$col) + 1]    
            A$cex <- gs$cex[(as.numeric(chr[subscripts])-1) %% length(gs$cex) + 1]
            A$pch <- gs$pch[(as.numeric(chr[subscripts])-1) %% length(gs$pch) + 1]
            A$fill <- gs$fill[(as.numeric(chr[subscripts])-1) %% length(gs$fill) + 1]
            A$x <- x
            A$y <- y
            do.call("panel.xyplot", A)
            #draw labels (if requested)
            if(gs$label$show) {
                gt<-gs$label
                names(gt)[which(names(gt)=="text")]<-"labels"
                gt$show<-NULL
                if(is.character(gt$x) | is.character(gt$y)) {
                    peak = which.max(y)
                    center = mean(range(x))
                    if (is.character(gt$x)) {
                        if(gt$x=="peak") {gt$x<-x[peak]}
                        if(gt$x=="center") {gt$x<-center}
                    }
                    if (is.character(gt$y)) {
                        if(gt$y=="peak") {gt$y<-y[peak]}
                    }
                }
                if(is.list(gt$x)) {
                    gt$x<-A$getgenpos(gt$x[[1]],gt$x[[2]])
                }
                do.call("panel.text", gt)
            }
        },
        xlab=xlab, ylab=ylab, 
        panel.extra=panel.extra, getgenpos=getGenPos, ...
    );
}

#select columns to include in manhattan plot
myTopHits.df <- arid1b %>% dplyr::select(chromosomenumber, txStart, pvaluebound)

#filter to only include chromosomes 1-25
myTopHits.df <- myTopHits.df %>% dplyr::filter(chromosomenumber %in% 1:25)

#make chromosomes plotted in order
myTopHits.df$chromosomenumber <- factor(myTopHits.df$chromosomenumber, levels = 1:25)

#omit NA values
myTopHits.df <- na.omit(myTopHits.df)

#make colors 
manhattancol <- c(replicate(9, c("lightgrey", "darkgrey")), "lightgrey", "#21908CFF", replicate(4, c("lightgrey", "darkgrey")))
 

manhattan <- myTopHits.df

#make manhattan plot
manhattan.plot(manhattan$chromosomenumber, manhattan$txStart, manhattan$pvaluebound, should.thin=F, col=manhattancol)
```

```
#export manhattan plots at 1000x500 pixels
```

#### 2.2 Make custom GSEA annotation with zebrafish genes and chromosomes

```
#import annotations for chromosome and gene name
chromosomeinfo <- readr::read_tsv(file="ncbi_refseqgenes")

#keep chromosome and gene name
chromosomeinfo <- chromosomeinfo %>% dplyr::select(chrom, name2)

#change column names
colnames(chromosomeinfo) <- c("chromosome", "genename")

#select only chromosomes 1-25
chromosomeinfo <- chromosomeinfo %>% dplyr::filter(chromosome %in% paste("chr", 1:25, sep=""))

#keep only distinct rows
chromosomeinfo <- distinct(chromosomeinfo)
```

#### 2.3 Check for enrichment of chromosome 20 genes in arid1b data

```
# Perform GSEA using clusterProfiler
#filter to only one comparison
GSEAgenes <- arid1b

#keep only the columns we need for GSEA
mydata.df.sub <- dplyr::select(GSEAgenes, LLgeneAbbrev, log2FoldChange)

#sort by abs(foldchange)
mydata.df.sub$log2FoldChange <- abs(mydata.df.sub$log2FoldChange)

# construct a named vector
mydata.gsea <- mydata.df.sub$log2FoldChange
names(mydata.gsea) <- as.character(mydata.df.sub$LLgeneAbbrev)
mydata.gsea <- sort(mydata.gsea, decreasing = TRUE)

# run GSEA using the 'GSEA' function from clusterProfiler
myGSEA.res <- GSEA(mydata.gsea, TERM2GENE=chromosomeinfo, verbose=FALSE, scoreType="pos", maxGSSize=2000, pvalueCutoff = 1, seed=TRUE)
myGSEA.df <- as_tibble(myGSEA.res@result)

# view results as an interactive table
datatable(myGSEA.df, 
          extensions = c('KeyTable', "FixedHeader"), 
          options = list(keys = TRUE, searchHighlight = TRUE, pageLength = 10, lengthMenu = c("10", "25", "50", "100"))) %>%
  formatRound(columns=c(2:10), digits=2)
```

No chromosome is enriched for differentially expressed genes in
arid1b mutants.

### 3 Individual analysis of arid1b

#### 3.1 Make bar plots

```
#import  normalized counts for arid1b data
arid1b_genecounts <- read.csv("arid1badbrain-allhomvshetNOb1hom3COMBAT_normalized_reads_gene_list.csv")[,2:19]

#pick out a genes to plot
geneofinterest <- unique(c("rln3a", "pmchl", "ucn3l", "pmch", "plxnb2b", "plxna2", "sema3aa", "per1a", "per3", "bcl6ab", "ctbp2a", "etv4", "hbp1", "jund", "klf7b", "rcor1", "sox9b", "tox3", "cremb", "csrnp1a", "dmbx1a", "foxp2", "kdm2ba", "onecutl", "pitx2", "shox2", "stat1a", "sall2", "srebf1", "stox1", "tcf15", "usf2", "zbtb22a", "zbtb47b", "znf362b", "znf395a", "znf740a", "ppp1caa", "ppp2r1ba", "ppp3ccb", "arid1b", "rln3a", "XLOC_033136", "cav1", "gabra6b", "pmch", "pmchl", "si:ch211-237l4.6", "kcnh4b", "dedd1", "kif5ab", "adcy7", "abcg1", "smc1a", "uacab", "acap3b", "per1a", "per3", "ucn3l", "cxadr", "s100b", "fkbp5", "ppp1caa", "ppp2r1ba", "ppp3ccb", "prickle2a", "foxp2", "fkbp5", "htr1aa", "isl1", "rln3b", "per1b", "cry2", "nr1d2b", "roraa", "rorab", "rorca", "rorcb"))


#make new columns with mean counts of het samples
genecountslogfc <- arid1b_genecounts
genecountslogfc$arid1bavg <- rowMeans(genecountslogfc[,c(4,6,8,10,12,13,14,15)])

#divide columns by mean counts of hets
genecountslogfc <- genecountslogfc %>% mutate(across(colnames(genecountslogfc)[4:18], function(x) x/arid1bavg))

#pull out data for a select gene
generepdata <- genecountslogfc %>% dplyr::filter(LLgeneAbbrev %in% geneofinterest)

#get rid of unnecessary columns
generepdata <- generepdata[,c(1,4:18)]

#pivot longer to make tidy
generepdata <- generepdata %>% pivot_longer(!LLgeneAbbrev, names_to = "sample", values_to = "foldchange")

#add condition information based on sampleName
generepdata$genotype <- generepdata$sample

#substitute condition name based on replicate
generepdata <- generepdata %>% mutate(genotype = str_replace_all(genotype, c("^het.*"="het", "^hom.*"="hom")))

#order data on plot
generepdata$genotype <- factor(generepdata$genotype, levels=c("het", "hom"))

#set colors
colors <- c("#472D7BFF", "#21908CFF") 

#make an empty list
plot_list = list()

#make all the plots
for (z in 1:length(geneofinterest)) {
  genedata2dpf <- generepdata %>% dplyr::filter(LLgeneAbbrev == geneofinterest[z])
  plot <- ggplot(genedata2dpf, aes(fill=genotype, y=foldchange, x=genotype)) + 
    geom_bar(position="dodge", stat="summary") +
    geom_point(position = position_dodge(width = .9)) + 
    labs(title = paste(geneofinterest[z])) +
    ylab("fold change") +
    theme_bw() +
    scale_fill_manual(values = colors)
  plot_list[[z]] <- plot
}

# Save plots to svg Makes a separate file for each plot.
for (i in 1:length(geneofinterest)) {
    file_name = paste("arid1b_foldchange_", geneofinterest[i], ".svg", sep="")
    svglite(file_name)
    print(plot_list[[i]])
    dev.off()
}
```

#### 3.2 Make volcano plot

```
#subset data to arid1b
myTopHits.df <- arid1b

#edit padj bound to e-20
myTopHits.df <- myTopHits.df %>% dplyr::mutate(padjbound = ifelse(padj < 1e-20, 1e-20, padj))

#add column about whether chr is 20
myTopHits.df <- myTopHits.df %>% dplyr::mutate(chrom = replace_na(chrom, "none"))
myTopHits.df <- myTopHits.df %>% mutate(chromosome = ifelse(chrom == "chr20", "chromosome 20", "other chromosome"))

#list of hormones
myTopHits.hormones <- myTopHits.df %>% dplyr::filter(LLgeneAbbrev %in% c("rln3a", "pmchl", "ucn3l", "pmch"))
myTopHits.hormones$category <- "hormones"

myTopHits.semaphorin <- myTopHits.df %>% dplyr::filter(LLgeneAbbrev %in% c("plxnb2b", "plxna2", "sema3aa"))
myTopHits.semaphorin$category <- "semaphorin pathway"

myTopHits.sleep <- myTopHits.df %>% dplyr::filter(LLgeneAbbrev %in% c("per1a", "per3", "per1b", "cry2", "nr1d2b", "roraa", "rorab", "rorca", "rorcb"))
myTopHits.sleep$category <- "sleep"

myTopHits.transcription <- myTopHits.df %>% dplyr::filter(LLgeneAbbrev %in% c("bcl6ab", "ctbp2a", "etv4", "hbp1", "jund", "klf7b", "rcor1", "sox9b", "tox3", "cremb", "csrnp1a", "dmbx1a", "foxp2", "kdm2ba", "onecutl", "pitx2", "shox2", "stat1a", "sall2", "srebf1", "stox1", "tcf15", "usf2", "zbtb22a", "zbtb47b", "znf362b", "znf395a", "znf740a"))
myTopHits.transcription$category <- "regulation of transcription"

myTopHits.phosphatase <- myTopHits.df %>% dplyr::filter(LLgeneAbbrev %in% c("ppp1caa", "ppp2r1ba", "ppp3ccb"))
myTopHits.phosphatase$category <- "protein phosphatase"

#genes to label
myTopHits.labels <- rbind(myTopHits.hormones, myTopHits.semaphorin, myTopHits.sleep, myTopHits.transcription, myTopHits.phosphatase)

#change order that points are plotted
myTopHits.labels <- myTopHits.labels %>% arrange(match(category,  c("regulation of transcription", "sleep",  "hormones", "semaphorin pathway", "protein phosphatase")), desc(category))

genestolabel <- c("arid1b", "rln3a", "XLOC_033136", "cav1", "gabra6b", "pmch", "pmchl", "si:ch211-237l4.6", "kcnh4b", "dedd1", "kif5ab", "adcy7", "abcg1", "smc1a", "uacab", "acap3b", "per1a", "per3", "ucn3l", "cxadr", "s100b", "fkbp5", "ppp1caa", "ppp2r1ba", "ppp3ccb", "prickle2a", "foxp2", "fkbp5")

#subset to only include labeled genes
myTopHits.labels.all <- myTopHits.df %>% dplyr::filter(LLgeneAbbrev %in% genestolabel)

#make all points other
myTopHits.df <- myTopHits.df %>% dplyr::mutate(mutation = "other")

#make the plot
arid1b_volcano <- ggplot() +
  annotate("rect", xmin = 1, xmax = 2.5, ymin = -log10(0.05), ymax = 6.5, 
           alpha=.15,   fill="grey") +
  annotate("rect", xmin = -1, xmax = -2.5, ymin = -log10(0.05), ymax = 6.5, 
           alpha=.15, fill="grey") +
  geom_point(data=myTopHits.df, aes(y=-log10(padjbound), x=log2FoldChange, shape = chromosome, color = mutation,  text = paste("Symbol:", LLgeneAbbrev)), size=2) +
  geom_point(data=myTopHits.labels, aes(y=-log10(padjbound), x = log2FoldChange, color=category, shape=chromosome),  size=2, show.legend = T) +
  theme_bw() +
  coord_cartesian(xlim = c(-2.6, 2.6), ylim = c(-0.5, 7), expand = FALSE) +
  ylab("-log10(padj)") + 
  xlab("log2 fold change") +
  geom_label_repel(data=myTopHits.labels.all, aes(x=log2FoldChange, y=-log10(padjbound), label=LLgeneAbbrev), force = 3, nudge_y = 1, size = 2.5, max.overlaps = Inf, show.legend = FALSE, color = "black") + #label selected genes
  theme_bw() +
  scale_color_manual(values = c("#AADC32FF",  "darkgrey", "#27AD81FF", "#472D7BFF","gold", "deepskyblue", "orange", "#D697FF"), name = "pathway") +
  scale_shape_manual(values=c(4, 16)) 

arid1b_volcano
```

```
#export 700x500
```

#### 3.3 DAVID pathway analysis

Get a list of the top 300 genes dysregulated in arid1b by padj to put
into DAVID.

```
#select arid1b genes that aren't on chromosome 20
arid1b_genesfordavid <- arid1b %>% dplyr::filter(chromosomenumber != 20)

#sort by pvalue
arid1b_genesfordavid <- arid1b_genesfordavid %>% dplyr::arrange(padj)

#get names of top 300 genes
arid1b_genesfordavid <- arid1b_genesfordavid$LLgeneAbbrev[1:300]

write.csv(arid1b_genesfordavid, "arid1b_output/arid1b_genesfordavid.csv")

# use the 'gost' function from the gprofiler2 package to run GO enrichment analysis
#genes shared between comparisons - both up and downregulated
gost.res.homwtall <- gost(arid1b_genesfordavid, organism = "drerio", correction_method = "fdr", significant = TRUE, evcodes = TRUE, ordered_query = TRUE)

#export to csv
write.csv(apply(gost.res.homwtall$result,2,as.character), file="arid1b_output/arid1b_GO-analysis.csv")

# produce a manhattan plot of enriched GO terms
gostplot(gost.res.homwtall, interactive = T, capped = T)
```

#### 3.4 arid1b GSEA with C5

```
# choose a specific msigdb collection/subcollection
dr_gsea_c5 <- msigdbr(species = "Danio rerio", category = "C5") %>% # choose  msigdb collection of interest
  dplyr::select(gs_name, gene_symbol) #just get the columns corresponding to signature name and gene symbols of genes in each signature 

# pull out data for arid1b
GSEAgenes <- arid1b %>% dplyr::filter(chromosomenumber != 20)
mydata.df.sub <- dplyr::select(GSEAgenes, LLgeneAbbrev, log2FoldChange)

#get rid of duplicates
mydata.df.sub <- mydata.df.sub %>% dplyr::filter(LLgeneAbbrev %in% mydata.df.sub$LLgeneAbbrev[!duplicated(mydata.df.sub$LLgeneAbbrev)])

# construct a named vector
mydata.gsea <- mydata.df.sub$log2FoldChange
names(mydata.gsea) <- as.character(mydata.df.sub$LLgeneAbbrev)
mydata.gsea <- sort(mydata.gsea, decreasing = TRUE)

# run GSEA with C5 in arid1b
myGSEA.res <- GSEA(mydata.gsea, TERM2GENE=dr_gsea_c5, verbose=FALSE, seed=TRUE)

#convert to DF
myGSEA.df <- as_tibble(myGSEA.res)

#look at top terms
datatable(myGSEA.df, 
          extensions = c('KeyTable', "FixedHeader"), 
          options = list(keys = TRUE, searchHighlight = TRUE, pageLength = 10, lengthMenu = c("10", "25", "50", "100"))) %>%
  formatRound(columns=c(2:10), digits=2)
```

```
#export
write.csv(myGSEA.df, "arid1b_output/arid1b-GSEA-c5.csv")
```

#### 3.5 Use ZFA anatomy to perform GSEA on arid1b

```
CNStermsGSEA <- read.csv("CNStermsGSEA.csv")[,2:3]
headtermsGSEA <- read.csv("headtermsGSEA.csv")[,2:3]
fullsetGSEA <- read.csv("fullsetGSEA.csv")[,2:3]

# Pull out just the columns corresponding to gene symbols and LogFC for at least one pairwise comparison for the enrichment analysis
GSEAgenes <- arid1b %>% dplyr::filter(chromosomenumber != 20)
mydata.df.sub <- dplyr::select(GSEAgenes, LLgeneAbbrev, log2FoldChange)

#get rid of duplicates
mydata.df.sub <- mydata.df.sub %>% dplyr::filter(LLgeneAbbrev %in% mydata.df.sub$LLgeneAbbrev[!duplicated(mydata.df.sub$LLgeneAbbrev)])

# construct a named vector
mydata.gsea <- mydata.df.sub$log2FoldChange
names(mydata.gsea) <- as.character(mydata.df.sub$LLgeneAbbrev)
mydata.gsea <- sort(mydata.gsea, decreasing = TRUE)

#options for doing GSEA using ZFA
#CNStermsGSEA
#headtermsGSEA
#fullsetGSEA

# run GSEA with CNS terms 
myGSEA.res <- GSEA(mydata.gsea, TERM2GENE=CNStermsGSEA, verbose=FALSE, seed=TRUE, minGSSize = 80)
myGSEA.df <- as_tibble(myGSEA.res@result)

#export file
write.csv(myGSEA.df, file="arid1b_output/arid1b_CNS_GSEA.csv")

# view results as an interactive table
datatable(myGSEA.df, 
          extensions = c('KeyTable', "FixedHeader"), 
          options = list(keys = TRUE, searchHighlight = TRUE, pageLength = 10, lengthMenu = c("10", "25", "50", "100"))) %>%
  formatRound(columns=c(2:10), digits=2)
```

```
#make network plot
myGSEA.res <- pairwise_termsim(myGSEA.res)
arid1bnetwork <- emapplot(myGSEA.res, color="NES", categorySize="p.adjust", showCategory = 2)

#get data of out network plot
arid1bnetwork <- ggplot_build(arid1bnetwork)
networkdata <- arid1bnetwork$plot$data

#get color
myheatcolors3 <- brewer.pal(name="RdBu", n=11)

#make network plot
arid1b_network <- ggraph(networkdata) + 
  geom_edge_link(alpha=.8, aes_(width=~I(width)), colour='darkgrey') + 
  geom_node_point(aes(colour = color, size=size)) +
  geom_node_text(aes(label=name), repel=TRUE) + 
  theme_void() +
  scale_color_gradientn(colors = myheatcolors3, limit=c(-1.5,1.5), name="NES")

arid1b_network
```

```
#export 700x500

#make heatmap
p3 <- heatplot(myGSEA.res, foldChange=mydata.gsea, showCategory = 2) + ggplot2::coord_flip()

#get data of out heatmap
p3 <- ggplot_build(p3)
heatmapdata <- p3$plot$data

#filter to only include fold changes > 0.5
genestoinclude <- heatmapdata %>% dplyr::filter(foldChange > 0.2 | foldChange < -0.2)
heatmapdata <- heatmapdata %>% dplyr::filter(Gene %in% genestoinclude$Gene)

#use pivot wider to make untidy table
heatmapdata <- heatmapdata %>% pivot_wider(names_from = categoryID, values_from  = foldChange, values_fill=NA) 

#make matrix
heatmapmatrix <- as.matrix(heatmapdata[,2:3])
rownames(heatmapmatrix) <- heatmapdata$Gene

#turn matrix into 1s and 0s and cluster
heatmapmatrix<-ifelse(abs(heatmapmatrix) > 0,1,0)
heatmapmatrix[is.na(heatmapmatrix)] = 0

 #cluster rows by pearson correlation
#hc <- hclust(as.dist(1-cor(heatmapmatrix, method="spearman")), method="average") #cluster columns by spearman correlation

#turn heatmap data back into tidy table
heatmapdata <- heatmapdata %>% pivot_longer(!Gene, names_to = "categoryID", values_to  = "foldChange") 
    
#change order of data to be plotted
heatmapdata$Gene <- factor(heatmapdata$Gene, levels = rev(unique(heatmapdata$Gene)))

#rename dataframe columns
colnames(heatmapdata) <- c("Gene", "categoryID", "log2foldchange")

singlecellheatmap <- ggplot(heatmapdata, aes(x = Gene, y = categoryID, fill = log2foldchange)) + 
  geom_tile(color="black") + 
  theme_classic() + 
  scale_fill_continuous_divergingx(palette = 'RdBu', mid = 0, na.value="white") +
  theme(axis.text.x = element_text(angle = 45, vjust = 1, hjust = 1), axis.title.x=element_blank(), axis.title.y=element_blank()) 

singlecellheatmap + ggplot2::coord_flip()
```

```
#export 1000x500
```

### 4 Session info

Packages and versions necessary to reproduce the results in this
report.

```
sessionInfo()
```

```
## R version 4.3.1 (2023-06-16 ucrt)
## Platform: x86_64-w64-mingw32/x64 (64-bit)
## Running under: Windows 10 x64 (build 19045)
## 
## Matrix products: default
## 
## 
## locale:
## [1] LC_COLLATE=English_United States.utf8 
## [2] LC_CTYPE=English_United States.utf8   
## [3] LC_MONETARY=English_United States.utf8
## [4] LC_NUMERIC=C                          
## [5] LC_TIME=English_United States.utf8    
## 
## time zone: America/New_York
## tzcode source: internal
## 
## attached base packages:
## [1] stats4    stats     graphics  grDevices utils     datasets  methods  
## [8] base     
## 
## other attached packages:
##  [1] eulerr_7.0.0                            
##  [2] svglite_2.1.2                           
##  [3] ggraph_2.1.0                            
##  [4] colorspace_2.1-0                        
##  [5] ggpubr_0.6.0                            
##  [6] lattice_0.22-5                          
##  [7] org.Dr.eg.db_3.18.0                     
##  [8] ensembldb_2.26.0                        
##  [9] AnnotationFilter_1.26.0                 
## [10] GenomicFeatures_1.54.1                  
## [11] ChIPseeker_1.38.0                       
## [12] orthogene_1.8.0                         
## [13] plotly_4.10.3                           
## [14] BaseSet_0.9.0                           
## [15] ontologyIndex_2.11                      
## [16] enrichplot_1.22.0                       
## [17] msigdbr_7.5.1                           
## [18] clusterProfiler_4.10.0                  
## [19] gprofiler2_0.2.2                        
## [20] GSVA_1.50.0                             
## [21] GSEABase_1.64.0                         
## [22] graph_1.80.0                            
## [23] annotate_1.80.0                         
## [24] XML_3.99-0.14                           
## [25] AnnotationDbi_1.64.0                    
## [26] DT_0.30                                 
## [27] RColorBrewer_1.1-3                      
## [28] ggrepel_0.9.4                           
## [29] scales_1.2.1                            
## [30] viridis_0.6.4                           
## [31] viridisLite_0.4.2                       
## [32] cowplot_1.1.1                           
## [33] DESeq2_1.42.0                           
## [34] SummarizedExperiment_1.32.0             
## [35] Biobase_2.62.0                          
## [36] MatrixGenerics_1.14.0                   
## [37] matrixStats_1.1.0                       
## [38] GenomicRanges_1.54.1                    
## [39] GenomeInfoDb_1.38.0                     
## [40] IRanges_2.36.0                          
## [41] S4Vectors_0.40.1                        
## [42] BiocGenerics_0.48.0                     
## [43] lubridate_1.9.3                         
## [44] forcats_1.0.0                           
## [45] stringr_1.5.0                           
## [46] dplyr_1.1.3                             
## [47] purrr_1.0.2                             
## [48] readr_2.1.4                             
## [49] tidyr_1.3.0                             
## [50] tibble_3.2.1                            
## [51] ggplot2_3.4.4                           
## [52] tidyverse_2.0.0                         
## [53] knitr_1.45                              
## [54] tinytex_0.48                            
## [55] rmarkdown_2.25                          
## [56] TxDb.Hsapiens.UCSC.hg38.knownGene_3.18.0
## [57] EnsDb.Hsapiens.v75_2.99.0               
## 
## loaded via a namespace (and not attached):
##   [1] ProtGenerics_1.34.0                    
##   [2] fs_1.6.3                               
##   [3] bitops_1.0-7                           
##   [4] HDO.db_0.99.1                          
##   [5] httr_1.4.7                             
##   [6] tools_4.3.1                            
##   [7] backports_1.4.1                        
##   [8] utf8_1.2.4                             
##   [9] R6_2.5.1                               
##  [10] HDF5Array_1.30.0                       
##  [11] lazyeval_0.2.2                         
##  [12] rhdf5filters_1.14.0                    
##  [13] withr_2.5.2                            
##  [14] prettyunits_1.2.0                      
##  [15] gridExtra_2.3                          
##  [16] cli_3.6.1                              
##  [17] scatterpie_0.2.1                       
##  [18] labeling_0.4.3                         
##  [19] sass_0.4.7                             
##  [20] systemfonts_1.0.5                      
##  [21] Rsamtools_2.18.0                       
##  [22] yulab.utils_0.1.0                      
##  [23] gson_0.1.0                             
##  [24] DOSE_3.28.0                            
##  [25] plotrix_3.8-2                          
##  [26] rstudioapi_0.15.0                      
##  [27] RSQLite_2.3.2                          
##  [28] TxDb.Hsapiens.UCSC.hg19.knownGene_3.2.2
##  [29] generics_0.1.3                         
##  [30] gridGraphics_0.5-1                     
##  [31] BiocIO_1.12.0                          
##  [32] crosstalk_1.2.0                        
##  [33] vroom_1.6.4                            
##  [34] gtools_3.9.4                           
##  [35] car_3.1-2                              
##  [36] homologene_1.4.68.19.3.27              
##  [37] GO.db_3.18.0                           
##  [38] Matrix_1.6-1.1                         
##  [39] fansi_1.0.5                            
##  [40] abind_1.4-5                            
##  [41] lifecycle_1.0.4                        
##  [42] yaml_2.3.7                             
##  [43] carData_3.0-5                          
##  [44] gplots_3.1.3                           
##  [45] rhdf5_2.46.0                           
##  [46] qvalue_2.34.0                          
##  [47] SparseArray_1.2.0                      
##  [48] BiocFileCache_2.10.1                   
##  [49] grid_4.3.1                             
##  [50] blob_1.2.4                             
##  [51] promises_1.2.1                         
##  [52] crayon_1.5.2                           
##  [53] beachmat_2.18.0                        
##  [54] KEGGREST_1.42.0                        
##  [55] pillar_1.9.0                           
##  [56] fgsea_1.28.0                           
##  [57] boot_1.3-28.1                          
##  [58] rjson_0.2.21                           
##  [59] codetools_0.2-19                       
##  [60] fastmatch_1.1-4                        
##  [61] glue_1.6.2                             
##  [62] ggfun_0.1.3                            
##  [63] data.table_1.14.8                      
##  [64] vctrs_0.6.4                            
##  [65] png_0.1-8                              
##  [66] treeio_1.26.0                          
##  [67] gtable_0.3.4                           
##  [68] cachem_1.0.8                           
##  [69] xfun_0.41                              
##  [70] S4Arrays_1.2.0                         
##  [71] mime_0.12                              
##  [72] tidygraph_1.2.3                        
##  [73] SingleCellExperiment_1.24.0            
##  [74] interactiveDisplayBase_1.40.0          
##  [75] ellipsis_0.3.2                         
##  [76] nlme_3.1-163                           
##  [77] ggtree_3.10.0                          
##  [78] bit64_4.0.5                            
##  [79] progress_1.2.2                         
##  [80] filelock_1.0.2                         
##  [81] bslib_0.5.1                            
##  [82] irlba_2.3.5.1                          
##  [83] KernSmooth_2.23-22                     
##  [84] DBI_1.1.3                              
##  [85] tidyselect_1.2.0                       
##  [86] bit_4.0.5                              
##  [87] compiler_4.3.1                         
##  [88] curl_5.1.0                             
##  [89] xml2_1.3.5                             
##  [90] DelayedArray_0.28.0                    
##  [91] shadowtext_0.1.2                       
##  [92] rtracklayer_1.62.0                     
##  [93] caTools_1.18.2                         
##  [94] rappdirs_0.3.3                         
##  [95] digest_0.6.33                          
##  [96] XVector_0.42.0                         
##  [97] htmltools_0.5.6.1                      
##  [98] pkgconfig_2.0.3                        
##  [99] sparseMatrixStats_1.14.0               
## [100] highr_0.10                             
## [101] dbplyr_2.4.0                           
## [102] fastmap_1.1.1                          
## [103] rlang_1.1.1                            
## [104] htmlwidgets_1.6.2                      
## [105] shiny_1.7.5.1                          
## [106] DelayedMatrixStats_1.24.0              
## [107] farver_2.1.1                           
## [108] jquerylib_0.1.4                        
## [109] jsonlite_1.8.7                         
## [110] BiocParallel_1.36.0                    
## [111] GOSemSim_2.28.0                        
## [112] BiocSingular_1.18.0                    
## [113] RCurl_1.98-1.12                        
## [114] magrittr_2.0.3                         
## [115] GenomeInfoDbData_1.2.11                
## [116] ggplotify_0.1.2                        
## [117] patchwork_1.1.3                        
## [118] Rhdf5lib_1.24.0                        
## [119] munsell_0.5.0                          
## [120] Rcpp_1.0.11                            
## [121] ggnewscale_0.4.9                       
## [122] ape_5.7-1                              
## [123] babelgene_22.9                         
## [124] stringi_1.8.1                          
## [125] zlibbioc_1.48.0                        
## [126] MASS_7.3-60                            
## [127] AnnotationHub_3.10.0                   
## [128] plyr_1.8.9                             
## [129] parallel_4.3.1                         
## [130] HPO.db_0.99.2                          
## [131] Biostrings_2.70.1                      
## [132] graphlayouts_1.0.1                     
## [133] splines_4.3.1                          
## [134] hms_1.1.3                              
## [135] locfit_1.5-9.8                         
## [136] igraph_1.5.1                           
## [137] ggsignif_0.6.4                         
## [138] reshape2_1.4.4                         
## [139] biomaRt_2.58.0                         
## [140] ScaledMatrix_1.10.0                    
## [141] BiocVersion_3.18.0                     
## [142] evaluate_0.23                          
## [143] BiocManager_1.30.22                    
## [144] tzdb_0.4.0                             
## [145] tweenr_2.0.2                           
## [146] httpuv_1.6.12                          
## [147] grr_0.9.5                              
## [148] polyclip_1.10-6                        
## [149] ggforce_0.4.1                          
## [150] rsvd_1.0.5                             
## [151] broom_1.0.5                            
## [152] xtable_1.8-4                           
## [153] restfulr_0.0.15                        
## [154] tidytree_0.4.5                         
## [155] MPO.db_0.99.7                          
## [156] rstatix_0.7.2                          
## [157] later_1.3.1                            
## [158] snow_0.4-4                             
## [159] aplot_0.2.2                            
## [160] GenomicAlignments_1.38.0               
## [161] memoise_2.0.1                          
## [162] timechange_0.2.0
```
