## Supplementary material for "Diencephalic and Neuropeptidergic Dysfunction in Zebrafish with Autism Risk Mutations": Data S1 - RNA-seq analysis files: deaf1_RNAseq.html

2dpf versus 6dpf Deaf1 RNAseq analysis


Code 

- Show All Code
- Hide All Code

### 2dpf versus 6dpf Deaf1 RNAseq analysis

###### Anna Moyer

#### *2023-11-15*

Make plots for Deaf1b RNAseq data collected from 2dpf and 6dpf
zebrafish head cuts. For 2dpf, there are three mutations (c207y, t238p,
and 23bpi) and each has wt, het, and hom samples. For 6dpf, there is
only the 23bpi mutation.

### 1 Setup

#### 1.1 Set seed and working directory

```
#set random seed
set.seed(123)

#set working directory
setwd("C:/Users/annam/Desktop/lab work/autismrnaseq/deaf1_2dpf-6dpf")
```

#### 1.2 Load packages

```
library("tidyverse") 
library("DESeq2") #finding differentially expressed genes 
library("cowplot") #arranging plots into grids
library("viridis") #viridis color schemes
library("scales") #use to get color schemes for viridis
library("ggrepel") #repels text labels on plots
library("RColorBrewer") #pick colors
library("DT") #interactive and searchable tables of our GSEA results
library("GSEABase") #functions and methods for Gene Set Enrichment Analysis
library("Biobase") #base functions for bioconductor; required by GSEABase
library("GSVA") #Gene Set Variation Analysis, a non-parametric and unsupervised method for estimating variation of gene set enrichment across samples.
library("gprofiler2") #tools for accessing the GO enrichment results using g:Profiler web resources
library("clusterProfiler") # provides a suite of tools for functional enrichment analysis
library("msigdbr") # access to msigdb collections directly within R
library("enrichplot") # great for making the standard GSEA enrichment plots
library("ontologyIndex") #for parsing obo files
library("BaseSet") #for importing gaf file
library("plotly") #make interactive plots
library("orthogene") #convert gene names between species
library("ChIPseeker") #for annotating and browsing chip-seq data
library("TxDb.Hsapiens.UCSC.hg38.knownGene", pos = .Machine$integer.max) #human gene annotation for chip-seq data
library("EnsDb.Hsapiens.v75", pos = .Machine$integer.max) #human gene annotation for chip-seq data
library("org.Dr.eg.db") #convert ensembl gene names to gene id
library("lattice") #used for making manhattan plot
library("ggpubr") #calculating correlation coefficient 
library("colorspace") #colors for heatmap
library("ggraph") #library for making network graphs
library("svglite") #export svgs in loop
```

### 2 Import files and run DESeq2

#### 2.1 Import files, make count matrix, and run DESeq for 2dpf

```
#import sample file with columns containing sample name, file name, condition, and replicate
sampleTable <- read_csv("input/input_deaf1together.csv")

#make conditions a factor
sampleTable$condition <- factor(sampleTable$condition,  levels = c("d23bpi_wt", "dc207y_wt", "dt238p_wt", "d23bpi_het", "dc207y_het", "dt238p_het", "d23bpi_hom", "dc207y_hom", "dt238p_hom"))

#create dds based on sample data
ddsHTSeq <-DESeqDataSetFromHTSeqCount(sampleTable=sampleTable, design=~condition)
dds <-DESeq(ddsHTSeq)

#prefilter (only keep genes that don't have counts of 0 in more than 3 samples)
keep <- rowSums(counts(dds) == 0) < 4
dds <- dds[keep,]

#export normalized counts for all samples
#each gene (with llgene ID, gene name, and entrez ID) and then normalized counts for each sample
countsdata <-counts(dds,normalized=TRUE)
namelist <-read.csv("input/LLgeneID_entrezID.csv")
##want to merge countsdata with namelist and gene_name
rownames(namelist) <- namelist[,1]
gene_name <-read.csv("input/llgeneid_genename.csv") ##this file has LLgeneID as column 1 and LLgeneAbbrev/gene name as column 2
rownames(gene_name) <- gene_name[,1]
genecounts<-merge(gene_name,namelist,by=0)
rownames(genecounts)<-genecounts[,1]
genecounts<-merge(genecounts,countsdata,by=0)
genecounts<-genecounts[,-(1:3)]

#export csv with normalized gene counts for each gene and each sample
write.csv(genecounts,file = "comparisons/normalized_reads_gene_list_2dpf.csv")

#plot PCA with all samples
plotPCA(rlog(dds),"condition")
```

The 2dpf samples cluster by clutch (ie, het, hom, and wt siblings
together) rather than different mutants clustering together.

#### 2.2 Make all pairwise comparisons

This uses a loop to make planned comparisons between wt/hom, wt/het,
and het/hom for all three 2 dpf mutants. It makes one csv file per
comparison and one summary file with all comparisons
(“deaf1DEGdata\_2dpf”).

```
#import data frame with the comparisons that we want to make
#should have "name" of comparison, first group for comparison, and second group for comparison
#"wt" group is in the third columnm, 
deaf1comparisons <- read.csv("input/deaf1_plannedcomparisons.csv")

#make an empty table to hold data
deaf1DEGdata <- setNames(data.frame(matrix(ncol = 9, nrow = 0)), c("LLgeneID", "baseMean", "log2FoldChange", "lfcSE", "stat", "pvalue", "padj", "LLgeneAbbrev", "comparison"))

#make a loop that performs each comparison
for (x in 1:length(deaf1comparisons$comparison.name)) {
  experimental <- deaf1comparisons[x,2] #name of experimental group
  control <- deaf1comparisons[x,3] #name of control group
  comparisonname <- deaf1comparisons[x,1] #name of comparison
  res <- results(dds, contrast=c("condition",experimental,control))
  res<-res[order(res$pvalue),]
  rlog<-rlog(dds)
  vst <-vst(dds)
  dataframe_res <-as.data.frame(res)
  dataframe_res$LLgeneID<-row.names(dataframe_res)
  dataframe_res<-dataframe_res[c(7,1:6)]
  res_gene <-inner_join(dataframe_res,gene_name,by="LLgeneID")
  res_gene<-res_gene[!is.na(res_gene$padj),]
  res_gene$comparison <- comparisonname #add column with comparison name
  write.csv(res_gene,file=paste0("comparisons/deaf1_", comparisonname, ".csv")) #export file
  deaf1DEGdata <- rbind(deaf1DEGdata, res_gene) #add comparison to table with all data
}

#export table with all comparisons
write.csv(deaf1DEGdata, "comparisons/deaf1DEGdata_2dpf.csv")

#make a copy of the 2dpf data
deaf1DEGdata_2dpf <- deaf1DEGdata
```

#### 2.3 Import files, make count matrix, and run DESeq for 6dpf

```
#import sample file with columns containing sample name, file name, condition, and replicate
sampleTable_6dpf <- read_csv("input/input_deaf1together_6dpf.csv")

#make conditions a factor
sampleTable_6dpf$condition <- factor(sampleTable_6dpf$condition,  levels = c("d23bpi_wt_6dpf", "d23bpi_het_6dpf", "d23bpi_hom_6dpf"))

#create dds based on sample data
ddsHTSeq_6dpf <-DESeqDataSetFromHTSeqCount(sampleTable=sampleTable_6dpf, design=~condition)
dds_6dpf <-DESeq(ddsHTSeq_6dpf)

#prefilter 
keep <- rowSums(counts(dds_6dpf) == 0) < 4
dds_6dpf <- dds_6dpf[keep,]

#export normalized counts for all samples
#each gene (with llgene ID, gene name, and entrez ID) and then normalized counts for each sample
countsdata_6dpf <-counts(dds_6dpf,normalized=TRUE)
genecounts_6dpf<-merge(gene_name,namelist,by=0)
rownames(genecounts_6dpf)<-genecounts_6dpf[,1]
genecounts_6dpf<-merge(genecounts_6dpf,countsdata_6dpf,by=0)
genecounts_6dpf<-genecounts_6dpf[,-(1:3)]

#export csv with normalized gene counts for each gene and each sample
write.csv(genecounts_6dpf,file = "comparisons/normalized_reads_gene_list_6dpf.csv")

#plot PCA with all samples
plotPCA(rlog(dds_6dpf),"condition")

#add rownames to genecounts
rownames(genecounts) <- genecounts$LLgeneID.y

#add rownames to genecounts_6dpf
rownames(genecounts_6dpf) <- genecounts_6dpf$LLgeneID.y

#merge 2dpf and 6dpf counts
genecounts <- merge(genecounts,genecounts_6dpf, by="row.names", all=TRUE)

#if NA in LLgeneAbbrev.x, get value from LLgeneAbbrev.y
genecounts <- genecounts %>% mutate(LLgeneAbbrev.x = ifelse(is.na(LLgeneAbbrev.x), LLgeneAbbrev.y, LLgeneAbbrev.x))
genecounts <- genecounts %>% mutate(LLgeneID.y.x = ifelse(is.na(LLgeneID.y.x), LLgeneID.y.y, LLgeneID.y.x))
genecounts <- genecounts %>% mutate(EntrezGeneID.x = ifelse(is.na(EntrezGeneID.x), EntrezGeneID.y, EntrezGeneID.x))

#get rid of extra columns
genecounts <- genecounts %>% dplyr::select(-c(Row.names, LLgeneAbbrev.y, LLgeneID.y.y, EntrezGeneID.y))

#rename columns
colnames(genecounts)[1:3] <- c("LLgeneAbbrev", "LLgeneID.y", "EntrezGeneID")

#export csv with normalized gene counts for both 2dpf and 6dpf
write.csv(genecounts,file = "comparisons/normalized_reads_gene_list_2dpf-6dpf.csv")
```

#### 2.4 Make all pairwise comparisons

This uses a loop to make planned comparisons between wt/hom, wt/het,
and het/hom for 23bp insertion 6dpf mutant. It makes one csv file per
comparison and one summary file with all comparisons
(“deaf1DEGdata”).

```
#import data frame with the comparisons that we want to make
#should have "name" of comparison, first group for comparison, and second group for comparison
#"wt" group is in the third column
deaf1comparisons_6dpf <- read.csv("input/deaf1_plannedcomparisons_6dpf.csv")

#make an empty table to hold data
deaf1DEGdata_6dpf <- setNames(data.frame(matrix(ncol = 9, nrow = 0)), c("LLgeneID", "baseMean", "log2FoldChange", "lfcSE", "stat", "pvalue", "padj", "LLgeneAbbrev", "comparison"))

#make a loop that performs each comparison
for (x in 1:length(deaf1comparisons_6dpf$comparison.name)) {
  experimental <- deaf1comparisons_6dpf[x,2] #name of experimental group
  control <- deaf1comparisons_6dpf[x,3] #name of control group
  comparisonname <- deaf1comparisons_6dpf[x,1] #name of comparison
  res <- results(dds_6dpf, contrast=c("condition",experimental,control))
  res<-res[order(res$pvalue),]
  rlog<-rlog(dds_6dpf)
  vst <-vst(dds_6dpf)
  dataframe_res <-as.data.frame(res)
  dataframe_res$LLgeneID<-row.names(dataframe_res)
  dataframe_res<-dataframe_res[c(7,1:6)]
  res_gene <-inner_join(dataframe_res,gene_name,by="LLgeneID")
  res_gene<-res_gene[!is.na(res_gene$padj),]
  res_gene$comparison <- comparisonname #add column with comparison name
  write.csv(res_gene,file=paste0("comparisons/deaf1_", comparisonname, ".csv")) #export file
  deaf1DEGdata_6dpf <- rbind(deaf1DEGdata_6dpf, res_gene) #add comparison to table with all data
}

#export table with all comparisons
write.csv(deaf1DEGdata_6dpf, "comparisons/deaf1DEGdata_6dpf.csv")

#combine 2dpf and 6dpf comparisons
deaf1DEGdata <- rbind(deaf1DEGdata_2dpf, deaf1DEGdata_6dpf)

#add some information to deaf1DEGdata
#limit padj to 1e-10
deaf1DEGdata <- deaf1DEGdata %>% mutate(padjbound = ifelse(deaf1DEGdata$padj < 1e-10, 1e-10, deaf1DEGdata$padj))

#limit pvalue to 1e-10
deaf1DEGdata <- deaf1DEGdata %>% mutate(pvaluebound = ifelse(deaf1DEGdata$pvalue < 1e-10, 1e-10, deaf1DEGdata$pvalue))

#import annotations for chromosome and TSS
chromosomeinfo <- readr::read_tsv(file="ncbi_refseqgenes")

#keep chromosome and gene name
chromosomeinfo <- chromosomeinfo %>% dplyr::select(chrom, name2, txStart)

deaf1DEGdata$chrom <- NA
deaf1DEGdata$txStart <- NA

#add chromosome and txStart information to deaf1DEGdata
for (x in 1:length(deaf1DEGdata$LLgeneAbbrev)) {
  gene <- deaf1DEGdata$LLgeneAbbrev[x]
  chromosomeinfosub <- chromosomeinfo %>% dplyr::filter(name2 == gene)
  chr <- chromosomeinfosub$chrom[1]
  txStart <- chromosomeinfosub$txStart[1]
  deaf1DEGdata$chrom[x] <- chr
  deaf1DEGdata$txStart[x] <- txStart
}

#add a column with chromosome number
deaf1DEGdata <- deaf1DEGdata %>% mutate(chromosomenumber = str_replace_all(chrom, c("chr1"="1", "chr2"="2", "chr3"="3", "chr4"="4", "chr5"="5", "chr6"="6", "chr7"="7", "chr8"="8", "chr9"="9", "chr10"="10", "chr11"="11", "chr12"="12", "chr13"="13", "chr14"="14", "chr15"="15", "chr16"="16", "chr17"="17", "chr18"="18", "chr19"="19", "chr20"="20", "chr21"="21", "chr22"="22", "chr23"="23", "chr24"="24", "chr25"="25")))

deaf1DEGdata <- distinct(deaf1DEGdata)

#export csv with DEG for both 2dpf and 6dpf
write.csv(deaf1DEGdata,file = "comparisons/Deaf1_DEG_2dpf-6dpf.csv")
```

### 3 Plotting DEGs by chromosome

Are differentially expressed genes located at a particular place in
the genome?

#### 3.1 Make Manhattan plot for 23bp 2dpf

This function is from: https://genome.sph.umich.edu/wiki/Code\_Sample:\_Generating\_Manhattan\_Plots\_in\_R

```
#import  comparisons (to skip running previous steps)
deaf1DEGdata <- read.csv("comparisons/Deaf1_DEG_2dpf-6dpf.csv")[,2:15]

#import  normalized counts (to skip running previous steps)
genecounts <- read.csv("comparisons/normalized_reads_gene_list_2dpf-6dpf.csv")[,2:57]

#function for making manhattan plots
manhattan.plot<-function(chr, pos, pvalue, 
    sig.level=NA, annotate=NULL, ann.default=list(),
    should.thin=T, thin.pos.places=2, thin.logp.places=2, 
    xlab="Chromosome", ylab=expression(-log[10](p-value)),
    col=c("gray","darkgray"), panel.extra=NULL, pch=20, cex=0.8,...) {

    if (length(chr)==0) stop("chromosome vector is empty")
    if (length(pos)==0) stop("position vector is empty")
    if (length(pvalue)==0) stop("pvalue vector is empty")

    #make sure we have an ordered factor
    if(!is.ordered(chr)) {
        chr <- ordered(chr)
    } else {
        chr <- chr[,drop=T]
    }

    #make sure positions are in kbp
    if (any(pos>1e6)) pos<-pos/1e6;

    #calculate absolute genomic position
    #from relative chromosomal positions
    posmin <- tapply(pos,chr, min);
    posmax <- tapply(pos,chr, max);
    posshift <- head(c(0,cumsum(posmax)),-1);
    names(posshift) <- levels(chr)
    genpos <- pos + posshift[chr];
    getGenPos<-function(cchr, cpos) {
        p<-posshift[as.character(cchr)]+cpos
        return(p)
    }

    #parse annotations
    grp <- NULL
    ann.settings <- list()
    label.default<-list(x="peak",y="peak",adj=NULL, pos=3, offset=0.5, 
        col=NULL, fontface=NULL, fontsize=NULL, show=F)
    parse.label<-function(rawval, groupname) {
        r<-list(text=groupname)
        if(is.logical(rawval)) {
            if(!rawval) {r$show <- F}
        } else if (is.character(rawval) || is.expression(rawval)) {
            if(nchar(rawval)>=1) {
                r$text <- rawval
            }
        } else if (is.list(rawval)) {
            r <- modifyList(r, rawval)
        }
        return(r)
    }

    if(!is.null(annotate)) {
        if (is.list(annotate)) {
            grp <- annotate[[1]]
        } else {
            grp <- annotate
        } 
        if (!is.factor(grp)) {
            grp <- factor(grp)
        }
    } else {
        grp <- factor(rep(1, times=length(pvalue)))
    }
  
    ann.settings<-vector("list", length(levels(grp)))
    ann.settings[[1]]<-list(pch=pch, col=col, cex=cex, fill=col, label=label.default)

    if (length(ann.settings)>1) { 
        lcols<-trellis.par.get("superpose.symbol")$col 
        lfills<-trellis.par.get("superpose.symbol")$fill
        for(i in 2:length(levels(grp))) {
            ann.settings[[i]]<-list(pch=pch, 
                col=lcols[(i-2) %% length(lcols) +1 ], 
                fill=lfills[(i-2) %% length(lfills) +1 ], 
                cex=cex, label=label.default);
            ann.settings[[i]]$label$show <- T
        }
        names(ann.settings)<-levels(grp)
    }
    for(i in 1:length(ann.settings)) {
        if (i>1) {ann.settings[[i]] <- modifyList(ann.settings[[i]], ann.default)}
        ann.settings[[i]]$label <- modifyList(ann.settings[[i]]$label, 
            parse.label(ann.settings[[i]]$label, levels(grp)[i]))
    }
    if(is.list(annotate) && length(annotate)>1) {
        user.cols <- 2:length(annotate)
        ann.cols <- c()
        if(!is.null(names(annotate[-1])) && all(names(annotate[-1])!="")) {
            ann.cols<-match(names(annotate)[-1], names(ann.settings))
        } else {
            ann.cols<-user.cols-1
        }
        for(i in seq_along(user.cols)) {
            if(!is.null(annotate[[user.cols[i]]]$label)) {
                annotate[[user.cols[i]]]$label<-parse.label(annotate[[user.cols[i]]]$label, 
                    levels(grp)[ann.cols[i]])
            }
            ann.settings[[ann.cols[i]]]<-modifyList(ann.settings[[ann.cols[i]]], 
                annotate[[user.cols[i]]])
        }
    }
    rm(annotate)

    #reduce number of points plotted
    if(should.thin) {
        thinned <- unique(data.frame(
            logp=round(-log10(pvalue),thin.logp.places), 
            pos=round(genpos,thin.pos.places), 
            chr=chr,
            grp=grp)
        )
        logp <- thinned$logp
        genpos <- thinned$pos
        chr <- thinned$chr
        grp <- thinned$grp
        rm(thinned)
    } else {
        logp <- -log10(pvalue)
    }
    rm(pos, pvalue)
    gc()

    #custom axis to print chromosome names
    axis.chr <- function(side,...) {
        if(side=="bottom") {
            panel.axis(side=side, outside=T,
                at=((posmax+posmin)/2+posshift),
                labels=levels(chr), 
                ticks=F, rot=0,
                check.overlap=F
            )
        } else if (side=="top" || side=="right") {
            panel.axis(side=side, draw.labels=F, ticks=F);
        }
        else {
            axis.default(side=side,...);
        }
     }

    #make sure the y-lim covers the range (plus a bit more to look nice)
    prepanel.chr<-function(x,y,...) { 
        A<-list();
        maxy<-ceiling(max(y, ifelse(!is.na(sig.level), -log10(sig.level), 0)))+.5;
        A$ylim=c(0,maxy);
        A;
    }

    xyplot(logp~genpos, chr=chr, groups=grp,
        axis=axis.chr, ann.settings=ann.settings, 
        prepanel=prepanel.chr, scales=list(axs="i"),
        panel=function(x, y, ..., getgenpos) {
            if(!is.na(sig.level)) {
                #add significance line (if requested)
                panel.abline(h=-log10(sig.level), lty=2);
            }
            panel.superpose(x, y, ..., getgenpos=getgenpos);
            if(!is.null(panel.extra)) {
                panel.extra(x,y, getgenpos, ...)
            }
        },
        panel.groups = function(x,y,..., subscripts, group.number) {
            A<-list(...)
            #allow for different annotation settings
            gs <- ann.settings[[group.number]]
            A$col.symbol <- gs$col[(as.numeric(chr[subscripts])-1) %% length(gs$col) + 1]    
            A$cex <- gs$cex[(as.numeric(chr[subscripts])-1) %% length(gs$cex) + 1]
            A$pch <- gs$pch[(as.numeric(chr[subscripts])-1) %% length(gs$pch) + 1]
            A$fill <- gs$fill[(as.numeric(chr[subscripts])-1) %% length(gs$fill) + 1]
            A$x <- x
            A$y <- y
            do.call("panel.xyplot", A)
            #draw labels (if requested)
            if(gs$label$show) {
                gt<-gs$label
                names(gt)[which(names(gt)=="text")]<-"labels"
                gt$show<-NULL
                if(is.character(gt$x) | is.character(gt$y)) {
                    peak = which.max(y)
                    center = mean(range(x))
                    if (is.character(gt$x)) {
                        if(gt$x=="peak") {gt$x<-x[peak]}
                        if(gt$x=="center") {gt$x<-center}
                    }
                    if (is.character(gt$y)) {
                        if(gt$y=="peak") {gt$y<-y[peak]}
                    }
                }
                if(is.list(gt$x)) {
                    gt$x<-A$getgenpos(gt$x[[1]],gt$x[[2]])
                }
                do.call("panel.text", gt)
            }
        },
        xlab=xlab, ylab=ylab, 
        panel.extra=panel.extra, getgenpos=getGenPos, ...
    );
}

#select columns to include in manhattan plot
myTopHits.df <- deaf1DEGdata %>% dplyr::select(chromosomenumber, txStart, pvaluebound, comparison)

#filter to only include chromosomes 1-25
myTopHits.df <- myTopHits.df %>% dplyr::filter(chromosomenumber %in% 1:25)

#make chromosomes plotted in order
myTopHits.df$chromosomenumber <- factor(myTopHits.df$chromosomenumber, levels = 1:25)

#omit NA values
myTopHits.df <- na.omit(myTopHits.df)

#make colors 
manhattancol <- c(replicate(12, c("lightgrey", "darkgrey")), "#21908CFF")
  
#filter comparisons to only include 23bp hom v wt comparisons
manhattan <- myTopHits.df %>% dplyr::filter(comparison == "d23bp_homvwt")

#make manhattan plot
manhattan.plot(manhattan$chromosomenumber, manhattan$txStart, manhattan$pvaluebound, should.thin=F, col=manhattancol)
```

```
#export manhattan plots at 1000x500 pixels
```

#### 3.2 Make Manhattan plot for c207y 2dpf

```
#filter comparisons to only include one comparison
manhattan <- myTopHits.df %>% dplyr::filter(comparison == "dc207y_homvwt")

#make manhattan plot
manhattan.plot(manhattan$chromosomenumber, manhattan$txStart, manhattan$pvaluebound, should.thin=F, col=manhattancol)
```

#### 3.3 Make Manhattan plot for t238p 2dpf

```
#filter comparisons to only include one comparison
manhattan <- myTopHits.df %>% dplyr::filter(comparison == "dt238p_homvwt")

#make manhattan plot
manhattan.plot(manhattan$chromosomenumber, manhattan$txStart, manhattan$pvaluebound, should.thin=F, col=manhattancol)
```

#### 3.4 Make Manhattan plot for 23bp 6dpf

```
#filter comparisons to only include one comparison
manhattan <- myTopHits.df %>% dplyr::filter(comparison == "d23bp_homvwt_6dpf")

#make manhattan plot
manhattan.plot(manhattan$chromosomenumber, manhattan$txStart, manhattan$pvaluebound, should.thin=F, col=manhattancol)
```

#### 3.5 Make Manhattan plot for mouse DEAF1 mutant data

```
#import DEAF1 mouse hippocampus
DEAF1mousedata <- read.csv("comparisonrnaseq/deaf1mousehippocampus.csv")

#keep only columns for manhattan
DEAF1mousedata <- DEAF1mousedata %>% dplyr::select(chromosome_name, start_position, PValue)

#make chromosomes plotted in order
DEAF1mousedata$chromosome_name <- factor(DEAF1mousedata$chromosome_name, levels = c(1:19, "X", "Y", "MT"))

#omit NA values
DEAF1mousedata <- na.omit(DEAF1mousedata)

#make colors for mouse manhattan
manhattancol_mouse <- c(replicate(3, c("lightgrey", "darkgrey")), "#21908CFF", replicate(8, c("lightgrey", "darkgrey")))

#make manhattan plot
manhattan.plot(DEAF1mousedata$chromosome_name, DEAF1mousedata$start_position, DEAF1mousedata$PValue, should.thin=F, col=manhattancol_mouse)
```

Genes on chromosome 25 look overrepresented compared to genes on
other chromosomes. Next will test whether these genes are formally
overrepresented using GSEA.

#### 3.6 Make custom GSEA annotation with zebrafish genes and chromosomes

```
#import annotations for chromosome and gene name
chromosomeinfo <- readr::read_tsv(file="ncbi_refseqgenes")

#keep chromosome and gene name
chromosomeinfo <- chromosomeinfo %>% dplyr::select(chrom, name2)

#change column names
colnames(chromosomeinfo) <- c("chromosome", "genename")

#select only chromosomes 1-25
chromosomeinfo <- chromosomeinfo %>% dplyr::filter(chromosome %in% paste("chr", 1:25, sep=""))

#keep only distinct rows
chromosomeinfo <- distinct(chromosomeinfo)
```

#### 3.7 Check for enrichment of chromosome 25 genes in 23bp 2dpf

```
# Perform GSEA using clusterProfiler
#filter to only one comparison
GSEAgenes <- deaf1DEGdata %>% dplyr::filter(comparison == "d23bp_homvwt")

#keep only the columns we need for GSEA
mydata.df.sub <- dplyr::select(GSEAgenes, LLgeneAbbrev, log2FoldChange)

#sort by abs(foldchange)
mydata.df.sub$log2FoldChange <- abs(mydata.df.sub$log2FoldChange)

# construct a named vector
mydata.gsea <- mydata.df.sub$log2FoldChange
names(mydata.gsea) <- as.character(mydata.df.sub$LLgeneAbbrev)
mydata.gsea <- sort(mydata.gsea, decreasing = TRUE)

# run GSEA using the 'GSEA' function from clusterProfiler
myGSEA.res <- GSEA(mydata.gsea, TERM2GENE=chromosomeinfo, verbose=FALSE, scoreType="pos", maxGSSize=2000)
myGSEA.df <- as_tibble(myGSEA.res@result)

# view results as an interactive table
datatable(myGSEA.df, 
          extensions = c('KeyTable', "FixedHeader"), 
          options = list(keys = TRUE, searchHighlight = TRUE, pageLength = 10, lengthMenu = c("10", "25", "50", "100"))) %>%
  formatRound(columns=c(2:10), digits=2)
```

```
# create enrichment plots using the enrichplot package
gseaplot2(myGSEA.res, 
          geneSetID = c(1), #can choose multiple signatures to overlay in this plot
         pvalue_table = FALSE, #can set this to FALSE for a cleaner plot
        title = myGSEA.res$Description[1]) #can also turn off this title
```

#### 3.8 Check for enrichment of chromosome 25 genes in c207y 2dpf

```
# Perform GSEA using clusterProfiler
#filter to only one comparison
GSEAgenes <- deaf1DEGdata %>% dplyr::filter(comparison == "dc207y_homvwt")

#keep only the columns we need for GSEA
mydata.df.sub <- dplyr::select(GSEAgenes, LLgeneAbbrev, log2FoldChange)

#sort by abs(foldchange)
mydata.df.sub$log2FoldChange <- abs(mydata.df.sub$log2FoldChange)

# construct a named vector
mydata.gsea <- mydata.df.sub$log2FoldChange
names(mydata.gsea) <- as.character(mydata.df.sub$LLgeneAbbrev)
mydata.gsea <- sort(mydata.gsea, decreasing = TRUE)

# run GSEA using the 'GSEA' function from clusterProfiler
myGSEA.res <- GSEA(mydata.gsea, TERM2GENE=chromosomeinfo, verbose=FALSE, scoreType="pos", maxGSSize=2000)
myGSEA.df <- as_tibble(myGSEA.res@result)

# view results as an interactive table
datatable(myGSEA.df, 
          extensions = c('KeyTable', "FixedHeader"), 
          options = list(keys = TRUE, searchHighlight = TRUE, pageLength = 10, lengthMenu = c("10", "25", "50", "100"))) %>%
  formatRound(columns=c(2:10), digits=2)
```

```
# create enrichment plots using the enrichplot package
gseaplot2(myGSEA.res, 
          geneSetID = c(1), #can choose multiple signatures to overlay in this plot
         pvalue_table = FALSE, #can set this to FALSE for a cleaner plot
        title = myGSEA.res$Description[1]) #can also turn off this title
```

#### 3.9 Check for enrichment of chromosome 25 genes in t238p 2dpf

```
# Perform GSEA using clusterProfiler
#filter to only one comparison
GSEAgenes <- deaf1DEGdata %>% dplyr::filter(comparison == "dt238p_homvwt")

#keep only the columns we need for GSEA
mydata.df.sub <- dplyr::select(GSEAgenes, LLgeneAbbrev, log2FoldChange)

#sort by abs(foldchange)
mydata.df.sub$log2FoldChange <- abs(mydata.df.sub$log2FoldChange)

# construct a named vector
mydata.gsea <- mydata.df.sub$log2FoldChange
names(mydata.gsea) <- as.character(mydata.df.sub$LLgeneAbbrev)
mydata.gsea <- sort(mydata.gsea, decreasing = TRUE)

# run GSEA using the 'GSEA' function from clusterProfiler
myGSEA.res <- GSEA(mydata.gsea, TERM2GENE=chromosomeinfo, verbose=FALSE, scoreType="pos", maxGSSize=2000)
myGSEA.df <- as_tibble(myGSEA.res@result)

# view results as an interactive table
datatable(myGSEA.df, 
          extensions = c('KeyTable', "FixedHeader"), 
          options = list(keys = TRUE, searchHighlight = TRUE, pageLength = 10, lengthMenu = c("10", "25", "50", "100"))) %>%
  formatRound(columns=c(2:10), digits=2)
```

```
# create enrichment plots using the enrichplot package
gseaplot2(myGSEA.res, 
          geneSetID = c(1), #can choose multiple signatures to overlay in this plot
         pvalue_table = FALSE, #can set this to FALSE for a cleaner plot
        title = myGSEA.res$Description[1]) #can also turn off this title
```

#### 3.10 Check for enrichment of chromosome 25 genes in 23bp 6dpf

```
# Perform GSEA using clusterProfiler
#filter to only one comparison
GSEAgenes <- deaf1DEGdata %>% dplyr::filter(comparison == "d23bp_homvwt_6dpf")

#keep only the columns we need for GSEA
mydata.df.sub <- dplyr::select(GSEAgenes, LLgeneAbbrev, log2FoldChange)

#sort by abs(foldchange)
mydata.df.sub$log2FoldChange <- abs(mydata.df.sub$log2FoldChange)

# construct a named vector
mydata.gsea <- mydata.df.sub$log2FoldChange
names(mydata.gsea) <- as.character(mydata.df.sub$LLgeneAbbrev)
mydata.gsea <- sort(mydata.gsea, decreasing = TRUE)

# run GSEA using the 'GSEA' function from clusterProfiler
myGSEA.res <- GSEA(mydata.gsea, TERM2GENE=chromosomeinfo, verbose=FALSE, scoreType="pos", maxGSSize=2000)
myGSEA.df <- as_tibble(myGSEA.res@result)

# view results as an interactive table
datatable(myGSEA.df, 
          extensions = c('KeyTable', "FixedHeader"), 
          options = list(keys = TRUE, searchHighlight = TRUE, pageLength = 10, lengthMenu = c("10", "25", "50", "100"))) %>%
  formatRound(columns=c(2:10), digits=2)
```

```
# create enrichment plots using the enrichplot package
gseaplot2(myGSEA.res, 
          geneSetID = c(1), #can choose multiple signatures to overlay in this plot
         pvalue_table = FALSE, #can set this to FALSE for a cleaner plot
        title = myGSEA.res$Description[1]) #can also turn off this title
```

#### 3.11 Check for enrichment of chromosome 7 genes in mouse dataset

```
#import DEAF1 mouse hippocampus
GSEAgenes <- read.csv("comparisonrnaseq/deaf1mousehippocampus.csv")

#keep only the columns we need for GSEA
mydata.df.sub <- dplyr::select(GSEAgenes, Symbol, FC_NKOvsControl)

#abs(foldchange)
mydata.df.sub$FC_NKOvsControl <- abs(mydata.df.sub$FC_NKOvsControl)

# construct a named vector
mydata.gsea <- mydata.df.sub$FC_NKOvsControl
names(mydata.gsea) <- as.character(mydata.df.sub$Symbol)
mydata.gsea <- sort(mydata.gsea, decreasing = TRUE)

#import annotations for mouse chromosome and gene name
chromosomeinfo <- readr::read_tsv(file="ncbirefseqgenes_mouse")

#keep chromosome and gene name
chromosomeinfo <- chromosomeinfo %>% dplyr::select(chrom, name2)

#change column names
colnames(chromosomeinfo) <- c("chromosome", "genename")

#select only chromosomes 1-19 + XYM
chromosomeinfo <- chromosomeinfo %>% dplyr::filter(chromosome %in% c(paste("chr", 1:19, sep=""), "chrM", "chrX", "chrY"))

#keep only distinct rows
chromosomeinfo <- distinct(chromosomeinfo)

# run GSEA using the 'GSEA' function from clusterProfiler
myGSEA.res <- GSEA(mydata.gsea, TERM2GENE=chromosomeinfo, verbose=FALSE, scoreType="pos", maxGSSize=20000, pvalueCutoff = 1, pAdjustMethod="none")
myGSEA.df <- as_tibble(myGSEA.res@result)

# view results as an interactive table
datatable(myGSEA.df, 
          extensions = c('KeyTable', "FixedHeader"), 
          options = list(keys = TRUE, searchHighlight = TRUE, pageLength = 10, lengthMenu = c("10", "25", "50", "100"))) %>%
  formatRound(columns=c(2:10), digits=2)
```

Genes on chromosome 7 are not statistically significantly enriched in
the DEAF1 RNAseq data.

#### 3.12 Plot fold change of genes on chromosome 25 in chromosomal order

```
#keep only 2dpf comparisons
myTopHits.df <- deaf1DEGdata %>% dplyr::filter(comparison %in% c("d23bp_homvwt", "dc207y_homvwt", "dt238p_homvwt"))

#filter to only include those with logfc > 1
myTopHits.df <- myTopHits.df %>% dplyr::filter(abs(log2FoldChange) > 1)

#get names of genes with logfc > 1
genestoplot <- unique(myTopHits.df$LLgeneAbbrev)

#filter to only include 2dpf homvwt comparisons
myTopHits.df <- deaf1DEGdata %>% dplyr::filter(comparison %in% c("d23bp_homvwt", "dc207y_homvwt", "dt238p_homvwt"))

#filter to only include genes on chr 25
myTopHits.df <- myTopHits.df %>% dplyr::filter(chrom == "chr25")

#change back to fold change
myTopHits.df <- myTopHits.df %>% mutate(foldchange = 2^log2FoldChange)

#change the order of the genes
myTopHits.df <- myTopHits.df %>% arrange(txStart)

#filter to include only those with log2fc >1
myTopHits.df <- myTopHits.df %>% dplyr::filter(LLgeneAbbrev %in% genestoplot)

#order comparisons
myTopHits.df$comparison <- factor(myTopHits.df$comparison, levels = c("d23bp_homvwt", "dc207y_homvwt", "dt238p_homvwt"))

#rename comparisons
levels(myTopHits.df$comparison) <- c("23d46i 2dpf", "c207y 2dpf", "t238p 2dpf")

#make genes into a factor
myTopHits.df$LLgeneAbbrev <- factor(myTopHits.df$LLgeneAbbrev, levels=unique(myTopHits.df$LLgeneAbbrev))

plot <- ggplot(myTopHits.df, aes(y=log2FoldChange, x=LLgeneAbbrev, fill=factor(ifelse(LLgeneAbbrev == "deaf1", "Highlighted", "Normal")))) + 
    geom_bar(position="dodge", stat="summary", ) +
    ylab("log2 fold change") +
    theme_bw() +
    facet_wrap(~comparison,  ncol=1) +
    theme(axis.text.x = element_text(angle = 45, hjust=1), legend.position = "none", axis.title.x=element_blank()) +
    scale_fill_manual(values = c("#21908CFF", "gray35")) 


plot
```

```
#export 1300x700
```

#### 3.13 Plot fold change of genes on chromosome 25 in by TSS

```
#filter to only include 2dpf homvwt comparisons
myTopHits.df <- deaf1DEGdata %>% dplyr::filter(comparison %in% c("d23bp_homvwt", "dc207y_homvwt", "dt238p_homvwt"))

#filter to only include genes on chr 25
myTopHits.df <- myTopHits.df %>% dplyr::filter(chrom == "chr25")

#change the order of the genes
myTopHits.df <- myTopHits.df %>% arrange(txStart)

#order comparisons
myTopHits.df$comparison <- factor(myTopHits.df$comparison, levels = c("d23bp_homvwt", "dc207y_homvwt", "dt238p_homvwt"))

#rename comparisons
levels(myTopHits.df$comparison) <- c("23d46i 2dpf", "c207y 2dpf", "t238p 2dpf")


chr25plot <- ggplot(myTopHits.df, aes(y=log2FoldChange, x=txStart, color=factor(ifelse(LLgeneAbbrev == "deaf1", "Highlighted", "Normal")))) + 
    geom_bar(stat="summary" ) +
    ylab("log2 fold change") +
    xlab("TSS base position") +
    theme_bw() +
    facet_wrap(~comparison,  ncol=1) +
    theme(axis.text.x = element_text(angle = 45, hjust=1), legend.position = "none") +
    scale_color_manual(values = c("#21908CFF", "gray35")) +
    scale_x_continuous(n.breaks = 20, labels = scales::comma, limits = c(0, 38000000), expand = c(0, 0))


chr25plot
```

```
#export 1300x700
```

It looks like there’s a cluster of histone genes on the distal end of
chromosome 25 that is upregulated in 23bp and downregulated in
t238p.

#### 3.14 Find genes that are consistently up or downregulated in all three 2dpf homvwt comparisons

```
 #filter to only include homvwt comparisons
myTopHits.df <- deaf1DEGdata %>% dplyr::filter(comparison %in% c("d23bp_homvwt", "dc207y_homvwt", "dt238p_homvwt"))

#filter to only include genes on chr 25
myTopHits.df <- myTopHits.df %>% dplyr::filter(chrom == "chr25")

#filter to only include those with pval <0.05
myTopHits.df <- myTopHits.df %>% dplyr::filter(pvalue < 0.05)

#get list of genes
genesofinterest <- unique(myTopHits.df$LLgeneAbbrev)

#get data for list of genes
myTopHits.df <- deaf1DEGdata %>% dplyr::filter(LLgeneAbbrev %in% genesofinterest)

#filter to only include homvwt comparisons
myTopHits.df <- myTopHits.df %>% dplyr::filter(comparison %in% c("d23bp_homvwt", "dc207y_homvwt", "dt238p_homvwt"))

#keep only the columns needed to make a heat map
myTopHits.df <- myTopHits.df %>% dplyr::select(LLgeneAbbrev, comparison, log2FoldChange)

#pivot wider
myTopHits.df <- myTopHits.df %>% pivot_wider(names_from=comparison, values_from=log2FoldChange, id_cols = LLgeneAbbrev, values_fn = ~ mean(.x, na.rm = TRUE))

#filter to keep those all + and >0.3 in two conditions
keepup <- rowSums(myTopHits.df[,2:4] > 0) > 2
keepup <- myTopHits.df[keepup,]
keepupfold <- rowSums(keepup[,2:4] > 0.3) > 1
keepup <- keepup[keepupfold,]
keepup <- na.omit(keepup)

#filter to keep those all - and <-0.3 in two conditions
keepdown <- rowSums(myTopHits.df[,2:4] < 0) > 2
keepdown <- myTopHits.df[keepdown,]
keepdownfold <- rowSums(keepdown[,2:4] < -0.3) > 1
keepdown <- keepdown[keepdownfold,]
keepdown <- na.omit(keepdown)

#make list of genes to include
chr25genestokeep <- c(keepdown$LLgeneAbbrev, keepup$LLgeneAbbrev)

#filter to only include homvwt comparisons
myTopHits.df <- deaf1DEGdata %>% dplyr::filter(comparison %in% c("d23bp_homvwt", "dc207y_homvwt", "dt238p_homvwt"))

#filter to only include genes on chr 25
myTopHits.df <- myTopHits.df %>% dplyr::filter(chrom == "chr25")

#make list of all chr25 genes
chr25genes <- unique(myTopHits.df$LLgeneAbbrev)

#get rid of tes (had NA value)
chr25genestokeep <- setdiff(chr25genestokeep, "tes")

#chr25 genes to get rid of
chr25genes <- setdiff(chr25genes, chr25genestokeep)

chr25genestokeep
```

```
##  [1] "ptpn9a"       "muc5.1"       "saal1"        "hdac10"       "zgc:153293"  
##  [6] "morc2"        "ppib"         "fam185a"      "tnnt2d"       "cdkn1bb"     
## [11] "zgc:193812"   "crabp1a"      "lrrc29"       "pamr1"        "pdia3"       
## [16] "magi2b"       "abtb2b"       "zgc:154077"   "ccdc113"      "cpne8"       
## [21] "muc5.2"       "LOC110438378" "snupn"        "cav1"         "btbd10a"     
## [26] "parvaa"       "galk2"        "dkk3b"        "dusp6"        "vac14"       
## [31] "tmem17"       "kitlga"       "tbc1d2b"      "bbs4"         "lrrk2"       
## [36] "e2f4"         "tnni2a.2"     "mob2a"        "deaf1"
```

#### 3.15 Make bubble plot of chr25 genes that are consistent between the three mutations

```
#comparisons to include in plot
comparisons <- c("dt238p_homvwt", "dc207y_homvwt", "d23bp_homvwt_6dpf", "d23bp_homvwt")

#filter to only include genes of interest
bubbleDEG <- deaf1DEGdata %>% dplyr::filter(LLgeneAbbrev %in% chr25genestokeep & comparison %in% comparisons)

#find middle of foldchange (to center color scale)
limit <- max(abs(bubbleDEG$log2FoldChange)) * c(-1, 1)

#get color
myheatcolors3 <- brewer.pal(name="RdBu", n=11)

#set order of genes along x axis
geneorder <- c(rev(sort(keepup$LLgeneAbbrev)), rev(sort(keepdown$LLgeneAbbrev)))

#order genes
bubbleDEG$LLgeneAbbrev <- factor(bubbleDEG$LLgeneAbbrev, levels = geneorder)

#order comparisons
bubbleDEG$comparison <- factor(bubbleDEG$comparison, levels = rev(c("d23bp_homvwt_6dpf", "d23bp_homvwt", "dt238p_homvwt", "dc207y_homvwt")))

#rename comparisons
levels(bubbleDEG$comparison) <- rev(c("23d46i 6dpf", "23d46i 2dpf", "t238p 2dpf", "c207y 2dpf"))

#rename key legends
colnames(bubbleDEG) <- c("LLgeneID", "baseMean", "log2FoldChange", "lfcSE", "stat", "pvalue", "padjn", "LLgeneAbbrev", "comparison", "padj")

# create 'bubble plot' to summarize y signatures across x phenotypes
bubbleplot_chr25 <- ggplot(bubbleDEG, aes(x=LLgeneAbbrev, y=comparison)) + 
  geom_point(aes(size=-log10(padj), color = log2FoldChange)) +
  scale_color_gradientn(colors = myheatcolors3, limit=limit) +
  theme_bw() +
  theme(axis.text.x = element_text(angle = 45, hjust = 1),
        axis.title.x = element_blank(),
        axis.title.y = element_blank())+
  annotate("rect", xmin = 0, xmax = 20.5, ymin = 0.25, ymax = 0.5, 
           alpha=1, fill="#21908CFF") +
  annotate("rect", xmin = 20.5, xmax = 40, ymin = 0.25, ymax = 0.5, 
           alpha=1, fill="#472D7BFF") 

bubbleplot_chr25 + ggplot2::coord_flip()
```

```
#export 1000x300 for horizontal
#export 500x700 for vertical
```

### 4 Bubble Plots

#### 4.1 Find genes that are shared between comparisons

```
#add a column that is 1 if pvalue is <0.05 and +FC, -1 if padj is <0.05 and -FC
deaf1DEGdata2 <- deaf1DEGdata %>% mutate(Venn = ifelse(pvalue < 0.05 & log2FoldChange > 0, 1, ifelse(pvalue < 0.05 & log2FoldChange < 0, -1, 0)))

#filter to only include homvwt comparisons
deaf1DEGdata2 <- deaf1DEGdata2 %>% dplyr::filter(comparison %in% c("d23bp_homvwt", "dc207y_homvwt", "dt238p_homvwt", "d23bp_homvwt_6dpf"))

#make a new table with only gene name, comparison name, and Venn column
venntable <- deaf1DEGdata2 %>% dplyr::select(LLgeneAbbrev, comparison, Venn)

#find duplicates in venntable
duplicategenes <- dplyr::filter(venntable %>%
  distinct() %>%
  group_by(LLgeneAbbrev, comparison) %>%
  dplyr::count(), n !=1)$LLgeneAbbrev

#remove duplicate rows from venntable
venntable <- venntable %>% dplyr::filter(!LLgeneAbbrev %in% duplicategenes)

#use pivot wider to make untidy table
venntable <- as.matrix(venntable %>% distinct() %>%
    pivot_wider(names_from = comparison, values_from  = Venn, values_fill=0) %>%
    column_to_rownames('LLgeneAbbrev'))

#convert NULL to 0
venntable[venntable=='NULL'] <- 0

#find sum of values of rows
rowsum <- rowSums(venntable)
venntable <- cbind(venntable, rowsum)

#convert to dataframe
venntable <- as.data.frame(venntable)

#sort by those with highest sum of absolute values of rows
venntable <- venntable %>% arrange(desc(abs(rowsum)))

#filter by those with 2+ shared upregulated genes
venntableup <- rownames(venntable %>% dplyr::filter(rowsum > 1))
venntableup <- setdiff(venntableup, chr25genes)

#filter by those with 2+ shared downregulated genes
venntabledown <- rownames(venntable %>% dplyr::filter(rowsum < -1))
venntabledown <- setdiff(venntabledown, chr25genes)

#make vector with upregulated and downregulated genes together
venntableall <- c(venntableup, venntabledown)

#export
write.csv(venntable, "functionalanalysis/venntable_sorted.csv")
write.csv(venntableup, file="functionalanalysis/venntableup.csv")
write.csv(venntabledown, file="functionalanalysis/venntabledown.csv")
```

#### 4.2 Make bubble plot of genes shared between mutants

```
#genes to include in plot
geneofinterest <- read.csv(file="genesfromsharedannotation_concise.csv", header=FALSE)

#comparisons to include in plot
comparisons <- c("dt238p_homvwt", "dc207y_homvwt", "d23bp_homvwt_6dpf", "d23bp_homvwt")

#filter to only include genes of interest
bubbleDEG <- deaf1DEGdata %>% dplyr::filter(LLgeneAbbrev %in% geneofinterest$V1 & comparison %in% comparisons)

#find middle of foldchange (to center color scale)
limit <- max(abs(bubbleDEG$log2FoldChange)) * c(-1, 1)

#get color
myheatcolors3 <- brewer.pal(name="RdBu", n=11)

#order genes
bubbleDEG$LLgeneAbbrev <- factor(bubbleDEG$LLgeneAbbrev, levels = geneofinterest$V1)

#order comparisons
bubbleDEG$comparison <- factor(bubbleDEG$comparison, levels = c("d23bp_homvwt_6dpf", "d23bp_homvwt", "dt238p_homvwt", "dc207y_homvwt"))

#rename comparisons
levels(bubbleDEG$comparison) <- c("23d46i 6dpf", "23d46i 2dpf", "t238p 2dpf", "c207y 2dpf")

#rename key legends
colnames(bubbleDEG) <- c("LLgeneID", "baseMean", "log2FoldChange", "lfcSE", "stat", "pvalue", "padjn", "LLgeneAbbrev", "comparison", "padj")

# create 'bubble plot' to summarize y signatures across x phenotypes
bubbleplot_alldeaf1 <- ggplot(bubbleDEG, aes(x=LLgeneAbbrev, y=comparison)) + 
  geom_point(aes(size=-log10(padj), color = log2FoldChange)) +
  scale_color_gradientn(colors = myheatcolors3, limit=limit) +
  theme_bw() +
  theme(axis.text.x = element_text(angle = 45, hjust = 1),
        axis.title.x = element_blank(),
        axis.title.y = element_blank())+
  annotate("rect", xmin = 0, xmax = 6.5, ymin = 0.25, ymax = 0.5, 
           alpha=1, fill="#FDE725FF") +
  annotate("rect", xmin = 6.5, xmax = 25.5, ymin = 0.25, ymax = 0.5, 
           alpha=1, fill="#472D7BFF") +
  annotate("rect", xmin = 25.5, xmax = 33, ymin = 0.25, ymax = 0.5, 
           alpha=1, fill="#21908CFF") 

bubbleplot_alldeaf1
```

```
#export 1000x300
#export 1000x500 for legend
```

#### 4.3 Find genes that are dysregulated in c207y but not in the other mutants

```
#add a column that is 1 if pvalue is <0.05 and +FC, -1 if padj is <0.05 and -FC
deaf1DEGdata2 <- deaf1DEGdata %>% mutate(Venn = ifelse(pvalue < 0.01 & log2FoldChange > 0, 1, ifelse(pvalue < 0.01 & log2FoldChange < 0, 1, 0)))

#filter to only include homvwt comparisons
deaf1DEGdata2 <- deaf1DEGdata2 %>% dplyr::filter(comparison %in% c("d23bp_homvwt", "dc207y_homvwt", "dt238p_homvwt", "d23bp_homvwt_6dpf"))

#make a new table with only gene name, comparison name, and Venn column
venntable <- deaf1DEGdata2 %>% dplyr::select(LLgeneAbbrev, comparison, Venn)

#find duplicates in venntable
duplicategenes <- dplyr::filter(venntable %>%
  distinct() %>%
  group_by(LLgeneAbbrev, comparison) %>%
  dplyr::count(), n !=1)$LLgeneAbbrev

#remove duplicate rows from venntable
venntable <- venntable %>% dplyr::filter(!LLgeneAbbrev %in% duplicategenes)

#use pivot wider to make untidy table
venntable <- as.matrix(venntable %>% distinct() %>%
    pivot_wider(names_from = comparison, values_from  = Venn, values_fill=0) %>%
    column_to_rownames('LLgeneAbbrev'))

#convert NULL to 0
venntable[venntable=='NULL'] <- 0

#find sum of values of rows
rowsum <- rowSums(venntable)
venntable <- cbind(venntable, rowsum)

#convert to dataframe
venntable <- as.data.frame(venntable)

#sort by those with highest sum of absolute values of rows
venntable <- venntable %>% arrange(desc(abs(rowsum)))

#filter by those with ==1 sig gene
venntable <- venntable %>% dplyr::filter(rowsum == 1)

#filter by those where c207 is 1
venntable <- venntable %>% dplyr::filter(dc207y_homvwt == 1)

#filter by those not on chr25
venntable_nochr25 <- setdiff(rownames(venntable), chr25genes)

#get all data for list of genes
c207list <- deaf1DEGdata %>% dplyr::filter(comparison == "dc207y_homvwt" & LLgeneAbbrev %in% venntable_nochr25)

#sort by p value
c207list <- c207list %>% arrange(pvalue)

c207list$LLgeneAbbrev
```

```
##  [1] "e2f4"              "shisal1b"          "rcvrnb"           
##  [4] "hps4"              "prox1b"            "abtb2b"           
##  [7] "zgc:153219"        "LOC110440168"      "brpf1"            
## [10] "dop1a"             "znf1089"           "urp2"             
## [13] "acmsd"             "col28a1a"          "rhcga"            
## [16] "vip"               "fah"               "s1pr3a"           
## [19] "ttll6"             "kcnv2b"            "phkg2"            
## [22] "si:ch211-219a15.3" "hhat"              "CABZ01114105.1"   
## [25] "XLOC_003689"       "kmt5ab"            "slco1e1"          
## [28] "cldn20"            "creb1a"            "wnt11f2"          
## [31] "XLOC_028644"       "FQ311928.1"        "armc4"            
## [34] "cacna2d3"          "adcy7"             "CU467646.6"       
## [37] "si:dkey-93h22.8"   "efna5a"            "inpp5d"           
## [40] "zgc:103564"        "fbxl3b"            "spsb4b"           
## [43] "taco1"             "fam20cb"           "LOC101883486"     
## [46] "fam219ab"          "XLOC_039632"       "crygm2e"          
## [49] "zswim6"
```

#### 4.4 Make bubble plot of genes specific to c207y

```
#comparisons to include in plot
comparisons <- c("dt238p_homvwt", "dc207y_homvwt", "d23bp_homvwt_6dpf", "d23bp_homvwt")

c207specificgenes <- c(sort(c("rcvrnb",  "prox1b",  "brpf1", "dop1a",  "efna5a", "fbxl3b", "spsb4b")), sort(c("hps4", "zgc:153219", "LOC110440168","znf1089", "urp2", "vip", "fah", "s1pr3a", "ttll6", "slco1e1", "armc4", "cacna2d3")))

#filter to only include genes of interest
bubbleDEG <- deaf1DEGdata %>% dplyr::filter(LLgeneAbbrev %in% c207specificgenes & comparison %in% comparisons)

#find middle of foldchange (to center color scale)
limit <- max(abs(bubbleDEG$log2FoldChange)) * c(-1, 1)

#order genes
bubbleDEG$LLgeneAbbrev <- factor(bubbleDEG$LLgeneAbbrev, levels = c207specificgenes)

#order comparisons
bubbleDEG$comparison <- factor(bubbleDEG$comparison, levels = c("d23bp_homvwt_6dpf", "d23bp_homvwt", "dt238p_homvwt", "dc207y_homvwt"))

#rename comparisons
levels(bubbleDEG$comparison) <- c("23d46i 6dpf", "23d46i 2dpf", "t238p 2dpf", "c207y 2dpf")

# create 'bubble plot' to summarize y signatures across x phenotypes
bubbleplot_c207y <- ggplot(bubbleDEG, aes(x=LLgeneAbbrev, y=comparison)) + 
  geom_point(aes(size=-log10(padj), color = log2FoldChange)) +
  scale_color_gradientn(colors = myheatcolors3, limit=limit) +
  theme_bw() +
  theme(axis.text.x = element_text(angle = 45, hjust = 1),
        axis.title.x = element_blank(),
        axis.title.y = element_blank())+
  annotate("rect", xmin = 0, xmax = 7.5, ymin = 0.25, ymax = 0.5, 
           alpha=1, fill="#472D7BFF") +
  annotate("rect", xmin = 7.5, xmax = 20, ymin = 0.25, ymax = 0.5, 
           alpha=1, fill="#21908CFF") 

bubbleplot_c207y
```

```
#export 700x300
```

#### 4.5 Find genes that are dysregulated in 2dpf 23bp but not in 6dpf 23bp

```
#add a column that is 1 if pvalue is <0.05 and +FC, -1 if padj is <0.05 and -FC
deaf1DEGdata2 <- deaf1DEGdata %>% mutate(Venn = ifelse(pvalue < 0.01 & log2FoldChange > 0, 1, ifelse(pvalue < 0.01 & log2FoldChange < 0, 1, 0)))

#filter to only include homvwt comparisons
deaf1DEGdata2 <- deaf1DEGdata2 %>% dplyr::filter(comparison %in% c("d23bp_homvwt", "d23bp_homvwt_6dpf", "dc207y_homvwt", "dt238p_homvwt"))

#make a new table with only gene name, comparison name, and Venn column
venntable <- deaf1DEGdata2 %>% dplyr::select(LLgeneAbbrev, comparison, Venn)

#find duplicates in venntable
duplicategenes <- dplyr::filter(venntable %>%
  distinct() %>%
  group_by(LLgeneAbbrev, comparison) %>%
  dplyr::count(), n !=1)$LLgeneAbbrev

#remove duplicate rows from venntable
venntable <- venntable %>% dplyr::filter(!LLgeneAbbrev %in% duplicategenes)

#use pivot wider to make untidy table
venntable <- as.matrix(venntable %>% distinct() %>%
    pivot_wider(names_from = comparison, values_from  = Venn, values_fill=0) %>%
    column_to_rownames('LLgeneAbbrev'))

#convert NULL to 0
venntable[venntable=='NULL'] <- 0

#find sum of values of rows
rowsum <- rowSums(venntable)
venntable <- cbind(venntable, rowsum)

#convert to dataframe
venntable <- as.data.frame(venntable)

#sort by those with highest sum of absolute values of rows
venntable <- venntable %>% arrange(desc(abs(rowsum)))

#filter by those with >1 significant gene
venntable <- venntable %>% dplyr::filter(rowsum >= 2)

#filter by those where 6dpf isn't different
venntable <- venntable %>% dplyr::filter(d23bp_homvwt_6dpf == 0)

#filter by those not on chr25
venntable_nochr25 <- setdiff(rownames(venntable), chr25genes)

#get all data for 6dpf genes
age23bp <- deaf1DEGdata %>% dplyr::filter(comparison == "d23bp_homvwt" & LLgeneAbbrev %in% venntable_nochr25)

#sort by pvalue
age23bp <- age23bp %>% arrange(pvalue)

age23bp$LLgeneAbbrev
```

```
##  [1] "CABZ01039863.1"   "cav1"             "mkrn1"            "evlb"            
##  [5] "hdac10"           "CABZ01039859.1"   "apc2"             "adarb2"          
##  [9] "slc25a22a"        "vldlr"            "col12a1a"         "rapgef1a"        
## [13] "dkk3b"            "chordc1b"         "draxin"           "igf2bp2a"        
## [17] "dusp6"            "zgc:158689"       "snx27a"           "NC_002333.17"    
## [21] "schip1"           "samd11"           "si:ch211-102c2.7" "tgfbr1b"         
## [25] "rab3ab"           "prkx"             "pank1b"           "pdgfra"          
## [29] "elovl7b"          "aff4"             "casp8ap2"         "zgc:162255"      
## [33] "si:ch211-276c2.4" "pbx4"             "si:ch211-239f4.1" "elovl4a"         
## [37] "CACFD1"           "SIPA1"            "ganab"            "rac3a"           
## [41] "CC2D1A"           "elovl2"           "LOC108182329"     "hipk2"           
## [45] "ano9b"            "LOC110438587"     "pde5ab"           "abi3b"           
## [49] "CABZ01092170.1"   "gas7a"            "ctcf"             "tnfaip8l2b"      
## [53] "fbln2"            "kdm6bb"           "rab11fip4a"       "zgc:110425"      
## [57] "map6b"            "CABZ01004876.1"   "tfe3a"            "msrb1a"          
## [61] "mapk3"            "ube2al"           "rsf1a"            "vac14"           
## [65] "polr3b"           "ldb2a"            "LOC100537384"     "rnasekb"         
## [69] "hs3st4"           "cenpf"            "gdpd4a"           "add1"            
## [73] "sacm1la"          "exoc5"            "gjd1a"            "rnf126"          
## [77] "kif5c"            "kitlga"           "CABZ01029822.1"   "myof"            
## [81] "pax10"            "prdm1b"           "pdia3"            "itpr1a"          
## [85] "si:dkeyp-120h9.1" "impa2"            "deaf1"
```

#### 4.6 Make bubble plot of genes that are dysregulated at 2dpf but not as 6dpf

```
#comparisons to include in plot
comparisons <- c("dt238p_homvwt", "dc207y_homvwt", "d23bp_homvwt_6dpf", "d23bp_homvwt")

#make list of age-specific genes to include
agespecificgenes <- c(sort(c("CABZ01039859.1", "CABZ01092170.1", "evlb", "hdac10", "pdia3", "polr3b")), sort(c("CABZ01039863.1", "chordc1b", "dkk3b", "draxin", "dusp6", "elovl2", "exoc5", "itpr1a", "mkrn1", "msrb1a", 'pbx4', "prkx", "rapgef1a", "ube2al")))

#filter to only include genes of interest
bubbleDEG <- deaf1DEGdata %>% dplyr::filter(LLgeneAbbrev %in% agespecificgenes & comparison %in% comparisons)

#find middle of foldchange (to center color scale)
limit <- max(abs(bubbleDEG$log2FoldChange)) * c(-1, 1)

#order genes
bubbleDEG$LLgeneAbbrev <- factor(bubbleDEG$LLgeneAbbrev, levels = agespecificgenes)

#order comparisons
bubbleDEG$comparison <- factor(bubbleDEG$comparison, levels = c("d23bp_homvwt_6dpf", "d23bp_homvwt", "dt238p_homvwt", "dc207y_homvwt"))

#rename comparisons
levels(bubbleDEG$comparison) <- c("23d46i 6dpf", "23d46i 2dpf", "t238p 2dpf", "c207y 2dpf")

# create 'bubble plot' to summarize y signatures across x phenotypes
bubbleplot_ages <- ggplot(bubbleDEG, aes(x=LLgeneAbbrev, y=comparison)) + 
  geom_point(aes(size=-log10(padj), color = log2FoldChange)) +
  scale_color_gradientn(colors = myheatcolors3, limit=limit) +
  theme_bw() +
  theme(axis.text.x = element_text(angle = 45, hjust = 1),
        axis.title.x = element_blank(),
        axis.title.y = element_blank())+
  annotate("rect", xmin = 0, xmax = 6.5, ymin = 0.25, ymax = 0.5, 
           alpha=1, fill="#472D7BFF") +
  annotate("rect", xmin = 6.5, xmax = 21, ymin = 0.25, ymax = 0.5, 
           alpha=1, fill="#21908CFF") 


bubbleplot_ages
```

```
#export 700x300
```

### 5 Gene Set Enrichment Analysis

#### 5.1 Bubble plot of eye pathways across 2 dpf mutants

```
#we want the zebrafish signatures
dr_gsea <- msigdbr(species = "Danio rerio") #gets all collections/signatures with zebrafish

#look at the categories and subcategories of signatures available
dr_gsea %>% 
  dplyr::distinct(gs_cat, gs_subcat) %>% 
  dplyr::arrange(gs_cat, gs_subcat)
```

```
# choose a specific msigdb collection/subcollection
dr_gsea_c5 <- msigdbr(species = "Danio rerio", category = "C5") %>% # choose  msigdb collection of interest
  dplyr::select(gs_name, gene_symbol) #just get the columns corresponding to signature name and gene symbols of genes in each signature 

# pull out data for dc207y
GSEAgenes <- deaf1DEGdata %>% dplyr::filter(comparison == "dc207y_homvwt")
GSEAgenes <- GSEAgenes %>% dplyr::filter(! LLgeneAbbrev %in% chr25genes)
mydata.df.sub <- dplyr::select(GSEAgenes, LLgeneAbbrev, log2FoldChange)

#get rid of duplicates
mydata.df.sub <- mydata.df.sub %>% dplyr::filter(LLgeneAbbrev %in% mydata.df.sub$LLgeneAbbrev[!duplicated(mydata.df.sub$LLgeneAbbrev)])

# construct a named vector
mydata.gsea <- mydata.df.sub$log2FoldChange
names(mydata.gsea) <- as.character(mydata.df.sub$LLgeneAbbrev)
mydata.gsea <- sort(mydata.gsea, decreasing = TRUE)

# run GSEA with C5 in c207y
myGSEA.res.c207y <- GSEA(mydata.gsea, TERM2GENE=dr_gsea_c5, verbose=FALSE, seed=TRUE, pvalueCutoff = 1)

# pull out data for 23bp
GSEAgenes <- deaf1DEGdata %>% dplyr::filter(comparison == "d23bp_homvwt")
GSEAgenes <- GSEAgenes %>% dplyr::filter(! LLgeneAbbrev %in% chr25genes)
mydata.df.sub <- dplyr::select(GSEAgenes, LLgeneAbbrev, log2FoldChange)

#get rid of duplicates
mydata.df.sub <- mydata.df.sub %>% dplyr::filter(LLgeneAbbrev %in% mydata.df.sub$LLgeneAbbrev[!duplicated(mydata.df.sub$LLgeneAbbrev)])

# construct a named vector
mydata.gsea <- mydata.df.sub$log2FoldChange
names(mydata.gsea) <- as.character(mydata.df.sub$LLgeneAbbrev)
mydata.gsea <- sort(mydata.gsea, decreasing = TRUE)

# run GSEA with C5 in 23bp
myGSEA.res.23bp <- GSEA(mydata.gsea, TERM2GENE=dr_gsea_c5, verbose=FALSE, seed=TRUE, pvalueCutoff = 1)

# pull out data for t238p
GSEAgenes <- deaf1DEGdata %>% dplyr::filter(comparison == "dt238p_homvwt")
GSEAgenes <- GSEAgenes %>% dplyr::filter(! LLgeneAbbrev %in% chr25genes)
mydata.df.sub <- dplyr::select(GSEAgenes, LLgeneAbbrev, log2FoldChange)

#get rid of duplicates
mydata.df.sub <- mydata.df.sub %>% dplyr::filter(LLgeneAbbrev %in% mydata.df.sub$LLgeneAbbrev[!duplicated(mydata.df.sub$LLgeneAbbrev)])

# construct a named vector
mydata.gsea <- mydata.df.sub$log2FoldChange
names(mydata.gsea) <- as.character(mydata.df.sub$LLgeneAbbrev)
mydata.gsea <- sort(mydata.gsea, decreasing = TRUE)

# run GSEA with C5 in t238p
myGSEA.res.t238p <- GSEA(mydata.gsea, TERM2GENE=dr_gsea_c5, verbose=FALSE, seed=TRUE)

#convert to data frame add column about mutations
myGSEA.df.t238p <- as_tibble(myGSEA.res.t238p@result)
myGSEA.df.c207y <- as_tibble(myGSEA.res.c207y@result)
myGSEA.df.23bp <- as_tibble(myGSEA.res.23bp@result)
myGSEA.df.t238p$mutation <- "t238p"
myGSEA.df.c207y$mutation <- "c207y"
myGSEA.df.23bp$mutation <- "23d46i"

#combine the GSEA data into one data frame
myGSEA2dpf <- rbind(myGSEA.df.t238p, myGSEA.df.c207y, myGSEA.df.23bp)

#export myGSEA2dpf
#write.csv(myGSEA2dpf, file="deaf1_2dpf_C5_GSEA.csv")

#import GSEA
myGSEA2dpf <- read.csv(file="deaf1_2dpf_C5_GSEA.csv")[,2:13]

#look at top terms
datatable(myGSEA2dpf, 
          extensions = c('KeyTable', "FixedHeader"), 
          options = list(keys = TRUE, searchHighlight = TRUE, pageLength = 10, lengthMenu = c("10", "25", "50", "100"))) %>%
  formatRound(columns=c(2:10), digits=2)
```

```
#list of eye-related terms
eyeterms <- c("GOBP_DETECTION_OF_LIGHT_STIMULUS", "HP_COLOR_VISION_DEFECT", "GOCC_PHOTORECEPTOR_INNER_SEGMENT", "HP_UNDETECTABLE_ELECTRORETINOGRAM", "GOBP_PHOTORECEPTOR_CELL_DIFFERENTIATION", "GOBP_RETINA_DEVELOPMENT_IN_CAMERA_TYPE_EYE", "GOBP_SENSORY_PERCEPTION_OF_LIGHT_STIMULUS", "GOBP_PHOTORECEPTOR_CELL_DEVELOPMENT", "HP_ABNORMAL_VISUAL_ELECTROPHYSIOLOGY", "GOBP_RETINA_MORPHOGENESIS_IN_CAMERA_TYPE_EYE", "GOBP_DETECTION_OF_LIGHT_STIMULUS_INVOLVED_IN_SENSORY_PERCEPTION", "HP_COLOR_VISION_DEFECT")

#filter to include only eye terms
myGSEA2dpf <- myGSEA2dpf %>% dplyr::filter(ID %in% eyeterms)

# create bubble plot of shared pathways
eyetermsbubble <- ggplot(myGSEA2dpf, aes(x=mutation, y=ID)) + 
  geom_point(aes(size=setSize, color = NES, alpha=-log10(p.adjust))) +
  scale_color_gradientn(colors = myheatcolors3, limit=c(-3,3)) +
  theme_bw() + 
  theme(axis.title.x=element_blank(), axis.title.y=element_blank())

eyetermsbubble
```

```
#export 900x500

#make plot of c207y visual 
gseaplot2(myGSEA.res.c207y, 
          geneSetID = c(91, 1640, 673), #can choose multiple signatures to overlay in this plot
         pvalue_table = FALSE)
```

```
#export 800x500

#make plot of t238p visual 
gseaplot2(myGSEA.res.t238p, 
          geneSetID = c(19, 20, 15), #can choose multiple signatures to overlay in this plot
         pvalue_table = FALSE)
```

#### 5.2 6 dpf GSEA C5

```
# pull out data for 23bp 6dpf
GSEAgenes <- deaf1DEGdata %>% dplyr::filter(comparison == "d23bp_homvwt_6dpf")
GSEAgenes <- GSEAgenes %>% dplyr::filter(! LLgeneAbbrev %in% chr25genes)
mydata.df.sub <- dplyr::select(GSEAgenes, LLgeneAbbrev, log2FoldChange)

#get rid of duplicates
mydata.df.sub <- mydata.df.sub %>% dplyr::filter(LLgeneAbbrev %in% mydata.df.sub$LLgeneAbbrev[!duplicated(mydata.df.sub$LLgeneAbbrev)])

# construct a named vector
mydata.gsea <- mydata.df.sub$log2FoldChange
names(mydata.gsea) <- as.character(mydata.df.sub$LLgeneAbbrev)
mydata.gsea <- sort(mydata.gsea, decreasing = TRUE)

# run GSEA with C5 in 23bp 6dpf
myGSEA.res.23bp6dpf <- GSEA(mydata.gsea, TERM2GENE=dr_gsea_c5, verbose=FALSE, seed=TRUE, pvalueCutoff = 1)

#convert to DF
myGSEA.df.23bp.6dpf <- as_tibble(myGSEA.res.23bp6dpf@result)

#export
#write.csv(myGSEA.res.23bp6dpf, file="GSEA6dpf.csv")

#import
myGSEA.df.23bp.6dpf <- read.csv("GSEA6dpf.csv")[,2:12]

#look at top terms
datatable(myGSEA.df.23bp.6dpf, 
          extensions = c('KeyTable', "FixedHeader"), 
          options = list(keys = TRUE, searchHighlight = TRUE, pageLength = 10, lengthMenu = c("10", "25", "50", "100"))) %>%
  formatRound(columns=c(2:10), digits=2)
```

```
#make list of terms to include in heat map
C5GSEAterms <- c("GOCC_NEURON_TO_NEURON_SYNAPSE", "GOCC_GLUTAMATERGIC_SYNAPSE", "GOCC_POSTSYNAPTIC_MEMBRANE", "GOCC_SYNAPTIC_MEMBRANE", "GOBP_POSITIVE_REGULATION_OF_INTERLEUKIN_1_PRODUCTION", "GOBP_RESPONSE_TO_CHEMOKINE", "GOBP_POSITIVE_REGULATION_OF_CYTOKINE_PRODUCTION")

#make heatmap
p3 <- heatplot(myGSEA.res.23bp6dpf, foldChange=mydata.gsea) + ggplot2::coord_flip()

#get data of out heatmap
p3 <- ggplot_build(p3)
heatmapdata <- p3$plot$data

#filter to only include results for subtypes of interest
heatmapdata <- heatmapdata %>% dplyr::filter(categoryID %in% C5GSEAterms)

#filter to only include fold changes > 0.5
genestoinclude <- heatmapdata %>% dplyr::filter(foldChange > 0.75 | foldChange < -0.75)
heatmapdata <- heatmapdata %>% dplyr::filter(Gene %in% genestoinclude$Gene)

#use pivot wider to make untidy table
heatmapdata <- heatmapdata %>% pivot_wider(names_from = categoryID, values_from  = foldChange, values_fill=NA) 

#turn heatmap data back into tidy table
heatmapdata <- heatmapdata %>% pivot_longer(!Gene, names_to = "categoryID", values_to  = "foldChange") 
    
#change order of data to be plotted
heatmapdata$categoryID <- factor(heatmapdata$categoryID, levels = rev(c("GOCC_POSTSYNAPTIC_MEMBRANE", "GOCC_SYNAPTIC_MEMBRANE", "GOCC_NEURON_TO_NEURON_SYNAPSE", "GOCC_GLUTAMATERGIC_SYNAPSE", "GOBP_POSITIVE_REGULATION_OF_INTERLEUKIN_1_PRODUCTION", "GOBP_RESPONSE_TO_CHEMOKINE", "GOBP_POSITIVE_REGULATION_OF_CYTOKINE_PRODUCTION" )))

#rename dataframe columns
colnames(heatmapdata) <- c("Gene", "categoryID", "log2foldchange")

C5_6dpf_heatmap <- ggplot(heatmapdata, aes(x = Gene, y = categoryID, fill = log2foldchange)) + 
  geom_tile(color="black") + 
  theme_classic() + 
  scale_fill_continuous_divergingx(palette = 'RdBu', mid = 0, na.value="white") +
  theme(axis.text.x = element_text(angle = 45, vjust = 1, hjust = 1), axis.title.x=element_blank(), axis.title.y=element_blank()) 

C5_6dpf_heatmap
```

```
#export 1100x350
```

#### 5.3 Zebrafish anatomy ontology - make gmt file

This uses the zebrafish anatomy ontology and expression information
to make lists of genes that are expressed within each anatomy term.

```
#make a vector of the relationship types included in the ZFA obo file
relationshipnames <- get_relation_names("zfaontology.OBO.txt")

#read in OBO file
zfaontology <- get_ontology("zfaontology.OBO.txt",  propagate_relationships=relationshipnames, extract_tags = "everything")

#read in gaf file
zfagaf<-readr::read_tsv("zfagaf.gaf") #import gaf
zfagaf <- zfagaf[37:235090,c(2:6)] #remove the header and unnecessary columns
colnames(zfagaf) <- c("zfingeneid", "genename", "relationship", "zfa", "evidence")

#make empty list
genezfalist <- list()

#use a loop to find all the ZFA terms associated with each gene and put them in a list
for (x in 1:length(zfagaf$genename)) {
    #get the first gene
    rowdata <- zfagaf[x,]
    gene <- rowdata$genename #name of gene
    zfaterm <- rowdata$zfa #name of ZFA term associated with gene
    ancestors <- get_ancestors(zfaontology, zfaterm)  #all ancestors of term
    ancestorsgene <- as.data.frame(matrix(data=NA, nrow=length(ancestors), ncol=0)) #empty data frame
    ancestorsgene$genename <- gene #add gene name to data frame
    ancestorsgene$ZFAterm <- ancestors #add parents of ZFA term to data frame
    genezfalist[[x]] <- ancestorsgene #put data frame in a list
}

#combine all the data frames from list into one data frame
genezfaassociation <- do.call(rbind, genezfalist)

#remove duplicate rows
genezfaassociation <- genezfaassociation %>% distinct()

#export file
write.csv(genezfaassociation, file="genezfaassociation.csv")

#pivot wider to make ZFA terms a column and each gene a row
pivot <- genezfaassociation <- genezfaassociation %>% 
  pivot_wider(
    names_from = genename, 
    values_from = genename
  )

pivot2 <- pivot

#add name of zfa ID to data frame
zfaontologid <- as.data.frame(zfaontology$name) #get ZFA ontology/id pairs
zfaontologid$id <- rownames(zfaontologid) #make column called id containing ZFA id
pivot2$description <- NA #add a column for description containing NAs

#add descriptions for ZFA IDs
for (x in 1:length(pivot2$description)) {
  firstterm <- as.character(pivot2[x,1]) #get ZFA term out of pivot
  filteredzfa <- zfaontologid %>% dplyr::filter(id == firstterm)
  pivot2$description[x] <- filteredzfa$`zfaontology$name`
}

#rearrange table
pivot2 <- cbind(pivot2$ZFAterm, pivot2$description, pivot2[,2:14213])

#shift to the left
pivot3 = as.data.frame(t(apply(pivot2,1, function(x) { return(c(x[!is.na(x)],x[is.na(x)]) )} )))
colnames(pivot3) = colnames(pivot2)

#get rid of ZFA terms that only have one or two genes (zfa terms must have three genes)
pivot4 <- pivot3
pivot4 <- pivot4 %>% drop_na(tbxta)
pivot4 <- pivot4 %>% drop_na(bglap)
pivot4[is.na(pivot4)] <- ""
colnames(pivot4)[1] <- "zfaterm"

#export gmt with at least 3 genes/zfa term
write.table(pivot4, file='zfa-geneexp-min.gmt', quote=FALSE, sep='\t', row.names =FALSE)

#pivot longer to make tidy for GSEA
pivot5 <- pivot4[,2:14214]
colnames(pivot5)[1] <- "zfaterm"
pivot5 <- pivot5 %>% 
  pivot_longer(
  !zfaterm,
  names_to = "extracolumn",
  values_to = "genename"
  )
pivot5 <- pivot5[,c(1,3)]
fullsetGSEA <- pivot5[!(is.na(pivot5$genename) | pivot5$genename==""), ]

#filter to only include descendants of head
headterms <- get_descendants(zfaontology, "ZFA:0001114") #get descendants of head
pivot5 <- pivot4 %>% dplyr::filter (zfaterm %in% headterms)
pivot5 <- pivot5[,2:14214]
colnames(pivot5)[1] <- "zfaterm"
pivot5 <- pivot5 %>% 
  pivot_longer(
  !zfaterm,
  names_to = "extracolumn",
  values_to = "genename"
  )
pivot5 <- pivot5[,c(1,3)]
headtermsGSEA <- pivot5[!(is.na(pivot5$genename) | pivot5$genename==""), ]

#get descendants of CNS
CNSterms <- get_descendants(zfaontology, "ZFA:0000012")
pivot5 <- pivot4 %>% dplyr::filter (zfaterm %in% CNSterms)
pivot5 <- pivot5[,2:14214]
colnames(pivot5)[1] <- "zfaterm"
pivot5 <- pivot5 %>% 
  pivot_longer(
  !zfaterm,
  names_to = "extracolumn",
  values_to = "genename"
  )
pivot5 <- pivot5[,c(1,3)]
CNStermsGSEA <- pivot5[!(is.na(pivot5$genename) | pivot5$genename==""), ]

#change NA to empty string for full gmt
pivot3[is.na(pivot3)] <- ""

#export full gmt
write.table(pivot3, file='zfa-geneexp.gmt', quote=FALSE, sep='\t', row.names =FALSE)

#token = upload_GMT_file(gmtfile = "zfa-geneexp-min.gmt")
#the token to use the gmt is "gp__3Toj_Qd5k_dG4"
```

#### 5.4 Use ZFA anatomy to perform GSEA on 6dpf 23bp DEGs

```
# Pull out just the columns corresponding to gene symbols and LogFC for at least one pairwise comparison for the enrichment analysis
GSEAgenes <- deaf1DEGdata %>% dplyr::filter(comparison == "d23bp_homvwt_6dpf")
mydata.df.sub <- dplyr::select(GSEAgenes, LLgeneAbbrev, log2FoldChange)

#get rid of duplicates
mydata.df.sub <- mydata.df.sub %>% dplyr::filter(LLgeneAbbrev %in% mydata.df.sub$LLgeneAbbrev[!duplicated(mydata.df.sub$LLgeneAbbrev)])

# construct a named vector
mydata.gsea <- mydata.df.sub$log2FoldChange
names(mydata.gsea) <- as.character(mydata.df.sub$LLgeneAbbrev)
mydata.gsea <- sort(mydata.gsea, decreasing = TRUE)

#options for doing GSEA using ZFA
#CNStermsGSEA
#headtermsGSEA
#fullsetGSEA

# run GSEA with CNS terms in 23bp
myGSEA.res <- GSEA(mydata.gsea, TERM2GENE=CNStermsGSEA, verbose=FALSE, seed=TRUE, minGSSize = 80)
myGSEA.df <- as_tibble(myGSEA.res@result)

#export file
#write.csv(myGSEA.df, file="functionalanalysis/6dpf_CNSterms_GSEA.csv")

#import file
myGSEA.df <- read.csv(file="functionalanalysis/6dpf_CNSterms_GSEA.csv")[,2:12]

# view results as an interactive table
datatable(myGSEA.df, 
          extensions = c('KeyTable', "FixedHeader"), 
          options = list(keys = TRUE, searchHighlight = TRUE, pageLength = 10, lengthMenu = c("10", "25", "50", "100"))) %>%
  formatRound(columns=c(2:10), digits=2)
```

```
#make network plot
myGSEA.res <- pairwise_termsim(myGSEA.res)
deaf1network <- emapplot(myGSEA.res, color="NES", categorySize="p.adjust")

#get data of out network plot
deaf1network <- ggplot_build(deaf1network)
networkdata <- deaf1network$plot$data

deaf1CNSterms_network <- ggraph(networkdata) + 
  geom_edge_link(alpha=.8, aes_(width=~I(width)), colour='darkgrey') + 
  geom_node_point(aes(colour = color, size=size)) +
  geom_node_text(aes(label=name), repel=TRUE) + 
  theme_void() +
  scale_color_gradientn(colors = myheatcolors3, limit=c(-2.5,2.5), name="NES")

deaf1CNSterms_network
```

```
#export 700x500

#make heatmap
p3 <- heatplot(myGSEA.res, foldChange=mydata.gsea) + ggplot2::coord_flip()

#get data of out heatmap
p3 <- ggplot_build(p3)
heatmapdata <- p3$plot$data

#filter to only include fold changes > 0.5
genestoinclude <- heatmapdata %>% dplyr::filter(foldChange > 0.5 | foldChange < -0.5)
heatmapdata <- heatmapdata %>% dplyr::filter(Gene %in% genestoinclude$Gene)

#use pivot wider to make untidy table
heatmapdata <- heatmapdata %>% pivot_wider(names_from = categoryID, values_from  = foldChange, values_fill=NA) 

#make matrix
heatmapmatrix <- as.matrix(heatmapdata[,2:12])
rownames(heatmapmatrix) <- heatmapdata$Gene

#turn matrix into 1s and 0s and cluster
heatmapmatrix<-ifelse(abs(heatmapmatrix) > 0,1,0)
heatmapmatrix[is.na(heatmapmatrix)] = 0
 #cluster rows by pearson correlation
hc <- hclust(as.dist(1-cor(heatmapmatrix, method="spearman")), method="average") #cluster columns by spearman correlation

#turn heatmap data back into tidy table
heatmapdata <- heatmapdata %>% pivot_longer(!Gene, names_to = "categoryID", values_to  = "foldChange") 
    
#change order of data to be plotted
heatmapdata$categoryID <- factor(heatmapdata$categoryID, levels = hc$labels)

#rename dataframe columns
colnames(heatmapdata) <- c("Gene", "categoryID", "log2foldchange")

zfaCNS_heatmap <- ggplot(heatmapdata, aes(x = Gene, y = categoryID, fill = log2foldchange)) + 
  geom_tile(color="black") + 
  theme_classic() + 
  scale_fill_continuous_divergingx(palette = 'RdBu', mid = 0, na.value="white") +
  theme(axis.text.x = element_text(angle = 45, vjust = 1, hjust = 1), axis.title.x=element_blank(), axis.title.y=element_blank()) 

zfaCNS_heatmap
```

```
#export 1000x400


# run GSEA with head terms in 23bp
myGSEA.res <- GSEA(mydata.gsea, TERM2GENE=headtermsGSEA, verbose=FALSE, seed=TRUE, minGSSize = 80)
myGSEA.df <- as_tibble(myGSEA.res@result)

# view results as an interactive table
datatable(myGSEA.df, 
          extensions = c('KeyTable', "FixedHeader"), 
          options = list(keys = TRUE, searchHighlight = TRUE, pageLength = 10, lengthMenu = c("10", "25", "50", "100"))) %>%
  formatRound(columns=c(2:10), digits=2)
```

```
#make network plot
myGSEA.res <- pairwise_termsim(myGSEA.res)
deaf1network <- emapplot(myGSEA.res, color="NES", categorySize="p.adjust")

#get data of out heatmap
deaf1network <- ggplot_build(deaf1network)
networkdata <- deaf1network$plot$data

deaf1headterms_network <- ggraph(networkdata) + 
  geom_edge_link(alpha=.8, aes_(width=~I(width)), colour='darkgrey') + 
  geom_node_point(aes(colour = color, size=size)) +
  geom_node_text(aes(label=name), repel=TRUE) + 
  theme_void() +
  scale_color_gradientn(colors = myheatcolors3, limit=c(-2.5,2.5), name = "NES")

deaf1headterms_network
```

```
#export 700x500
```

#### 5.5 Use single cell markers from 5 dpf zebrafish brain to perform GSEA on 6dpf 23bp insertion

```
#import single cell markers
singlecellmarkers <- read.csv("zf5dpf_markersforGSEA.csv")

#select relevant columns
singlecellmarkers <- singlecellmarkers %>% dplyr::select(cluster.description, gene)

# Pull out just the columns corresponding to gene symbols and LogFC
GSEAgenes <- deaf1DEGdata %>% dplyr::filter(comparison == "d23bp_homvwt_6dpf")
mydata.df.sub <- dplyr::select(GSEAgenes, LLgeneAbbrev, log2FoldChange)

#get rid of duplicates
mydata.df.sub <- mydata.df.sub %>% dplyr::filter(LLgeneAbbrev %in% mydata.df.sub$LLgeneAbbrev[!duplicated(mydata.df.sub$LLgeneAbbrev)])

# construct a named vector
mydata.gsea <- mydata.df.sub$log2FoldChange
names(mydata.gsea) <- as.character(mydata.df.sub$LLgeneAbbrev)
mydata.gsea <- sort(mydata.gsea, decreasing = TRUE)

# run GSEA with single cell markers in 6dpf 23bp
myGSEA.res <- GSEA(mydata.gsea, TERM2GENE=singlecellmarkers, verbose=FALSE, seed=TRUE, minGSSize = 80)
myGSEA.df <- as_tibble(myGSEA.res@result)

#export
#write.csv(myGSEA.df, file="deaf1-singlecellGSEA.csv")

#import
myGSEA.df <- read.csv(file="deaf1-singlecellGSEA.csv")[,2:12]

datatable(myGSEA.df, 
          extensions = c('KeyTable', "FixedHeader"), 
          options = list(keys = TRUE, searchHighlight = TRUE, pageLength = 10, lengthMenu = c("10", "25", "50", "100"))) %>%
  formatRound(columns=c(2:10), digits=2)
```

```
#make network plot
myGSEA.res <- pairwise_termsim(myGSEA.res)
deaf1network <- emapplot(myGSEA.res, color="NES", categorySize="p.adjust", showCategory = 57)

#get data of out network plot
deaf1network <- ggplot_build(deaf1network)
networkdata <- deaf1network$plot$data

#make network plot
deaf1singlecell_network <- ggraph(networkdata) + 
  geom_edge_link(alpha=.8, aes_(width=~I(width)), colour='darkgrey') + 
  geom_node_point(aes(colour = color, size=size)) +
  geom_node_text(aes(label=name), repel=TRUE) + 
  theme_void() +
  scale_color_gradientn(colors = myheatcolors3, limit=c(-3.5,3.5), name="NES")

deaf1singlecell_network
```

```
#export 700x500

#repeat 6dpf GSEA but with neuronal clusters only
nonneuronalclusters <- c("#N/A", "cartilage #2", "cartilage #1", "cartilage #3", "cornea", "cranial ganglion", "cranial ganglion (vagal) #1", "cranial ganglion (vagal) #2", "dermal bone", "epidermis (progenitors)", "epithelium (cornea, progenitors)", "erythrocytes", "eye (cornea), cartilage", "eye (cornea, epithelium)", "eye (non retina)", "glia (eye)", "lens #1", "lens #2", "muscle", "neural crest derived (pigment/xanthophore)", "neutrophils", "otic vesicle", "otic/lateral line", "pigment cell (iridophore)", "retina (amacrine, gabaergic)", "retina (cone bipolar cells)", "retina (cones)", "retina (muller glia)", "retina (photoreceptor precursor cells)", "retina (RGC)", "retina (rods)", "retina (RPE differentiation)", "retina (RPE)", "retina neurons (horizontal?)", "rostral blood island (myeloid) #1", "rostral blood island (myeloid) #2", "pharangeal arch/pectoral fin (mesoderm)")

#make GSEA set for CNS single cell data
singlecellmarkers_CNS <- singlecellmarkers %>% dplyr::filter(! cluster.description %in% nonneuronalclusters)

# run GSEA with single cell markers in 6dpf 23bp
myGSEA.res <- GSEA(mydata.gsea, TERM2GENE=singlecellmarkers_CNS, verbose=FALSE, seed=TRUE, minGSSize = 80)
myGSEA.df <- as_tibble(myGSEA.res@result)

#export
#write.csv(myGSEA.df, file="deaf1-singlecellGSEA-CNS.csv")

#import
myGSEA.df <- read.csv(file="deaf1-singlecellGSEA-CNS.csv")[,2:12]

#view data table
datatable(myGSEA.df, 
          extensions = c('KeyTable', "FixedHeader"), 
          options = list(keys = TRUE, searchHighlight = TRUE, pageLength = 10, lengthMenu = c("10", "25", "50", "100"))) %>%
  formatRound(columns=c(2:10), digits=2)
```

```
#make network plot
myGSEA.res <- pairwise_termsim(myGSEA.res)
deaf1network <- emapplot(myGSEA.res, color="NES", categorySize="p.adjust", showCategory = 33)

#get data of out network plot
deaf1network <- ggplot_build(deaf1network)
networkdata <- deaf1network$plot$data

#make new network plot
deaf1singlecellCNS_network <- ggraph(networkdata) + 
  geom_edge_link(alpha=.8, aes_(width=~I(width)), colour='darkgrey') + 
  geom_node_point(aes(colour = color, size=size)) +
  geom_node_text(aes(label=name), repel=TRUE) + 
  theme_void() +
  scale_color_gradientn(colors = myheatcolors3, limit=c(-3,3), name="NES")

deaf1singlecellCNS_network
```

```
#export 700x500

#list of neuronal subtypes to graph
neuronalsubtypes <- c("dorsal diencephalon (thalamus) #1", "dorsal diencephalon (thalamus) #2", "diencephalon (tuberculum; gabaergic)", "optic tectum (gabaergic) #1", "mid-hind boundary (gabaergic) #1", "dorsal diencephalon (thalamus) #3", "mid-hind boundary (gabaergic) #2", "dorsal diencephalon (thalamus) #4", "diencephalon", "neurons (dienc, glutamatergic, ZLI)", "optic tectum (gabaergic) #2", "midbrain #2", "hypothalamus #1", "hypothalamus #5","neurons (midbrain?, glutamatergic)", "hypothalamus #6")

#make heatmap
p3 <- heatplot(myGSEA.res, foldChange=mydata.gsea) + ggplot2::coord_flip()

#get data of out heatmap
p3 <- ggplot_build(p3)
heatmapdata <- p3$plot$data

#filter to only include results for subtypes of interest
heatmapdata <- heatmapdata %>% dplyr::filter(categoryID %in% neuronalsubtypes)

#filter to only include fold changes > 0.5
genestoinclude <- heatmapdata %>% dplyr::filter(foldChange > 0.5 | foldChange < -0.5)
heatmapdata <- heatmapdata %>% dplyr::filter(Gene %in% genestoinclude$Gene)

#use pivot wider to make untidy table
heatmapdata <- heatmapdata %>% pivot_wider(names_from = categoryID, values_from  = foldChange, values_fill=NA) 

#make matrix
heatmapmatrix <- as.matrix(heatmapdata[,2:15])
rownames(heatmapmatrix) <- heatmapdata$Gene

#turn matrix into 1s and 0s and cluster
heatmapmatrix<-ifelse(abs(heatmapmatrix) > 0,1,0)
heatmapmatrix[is.na(heatmapmatrix)] = 0
#cluster your selected genes
hr <- hclust(as.dist(1-cor(t(heatmapmatrix), method="pearson")), method="complete") #cluster rows by pearson correlation
hc <- hclust(as.dist(1-cor(heatmapmatrix, method="spearman")), method="average") #cluster columns by spearman correlation

#turn heatmap data back into tidy table
heatmapdata <- heatmapdata %>% pivot_longer(!Gene, names_to = "categoryID", values_to  = "foldChange") 
    
#change order of data to be plotted
heatmapdata$categoryID <- factor(heatmapdata$categoryID, levels = hc$labels)
heatmapdata$Gene <- factor(heatmapdata$Gene, levels = hr$labels)

#rename dataframe columns
colnames(heatmapdata) <- c("Gene", "categoryID", "log2foldchange")

singlecellheatmap <- ggplot(heatmapdata, aes(x = Gene, y = categoryID, fill = log2foldchange)) + 
  geom_tile(color="black") + 
  theme_classic() + 
  scale_fill_continuous_divergingx(palette = 'RdBu', mid = 0, na.value="white") +
  theme(axis.text.x = element_text(angle = 45, vjust = 1, hjust = 1), axis.title.x=element_blank(), axis.title.y=element_blank()) 

singlecellheatmap
```

```
#export 1300x400
```

### 6 Chip-seq data and DEAF1 binding sites

#### 6.1 Import human DEAF1 chip-seq data and convert to zebrafish gene names

DEAF1 Chip-seq data was downloaded from this URL: https://www.encodeproject.org/files/ENCFF813TRY/ on
10/30/2023. Accession is ENCFF813TRY. Peaks are annotated to human gene
names using ChIPSeeker. The goal is to identify

```
#import data
samplefiles <- list.files("deaf1bindingsites", pattern= ".bed", full.names=T)
samplefiles <- as.list(samplefiles)
names(samplefiles) <- c("DEAF1")

#import DEAF1 binding data
peak <- readPeakFile(samplefiles[[1]])

#import human gene annotation data
txdb <- TxDb.Hsapiens.UCSC.hg38.knownGene

#retrieve annotations for peak calls: tss is 2000 downstream and upstream, flank distance for nearby genes is 250bp
peakAnnoList <- lapply(samplefiles, annotatePeak, TxDb=txdb, 
                       tssRegion=c(-2000, 2000), verbose=FALSE, addFlankGeneInfo=TRUE, flankDistance = 250)

#make heat map of binding over promoters
promoter <- getPromoters(TxDb=txdb, upstream=2000, downstream=2000)
tagMatrix <- getTagMatrix(peak, windows=promoter)
```

```
## >> preparing start_site regions by gene... 2023-11-15 5:20:38 PM
## >> preparing tag matrix...  2023-11-15 5:20:38 PM
```

```
#make graph of binding versus distance from TSS
plotAvgProf(tagMatrix, xlim=c(-2000, 2000),
            xlab="Genomic Region (5'->3')", ylab = "Read Count Frequency")
```

```
## >> plotting figure...             2023-11-15 5:20:44 PM
```

```
#export 700x500

#This graph makes it seem like most DEAF1 binding at the promoter is between -500 and ~+500

#make bar plot of where DEAF1 is binding
plotAnnoBar(peakAnnoList)
```

```
#export 700x500

#DEAF1 is primarily binding to promoters, within 1kb of TSS

#Distribution of TF-binding loci relative to TSS
plotDistToTSS(peakAnnoList, title="Distribution of transcription factor-binding loci \n relative to TSS")
```

```
# save annotations for each peak call
deaf1_annot <- data.frame(peakAnnoList[["DEAF1"]]@anno)

#get entrez gene ids
entrez <- deaf1_annot$geneId

# Return the gene symbol for the set of Entrez IDs
annotations_edb <- AnnotationDbi::select(EnsDb.Hsapiens.v75,
                                         keys = entrez,
                                         columns = c("GENENAME"),
                                         keytype = "ENTREZID")

# Change IDs to character type to merge
annotations_edb$ENTREZID <- as.character(annotations_edb$ENTREZID)

#add geneID annotations
deaf1_annot <- deaf1_annot %>% 
  left_join(annotations_edb, by=c("geneId"="ENTREZID"))

#export file with peak annotations
write.csv(deaf1_annot, file="deaf1_chipseq_geneannotation.csv")

#filter to only include DEAF1 binding at TSS (promoters)
deaf1_promoterpeaks <- deaf1_annot %>% dplyr::filter(annotation %in% c("Promoter (<=1kb)", "Promoter (1-2kb)"))

#keep only columns with gene names
deaf1_promoterpeaks <- deaf1_promoterpeaks %>% dplyr::select(geneId, flank_geneIds)

#split flank_txIds by semicolon
deaf1_promoterpeaks <- deaf1_promoterpeaks %>% separate_wider_delim(flank_geneIds, ";", names = c(paste("x", 1:15)), too_many = "drop", too_few = "align_start")

#turn df into vector
deaf1_promoterpeaks <- unlist(deaf1_promoterpeaks, use.names=FALSE)

#remove duplicate values
deaf1_promoterpeaks <- unique(deaf1_promoterpeaks)

#remove NA
deaf1_promoterpeaks <- na.omit(deaf1_promoterpeaks)

# Return the gene symbol for the set of Entrez IDs
annotations_promoter <- AnnotationDbi::select(EnsDb.Hsapiens.v75,
                                         keys = deaf1_promoterpeaks,
                                         columns = c("GENENAME"),
                                         keytype = "ENTREZID")


#convert zebrafish genes to mouse genes(output is a dictionary)
mouse_zfish_dict <- orthogene::convert_orthologs(gene_df=annotations_promoter,
                                              gene_input="GENENAME",
                                              gene_output="dict",
                                              input_species="human",
                                              output_species="zebrafish",
                                              non121_strategy="kbs",
                                              method="gprofiler")


#change ENSDARG00000089860 to atxn7l1
mouse_zfish_dict["ATXN7L1"] <- "atxn7l1"

#get list of zebrafish genes with human CHIPSeq binding
deaf1_chipseq_genes <- unique(unname(mouse_zfish_dict))

#add EIF4G3a and EIF4G3b (EIF4G3 incorrectly converted from human)
deaf1_chipseq_genes <- c(deaf1_chipseq_genes, "eif4g3a", "eif4g3b")

#make GSEA set for deaf1 chipseq
deaf1chipseqGSEA <- as.data.frame(deaf1_chipseq_genes)
deaf1chipseqGSEA$term <- "DEAF1chipseq"
deaf1chipseqGSEA <- deaf1chipseqGSEA %>% dplyr::select(term, deaf1_chipseq_genes)
```

#### 6.2 Volcano plot for 6dpf 23bp insertion with points labeled by binding site & chip-seq data

List of zebrafish genes with DEAF1 motifs within 100 upstream of the
TSS was acquired by using https://rsat.france-bioinformatique.fr/metazoa/genome-scale-dna-pattern\_form.cgi.
Organism was Danio rerio GRCz10. Feature type was gene. Sequence type
was upstream between -100 and -1. Sequence label was gene name. Overlap
with upstream ORFs was allowed. Search strand was both strands. Match
counts was set to 1. Prevent overlapping matches was unchecked.
Substitutions was 0. Query patterns were: TCGNNNNNTCG TCGNNNNNNTCG
TCGNNNNNNNTCG TCGNNNNNNNNTCG TCGNNNNNNNNNTCG TCGNNNNNNNNNNTCG
TCGNNNNNNNNNNNTCG

The server command was: $RSAT/perl-scripts/retrieve-seq -all
-nocomment -org Danio\_rerio\_GRCz10 -feattype gene -type upstream -format
fasta -label name -from -100 -to -1 | $RSAT/perl-scripts/dna-pattern
-nolimits -v -format fasta -pl
$RSAT/public\_html/tmp/www-data/2023/11/15/gs-dna-pattern\_2023-11-15.055318\_5Nh2ac\_patterns.txt
-return counts -th 1 -subst 0 | $RSAT/perl-scripts/add-gene-info -info
descr -org Danio\_rerio\_GRCz10 | $RSAT/perl-scripts/add-linenb
dna-pattern -nolimits -v -format fasta -pl
$RSAT/public\_html/tmp/www-data/2023/11/15/gs-dna-pattern\_2023-11-15.055318\_5Nh2ac\_patterns.txt
-return counts -th 1 -subst 0 Citation: van Helden et al. (2000). Yeast
16(2), 177-187. Input format fasta Pattern file
$RSAT/public\_html/tmp/www-data/2023/11/15/gs-dna-pattern\_2023-11-15.055318\_5Nh2ac\_patterns.txt
Search method regexp Threshold 1 Allowed substitutions 0 Return fields
counts Patterns seq id score TCGNNNNNTCG TCGNNNNNTCG 1 TCGNNNNNNTCG
TCGNNNNNNTCG 1 TCGNNNNNNNTCG TCGNNNNNNNTCG 1 TCGNNNNNNNNTCG
TCGNNNNNNNNTCG 1 TCGNNNNNNNNNTCG TCGNNNNNNNNNTCG 1 TCGNNNNNNNNNNTCG
TCGNNNNNNNNNNTCG 1 TCGNNNNNNNNNNNTCG TCGNNNNNNNNNNNTCG 1

The resulting file is
gs-dna-pattern\_2023-11-15.055318\_5Nh2ac.dnapat

```
#import genes with binding sites within 100 of TSS
rsatbinding <- read.table(file="deaf1bindingsites/gs-dna-pattern_2023-11-15.055318_5Nh2ac.dnapat.txt", fill=TRUE)

#make a list of the unique binding sites
rsatbinding <- unique(rsatbinding$V1)

#add deaf1 (has DEAF1 binding motif but TSS annotation is incorrect in assembly)
rsatbinding <- c(rsatbinding, "deaf1")

#make GSEA set for deaf1 binding sites
deaf1bindingGSEA <- as.data.frame(rsatbinding)
deaf1bindingGSEA$term <- "DEAF1motif"
deaf1bindingGSEA <- deaf1bindingGSEA %>% dplyr::select(term, rsatbinding)

#make a list of genes shared between zebrafish binding site motif (rsatbinding) and human DEAF1 chipseq
sharedgenes <- intersect(rsatbinding, deaf1_chipseq_genes)

#make a list of genes in rsatbinding but not in deaf1 chipseq
rsatbinding <- rsatbinding[!rsatbinding %in% sharedgenes]

#make a list of genes in deaf1 chipseq but not in rsatbinding
deaf1_chipseq_genes <- deaf1_chipseq_genes[!deaf1_chipseq_genes %in% sharedgenes]

#add new column to deaf1DEGdata with information about binding site
deaf1DEGdata <- deaf1DEGdata %>% mutate(bindingsite = ifelse(LLgeneAbbrev %in% rsatbinding, "motif near TSS", ifelse(LLgeneAbbrev %in% deaf1_chipseq_genes, "ChIP-Seq peak", ifelse(LLgeneAbbrev %in% sharedgenes, "both", "neither"))))

#filter comparisons to only include 23bp hom v wt comparisons
myTopHits.df <- deaf1DEGdata %>% dplyr::filter(comparison %in% c("d23bp_homvwt_6dpf"))

#add column about whether chr is chr25
myTopHits.df <- myTopHits.df %>% dplyr::mutate(chrom = replace_na(chrom, "none"))
myTopHits.df <- myTopHits.df %>% mutate(chromosome = ifelse(chrom == "chr25", "chromosome 25", "other chromosome"))

#change order that points are plotted
myTopHits.df <- myTopHits.df %>% arrange(match(bindingsite, c("neither", "ChIP-Seq peak", "motif near TSS", "both")), desc(bindingsite))

genestolabel <- c("sec22a", "creb1b", "nfyal", "atxn7l1", "galk2", "nfyal", "cdkn1bb", "ambra1b", "khk", "muc5.1", "znf319b", "ccl39.3", "LOC100330348", "map1lc3b", "ccl39.6", "ccl39.7", "usb1", "cdc27",  "deaf1")

#subset to only include labeled genes
myTopHits.labels <- myTopHits.df %>% dplyr::filter(LLgeneAbbrev %in% genestolabel)

#make the plot
deaf1_6dpf_volcano <- ggplot(myTopHits.df) +
  aes(y=-log10(padjbound), x=log2FoldChange, text = paste("Symbol:", LLgeneAbbrev), color=bindingsite, shape=chromosome) +
  annotate("rect", xmin = 1, xmax = 7, ymin = -log10(0.05), ymax = 10, 
           alpha=.15,   fill="grey") +
  annotate("rect", xmin = -1, xmax = -7, ymin = -log10(0.05), ymax = 10, 
           alpha=.15, fill="grey") +
  geom_point(size=2) +
  theme_bw() +
  coord_cartesian(xlim = c(-8, 8), ylim = c(-0.5, 10.5), expand = FALSE) +
  ylab("-log10(padj)") + 
  xlab("log2 fold change") +
  geom_label_repel(data=myTopHits.labels, aes(x=log2FoldChange, y=-log10(padjbound), label=LLgeneAbbrev), force = 2, nudge_y = -1, size = 2.5, max.overlaps = Inf, show.legend = FALSE, color = "black") + #label selected genes
  theme_bw() +
  scale_color_manual(values = c("#AADC32FF", "#27AD81FF", "#472D7BFF", "darkgrey"), name = "DEAF1 motif/  \n binding near TSS") +
  scale_shape_manual(values=c(4, 16)) 

deaf1_6dpf_volcano
```

```
#export 700x500
```

#### 6.3 Volcano plot for 2dpf 23bp ins

```
#filter comparisons to only include 23bp hom v wt comparisons
myTopHits.df <- deaf1DEGdata %>% dplyr::filter(comparison %in% c("d23bp_homvwt"))

#add column about whether chr is chr25
myTopHits.df <- myTopHits.df %>% dplyr::mutate(chrom = replace_na(chrom, "none"))
myTopHits.df <- myTopHits.df %>% mutate(chromosome = ifelse(chrom == "chr25", "chromosome 25", "other chromosome"))

#change order that points are plotted
myTopHits.df <- myTopHits.df %>% arrange(match(bindingsite, c("neither", "ChIP-Seq peak", "motif near TSS", "both")), desc(bindingsite))

genestolabel <- c("khk", "muc5.1", "sec22a", "ambra1b", "znf319b", "ptpn9a", "saal1", "ptdss2", "muc5.2", "tmem138", "CABZ01039863.1", "LOC110438378", "cav1", "btbd10a", "snupn", "evlb", "mkrn1", "klhdc4", "erp44", "nfyal", "chordc1b")

#subset to only include labeled genes
myTopHits.labels <- myTopHits.df %>% dplyr::filter(LLgeneAbbrev %in% genestolabel)

#make the plot
deaf1_2dpf_23bp_volcano <- ggplot(myTopHits.df) +
  aes(y=-log10(padjbound), x=log2FoldChange, text = paste("Symbol:", LLgeneAbbrev), color=bindingsite, shape=chromosome) +
  annotate("rect", xmin = 1, xmax = 5, ymin = -log10(0.05), ymax = 10, 
           alpha=.15,   fill="grey") +
  annotate("rect", xmin = -1, xmax = -5, ymin = -log10(0.05), ymax = 10, 
           alpha=.15, fill="grey") +
  geom_point(size=2) +
  theme_bw() +
  coord_cartesian(xlim = c(-6, 6), ylim = c(-0.5, 10.5), expand = FALSE) +
  ylab("-log10(padj)") + 
  xlab("log2 fold change") +
  geom_label_repel(data=myTopHits.labels, aes(x=log2FoldChange, y=-log10(padjbound), label=LLgeneAbbrev), force = 2, nudge_y = -1, size = 2.5, max.overlaps = Inf, show.legend = FALSE, color = "black") + #label selected genes
  theme_bw() +
  scale_color_manual(values = c("#AADC32FF", "#27AD81FF", "#472D7BFF", "darkgrey"), name = "DEAF1 motif/  \n binding near TSS") +
  scale_shape_manual(values=c(4, 16)) 

deaf1_2dpf_23bp_volcano
```

```
#export 700x500
```

#### 6.4 Volcano plot for 2dpf c207y

```
#filter comparisons to only include c207y hom v wt comparison
myTopHits.df <- deaf1DEGdata %>% dplyr::filter(comparison %in% c("dc207y_homvwt"))

#add column about whether chr is chr25
myTopHits.df <- myTopHits.df %>% dplyr::mutate(chrom = replace_na(chrom, "none"))
myTopHits.df <- myTopHits.df %>% mutate(chromosome = ifelse(chrom == "chr25", "chromosome 25", "other chromosome"))

#change order that points are plotted
myTopHits.df <- myTopHits.df %>% arrange(match(bindingsite, c("neither", "ChIP-Seq peak", "motif near TSS", "both")), desc(bindingsite))

genestolabel <- c("khk", "sec22a", "ptdss2", "nfyal", "snupn", "btbd10a", "cav1", "mkrn1", "CABZ01039863.1", "hdac10", "CABZ01039859.1", "atxn7l1", "ccnd2a", "zgc:77158", "dus2", "slc25a22a", "dkk3b", "cpt1ab", "deaf1", "dusp6", "setd6", "dusp8a")

#subset to only include labeled genes
myTopHits.labels <- myTopHits.df %>% dplyr::filter(LLgeneAbbrev %in% genestolabel)

#make the plot
deaf1_2dpf_c207y_volcano <- ggplot(myTopHits.df) +
  aes(y=-log10(padjbound), x=log2FoldChange, text = paste("Symbol:", LLgeneAbbrev), color=bindingsite, shape=chromosome) +
  annotate("rect", xmin = 1, xmax = 5, ymin = -log10(0.05), ymax = 10, 
           alpha=.15,   fill="grey") +
  annotate("rect", xmin = -1, xmax = -5, ymin = -log10(0.05), ymax = 10, 
           alpha=.15, fill="grey") +
  geom_point(size=2) +
  theme_bw() +
  coord_cartesian(xlim = c(-6, 6), ylim = c(-0.5, 10.5), expand = FALSE) +
  ylab("-log10(padj)") + 
  xlab("log2 fold change") +
  geom_label_repel(data=myTopHits.labels, aes(x=log2FoldChange, y=-log10(padjbound), label=LLgeneAbbrev), force = 2, nudge_y = -1, size = 2.5, max.overlaps = Inf, show.legend = FALSE, color = "black") + #label selected genes
  theme_bw() +
  scale_color_manual(values = c("#AADC32FF", "#27AD81FF", "#472D7BFF", "darkgrey"), name = "DEAF1 motif/  \n binding near TSS") +
  scale_shape_manual(values=c(4, 16)) 

deaf1_2dpf_c207y_volcano
```

```
#export 700x500
```

#### 6.5 Volcano plot for 2dpf t238p

```
#filter comparisons to only include t238p hom v wt comparison
myTopHits.df <- deaf1DEGdata %>% dplyr::filter(comparison %in% c("dt238p_homvwt"))

#add column about whether chr is chr25
myTopHits.df <- myTopHits.df %>% dplyr::mutate(chrom = replace_na(chrom, "none"))
myTopHits.df <- myTopHits.df %>% mutate(chromosome = ifelse(chrom == "chr25", "chromosome 25", "other chromosome"))

#change order that points are plotted
myTopHits.df <- myTopHits.df %>% arrange(match(bindingsite, c("neither", "ChIP-Seq peak", "motif near TSS", "both")), desc(bindingsite))

genestolabel <- c("khk", "sec22a", "hdac10", "slc25a22a", "deaf1", "atxn7l", "brd9", "myo6a", "chchd3b", "unc45a", "tgfbr1b", "nuak1b", "mta1", "calm1a", "id3", "zgc:163040", "cdc27")

#subset to only include labeled genes
myTopHits.labels <- myTopHits.df %>% dplyr::filter(LLgeneAbbrev %in% genestolabel)

#make the plot
deaf1_2dpf_t238p_volcano <- ggplot(myTopHits.df) +
  aes(y=-log10(padjbound), x=log2FoldChange, text = paste("Symbol:", LLgeneAbbrev), color=bindingsite, shape=chromosome) +
  annotate("rect", xmin = 1, xmax = 5, ymin = -log10(0.05), ymax = 10, 
           alpha=.15,   fill="grey") +
  annotate("rect", xmin = -1, xmax = -5, ymin = -log10(0.05), ymax = 10, 
           alpha=.15, fill="grey") +
  geom_point(size=2) +
  theme_bw() +
  coord_cartesian(xlim = c(-6, 6), ylim = c(-0.5, 10.5), expand = FALSE) +
  ylab("-log10(padj)") + 
  xlab("log2 fold change") +
  geom_label_repel(data=myTopHits.labels, aes(x=log2FoldChange, y=-log10(padjbound), label=LLgeneAbbrev), force = 2, nudge_y = -1, size = 2.5, max.overlaps = Inf, show.legend = FALSE, color = "black") + #label selected genes
  theme_bw() +
  scale_color_manual(values = c("#AADC32FF", "#27AD81FF", "#472D7BFF", "darkgrey"), name = "DEAF1 motif/  \n binding near TSS") +
  scale_shape_manual(values=c(4, 16)) 

deaf1_2dpf_t238p_volcano
```

```
#export 700x500
```

### 7 Fold Change Plots

#### 7.1 Make plots of selected genes across mutants (fold change)

```
#pick out a genes to plot
geneofinterest <- c("deaf1", "cdkn1bb", "eif4g3b", "eif4g3a", "rac3a", "rac3b", "khk", "sec22a", "creb1b", "nfyal", "atxn7l1", "cdc27", "hdac10", "cdc27")

#make new columns with mean counts of wt
genecountslogfc <- genecounts
genecountslogfc$c208yavg <- rowMeans(genecountslogfc[,12:16])
genecountslogfc$t238pavg <- rowMeans(genecountslogfc[,17:20])
genecountslogfc$d23bpavg <- rowMeans(genecountslogfc[,33:37])
genecountslogfc$d23bpavg6dpf <- rowMeans(genecountslogfc[,53:56])

#divide columns by mean counts of wt
genecountslogfc <- genecountslogfc %>% mutate(across(colnames(genecountslogfc)[4:16], function(x) x/c208yavg))
genecountslogfc <- genecountslogfc %>% mutate(across(colnames(genecountslogfc)[17:29], function(x) x/t238pavg))
genecountslogfc <- genecountslogfc %>% mutate(across(colnames(genecountslogfc)[30:42], function(x) x/d23bpavg))
genecountslogfc <- genecountslogfc %>% mutate(across(colnames(genecountslogfc)[43:56], function(x) x/d23bpavg6dpf))

#pull out data for a select gene
generepdata <- genecountslogfc %>% dplyr::filter(LLgeneAbbrev %in% geneofinterest)

#get rid of unnecessary columns
generepdata <- generepdata[,c(1,4:56)]

#pivot longer to make tidy
generepdata <- generepdata %>% pivot_longer(!LLgeneAbbrev, names_to = "sample", values_to = "foldchange")

#add condition information based on sampleName
generepdata$condition <- generepdata$sample

#substitute condition name based on replicate
generepdata <- generepdata %>% mutate(condition = str_replace_all(condition, c("^chet.*"="c207y_het_2dpf", "^chom.*"="c207y_hom_2dpf", "^cwt.*" = "c207y_wt_2dpf", "^twt.*" = "t238p_wt_2dpf", "^thet.*" = "t238p_het_2dpf", "^thom.*" = "t238p_hom_2dpf", "^dhet.*" = "23d46i_het_2dpf", "^dwt.*" = "23d46i_wt_2dpf", "^dhom.*" = "23d46i_hom_2dpf", "^het.*" = "23d46i_het_6dpf", "^hom.*" = "23d46i_hom_6dpf", "^wt.*" = "23d46i_wt_6dpf")))

#split condition into three columns
generepdata <- generepdata %>% separate(condition, sep = "_", remove = FALSE, c('mutation', 'genotype', 'age'))

#make genotype a factor
generepdata$genotype <- factor(generepdata$genotype, levels = c("wt", "het", "hom"))

#make plots manually split by age
colors <- c("#472D7BFF", "#FDE725FF", "#21908CFF") #set colors

#make an empty list
plot_list = list()

#make all the plots
for (z in 1:length(geneofinterest)) {
  genedata2dpf <- generepdata %>% dplyr::filter(LLgeneAbbrev == geneofinterest[z] & age =="2dpf")
  plot <- ggplot(genedata2dpf, aes(fill=genotype, y=foldchange, x=mutation)) + 
    geom_bar(position="dodge", stat="summary") +
    geom_point(position = position_dodge(width = .9)) + 
    labs(title = paste(genedata2dpf$LLgeneAbbrev[z], ": 2dpf")) +
    ylab("fold change") +
    theme_bw() +
    scale_fill_manual(values = colors)
  genedata23bp <- generepdata %>% dplyr::filter(LLgeneAbbrev == geneofinterest[z] & mutation =="23d46i") 
  yrange <- layer_scales(plot)$y$range$range
  plot1 <- ggplot(genedata23bp, aes(fill=genotype, y=foldchange, x=age)) + 
    geom_bar(position="dodge", stat="summary") +
    geom_point(position = position_dodge(width = .9)) +
    labs(title = paste(genedata2dpf$LLgeneAbbrev[z], ": 23d46i")) +
    ylab("fold change") +
    ylim(yrange[1], yrange[2]) +
    theme_bw() +
    scale_fill_manual(values = colors)
  plot_list[[z]] <- plot_grid(plot + theme(legend.position="none"), plot1 )
}

# Save plots to svg Makes a separate file for each plot.
for (i in 1:length(geneofinterest)) {
    file_name = paste("deaf1_foldchange_", geneofinterest[i], ".svg", sep="")
    svglite(file_name)
    print(plot_list[[i]])
    dev.off()
}
```

### 8 Scatterplots

#### 8.1 Make scatterplot comparing log2 fold change of 23bp insertion 2dpf to 6dpf

```
#subset deaf1DEGdata to only include comparison, logfc, gene name, and pvalue
deaf1DEGdata_scatter <- deaf1DEGdata %>% dplyr::select(log2FoldChange, pvaluebound, chrom, bindingsite, comparison, LLgeneAbbrev)

#add column about whether chr is chr25
deaf1DEGdata_scatter <- deaf1DEGdata_scatter %>% dplyr::mutate(chrom = replace_na(chrom, "none"))
deaf1DEGdata_scatter <- deaf1DEGdata_scatter %>% mutate(chromosome = ifelse(chrom == "chr25", "chromosome 25", "other chromosome"))


#find duplicates
duplicategenes <- dplyr::filter(deaf1DEGdata_scatter %>%
  distinct() %>%
  group_by(LLgeneAbbrev, comparison) %>%
  dplyr::count(), n !=1)$LLgeneAbbrev

#remove duplicate rows from deaf1DEGdata_scatter
deaf1DEGdata_scatter <- deaf1DEGdata_scatter %>% dplyr::filter(!LLgeneAbbrev %in% duplicategenes)

#use pivot wider to make untidy table
deaf1DEGdata_scatter <- deaf1DEGdata_scatter %>% distinct() %>%
    pivot_wider(names_from = comparison, values_from  = c(log2FoldChange, pvaluebound), values_fill=NA) 
    
#add rownames
row.names(deaf1DEGdata_scatter) <- deaf1DEGdata_scatter$LLgeneAbbrev

#genes to label
genestolabel <- c("LOC110438378", "ndufa9b", "muc5d", "sec22a", "ccl39.3", "muc5.1", "khk", "ambra1b", "tmem138", "col21a1", "CABZ01039863.1", "atxn7l1", "znf319b", "ccl39.6", "ccl39.7", "XLOC_042595", "nfyal", "cyp27b1")

#subset deaf1DEGdata_scatter to only include labeled genes
myTopHits.labels <- deaf1DEGdata_scatter %>% dplyr::filter(LLgeneAbbrev %in% genestolabel)

#change order that points are plotted
deaf1DEGdata_scatter <- deaf1DEGdata_scatter %>% arrange(match(bindingsite, c("neither", "ChIP-Seq peak", "motif near TSS", "both")), desc(bindingsite))

scatterplot_deaf1_ages <- ggplot() +
  geom_point(data=deaf1DEGdata_scatter, aes(x=log2FoldChange_d23bp_homvwt, y=log2FoldChange_d23bp_homvwt_6dpf, text = paste("Symbol:", LLgeneAbbrev), color=bindingsite, shape=chromosome)) + #plot data points 
  geom_hline(yintercept=0, linetype = 'dotted') + #add horizontal line 
  geom_vline(xintercept=0, linetype = 'dotted') + #add vertical line 
  theme_bw() +
  scale_color_manual(values = c("#AADC32FF", "#27AD81FF", "#472D7BFF", "darkgrey"), name = "DEAF1 motif/  \n binding near TSS") +
  scale_shape_manual(values=c(4, 16)) +
  geom_label_repel(data=myTopHits.labels, aes(x=log2FoldChange_d23bp_homvwt, y=log2FoldChange_d23bp_homvwt_6dpf, label=LLgeneAbbrev), force = 1, nudge_y = .5, size = 2.5, max.overlaps = Inf, show.legend = FALSE, color = "black")  + #label selected genes
  ylab("log2 fold change 23d46i 6dpf") +
  xlab("log2 fold change 23d46i 2dpf")

scatterplot_deaf1_ages
```

```
#export 700x500
```

#### 8.2 Make scatterplot comparing log2 fold change of 23bp insertion v t238p

```
#genes to label
genestolabel <- c("LOC110438378", "CABZ01039863.1", "atxn7l1", "khk", "muc5.1", "sec22a", "nfyal", "unc45a", "apoa1b", "deaf1", "zgc:163040")

#subset deaf1DEGdata_scatter to only include labeled genes
myTopHits.labels <- deaf1DEGdata_scatter %>% dplyr::filter(LLgeneAbbrev %in% genestolabel)


scatterplot_deaf1_t238p <- ggplot() +
  geom_point(data=deaf1DEGdata_scatter, aes(x=log2FoldChange_d23bp_homvwt, y=log2FoldChange_dt238p_homvwt, text = paste("Symbol:", LLgeneAbbrev), color=bindingsite, shape=chromosome)) + #plot data points 
  geom_hline(yintercept=0, linetype = 'dotted') + #add horizontal line 
  geom_vline(xintercept=0, linetype = 'dotted') + #add vertical line 
  theme_bw() +
  scale_color_manual(values = c("#AADC32FF", "#27AD81FF", "#472D7BFF", "darkgrey"), name = "DEAF1 motif/  \n binding near TSS") +
  scale_shape_manual(values=c(4, 16)) +
  geom_label_repel(data=myTopHits.labels, aes(x=log2FoldChange_d23bp_homvwt, y=log2FoldChange_dt238p_homvwt, label=LLgeneAbbrev), force = 1, nudge_y = .5, size = 2.5, max.overlaps = Inf, show.legend = FALSE, color = "black")  + #label selected genes
  ylab("log2 fold change t238p 2dpf") +
  xlab("log2 fold change 23d46i 2dpf")

scatterplot_deaf1_t238p
```

```
#export 700x500
```

#### 8.3 Make scatterplot comparing log2 fold change of 23bp insertion v c207y

```
#genes to label
genestolabel <- c("CABZ01039863.1", "atxn7l1", "khk", "muc5.1", "sec22a", "nfyal",   "deaf1", "zgc:163040", "cav1", "CABZ01039859.1", "actn3a")

#subset deaf1DEGdata_scatter to only include labeled genes
myTopHits.labels <- deaf1DEGdata_scatter %>% dplyr::filter(LLgeneAbbrev %in% genestolabel)


scatterplot_deaf1_c207y <- ggplot() +
  geom_point(data=deaf1DEGdata_scatter, aes(x=log2FoldChange_d23bp_homvwt, y=log2FoldChange_dc207y_homvwt, text = paste("Symbol:", LLgeneAbbrev), color=bindingsite, shape=chromosome)) + #plot data points 
  geom_hline(yintercept=0, linetype = 'dotted') + #add horizontal line 
  geom_vline(xintercept=0, linetype = 'dotted') + #add vertical line 
  theme_bw() +
  scale_color_manual(values = c("#AADC32FF", "#27AD81FF", "#472D7BFF", "darkgrey"), name = "DEAF1 motif/  \n binding near TSS") +
  scale_shape_manual(values=c(4, 16)) +
  geom_label_repel(data=myTopHits.labels, aes(x=log2FoldChange_d23bp_homvwt, y=log2FoldChange_dc207y_homvwt, label=LLgeneAbbrev), force = 1, nudge_y = .5, size = 2.5, max.overlaps = Inf, show.legend = FALSE, color = "black")  + #label selected genes
  ylab("log2 fold change c207y 2dpf") +
  xlab("log2 fold change 23d46i 2dpf")

scatterplot_deaf1_c207y
```

```
#export 700x500
```

### 9 Motif discovery

#### 9.1 List of dysregulated genes

Get promoter sequences for dysregulated genes.

The list of shared upregulated and downregulated genes (venntableup
and venntable down) was submitted to http://rsat.sb-roscoff.fr/retrieve-ensembl-seq\_form.cgi
to retrieve promoter sequences. Query organism was zebrafish. Ensembl
database version was 110. Single organism was selected. Sequence type
was upstream/downstream. Mask repeats and mask coding sequences were
deselected. Avoid redundant sequences due to alternative transcripts was
selected. Sequence position was upstream -300 to -1. Relative to feature
was gene. Prevent overlap with neighboring was none. Sequences for
upregulated genes are in the file:
retrieve-ensembl-seq\_2023-11-15.224451\_lUoDXG and sequences for
downregulated genes are in the file:
retrieve-ensembl-seq\_2023-11-15.224849\_6kZwHv Sequences for both are in
the file: retrieve-ensembl-seq\_2023-11-15.225255\_nHSvhx

### 10 Session info

Packages and versions necessary to reproduce the results in this
report.

```
sessionInfo()
```

```
## R version 4.3.1 (2023-06-16 ucrt)
## Platform: x86_64-w64-mingw32/x64 (64-bit)
## Running under: Windows 10 x64 (build 19045)
## 
## Matrix products: default
## 
## 
## locale:
## [1] LC_COLLATE=English_United States.utf8 
## [2] LC_CTYPE=English_United States.utf8   
## [3] LC_MONETARY=English_United States.utf8
## [4] LC_NUMERIC=C                          
## [5] LC_TIME=English_United States.utf8    
## 
## time zone: America/New_York
## tzcode source: internal
## 
## attached base packages:
## [1] stats4    stats     graphics  grDevices utils     datasets  methods  
## [8] base     
## 
## other attached packages:
##  [1] svglite_2.1.2                           
##  [2] ggraph_2.1.0                            
##  [3] colorspace_2.1-0                        
##  [4] ggpubr_0.6.0                            
##  [5] lattice_0.22-5                          
##  [6] org.Dr.eg.db_3.18.0                     
##  [7] ensembldb_2.26.0                        
##  [8] AnnotationFilter_1.26.0                 
##  [9] GenomicFeatures_1.54.1                  
## [10] ChIPseeker_1.38.0                       
## [11] orthogene_1.8.0                         
## [12] plotly_4.10.3                           
## [13] BaseSet_0.9.0                           
## [14] ontologyIndex_2.11                      
## [15] enrichplot_1.22.0                       
## [16] msigdbr_7.5.1                           
## [17] clusterProfiler_4.10.0                  
## [18] gprofiler2_0.2.2                        
## [19] GSVA_1.50.0                             
## [20] GSEABase_1.64.0                         
## [21] graph_1.80.0                            
## [22] annotate_1.80.0                         
## [23] XML_3.99-0.14                           
## [24] AnnotationDbi_1.64.0                    
## [25] DT_0.30                                 
## [26] RColorBrewer_1.1-3                      
## [27] ggrepel_0.9.4                           
## [28] scales_1.2.1                            
## [29] viridis_0.6.4                           
## [30] viridisLite_0.4.2                       
## [31] cowplot_1.1.1                           
## [32] DESeq2_1.42.0                           
## [33] SummarizedExperiment_1.32.0             
## [34] Biobase_2.62.0                          
## [35] MatrixGenerics_1.14.0                   
## [36] matrixStats_1.1.0                       
## [37] GenomicRanges_1.54.1                    
## [38] GenomeInfoDb_1.38.0                     
## [39] IRanges_2.36.0                          
## [40] S4Vectors_0.40.1                        
## [41] BiocGenerics_0.48.0                     
## [42] lubridate_1.9.3                         
## [43] forcats_1.0.0                           
## [44] stringr_1.5.0                           
## [45] dplyr_1.1.3                             
## [46] purrr_1.0.2                             
## [47] readr_2.1.4                             
## [48] tidyr_1.3.0                             
## [49] tibble_3.2.1                            
## [50] ggplot2_3.4.4                           
## [51] tidyverse_2.0.0                         
## [52] knitr_1.45                              
## [53] tinytex_0.48                            
## [54] rmarkdown_2.25                          
## [55] TxDb.Hsapiens.UCSC.hg38.knownGene_3.18.0
## [56] EnsDb.Hsapiens.v75_2.99.0               
## 
## loaded via a namespace (and not attached):
##   [1] ProtGenerics_1.34.0                    
##   [2] fs_1.6.3                               
##   [3] bitops_1.0-7                           
##   [4] HDO.db_0.99.1                          
##   [5] httr_1.4.7                             
##   [6] tools_4.3.1                            
##   [7] backports_1.4.1                        
##   [8] utf8_1.2.4                             
##   [9] R6_2.5.1                               
##  [10] HDF5Array_1.30.0                       
##  [11] lazyeval_0.2.2                         
##  [12] rhdf5filters_1.14.0                    
##  [13] withr_2.5.2                            
##  [14] prettyunits_1.2.0                      
##  [15] gridExtra_2.3                          
##  [16] cli_3.6.1                              
##  [17] scatterpie_0.2.1                       
##  [18] labeling_0.4.3                         
##  [19] sass_0.4.7                             
##  [20] systemfonts_1.0.5                      
##  [21] Rsamtools_2.18.0                       
##  [22] yulab.utils_0.1.0                      
##  [23] gson_0.1.0                             
##  [24] DOSE_3.28.0                            
##  [25] plotrix_3.8-2                          
##  [26] rstudioapi_0.15.0                      
##  [27] RSQLite_2.3.2                          
##  [28] TxDb.Hsapiens.UCSC.hg19.knownGene_3.2.2
##  [29] generics_0.1.3                         
##  [30] gridGraphics_0.5-1                     
##  [31] BiocIO_1.12.0                          
##  [32] crosstalk_1.2.0                        
##  [33] vroom_1.6.4                            
##  [34] gtools_3.9.4                           
##  [35] car_3.1-2                              
##  [36] homologene_1.4.68.19.3.27              
##  [37] GO.db_3.18.0                           
##  [38] Matrix_1.6-1.1                         
##  [39] fansi_1.0.5                            
##  [40] abind_1.4-5                            
##  [41] lifecycle_1.0.4                        
##  [42] yaml_2.3.7                             
##  [43] carData_3.0-5                          
##  [44] gplots_3.1.3                           
##  [45] rhdf5_2.46.0                           
##  [46] qvalue_2.34.0                          
##  [47] SparseArray_1.2.0                      
##  [48] BiocFileCache_2.10.1                   
##  [49] grid_4.3.1                             
##  [50] blob_1.2.4                             
##  [51] promises_1.2.1                         
##  [52] crayon_1.5.2                           
##  [53] beachmat_2.18.0                        
##  [54] KEGGREST_1.42.0                        
##  [55] pillar_1.9.0                           
##  [56] fgsea_1.28.0                           
##  [57] boot_1.3-28.1                          
##  [58] rjson_0.2.21                           
##  [59] codetools_0.2-19                       
##  [60] fastmatch_1.1-4                        
##  [61] glue_1.6.2                             
##  [62] ggfun_0.1.3                            
##  [63] data.table_1.14.8                      
##  [64] vctrs_0.6.4                            
##  [65] png_0.1-8                              
##  [66] treeio_1.26.0                          
##  [67] gtable_0.3.4                           
##  [68] cachem_1.0.8                           
##  [69] xfun_0.41                              
##  [70] S4Arrays_1.2.0                         
##  [71] mime_0.12                              
##  [72] tidygraph_1.2.3                        
##  [73] SingleCellExperiment_1.24.0            
##  [74] interactiveDisplayBase_1.40.0          
##  [75] ellipsis_0.3.2                         
##  [76] nlme_3.1-163                           
##  [77] ggtree_3.10.0                          
##  [78] bit64_4.0.5                            
##  [79] progress_1.2.2                         
##  [80] filelock_1.0.2                         
##  [81] bslib_0.5.1                            
##  [82] irlba_2.3.5.1                          
##  [83] KernSmooth_2.23-22                     
##  [84] DBI_1.1.3                              
##  [85] tidyselect_1.2.0                       
##  [86] bit_4.0.5                              
##  [87] compiler_4.3.1                         
##  [88] curl_5.1.0                             
##  [89] xml2_1.3.5                             
##  [90] DelayedArray_0.28.0                    
##  [91] shadowtext_0.1.2                       
##  [92] rtracklayer_1.62.0                     
##  [93] caTools_1.18.2                         
##  [94] rappdirs_0.3.3                         
##  [95] digest_0.6.33                          
##  [96] XVector_0.42.0                         
##  [97] htmltools_0.5.6.1                      
##  [98] pkgconfig_2.0.3                        
##  [99] sparseMatrixStats_1.14.0               
## [100] highr_0.10                             
## [101] dbplyr_2.4.0                           
## [102] fastmap_1.1.1                          
## [103] rlang_1.1.1                            
## [104] htmlwidgets_1.6.2                      
## [105] shiny_1.7.5.1                          
## [106] DelayedMatrixStats_1.24.0              
## [107] farver_2.1.1                           
## [108] jquerylib_0.1.4                        
## [109] jsonlite_1.8.7                         
## [110] BiocParallel_1.36.0                    
## [111] GOSemSim_2.28.0                        
## [112] BiocSingular_1.18.0                    
## [113] RCurl_1.98-1.12                        
## [114] magrittr_2.0.3                         
## [115] GenomeInfoDbData_1.2.11                
## [116] ggplotify_0.1.2                        
## [117] patchwork_1.1.3                        
## [118] Rhdf5lib_1.24.0                        
## [119] munsell_0.5.0                          
## [120] Rcpp_1.0.11                            
## [121] ggnewscale_0.4.9                       
## [122] ape_5.7-1                              
## [123] babelgene_22.9                         
## [124] stringi_1.8.1                          
## [125] zlibbioc_1.48.0                        
## [126] MASS_7.3-60                            
## [127] AnnotationHub_3.10.0                   
## [128] plyr_1.8.9                             
## [129] parallel_4.3.1                         
## [130] HPO.db_0.99.2                          
## [131] Biostrings_2.70.1                      
## [132] graphlayouts_1.0.1                     
## [133] splines_4.3.1                          
## [134] hms_1.1.3                              
## [135] locfit_1.5-9.8                         
## [136] igraph_1.5.1                           
## [137] ggsignif_0.6.4                         
## [138] reshape2_1.4.4                         
## [139] biomaRt_2.58.0                         
## [140] ScaledMatrix_1.10.0                    
## [141] BiocVersion_3.18.0                     
## [142] evaluate_0.23                          
## [143] BiocManager_1.30.22                    
## [144] tzdb_0.4.0                             
## [145] tweenr_2.0.2                           
## [146] httpuv_1.6.12                          
## [147] grr_0.9.5                              
## [148] polyclip_1.10-6                        
## [149] ggforce_0.4.1                          
## [150] rsvd_1.0.5                             
## [151] broom_1.0.5                            
## [152] xtable_1.8-4                           
## [153] restfulr_0.0.15                        
## [154] tidytree_0.4.5                         
## [155] MPO.db_0.99.7                          
## [156] rstatix_0.7.2                          
## [157] later_1.3.1                            
## [158] snow_0.4-4                             
## [159] aplot_0.2.2                            
## [160] GenomicAlignments_1.38.0               
## [161] memoise_2.0.1                          
## [162] timechange_0.2.0
```
