## Supplementary material for "Diencephalic and Neuropeptidergic Dysfunction in Zebrafish with Autism Risk Mutations": Data S1 - RNA-seq analysis files: STREME Results_deaf1allgenes_landscape.pdf

**Sensitive, Thorough, Rapid, Enriched Motif Elicitation**

For further information on how to interpret these results please access https://meme-suite.org/meme/doc/streme.html.
To get a copy of the MEME software please access https://meme-suite.org.

If you use STREME in your research, please cite the following paper:
Timothy L. Bailey, "STREME: accurate and versatile sequence motif discovery", *Bioinformatics*, Mar. 24, 2021. [full text]

DISCOVERED MOTIFS | INPUTS & SETTINGS | PROGRAM INFORMATION | MOTIFS IN MEME TEXT FORMAT | MATCHING SEQUENCES | MATCHING SITES | RESULTS IN XML FORMAT

### DISCOVERED MOTIFS

Next Top

| Motif | Logo | RC Logo | P-value | E-value | Sites | More | Submit/Download | Positional Distribution | Matches per Sequence |
|---|---|---|---|---|---|---|---|---|---|
| 1-GACCAATCCC | 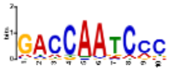 | 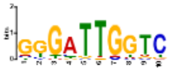 | 1.3e-002 | 8.9e-002 | 64 (15.9%) | ↧ | ⇢ | 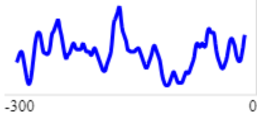 | 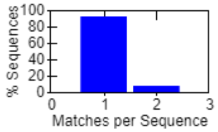 |
| 2-RTAGSCTAY | 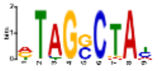 | 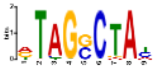 | 1.3e-002 | 8.9e-002 | 36 (9.0%) | ↧ | ⇢ | 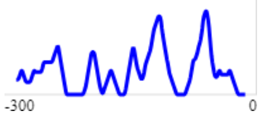 | 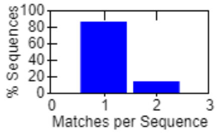 |
| 3-AGCTCCGCCYH | 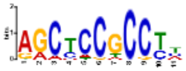 | 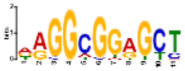 | 2.7e-002 | 1.9e-001 | 61 (15.2%) | ↧ | ⇢ | 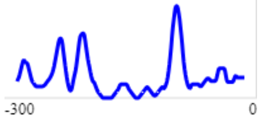 | 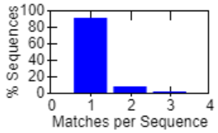 |
| 4-TAAATAAATAA | 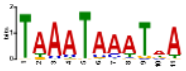 | 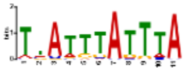 | 3.3e-002 | 2.3e-001 | 143 (35.6%) | ↧ | ⇢ | 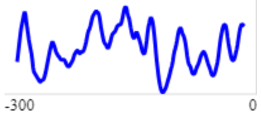 | 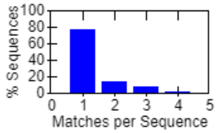 |
| 5-AGCGACTGTAAARCY | 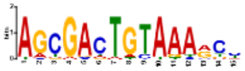 | 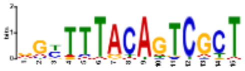 | 5.8e-002 | 4.0e-001 | 32 (8.0%) | ↧ | ⇢ | 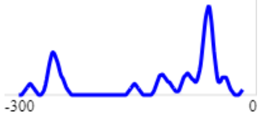 | 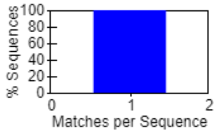 |
| 6-ACCTAGTTGA | 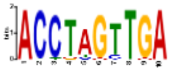 | 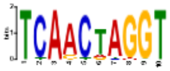 | 1.2e-001 | 8.4e-001 | 36 (9.0%) | ↧ | ⇢ | 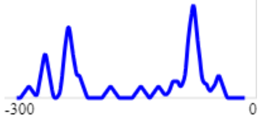 | 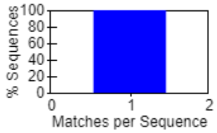 |

**Stopped because 3 consecutive motifs exceeded the p-value threshold (0.05).**
**STREME ran for 19.3 seconds.**

| Motif | Logo | RC Logo | P-value | E-value | Sites | More | Submit/Download | Positional Distribution | Matches per Sequence |
|---|---|---|---|---|---|---|---|---|---|
| 7-CACACAHACACAC | 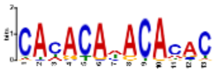 | 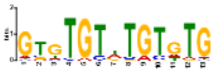 | 3.7e-001 | 2.6e+000 | 108 (26.9%) | ↧ | ⇢ | 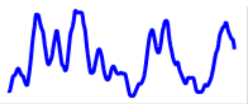 | 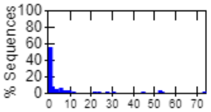 |

**Stopped because 3 consecutive motifs exceeded the p-value threshold (0.05).**
**STREME ran for 19.3 seconds.**

### Inputs & Settings

Previous Next Top

#### Sequences

| Role | Source | Alphabet | Sequence Count | Total Size |
|---|---|---|---|---|
| Positive (primary) Sequences | retrieve-ensembl-seq_2023-11-15.225255_nHSvhx.txt | DNA | 402 | 120600 |
| Negative (control) Sequences | 2-Order Shuffled Positive Sequences | DNA | 402 | 120600 |

#### Background Model

**Source:** built from the negative (control) sequences

**Order:** 2 (only order-0 shown)

| Name | Freq. | Bg. | | | | Bg. | Freq. | Name |
|---|---|---|---|---|---|---|---|---|
| Adenine | 0.307 | 0.307 | A | ~ | T | 0.307 | 0.307 | Thymine |
| Cytosine | 0.193 | 0.193 | C | ~ | G | 0.193 | 0.193 | Guanine |

#### Other Settings

| | |
|---|---|
| **Strand Handling** | Both the given and reverse complement strands are processed. |
| **Objective Function** | Differential Enrichment |
| **Statistical Test** | Fisher Exact Test |
| **Minimum Motif Width** | 5 |
| **Maximum Motif Width** | 15 |
| **Sequence Shuffling** | Negative sequences are positives shuffled preserving 3-mer frequencies. |
| **Test Set** | 10% of the input sequences were randomly assigned to the test set. |
| **Word Evaluation** | Up to 25 words of each width from 5 to 15 were evaluated to find seeds. |
| **Seed Refinement** | Up to 4 seeds of each width from 5 to 15 were further refined. |
| **Refinement Iterations** | Up to 20 iterations were allowed when refining a seed. |
| **Random Number Seed** | 0 |
| **Trimming of Control Sequences** | Trimming of control sequences was allowed. |
| **Total Length** | The total length of each sequence set was limited to 4.00e+6. |
| **Maximum Motif p-value** | Stop when the p-value is greater than 0.05 for 3 consecutive motifs. |
| **Maximum Motifs to Find** | No maximum number of motifs. |
| **Maximum Run Time** | 14400 seconds. |

Previous Top

**STREME version**
5.5.4 (Release date: Fri Jun 16 12:19:08 2023 -0700)

**Reference**
Timothy L. Bailey, "STREME: accurate and versatile sequence motif discovery", *Bioinformatics*, Mar. 24, 2021. [full text]

**Command line**

```
streme --verbosity 1 --oc . --dna --totallength 4000000 --time 14400 --minw 5 --maxw 15 --thresh 0.05 --align right --p retrieve-ensembl-seq_2023-11-15.225255_nHSvhx.txt
```
