## Supplementary material for "Diencephalic and Neuropeptidergic Dysfunction in Zebrafish with Autism Risk Mutations": Data S1 - RNA-seq analysis files: kmt5b_hdlbpa_rnaseq.html

kmt5b versus hdlbpa data


Code 

- Show All Code
- Hide All Code

### kmt5b versus hdlbpa data

###### Anna Moyer

#### *2023-11-28*

Compare kmt5b and hdlbpa mutant RNAseq data from 6dpf zebrafish
heads.

### 1 Setup

#### 1.1 Set seed and working directory

```
#set random seed
set.seed(123)

#set working directory
setwd("C:/Users/annam/Desktop/lab work/autismrnaseq/kmt5b-hdlbp/kmt5b-hdlbp")
```

#### 1.2 Load packages

```
library("tidyverse") 
library("DESeq2") #finding differentially expressed genes 
library("cowplot") #arranging plots into grids
library("viridis") #viridis color schemes
library("scales") #use to get color schemes for viridis
library("ggrepel") #repels text labels on plots
library("RColorBrewer") #pick colors
library("DT") #interactive and searchable tables of our GSEA results
library("GSEABase") #functions and methods for Gene Set Enrichment Analysis
library("Biobase") #base functions for bioconductor; required by GSEABase
library("GSVA") #Gene Set Variation Analysis, a non-parametric and unsupervised method for estimating variation of gene set enrichment across samples.
library("gprofiler2") #tools for accessing the GO enrichment results using g:Profiler web resources
library("clusterProfiler") # provides a suite of tools for functional enrichment analysis
library("msigdbr") # access to msigdb collections directly within R
library("enrichplot") # great for making the standard GSEA enrichment plots
library("ontologyIndex") #for parsing obo files
library("BaseSet") #for importing gaf file
library("plotly") #make interactive plots
library("orthogene") #convert gene names between species
library("ChIPseeker") #for annotating and browsing chip-seq data
library("TxDb.Hsapiens.UCSC.hg38.knownGene", pos = .Machine$integer.max) #human gene annotation for chip-seq data
library("EnsDb.Hsapiens.v75", pos = .Machine$integer.max) #human gene annotation for chip-seq data
library("org.Dr.eg.db") #convert ensembl gene names to gene id
library("lattice") #used for making manhattan plot
library("ggpubr") #calculating correlation coefficient 
library("colorspace") #colors for heatmap
library("ggraph") #library for making network graphs
library("svglite") #export svgs in loop
library("eulerr") #make venn diagrams
```

#### 1.3 Import files

```
#import  deseq output for kmt5b data
kmt5b <- read.csv("kmt5b-homvswt3x3_allresults_wt_hom-with-normalized.csv")[,2:9]

#import  normalized counts for kmt5b data
kmt5b_genecounts <- read.csv("kmt5b-homvswt3x3_normalized_reads_gene_list.csv")[,2:10]

#import  deseq output for hdlbpa
hdlbpa <- read.csv("hdlbpa-homvswt_allresults_wt_hom-with-normalized.csv")[,2:9]

#import  normalized counts for hdlbpa
hdlbpa_genecounts <- read.csv("hdlbpa-homvswt_normalized_reads_gene_list.csv")[,2:12]

#add columns to old and new data about whether data is old or new
kmt5b$mutation <- "kmt5b"
hdlbpa$mutation <- "hdlbpa"

#combine two mutations
kmt5b_hdlbpa <- rbind(kmt5b, hdlbpa)

#add some information 
#limit padj to 1e-10
kmt5b_hdlbpa <- kmt5b_hdlbpa %>% mutate(padjbound = ifelse(kmt5b_hdlbpa$padj < 1e-10, 1e-10, kmt5b_hdlbpa$padj))

#limit pvalue to 1e-10
kmt5b_hdlbpa <- kmt5b_hdlbpa %>% mutate(pvaluebound = ifelse(kmt5b_hdlbpa$pvalue < 1e-10, 1e-10, kmt5b_hdlbpa$pvalue))

#import annotations for chromosome and TSS
chromosomeinfo <- readr::read_tsv(file="ncbi_refseqgenes")

#keep chromosome and gene name
chromosomeinfo <- chromosomeinfo %>% dplyr::select(chrom, name2, txStart)

kmt5b_hdlbpa$chrom <- NA
kmt5b_hdlbpa$txStart <- NA

#add chromosome and txStart information
for (x in 1:length(kmt5b_hdlbpa$LLgeneAbbrev)) {
  gene <- kmt5b_hdlbpa$LLgeneAbbrev[x]
  chromosomeinfosub <- chromosomeinfo %>% dplyr::filter(name2 == gene)
  chr <- chromosomeinfosub$chrom[1]
  txStart <- chromosomeinfosub$txStart[1]
  kmt5b_hdlbpa$chrom[x] <- chr
  kmt5b_hdlbpa$txStart[x] <- txStart
}

#add chromosome number for RASGRF1
kmt5b_hdlbpa <- kmt5b_hdlbpa %>% mutate(chrom = ifelse(LLgeneAbbrev == "RASGRF1", "chr18", chrom))

#add a column with chromosome number
kmt5b_hdlbpa <- kmt5b_hdlbpa %>% mutate(chromosomenumber = str_replace_all(chrom, c("chr1"="1", "chr2"="2", "chr3"="3", "chr4"="4", "chr5"="5", "chr6"="6", "chr7"="7", "chr8"="8", "chr9"="9", "chr10"="10", "chr11"="11", "chr12"="12", "chr13"="13", "chr14"="14", "chr15"="15", "chr16"="16", "chr17"="17", "chr18"="18", "chr19"="19", "chr20"="20", "chr21"="21", "chr22"="22", "chr23"="23", "chr24"="24", "chr25"="25")))

kmt5b_hdlbpa <- distinct(kmt5b_hdlbpa)

#export csv with DEG for both 2dpf and 6dpf
write.csv(kmt5b_hdlbpa,file = "kmt5b_hdlbpa_combined.csv")
```

### 2 Plotting DEGs by chromosome

Are differentially expressed genes located at a particular place in
the genome?

#### 2.1 Make Manhattan plot for kmt5b data

This function is from: https://genome.sph.umich.edu/wiki/Code\_Sample:\_Generating\_Manhattan\_Plots\_in\_R

```
#import  comparisons (to skip running previous steps)
kmt5b_hdlbpa <- read.csv("kmt5b_hdlbpa_combined.csv")[,2:15]

#function for making manhattan plots
manhattan.plot<-function(chr, pos, pvalue, 
    sig.level=NA, annotate=NULL, ann.default=list(),
    should.thin=T, thin.pos.places=2, thin.logp.places=2, 
    xlab="Chromosome", ylab=expression(-log[10](p-value)),
    col=c("gray","darkgray"), panel.extra=NULL, pch=20, cex=0.8,...) {

    if (length(chr)==0) stop("chromosome vector is empty")
    if (length(pos)==0) stop("position vector is empty")
    if (length(pvalue)==0) stop("pvalue vector is empty")

    #make sure we have an ordered factor
    if(!is.ordered(chr)) {
        chr <- ordered(chr)
    } else {
        chr <- chr[,drop=T]
    }

    #make sure positions are in kbp
    if (any(pos>1e6)) pos<-pos/1e6;

    #calculate absolute genomic position
    #from relative chromosomal positions
    posmin <- tapply(pos,chr, min);
    posmax <- tapply(pos,chr, max);
    posshift <- head(c(0,cumsum(posmax)),-1);
    names(posshift) <- levels(chr)
    genpos <- pos + posshift[chr];
    getGenPos<-function(cchr, cpos) {
        p<-posshift[as.character(cchr)]+cpos
        return(p)
    }

    #parse annotations
    grp <- NULL
    ann.settings <- list()
    label.default<-list(x="peak",y="peak",adj=NULL, pos=3, offset=0.5, 
        col=NULL, fontface=NULL, fontsize=NULL, show=F)
    parse.label<-function(rawval, groupname) {
        r<-list(text=groupname)
        if(is.logical(rawval)) {
            if(!rawval) {r$show <- F}
        } else if (is.character(rawval) || is.expression(rawval)) {
            if(nchar(rawval)>=1) {
                r$text <- rawval
            }
        } else if (is.list(rawval)) {
            r <- modifyList(r, rawval)
        }
        return(r)
    }

    if(!is.null(annotate)) {
        if (is.list(annotate)) {
            grp <- annotate[[1]]
        } else {
            grp <- annotate
        } 
        if (!is.factor(grp)) {
            grp <- factor(grp)
        }
    } else {
        grp <- factor(rep(1, times=length(pvalue)))
    }
  
    ann.settings<-vector("list", length(levels(grp)))
    ann.settings[[1]]<-list(pch=pch, col=col, cex=cex, fill=col, label=label.default)

    if (length(ann.settings)>1) { 
        lcols<-trellis.par.get("superpose.symbol")$col 
        lfills<-trellis.par.get("superpose.symbol")$fill
        for(i in 2:length(levels(grp))) {
            ann.settings[[i]]<-list(pch=pch, 
                col=lcols[(i-2) %% length(lcols) +1 ], 
                fill=lfills[(i-2) %% length(lfills) +1 ], 
                cex=cex, label=label.default);
            ann.settings[[i]]$label$show <- T
        }
        names(ann.settings)<-levels(grp)
    }
    for(i in 1:length(ann.settings)) {
        if (i>1) {ann.settings[[i]] <- modifyList(ann.settings[[i]], ann.default)}
        ann.settings[[i]]$label <- modifyList(ann.settings[[i]]$label, 
            parse.label(ann.settings[[i]]$label, levels(grp)[i]))
    }
    if(is.list(annotate) && length(annotate)>1) {
        user.cols <- 2:length(annotate)
        ann.cols <- c()
        if(!is.null(names(annotate[-1])) && all(names(annotate[-1])!="")) {
            ann.cols<-match(names(annotate)[-1], names(ann.settings))
        } else {
            ann.cols<-user.cols-1
        }
        for(i in seq_along(user.cols)) {
            if(!is.null(annotate[[user.cols[i]]]$label)) {
                annotate[[user.cols[i]]]$label<-parse.label(annotate[[user.cols[i]]]$label, 
                    levels(grp)[ann.cols[i]])
            }
            ann.settings[[ann.cols[i]]]<-modifyList(ann.settings[[ann.cols[i]]], 
                annotate[[user.cols[i]]])
        }
    }
    rm(annotate)

    #reduce number of points plotted
    if(should.thin) {
        thinned <- unique(data.frame(
            logp=round(-log10(pvalue),thin.logp.places), 
            pos=round(genpos,thin.pos.places), 
            chr=chr,
            grp=grp)
        )
        logp <- thinned$logp
        genpos <- thinned$pos
        chr <- thinned$chr
        grp <- thinned$grp
        rm(thinned)
    } else {
        logp <- -log10(pvalue)
    }
    rm(pos, pvalue)
    gc()

    #custom axis to print chromosome names
    axis.chr <- function(side,...) {
        if(side=="bottom") {
            panel.axis(side=side, outside=T,
                at=((posmax+posmin)/2+posshift),
                labels=levels(chr), 
                ticks=F, rot=0,
                check.overlap=F
            )
        } else if (side=="top" || side=="right") {
            panel.axis(side=side, draw.labels=F, ticks=F);
        }
        else {
            axis.default(side=side,...);
        }
     }

    #make sure the y-lim covers the range (plus a bit more to look nice)
    prepanel.chr<-function(x,y,...) { 
        A<-list();
        maxy<-ceiling(max(y, ifelse(!is.na(sig.level), -log10(sig.level), 0)))+.5;
        A$ylim=c(0,maxy);
        A;
    }

    xyplot(logp~genpos, chr=chr, groups=grp,
        axis=axis.chr, ann.settings=ann.settings, 
        prepanel=prepanel.chr, scales=list(axs="i"),
        panel=function(x, y, ..., getgenpos) {
            if(!is.na(sig.level)) {
                #add significance line (if requested)
                panel.abline(h=-log10(sig.level), lty=2);
            }
            panel.superpose(x, y, ..., getgenpos=getgenpos);
            if(!is.null(panel.extra)) {
                panel.extra(x,y, getgenpos, ...)
            }
        },
        panel.groups = function(x,y,..., subscripts, group.number) {
            A<-list(...)
            #allow for different annotation settings
            gs <- ann.settings[[group.number]]
            A$col.symbol <- gs$col[(as.numeric(chr[subscripts])-1) %% length(gs$col) + 1]    
            A$cex <- gs$cex[(as.numeric(chr[subscripts])-1) %% length(gs$cex) + 1]
            A$pch <- gs$pch[(as.numeric(chr[subscripts])-1) %% length(gs$pch) + 1]
            A$fill <- gs$fill[(as.numeric(chr[subscripts])-1) %% length(gs$fill) + 1]
            A$x <- x
            A$y <- y
            do.call("panel.xyplot", A)
            #draw labels (if requested)
            if(gs$label$show) {
                gt<-gs$label
                names(gt)[which(names(gt)=="text")]<-"labels"
                gt$show<-NULL
                if(is.character(gt$x) | is.character(gt$y)) {
                    peak = which.max(y)
                    center = mean(range(x))
                    if (is.character(gt$x)) {
                        if(gt$x=="peak") {gt$x<-x[peak]}
                        if(gt$x=="center") {gt$x<-center}
                    }
                    if (is.character(gt$y)) {
                        if(gt$y=="peak") {gt$y<-y[peak]}
                    }
                }
                if(is.list(gt$x)) {
                    gt$x<-A$getgenpos(gt$x[[1]],gt$x[[2]])
                }
                do.call("panel.text", gt)
            }
        },
        xlab=xlab, ylab=ylab, 
        panel.extra=panel.extra, getgenpos=getGenPos, ...
    );
}

#select columns to include in manhattan plot
myTopHits.df <- kmt5b_hdlbpa %>% dplyr::select(chromosomenumber, txStart, pvaluebound, mutation)

#filter to only include chromosomes 1-25
myTopHits.df <- myTopHits.df %>% dplyr::filter(chromosomenumber %in% 1:25)

#make chromosomes plotted in order
myTopHits.df$chromosomenumber <- factor(myTopHits.df$chromosomenumber, levels = 1:25)

#omit NA values
myTopHits.df <- na.omit(myTopHits.df)

#make colors 
manhattancol <- c(replicate(8, c("lightgrey", "darkgrey")), "lightgrey", "#21908CFF", replicate(4, c("lightgrey", "darkgrey")))
  
#filter comparisons to only include old data
manhattan <- myTopHits.df %>% dplyr::filter(mutation == "kmt5b")

#make manhattan plot
manhattan.plot(manhattan$chromosomenumber, manhattan$txStart, manhattan$pvaluebound, should.thin=F, col=manhattancol)
```

```
#export manhattan plots at 1000x500 pixels
```

#### 2.2 Make Manhattan plot for hdlbpa

```
#filter comparisons to only include old data
manhattan <- myTopHits.df %>% dplyr::filter(mutation == "hdlbpa")

#make colors 
manhattancol <- c(replicate(2, c("lightgrey", "darkgrey")), "lightgrey", "#21908CFF", replicate(9, c("lightgrey", "darkgrey")))
  

#make manhattan plot
manhattan.plot(manhattan$chromosomenumber, manhattan$txStart, manhattan$pvaluebound, should.thin=F, col=manhattancol)
```

```
#export manhattan plots at 1000x500 pixels
```

#### 2.3 Make custom GSEA annotation with zebrafish genes and chromosomes

```
#import annotations for chromosome and gene name
chromosomeinfo <- readr::read_tsv(file="ncbi_refseqgenes")

#keep chromosome and gene name
chromosomeinfo <- chromosomeinfo %>% dplyr::select(chrom, name2)

#change column names
colnames(chromosomeinfo) <- c("chromosome", "genename")

#select only chromosomes 1-25
chromosomeinfo <- chromosomeinfo %>% dplyr::filter(chromosome %in% paste("chr", 1:25, sep=""))

#keep only distinct rows
chromosomeinfo <- distinct(chromosomeinfo)
```

#### 2.4 Check for enrichment of chromosome 18 genes in kmt5b data

```
# Perform GSEA using clusterProfiler
#filter to only one comparison
GSEAgenes <- kmt5b_hdlbpa %>% dplyr::filter(mutation == "kmt5b")

#keep only the columns we need for GSEA
mydata.df.sub <- dplyr::select(GSEAgenes, LLgeneAbbrev, log2FoldChange)

#sort by abs(foldchange)
mydata.df.sub$log2FoldChange <- abs(mydata.df.sub$log2FoldChange)

# construct a named vector
mydata.gsea <- mydata.df.sub$log2FoldChange
names(mydata.gsea) <- as.character(mydata.df.sub$LLgeneAbbrev)
mydata.gsea <- sort(mydata.gsea, decreasing = TRUE)

# run GSEA using the 'GSEA' function from clusterProfiler
myGSEA.res <- GSEA(mydata.gsea, TERM2GENE=chromosomeinfo, verbose=FALSE, scoreType="pos", maxGSSize=2000, pvalueCutoff = 1, seed=TRUE)
myGSEA.df <- as_tibble(myGSEA.res@result)

# view results as an interactive table
datatable(myGSEA.df, 
          extensions = c('KeyTable', "FixedHeader"), 
          options = list(keys = TRUE, searchHighlight = TRUE, pageLength = 10, lengthMenu = c("10", "25", "50", "100"))) %>%
  formatRound(columns=c(2:10), digits=2)
```

```
# create enrichment plots using the enrichplot package
gseaplot2(myGSEA.res, 
          geneSetID = c(1), #can choose multiple signatures to overlay in this plot
         pvalue_table = FALSE, #can set this to FALSE for a cleaner plot
        title = myGSEA.res$Description[1]) #can also turn off this title
```

Chromosome 18 genes are enriched in the kmt5b mutants.

#### 2.5 Check for enrichment of chromosome 6 genes in hdlbpa data

```
# Perform GSEA using clusterProfiler
#filter to only one comparison
GSEAgenes <- kmt5b_hdlbpa %>% dplyr::filter(mutation == "hdlbpa")

#keep only the columns we need for GSEA
mydata.df.sub <- dplyr::select(GSEAgenes, LLgeneAbbrev, log2FoldChange)

#sort by abs(foldchange)
mydata.df.sub$log2FoldChange <- abs(mydata.df.sub$log2FoldChange)

# construct a named vector
mydata.gsea <- mydata.df.sub$log2FoldChange
names(mydata.gsea) <- as.character(mydata.df.sub$LLgeneAbbrev)
mydata.gsea <- sort(mydata.gsea, decreasing = TRUE)

# run GSEA using the 'GSEA' function from clusterProfiler
myGSEA.res <- GSEA(mydata.gsea, TERM2GENE=chromosomeinfo, verbose=FALSE, scoreType="pos", maxGSSize=2000, pvalueCutoff = 1, seed=TRUE)
myGSEA.df <- as_tibble(myGSEA.res@result)

# view results as an interactive table
datatable(myGSEA.df, 
          extensions = c('KeyTable', "FixedHeader"), 
          options = list(keys = TRUE, searchHighlight = TRUE, pageLength = 10, lengthMenu = c("10", "25", "50", "100"))) %>%
  formatRound(columns=c(2:10), digits=2)
```

```
# create enrichment plots using the enrichplot package
gseaplot2(myGSEA.res, 
          geneSetID = c(1), #can choose multiple signatures to overlay in this plot
         pvalue_table = FALSE, #can set this to FALSE for a cleaner plot
        title = myGSEA.res$Description[1]) #can also turn off this title
```

Chromosome 6 genes are enriched in the hdlbpa mutant.

### 3 Find genes that are shared between kmt5b and hdlbpa

#### 3.1 Make euler diagram of genes that are shared between the two mutants

```
#genes significant in kmt5b
kmt5bgenes <- (kmt5b_hdlbpa %>% dplyr::filter(pvalue < 0.05 & mutation == "kmt5b" & abs(log2FoldChange) > 0))$LLgeneAbbrev

#number significant genes in kmt5b
numberkmt5b <- length(kmt5bgenes)

#genes significant in hdlbpa
hdlbpagenes <- (kmt5b_hdlbpa %>% dplyr::filter(pvalue < 0.05 & mutation == "hdlbpa"  & abs(log2FoldChange) > 0))$LLgeneAbbrev

#number significant genes in hdlbpa
numberhdlbpa <- length(hdlbpagenes)

#shared genes
sharedgenes <- intersect(kmt5bgenes, hdlbpagenes)

#number of shared genes
numbershared <- length(sharedgenes)

#make data for venn diagram
fit1 <- euler(c("kmt5b" = 5207-675, "hdlbpa" = 2154-675, "kmt5b&hdlbpa" = 675))

#plot
euler <- plot(fit1, quantities=TRUE)

euler
```

```
#export 700x500

sharedgenes
```

```
##   [1] "cngb1a"               "slc6a6b"              "irs4a"               
##   [4] "mbd4"                 "pitpnab"              "phlda1"              
##   [7] "carmil2"              "fgd4a"                "XLOC_022740"         
##  [10] "BX663516.2"           "sh3gl2b"              "dyrk4"               
##  [13] "ppp1r21"              "CABZ01099795.1"       "CR318607.1"          
##  [16] "scp2b"                "gatad2b"              "FO082781.1"          
##  [19] "XLOC_018182"          "larp4ab"              "odc1"                
##  [22] "nicn1"                "rbm24b"               "khdc4"               
##  [25] "FP016056.1"           "ube2d4"               "wwp1"                
##  [28] "nrarpb"               "cdc37"                "si:dkey-237i9.1"     
##  [31] "rps2"                 "XLOC_014336"          "pik3r3b"             
##  [34] "nrip1a"               "si:ch211-125e6.5"     "setdb2"              
##  [37] "tspan17"              "abcf2a"               "CABZ01055347.1"      
##  [40] "hlcs"                 "uggt1"                "ddb1"                
##  [43] "inpp5kb"              "lpar1"                "slc6a3"              
##  [46] "emp2"                 "ppfia4"               "nsa2"                
##  [49] "fam169aa"             "GGT7"                 "gpr75"               
##  [52] "osbpl9"               "lrtm2a"               "tab1"                
##  [55] "rap1b"                "LOC100332847"         "foxp4"               
##  [58] "CR407584.1"           "ralgps2"              "si:ch73-46j18.5"     
##  [61] "unkl"                 "sprn2"                "thap1"               
##  [64] "ap1ar"                "pcmtl"                "agmo"                
##  [67] "slc38a5a"             "eno3"                 "rab4b"               
##  [70] "XLOC_033366"          "cul4a"                "tcn2"                
##  [73] "tnksb"                "tmem74b"              "srl"                 
##  [76] "CABZ01084081.1"       "rps16"                "raf1b"               
##  [79] "nat16"                "rp1l1b"               "prr5a"               
##  [82] "rps15"                "il10rb"               "lrit1b"              
##  [85] "cntn1a"               "ppp3cca"              "med17"               
##  [88] "fbxo7"                "rgs5a"                "XLOC_034149"         
##  [91] "igf2bp2a"             "fgfr1b"               "ghitm"               
##  [94] "march8"               "hbbe2"                "slc6a9"              
##  [97] "si:dkeyp-1h4.6"       "gorasp2"              "slc7a11"             
## [100] "march2"               "keap1b"               "slc2a1b"             
## [103] "dab2ipb"              "ap1m1"                "tab3"                
## [106] "sap130a"              "LOC101884451"         "BX000438.2"          
## [109] "pde8a"                "BX323076.1"           "spon1a"              
## [112] "syvn1"                "shisa6"               "patz1"               
## [115] "tead3a"               "tpst1l"               "ubash3ba"            
## [118] "sumo3b"               "cmbl"                 "insyn1"              
## [121] "rab5ab"               "tmem11"               "arhgef18b"           
## [124] "XLOC_034498"          "XLOC_036173"          "zgc:110796"          
## [127] "wwc1"                 "si:dkey-280e21.3"     "agla"                
## [130] "CABZ01058046.1"       "si:ch73-167i17.6"     "actc1c"              
## [133] "si:ch211-212k18.5"    "crygm1"               "mybpc2a"             
## [136] "hint3"                "eif2b3"               "ddx21"               
## [139] "ckmt2b"               "fuca2"                "cadm4"               
## [142] "pfklb"                "ppp4r3b"              "tmc6b"               
## [145] "dnpep"                "grid2ipb"             "vps9d1"              
## [148] "rbpja"                "atp2a2a"              "smim29"              
## [151] "mrtfab"               "ubap2a"               "scarb2a"             
## [154] "mylz3"                "PDPR"                 "CU570781.3"          
## [157] "ppp2r5b"              "lxn"                  "blcap"               
## [160] "bnip3la"              "appl1"                "egln2"               
## [163] "efl1"                 "si:ch211-244c8.4"     "guca1d"              
## [166] "smad5"                "si:ch211-152c2.3"     "etv4"                
## [169] "rundc3aa"             "fam168a"              "LOC103909284"        
## [172] "actc1b"               "si:ch211-276a17.5"    "FP101914.1"          
## [175] "slco5a1b"             "gas6"                 "zbtb18"              
## [178] "nfia"                 "crb2a"                "clcn4"               
## [181] "si:dkey-111e8.5"      "aamp"                 "mxd1"                
## [184] "si:dkey-172h23.2"     "sybu"                 "nudcd3"              
## [187] "rbpjb"                "dyrk1aa"              "lrfn5b"              
## [190] "ehd3"                 "slc25a22a"            "tmpob"               
## [193] "aldh9a1a.1"           "ch25hl3"              "abca3b"              
## [196] "ccnt1"                "col9a1a"              "fhl2b"               
## [199] "si:dkey-79d12.5"      "dusp5"                "MAPK8IP1 (1 of many)"
## [202] "CZQB01141835.1"       "igfbp5a"              "lin7c"               
## [205] "rps12"                "lenep"                "szrd1"               
## [208] "smn1"                 "rap2ab"               "iars2"               
## [211] "exosc1"               "scamp5a"              "sp9"                 
## [214] "hbbe1.3"              "XLOC_008233"          "gcdhb"               
## [217] "copg2"                "stt3a"                "ccdc92"              
## [220] "ssr1"                 "mrpl44"               "si:dkey-244a7.1"     
## [223] "faxdc2"               "pgam2"                "crata"               
## [226] "oca2"                 "rps27.2"              "RNF208"              
## [229] "srsf7b"               "ndrg3a"               "gata2a"              
## [232] "dnajc5ga"             "paqr7b"               "matn1"               
## [235] "rb1cc1"               "rplp0"                "gria2a"              
## [238] "l3mbtl1a"             "cct2"                 "atn1"                
## [241] "XLOC_013854"          "hsd20b2"              "zgc:165461"          
## [244] "smg8"                 "BX294434.1"           "prx"                 
## [247] "degs1"                "scml4"                "cdc40"               
## [250] "slc25a5"              "pbx1a"                "FQ311908.1"          
## [253] "terfa"                "bhmt"                 "sgsh"                
## [256] "s1pr1"                "grm2a"                "e2f8"                
## [259] "pho"                  "si:dkey-11o15.7"      "zgc:153031"          
## [262] "mpnd"                 "itgb8"                "abi3a"               
## [265] "fam20b"               "arid5b"               "cat"                 
## [268] "ccnd2b"               "sema6ba"              "zgc:101853"          
## [271] "gys1"                 "abhd17ab"             "ndrg1b"              
## [274] "epgn"                 "dnajb1a"              "si:cabz01007794.1"   
## [277] "soul3"                "lypc"                 "sparta"              
## [280] "appb"                 "epn2"                 "FP101882.1"          
## [283] "trmt10c"              "rmdn1"                "pcsk7"               
## [286] "sae1"                 "cdk11b"               "pcdh12"              
## [289] "sall2"                "CABZ01084603.1"       "fosaa"               
## [292] "slc16a3"              "prickle2b"            "drosha"              
## [295] "nlrp16"               "smyd1a"               "desma"               
## [298] "lingo2b"              "rims4"                "tkta"                
## [301] "si:dkey-174m14.3"     "ptpro"                "aacs"                
## [304] "osbpl3a"              "igf2bp3"              "nexmifb"             
## [307] "samd7"                "pam"                  "plekhj1"             
## [310] "si:ch73-362m14.2"     "etf1b"                "fgfbp2b"             
## [313] "si:ch211-149b19.3"    "si:dkey-11o15.5"      "CU633745.1"          
## [316] "psd3l"                "vsnl1b"               "tnxba"               
## [319] "dachc"                "cyp2p6"               "gnb4b"               
## [322] "mkrn1"                "ccni"                 "zgc:56304"           
## [325] "srxn1"                "crabp2a"              "cirbpa"              
## [328] "pdca"                 "si:ch211-69b7.6"      "ipo13"               
## [331] "si:dkey-102c8.3"      "abcf1"                "thrb"                
## [334] "calr3b"               "pygl"                 "CABZ01079427.1"      
## [337] "lhx5"                 "zgc:114045"           "gpd1b"               
## [340] "garem"                "krt1-19d"             "styk1b"              
## [343] "XLOC_018236"          "cdk14"                "bada"                
## [346] "med30"                "epas1b"               "chd6"                
## [349] "pgp"                  "CU659670.1"           "dpp7"                
## [352] "cenpj"                "XLOC_011873"          "dglucy"              
## [355] "hmgcrb"               "pfn2l"                "dr1"                 
## [358] "palm3"                "cd74a"                "tuba8l"              
## [361] "map4k4"               "tmem56b"              "caskb"               
## [364] "slc44a1a"             "ptk2ab"               "ing3"                
## [367] "plekhh1"              "wasf3b"               "ubb"                 
## [370] "vclb"                 "ttc3"                 "slc25a44a"           
## [373] "tubgcp2"              "nfasca"               "tmem182a"            
## [376] "pccb"                 "zgc:174863"           "metap1"              
## [379] "XLOC_007078"          "eevs"                 "tuba8l2"             
## [382] "tspan4a"              "plxna1a"              "fam171a2a"           
## [385] "alad"                 "ncalda"               "tnfaip3"             
## [388] "hbae3"                "spi1a"                "cacng1a"             
## [391] "vps11"                "lactb"                "CU633479.5"          
## [394] "palm2"                "abcg2a"               "si:ch211-180a12.2"   
## [397] "aldoab"               "cadm2b"               "ncam2"               
## [400] "nosip"                "si:dkey-164f24.2"     "ift80"               
## [403] "myl13"                "alx1"                 "CABZ01072952.1"      
## [406] "LOC100330861"         "ldlra"                "klhl13"              
## [409] "cbln1"                "si:ch211-230g15.5"    "ddx47"               
## [412] "map3k14a"             "atox1"                "si:dkey-18j18.3"     
## [415] "LOC100332535"         "rnf44"                "znf367"              
## [418] "retsat"               "fbxl4"                "camta2"              
## [421] "vim"                  "clul1"                "aldob"               
## [424] "ptmaa"                "stk25a"               "wbp2nl"              
## [427] "CABZ01044281.1"       "cldn1"                "prmt8b"              
## [430] "msl1a"                "CABZ01118575.1"       "mpzl1l"              
## [433] "BX908782.2"           "golga5"               "krt92"               
## [436] "ppp1r9a"              "SAMD5"                "PAQR9"               
## [439] "plpp1a"               "scaf4a"               "ndst3"               
## [442] "ascc1"                "si:dkey-247k7.2"      "cdh24a"              
## [445] "smad9"                "im:7145024"           "btbd9"               
## [448] "vkorc1"               "tnnc1b"               "cenpf"               
## [451] "dpp3"                 "il17rc"               "ypel5"               
## [454] "pdgfrl"               "mical3b"              "mboat1"              
## [457] "si:ch1073-157b13.1"   "rnf185"               "ubap2b"              
## [460] "amigo1"               "nckap5l"              "ptprz1b"             
## [463] "zranb1a"              "p4hb"                 "pax6b"               
## [466] "RERG"                 "slc36a4"              "gnrh2"               
## [469] "eef1a1b"              "ckmb"                 "ndnfl"               
## [472] "tmem38a"              "pde6gb"               "kif15"               
## [475] "scai"                 "meis3"                "fndc7a"              
## [478] "col27a1b"             "SHKBP1"               "hoxd3a"              
## [481] "sik3"                 "slc9a6b"              "tnnt3b"              
## [484] "cfl1"                 "CABZ01117348.1"       "sgpp1"               
## [487] "slc1a2b"              "CABZ01069184.1"       "bsdc1"               
## [490] "dkk2"                 "banf1"                "wbp1"                
## [493] "kcnk1a"               "pkdcca"               "nmi"                 
## [496] "CU862020.1"           "si:ch211-241b2.1"     "zgc:171740"          
## [499] "marveld2b"            "atp1b1b"              "rtn4b"               
## [502] "si:dkey-31f5.11"      "nxph1"                "acta1b"              
## [505] "stat2"                "fxyd6l"               "CT025690.1"          
## [508] "si:ch211-108d22.2"    "ca6"                  "oxsr1b"              
## [511] "defbl1"               "gltpa"                "si:dkey-103g5.3"     
## [514] "plxdc2"               "gapdh"                "fxr1"                
## [517] "usp43a"               "celf2"                "pgm3"                
## [520] "slc24a2"              "rnf130"               "hepacama"            
## [523] "zgc:153115"           "zgc:110540"           "tmem43"              
## [526] "tent5ab"              "xpnpep3"              "hbae1.3"             
## [529] "dab1a"                "zgc:153704"           "pard6a"              
## [532] "clptm1l"              "alg6"                 "stat3"               
## [535] "capzb"                "smg6"                 "fut9b"               
## [538] "snrpf"                "mrpl13"               "trim9"               
## [541] "trib2"                "il13ra1"              "rab8a"               
## [544] "gnb2"                 "cpxm1a"               "ndst2a"              
## [547] "ncaldb"               "si:ch73-21g5.7"       "si:ch211-80h18.1"    
## [550] "snx8b"                "prnpa"                "glcea"               
## [553] "nfic"                 "lrfn2b"               "LOC101883507"        
## [556] "CABZ01054965.1"       "cdc6"                 "tcf7l1a"             
## [559] "rint1"                "XLOC_003678"          "slc6a8"              
## [562] "cryba2a"              "FP017274.1"           "cers2a"              
## [565] "grm2b"                "slc19a2"              "kif11"               
## [568] "fam117bb"             "cx23"                 "CABZ01087514.1"      
## [571] "XLOC_000666"          "LO018029.1"           "kif3b"               
## [574] "glceb"                "XLOC_010247"          "herc2"               
## [577] "cx44.2"               "copb2"                "ccdc174"             
## [580] "si:dkey-94e7.2"       "preb"                 "cog6"                
## [583] "eif2b4"               "txlnba"               "rnf165a"             
## [586] "eif4enif1"            "robo4"                "zranb2"              
## [589] "hsdl2"                "mrpl45"               "fbp1a"               
## [592] "osr2"                 "si:dkey-51a16.9"      "esco2"               
## [595] "prss16"               "CABZ01068248.1"       "myh14"               
## [598] "rsu1"                 "vps72b"               "zgc:101040"          
## [601] "tmem198b"             "dcc"                  "gnptg"               
## [604] "znf526"               "si:dkey-239i20.4"     "pik3ip1"             
## [607] "tnrc6c2"              "dop1a"                "aup1"                
## [610] "si:ch211-105j21.9"    "cpeb2"                "zgc:162945"          
## [613] "tmem178b"             "MTERF4"               "CU633479.1"          
## [616] "si:dkey-34d22.1"      "gipr"                 "ppm1db"              
## [619] "rgn"                  "stx6"                 "aanat2"              
## [622] "tbc1d16"              "nectin1a"             "MYO1D"               
## [625] "kras"                 "CABZ01067151.2"       "tap2t"               
## [628] "grk6"                 "crygm2d7"             "ajuba"               
## [631] "spata20"              "LOC100150882"         "nnt2"                
## [634] "prkcbb"               "nccrp1"               "si:ch1073-159d7.13"  
## [637] "ntmt1"                "znf750"               "slc7a8b"             
## [640] "crtac1a"              "mrpl16"               "cthrc1b"             
## [643] "si:dkey-195m11.8"     "si:ch211-117c9.5"     "slc45a1"             
## [646] "aifm5"                "si:ch211-241e1.5"     "cox5b2"              
## [649] "si:dkey-205h13.1"     "cndp2"                "purbb"               
## [652] "lgmn"                 "zbtb12.2"             "pcmtd2"              
## [655] "LOC101884666"         "rasd1"                "lim2.4"              
## [658] "cyp2v1"               "sipa1l2"              "guca1b"              
## [661] "fgf13a"               "fgf14"                "chodl"               
## [664] "vasnb"                "CABZ01037298.1"       "bbs12"               
## [667] "pbrm1"                "cldnb"                "phldb2a"             
## [670] "fam13a"               "prr33"                "abcb5"               
## [673] "guk1b"                "smarca5"              "id1"
```

#### 3.2 Bubble plot of select shared genes

```
downgenes <- sort(c("dcc", "gnrh2", "dyrk1aa", "irs4a", "mbd4", "ndrg3a", "RERG", "s1pr1", "zgc:101853"))

upgenes <- sort(c("ap1ar", "ap1m1", "cenpf", "CR407584.1", "dr1", "fosaa", "larp4ab", "palm2", "palm3", "pcdh12", "pik3r3b", "rab8a", "rims4", "si:ch211-125e6.5", "si:ch211-69b7.6", "si:ch73-21g5.7", "si:dkey-18j18.3", "si:dkey-79d12.5", "slc6a8", "slc7a11", "slc6a6b", "XLOC_000523", "jund"))

sharedgenes <- c(downgenes, upgenes)

#filter to only include genes of interest
bubbleDEG <- kmt5b_hdlbpa %>% dplyr::filter(LLgeneAbbrev %in% sharedgenes)

#find middle of foldchange (to center color scale)
limit <- max(abs(bubbleDEG$log2FoldChange)) * c(-1, 1)

#get color
myheatcolors3 <- brewer.pal(name="RdBu", n=11)

#order genes
bubbleDEG$LLgeneAbbrev <- factor(bubbleDEG$LLgeneAbbrev, levels = sharedgenes)

# create 'bubble plot' to summarize y signatures across x phenotypes
bubbleplot_shared <- ggplot(bubbleDEG, aes(x=LLgeneAbbrev, y=mutation)) + 
  geom_point(aes(size=-log10(padj), color = log2FoldChange)) +
  scale_color_gradientn(colors = myheatcolors3, limit=limit) +
  theme_bw() +
  theme(axis.text.x = element_text(angle = 45, hjust = 1),
        axis.title.x = element_blank(),
        axis.title.y = element_blank())+
  annotate("rect", xmin = 0, xmax = 9.5, ymin = 0.25, ymax = 0.5, 
           alpha=1, fill="#472D7BFF") +
  annotate("rect", xmin = 9.5, xmax = 33, ymin = 0.25, ymax = 0.5, 
           alpha=1, fill="#21908CFF") 

bubbleplot_shared
```

```
#export 800x300 for horizontal
```

#### 3.3 Make scatterplot comparing two mutants

```
#subset to only include comparison, logfc, gene name, and pvalue
scatterdata <- kmt5b_hdlbpa %>% dplyr::select(log2FoldChange, pvaluebound, chrom, mutation, LLgeneAbbrev)

#add column about whether chr is 18 or 6
scatterdata <- scatterdata %>% dplyr::mutate(chrom = replace_na(chrom, "none"))
scatterdata <- scatterdata %>% mutate(chromosome = ifelse(chrom == "chr18", "chromosome 18", ifelse(chrom == "chr6", "chromosome 6", "other chromosome")))

#find duplicates
duplicategenes <- dplyr::filter(scatterdata %>%
  distinct() %>%
  group_by(LLgeneAbbrev, mutation) %>%
  dplyr::count(), n !=1)$LLgeneAbbrev

#remove duplicate rows 
scatterdata <- scatterdata %>% dplyr::filter(!LLgeneAbbrev %in% duplicategenes)

#use pivot wider to make untidy table
scatterdata <- scatterdata %>% distinct() %>%
    pivot_wider(names_from = mutation, values_from  = c(log2FoldChange, pvaluebound), values_fill=NA) 
    
#add rownames
row.names(scatterdata) <- scatterdata$LLgeneAbbrev

#genes to label
genestolabel <- c("si:ch211-125e6.5", "XLOC_000523", "CR847844.1")

#subset to only include labeled genes
myTopHits.labels <- scatterdata %>% dplyr::filter(LLgeneAbbrev %in% genestolabel)

#change order that points are plotted
scatterdata <- scatterdata %>% arrange(match(chromosome, c("other chromosome", "chromosome 6", "chromosome 18")), desc(chromosome))

scatterplot <- ggplot() +
  geom_point(data=scatterdata, aes(x=log2FoldChange_kmt5b, y=log2FoldChange_hdlbpa, color=chromosome, text = paste("Symbol:", LLgeneAbbrev))) + #plot data points 
  geom_hline(yintercept=0, linetype = 'dotted') + #add horizontal line 
  geom_vline(xintercept=0, linetype = 'dotted') + #add vertical line 
  theme_bw() +
  scale_color_manual(values = c("#AADC32FF", "#27AD81FF", "#472D7BFF", "darkgrey"), name = "chromosome") +
  geom_label_repel(data=myTopHits.labels, aes(x=log2FoldChange_kmt5b, y=log2FoldChange_hdlbpa, label=LLgeneAbbrev), force = 1, nudge_y = .5, size = 2.5, max.overlaps = Inf, show.legend = FALSE, color = "black")  + #label selected genes
  ylab("log2 fold change hdlbpa") +
  xlab("log2 fold change kmt5b")

scatterplot
```

```
#export 700x500
```

### 4 Individual analysis of kmt5b

#### 4.1 Make bar plots

```
#import  normalized counts for kmt5b data
kmt5b_genecounts <- read.csv("kmt5b-homvswt3x3_normalized_reads_gene_list.csv")[,2:10]

#pick out a genes to plot
geneofinterest <- c(sharedgenes, "grb10b", "aurkb", "c1ql3a")


#make new columns with mean counts of wt
genecountslogfc <- kmt5b_genecounts
genecountslogfc$kmt5bavg <- rowMeans(genecountslogfc[,7:9])

#divide columns by mean counts of wt
genecountslogfc <- genecountslogfc %>% mutate(across(colnames(genecountslogfc)[4:9], function(x) x/kmt5bavg))

#pull out data for a select gene
generepdata <- genecountslogfc %>% dplyr::filter(LLgeneAbbrev %in% geneofinterest)

#get rid of unnecessary columns
generepdata <- generepdata[,c(1,4:9)]

#pivot longer to make tidy
generepdata <- generepdata %>% pivot_longer(!LLgeneAbbrev, names_to = "sample", values_to = "foldchange")

#add condition information based on sampleName
generepdata$genotype <- generepdata$sample

#substitute condition name based on replicate
generepdata <- generepdata %>% mutate(genotype = str_replace_all(genotype, c("^wt.*"="wt", "^hom.*"="hom")))

#order data on plot
generepdata$genotype <- factor(generepdata$genotype, levels=c("wt", "hom"))

#set colors
colors <- c("#472D7BFF", "#21908CFF") 

#make an empty list
plot_list = list()

#make all the plots
for (z in 1:length(geneofinterest)) {
  genedata2dpf <- generepdata %>% dplyr::filter(LLgeneAbbrev == geneofinterest[z])
  plot <- ggplot(genedata2dpf, aes(fill=genotype, y=foldchange, x=genotype)) + 
    geom_bar(position="dodge", stat="summary") +
    geom_point(position = position_dodge(width = .9)) + 
    labs(title = paste(geneofinterest[z])) +
    ylab("fold change") +
    theme_bw() +
    scale_fill_manual(values = colors)
  plot_list[[z]] <- plot
}

# Save plots to svg Makes a separate file for each plot.
for (i in 1:length(geneofinterest)) {
    file_name = paste("kmt5b_foldchange_", geneofinterest[i], ".svg", sep="")
    svglite(file_name)
    print(plot_list[[i]])
    dev.off()
}
```

#### 4.2 Make volcano plot

```
#subset data to kmt5b
myTopHits.df <- kmt5b_hdlbpa %>% dplyr::filter(mutation == "kmt5b")

#edit padj bound to e-20
myTopHits.df <- myTopHits.df %>% dplyr::mutate(padjbound = ifelse(padj < 1e-20, 1e-20, padj))

#add column about whether chr is 18
myTopHits.df <- myTopHits.df %>% dplyr::mutate(chrom = replace_na(chrom, "none"))
myTopHits.df <- myTopHits.df %>% mutate(chromosome = ifelse(chrom == "chr18", "chromosome 18", "other chromosome"))

#list of insulin genes
myTopHits.insulin <- myTopHits.df %>% dplyr::filter(LLgeneAbbrev %in% c("grb10b", "irs4a", "socs2", "igf2b", "pik3r3b", "raf1b", "rapgef1a", "rhebl1", "pik3cb", "ppp1cbl", "rps6"))
myTopHits.insulin$category <- "insulin"

myTopHits.glycolysis <- myTopHits.df %>% dplyr::filter(LLgeneAbbrev %in% c("adh5", "eno1a", "eno3", "pgk1", "tpila"))
myTopHits.glycolysis$category <- "glycolysis"

myTopHits.cellcycle <- myTopHits.df %>% dplyr::filter(LLgeneAbbrev %in% c("setdb2", "thap1", "anapc2", "cdc37"))
myTopHits.cellcycle$category <- "cell cycle"

myTopHits.synapse <- myTopHits.df %>% dplyr::filter(LLgeneAbbrev %in% c("adora1b", "chata", "gpc2", "pmchl", "syn2b", "syt1a", "tmem163a"))
myTopHits.synapse$category <- "synapse"

myTopHits.translation <- myTopHits.df %>% dplyr::filter(LLgeneAbbrev %in% c("eif3ea", "eif3f", "eif3g", "eif3i", "faua", "rpl10", "rpl13", "rpl18a", "rpl19", "rpl24", "rpl26", "rpl27a", "rpl3", "rpl37", "rpl6", "rpl7", "rpl7a", "rpl8", "rps15", "rps19", "rps2", "rps26", "rps38", "rps3a", "rps4x", "rps5", "rps9", "rpsa", "rplp2", "rplp2l"))
myTopHits.translation$category <- "translation"

#genes to label
myTopHits.labels <- rbind(myTopHits.insulin, myTopHits.glycolysis, myTopHits.cellcycle, myTopHits.synapse, myTopHits.translation)

#change order that points are plotted
myTopHits.labels <- myTopHits.labels %>% arrange(match(category,  c("translation", "glycolysis",  "insulin", "cell cycle", "synapse")), desc(category))

genestolabel <- c("si:dkey-242h9.3", "cd82b", "bcar1", "irs4a", "mbd4", "carmil2", "nars", "cngb1a", "RASGRF1", "mlc1", "grb10b", "slc6a6b", "XLOC_024221", "lmnb2", "opn1lw1", "b3gnt2l", "XLOC_013061", "grb10b", "irs4a", "socs2", "igf2b", "pik3r3b", "raf1b", "rapgef1a", "rhebl1", "pik3cb", "ppp1cbl", "rps6", "adh5", "eno1a", "eno3", "pgk1", "tpila", "setdb2", "thap1", "anapc2", "cdc37", "adora1b", "chata", "gpc2", "pmchl", "syn2b", "syt1a", "tmem163a", "eif3ea", "eif3f", "eif3g", "eif3i")

genestolabel <- c("si:dkey-242h9.3", "cd82b", "bcar1", "irs4a", "mbd4", "carmil2", "nars", "cngb1a", "RASGRF1", "mlc1", "grb10b", "slc6a6b", "XLOC_024221", "lmnb2", "opn1lw1", "b3gnt2l", "XLOC_013061")

#subset to only include labeled genes
myTopHits.labels.all <- myTopHits.df %>% dplyr::filter(LLgeneAbbrev %in% genestolabel)

#make all points other
myTopHits.df <- myTopHits.df %>% dplyr::mutate(mutation = "other")

#make the plot
kmt5b_volcano <- ggplot() +
  annotate("rect", xmin = 1, xmax = 5.5, ymin = -log10(0.05), ymax = 20, 
           alpha=.15,   fill="grey") +
  annotate("rect", xmin = -1, xmax = -5.5, ymin = -log10(0.05), ymax = 20, 
           alpha=.15, fill="grey") +
  geom_point(data=myTopHits.df, aes(y=-log10(padjbound), x=log2FoldChange, shape = chromosome, color = mutation), size=2) +
  geom_point(data=myTopHits.labels, aes(y=-log10(padjbound), x = log2FoldChange, color=category, shape=chromosome),  size=2, show.legend = T) +
  theme_bw() +
  coord_cartesian(xlim = c(-6, 6), ylim = c(-0.5, 20.5), expand = FALSE) +
  ylab("-log10(padj)") + 
  xlab("log2 fold change") +
  geom_label_repel(data=myTopHits.labels.all, aes(x=log2FoldChange, y=-log10(padjbound), label=LLgeneAbbrev), force = 2, nudge_y = -1, size = 2.5, max.overlaps = Inf, show.legend = FALSE, color = "black") + #label selected genes
  theme_bw() +
  scale_color_manual(values = c("#AADC32FF", "#27AD81FF", "#472D7BFF", "darkgrey", "gold", "deepskyblue", "orange", "#D697FF"), name = "pathway") +
  scale_shape_manual(values=c(4, 16)) 

kmt5b_volcano
```

```
#export 700x500
```

#### 4.3 DAVID pathway analysis

Get a list of the top 300 genes dysregulated in kmt5b by padj to put
into DAVID.

```
#select kmt5b genes that aren't on chromosome 18
kmt5b_genesfordavid <- kmt5b_hdlbpa %>% dplyr::filter(mutation == "kmt5b" & chromosomenumber != 18)

#sort by pvalue
kmt5b_genesfordavid <- kmt5b_genesfordavid %>% dplyr::arrange(padj)

#get names of top 300 genes
kmt5b_genesfordavid <- kmt5b_genesfordavid$LLgeneAbbrev[1:300]

#write.csv(kmt5b_genesfordavid, "kmt5b_output/kmt5b_genesfordavid.csv")

# use the 'gost' function from the gprofiler2 package to run GO enrichment analysis
#genes shared between comparisons - both up and downregulated
gost.res.homwtall <- gost(kmt5b_genesfordavid, organism = "drerio", correction_method = "fdr", significant = TRUE, evcodes = TRUE, ordered_query = TRUE)

#export to csv
write.csv(apply(gost.res.homwtall$result,2,as.character), file="kmt5b_output/kmt5b_GO-analysis.csv")

# produce a manhattan plot of enriched GO terms
gostplot(gost.res.homwtall, interactive = T, capped = T)
```

#### 4.4 kmt5b GSEA with C5

```
#we want the zebrafish signatures
dr_gsea <- msigdbr(species = "Danio rerio") #gets all collections/signatures with zebrafish

#look at the categories and subcategories of signatures available
dr_gsea %>% 
  dplyr::distinct(gs_cat, gs_subcat) %>% 
  dplyr::arrange(gs_cat, gs_subcat)
```

```
# choose a specific msigdb collection/subcollection
dr_gsea_c5 <- msigdbr(species = "Danio rerio", category = "C5") %>% # choose  msigdb collection of interest
dplyr::select(gs_name, gene_symbol) #just get the columns corresponding to signature name and gene symbols of genes in each signature 

# pull out data for kmt5b
GSEAgenes <- kmt5b_hdlbpa %>% dplyr::filter(mutation == "kmt5b" & chromosomenumber != 18)
mydata.df.sub <- dplyr::select(GSEAgenes, LLgeneAbbrev, log2FoldChange)

#get rid of duplicates
mydata.df.sub <- mydata.df.sub %>% dplyr::filter(LLgeneAbbrev %in% mydata.df.sub$LLgeneAbbrev[!duplicated(mydata.df.sub$LLgeneAbbrev)])

# construct a named vector
mydata.gsea <- mydata.df.sub$log2FoldChange
names(mydata.gsea) <- as.character(mydata.df.sub$LLgeneAbbrev)
mydata.gsea <- sort(mydata.gsea, decreasing = TRUE)

# run GSEA with C5 in kmt5b
myGSEA.res <- GSEA(mydata.gsea, TERM2GENE=dr_gsea_c5, verbose=FALSE, seed=TRUE)

#convert to DF
myGSEA.df <- as_tibble(myGSEA.res)

#look at top terms
datatable(myGSEA.df, 
          extensions = c('KeyTable', "FixedHeader"), 
          options = list(keys = TRUE, searchHighlight = TRUE, pageLength = 10, lengthMenu = c("10", "25", "50", "100"))) %>%
  formatRound(columns=c(2:10), digits=2)
```

```
#export
#write.csv(myGSEA.df, "kmt5b_output/kmt5b-GSEA-c5.csv")
```

#### 4.5 Make network plot from C5 kmt5b GSEA

```
#import data
myGSEA.df <- read.csv("kmt5b_output/kmt5b-GSEA-c5.csv")[,2:12]

#list of terms to include in network
networkterms <- c("GOCC_CYTOSOLIC_RIBOSOME", "GOCC_RIBOSOMAL_SUBUNIT", "GOBP_CYTOPLASMIC_TRANSLATION", "GOMF_STRUCTURAL_CONSTITUENT_OF_RIBOSOME", "GOCC_RIBOSOME", "GOCC_CYTOSOLIC_LARGE_RIBOSOMAL_SUBUNIT", "GOCC_LARGE_RIBOSOMAL_SUBUNIT", "GOCC_CYTOSOLIC_SMALL_RIBOSOMAL_SUBUNIT", "GOCC_SMALL_RIBOSOMAL_SUBUNIT", "GOCC_POLYSOME", "GOCC_MITOCHONDRIAL_PROTEIN_CONTAINING_COMPLEX", "GOBP_DNA_REPLICATION", "GOBP_DNA_REPAIR", "GOCC_MITOCHONDRIAL_MATRIX", "HP_ABNORMAL_MYELINATION", "GOBP_DNA_DEPENDENT_DNA_REPLICATION", "GOBP_AEROBIC_RESPIRATION", "HP_ABNORMAL_CNS_MYELINATION", "HP_CEREBELLAR_HYPOPLASIA", "GOBP_DOUBLE_STRAND_BREAK_REPAIR", "HP_APLASIA_HYPOPLASIA_OF_THE_CEREBELLUM", "HP_HYPOPLASIA_OF_THE_CORPUS_CALLOSUM", "GOCC_INNER_MITOCHONDRIAL_MEMBRANE_PROTEIN_COMPLEX", "HP_ABNORMALITY_OF_THE_MITOCHONDRION", "GOBP_ATP_SYNTHESIS_COUPLED_ELECTRON_TRANSPORT", "GOBP_OXIDATIVE_PHOSPHORYLATION", "GOMF_ORGANIC_ANION_TRANSMEMBRANE_TRANSPORTER_ACTIVITY", "GOMF_SECONDARY_ACTIVE_TRANSMEMBRANE_TRANSPORTER_ACTIVITY", "GOBP_ANION_TRANSPORT", "GOBP_SODIUM_ION_TRANSMEMBRANE_TRANSPORT", "GOBP_IMPORT_ACROSS_PLASMA_MEMBRANE", "GOBP_MITOTIC_CELL_CYCLE_PHASE_TRANSITION", "GOBP_REGULATION_OF_CELL_CYCLE_PHASE_TRANSITION", "GOBP_CELL_CYCLE_PHASE_TRANSITION")

myGSEA.res.filter <- myGSEA.res
myGSEA.res.filter@result <- myGSEA.res.filter@result %>% dplyr::filter(ID %in% networkterms)

#make network plot
myGSEA.res.filter <- pairwise_termsim(myGSEA.res.filter)
kmt5bnetwork <- emapplot(myGSEA.res.filter, color="NES", categorySize="p.adjust", showCategory=length(networkterms))

#get data of out network plot
kmt5bnetwork <- ggplot_build(kmt5bnetwork)
networkdata <- kmt5bnetwork$plot$data

kmt5b_c5_network <- ggraph(networkdata) + 
  geom_edge_link(alpha=.8, aes_(width=~I(width)), colour='darkgrey') + 
  geom_node_point(aes(colour = color, size=size)) +
  geom_node_text(aes(label=name), repel=TRUE) + 
  theme_void() +
  scale_color_gradientn(colors = myheatcolors3, limit=c(-3.5,3.5), name="NES")

kmt5b_c5_network
```

```
#export 1200x800

#make heatmap
p3 <- heatplot(myGSEA.res, foldChange=mydata.gsea, showCategory=749) + ggplot2::coord_flip()

#get data of out heatmap
p3 <- ggplot_build(p3)
heatmapdata <- p3$plot$data

#filter to only include fold changes > 0.5
genestoinclude <- heatmapdata %>% dplyr::filter(foldChange > 0.5 | foldChange < -0.5)
heatmapdata <- heatmapdata %>% dplyr::filter(Gene %in% genestoinclude$Gene)

#choose which clusters to keep in heatmap
heatmapdata <- heatmapdata %>% dplyr::filter(categoryID %in% c("HP_ABNORMAL_MYELINATION", "HP_ABNORMAL_CNS_MYELINATION", "HP_CEREBELLAR_HYPOPLASIA", "HP_APLASIA_HYPOPLASIA_OF_THE_CEREBELLUM", "HP_HYPOPLASIA_OF_THE_CORPUS_CALLOSUM", "GOBP_MITOTIC_CELL_CYCLE_PHASE_TRANSITION", "GOBP_REGULATION_OF_CELL_CYCLE_PHASE_TRANSITION", "GOBP_CELL_CYCLE_PHASE_TRANSITION"))

#use pivot wider to make untidy table
heatmapdata <- heatmapdata %>% pivot_wider(names_from = categoryID, values_from  = foldChange, values_fill=NA) 

#make matrix
heatmapmatrix <- as.matrix(heatmapdata[,2:9])
rownames(heatmapmatrix) <- heatmapdata$Gene

#turn matrix into 1s and 0s and cluster
heatmapmatrix<-ifelse(abs(heatmapmatrix) > 0,1,0)
heatmapmatrix[is.na(heatmapmatrix)] = 0

#cluster your selected genes
hc <- hclust(as.dist(1-cor(heatmapmatrix, method="spearman")), method="average") #cluster columns by spearman correlation

#turn heatmap data back into tidy table
heatmapdata <- heatmapdata %>% pivot_longer(!Gene, names_to = "categoryID", values_to  = "foldChange") 
    
#change order of data to be plotted
heatmapdata$categoryID <- factor(heatmapdata$categoryID, levels = hc$labels)
heatmapdata$Gene <- factor(heatmapdata$Gene, levels = rev(unique(heatmapdata$Gene)))

#rename dataframe columns
colnames(heatmapdata) <- c("Gene", "categoryID", "log2foldchange")

kmt5b_c5_heatmap <- ggplot(heatmapdata, aes(x = Gene, y = categoryID, fill = log2foldchange)) + 
  geom_tile(color="black") + 
  theme_classic() + 
  scale_fill_continuous_divergingx(palette = 'RdBu', mid = 0, na.value="white") +
  theme(axis.text.x = element_text(angle = 45, vjust = 1, hjust = 1), axis.title.x=element_blank(), axis.title.y=element_blank()) 

kmt5b_c5_heatmap + ggplot2::coord_flip()
```

```
#export 700x2000
```

#### 4.6 Use ZFA anatomy to perform GSEA on kmt5b

```
CNStermsGSEA <- read.csv("CNStermsGSEA.csv")[,2:3]
headtermsGSEA <- read.csv("headtermsGSEA.csv")[,2:3]
fullsetGSEA <- read.csv("fullsetGSEA.csv")[,2:3]

# Pull out just the columns corresponding to gene symbols and LogFC for at least one pairwise comparison for the enrichment analysis
GSEAgenes <- kmt5b_hdlbpa %>% dplyr::filter(mutation == "kmt5b" & chromosomenumber != 18)
mydata.df.sub <- dplyr::select(GSEAgenes, LLgeneAbbrev, log2FoldChange)

#get rid of duplicates
mydata.df.sub <- mydata.df.sub %>% dplyr::filter(LLgeneAbbrev %in% mydata.df.sub$LLgeneAbbrev[!duplicated(mydata.df.sub$LLgeneAbbrev)])

# construct a named vector
mydata.gsea <- mydata.df.sub$log2FoldChange
names(mydata.gsea) <- as.character(mydata.df.sub$LLgeneAbbrev)
mydata.gsea <- sort(mydata.gsea, decreasing = TRUE)

#options for doing GSEA using ZFA
#CNStermsGSEA
#headtermsGSEA
#fullsetGSEA

# run GSEA with CNS terms 
myGSEA.res <- GSEA(mydata.gsea, TERM2GENE=CNStermsGSEA, verbose=FALSE, seed=TRUE, minGSSize = 80, pvalueCutoff = 1)
myGSEA.df <- as_tibble(myGSEA.res@result)


# view results as an interactive table
datatable(myGSEA.df, 
          extensions = c('KeyTable', "FixedHeader"), 
          options = list(keys = TRUE, searchHighlight = TRUE, pageLength = 10, lengthMenu = c("10", "25", "50", "100"))) %>%
  formatRound(columns=c(2:10), digits=2)
```

The only term that is significant for head terms is lens. No terms
are significant for CNS terms.

#### 4.7 Use single cell markers from 5 dpf zebrafish brain to perform GSEA on kmt5b data

```
#import single cell markers
singlecellmarkers <- read.csv("zf5dpf_markersforGSEA.csv")

#select relevant columns
singlecellmarkers <- singlecellmarkers %>% dplyr::select(cluster.description, gene)

# Pull out just the columns corresponding to gene symbols and LogFC
GSEAgenes <- kmt5b_hdlbpa %>% dplyr::filter(mutation == "kmt5b" & chromosomenumber != 18)
mydata.df.sub <- dplyr::select(GSEAgenes, LLgeneAbbrev, log2FoldChange)

#get rid of duplicates
mydata.df.sub <- mydata.df.sub %>% dplyr::filter(LLgeneAbbrev %in% mydata.df.sub$LLgeneAbbrev[!duplicated(mydata.df.sub$LLgeneAbbrev)])

# construct a named vector
mydata.gsea <- mydata.df.sub$log2FoldChange
names(mydata.gsea) <- as.character(mydata.df.sub$LLgeneAbbrev)
mydata.gsea <- sort(mydata.gsea, decreasing = TRUE)

# run GSEA with single cell markers in 6dpf 23bp
myGSEA.res <- GSEA(mydata.gsea, TERM2GENE=singlecellmarkers, verbose=FALSE, seed=TRUE, minGSSize = 80)
myGSEA.df <- as_tibble(myGSEA.res@result)

#export
#write.csv(myGSEA.df, file="kmt5b_output/kmt5b-singlecellGSEA.csv")

#import
myGSEA.df <- read.csv(file="kmt5b_output/kmt5b-singlecellGSEA.csv")[,2:12]

datatable(myGSEA.df, 
          extensions = c('KeyTable', "FixedHeader"), 
          options = list(keys = TRUE, searchHighlight = TRUE, pageLength = 10, lengthMenu = c("10", "25", "50", "100"))) %>%
  formatRound(columns=c(2:10), digits=2)
```

```
#make network plot
myGSEA.res <- pairwise_termsim(myGSEA.res)
kmt5bnetwork <- emapplot(myGSEA.res, color="NES", categorySize="p.adjust", showCategory = 32)

#get data of out network plot
kmt5bnetwork <- ggplot_build(kmt5bnetwork)
networkdata <- kmt5bnetwork$plot$data

#make network plot
kmt5b_singlecell_network <- ggraph(networkdata) + 
  geom_edge_link(alpha=.8, aes_(width=~I(width)), colour='darkgrey') + 
  geom_node_point(aes(colour = color, size=size)) +
  geom_node_text(aes(label=name), repel=TRUE) + 
  theme_void() +
  scale_color_gradientn(colors = myheatcolors3, limit=c(-3.5,3.5), name="NES")

kmt5b_singlecell_network
```

```
#export 700x500

#make heatmap
p3 <- heatplot(myGSEA.res, foldChange=mydata.gsea, showCategory = 33) + ggplot2::coord_flip()

#get data of out heatmap
p3 <- ggplot_build(p3)
heatmapdata <- p3$plot$data

#filter to only include fold changes > 0.5
genestoinclude <- heatmapdata %>% dplyr::filter(foldChange > 0.5 | foldChange < -0.5)
heatmapdata <- heatmapdata %>% dplyr::filter(Gene %in% genestoinclude$Gene)

#filter to only include clusters to go in heatmap
heatmapdata <- heatmapdata %>% dplyr::filter(categoryID %in% c("diencephalon (tuberculum; gabaergic)", "granule cells", "hindbrain", "hypothalamus #3", "mid-hind boundary (gabaergic) #1", "mid-hind boundary (gabaergic) #2", "neurons #4", "neurons (ganglion)", "neurons (glutamatergic) #1", "neurons (glutamatergic) #2", "neurons (midbrain?, glutamatergic)", "olfactory bulb", "oligodendrocytes #1", "optic tectum (gabaergic) #1", "progenitors #1", "progenitors #2", "progenitors #3", "progenitors (cycling)", "progenitors/neurons (differentiating)", "ventral forebrain (dienc, hyp, poa; gabaergic) #1"))

#use pivot wider to make untidy table
heatmapdata <- heatmapdata %>% pivot_wider(names_from = categoryID, values_from  = foldChange, values_fill=NA) 

#make matrix
heatmapmatrix <- as.matrix(heatmapdata[,2:21])
rownames(heatmapmatrix) <- heatmapdata$Gene

#turn matrix into 1s and 0s and cluster
heatmapmatrix<-ifelse(abs(heatmapmatrix) > 0,1,0)
heatmapmatrix[is.na(heatmapmatrix)] = 0

 #cluster rows by pearson correlation
hc <- hclust(as.dist(1-cor(heatmapmatrix, method="spearman")), method="average") #cluster columns by spearman correlation

#turn heatmap data back into tidy table
heatmapdata <- heatmapdata %>% pivot_longer(!Gene, names_to = "categoryID", values_to  = "foldChange") 
    
#change order of data to be plotted
heatmapdata$categoryID <- factor(heatmapdata$categoryID, levels = hc$labels)
heatmapdata$Gene <- factor(heatmapdata$Gene, levels = rev(unique(heatmapdata$Gene)))

#rename dataframe columns
colnames(heatmapdata) <- c("Gene", "categoryID", "log2foldchange")

singlecellheatmap <- ggplot(heatmapdata, aes(x = Gene, y = categoryID, fill = log2foldchange)) + 
  geom_tile(color="black") + 
  theme_classic() + 
  scale_fill_continuous_divergingx(palette = 'RdBu', mid = 0, na.value="white") +
  theme(axis.text.x = element_text(angle = 45, vjust = 1, hjust = 1), axis.title.x=element_blank(), axis.title.y=element_blank()) 

singlecellheatmap + ggplot2::coord_flip()
```

```
#export 800x2500


#repeat 6dpf GSEA but with neuronal clusters only
nonneuronalclusters <- c("#N/A", "cartilage #2", "cartilage #1", "cartilage #3", "cornea", "cranial ganglion", "cranial ganglion (vagal) #1", "cranial ganglion (vagal) #2", "dermal bone", "epidermis (progenitors)", "epithelium (cornea, progenitors)", "erythrocytes", "eye (cornea), cartilage", "eye (cornea, epithelium)", "eye (non retina)", "glia (eye)", "lens #1", "lens #2", "muscle", "neural crest derived (pigment/xanthophore)", "neutrophils", "otic vesicle", "otic/lateral line", "pigment cell (iridophore)", "retina (amacrine, gabaergic)", "retina (cone bipolar cells)", "retina (cones)", "retina (muller glia)", "retina (photoreceptor precursor cells)", "retina (RGC)", "retina (rods)", "retina (RPE differentiation)", "retina (RPE)", "retina neurons (horizontal?)", "rostral blood island (myeloid) #1", "rostral blood island (myeloid) #2", "pharangeal arch/pectoral fin (mesoderm)")

#make GSEA set for CNS single cell data
singlecellmarkers_CNS <- singlecellmarkers %>% dplyr::filter(! cluster.description %in% nonneuronalclusters)

# run GSEA with single cell markers in 6dpf 23bp
myGSEA.res <- GSEA(mydata.gsea, TERM2GENE=singlecellmarkers_CNS, verbose=FALSE, seed=TRUE, minGSSize = 80)
myGSEA.df <- as_tibble(myGSEA.res@result)

#export
#write.csv(myGSEA.df, file="kmt5b_output/kmt5b-singlecellGSEA-CNS.csv")

#import
myGSEA.df <- read.csv(file="kmt5b_output/kmt5b-singlecellGSEA-CNS.csv")[,2:12]

#view data table
datatable(myGSEA.df, 
          extensions = c('KeyTable', "FixedHeader"), 
          options = list(keys = TRUE, searchHighlight = TRUE, pageLength = 10, lengthMenu = c("10", "25", "50", "100"))) %>%
  formatRound(columns=c(2:10), digits=2)
```

```
#make network plot
myGSEA.res <- pairwise_termsim(myGSEA.res)
kmt5bnetwork <- emapplot(myGSEA.res, color="NES", categorySize="p.adjust", showCategory = 33)

#get data of out network plot
kmt5bnetwork <- ggplot_build(kmt5bnetwork)
networkdata <- kmt5bnetwork$plot$data

#make new network plot
kmt5b_singlecellCNS_network <- ggraph(networkdata) + 
  geom_edge_link(alpha=.8, aes_(width=~I(width)), colour='darkgrey') + 
  geom_node_point(aes(colour = color, size=size)) +
  geom_node_text(aes(label=name), repel=TRUE) + 
  theme_void() +
  scale_color_gradientn(colors = myheatcolors3, limit=c(-3.5,3.5), name="NES")

kmt5b_singlecellCNS_network
```

```
#export 700x500
```

### 5 Individual analysis of hdlbpa

#### 5.1 Make bar plots

```
#pick out a genes to plot
geneofinterest <- c(sharedgenes, "mki67")

#import  normalized counts for hdlbpa
hdlbpa_genecounts <- read.csv("hdlbpa-homvswt_normalized_reads_gene_list.csv")[,2:12]

#make new columns with mean counts of wt
genecountslogfc <- hdlbpa_genecounts
genecountslogfc$hdlbpaavg <- rowMeans(genecountslogfc[,8:11])

#divide columns by mean counts of wt
genecountslogfc <- genecountslogfc %>% mutate(across(colnames(genecountslogfc)[4:11], function(x) x/hdlbpaavg))

#pull out data for a select gene
generepdata <- genecountslogfc %>% dplyr::filter(LLgeneAbbrev %in% geneofinterest)

#get rid of unnecessary columns
generepdata <- generepdata[,c(1,4:11)]

#pivot longer to make tidy
generepdata <- generepdata %>% pivot_longer(!LLgeneAbbrev, names_to = "sample", values_to = "foldchange")

#add condition information based on sampleName
generepdata$genotype <- generepdata$sample

#substitute condition name based on replicate
generepdata <- generepdata %>% mutate(genotype = str_replace_all(genotype, c("^wt.*"="wt", "^hom.*"="hom")))

#order data on plot
generepdata$genotype <- factor(generepdata$genotype, levels=c("wt", "hom"))

#set colors
colors <- c("#472D7BFF", "#21908CFF") 

#make an empty list
plot_list = list()

#make all the plots
for (z in 1:length(geneofinterest)) {
  genedata2dpf <- generepdata %>% dplyr::filter(LLgeneAbbrev == geneofinterest[z])
  plot <- ggplot(genedata2dpf, aes(fill=genotype, y=foldchange, x=genotype)) + 
    geom_bar(position="dodge", stat="summary") +
    geom_point(position = position_dodge(width = .9)) + 
    labs(title = paste(geneofinterest[z])) +
    ylab("fold change") +
    theme_bw() +
    scale_fill_manual(values = colors)
  plot_list[[z]] <- plot
}

# Save plots to svg Makes a separate file for each plot.
for (i in 1:length(geneofinterest)) {
    file_name = paste("hdlbpa_foldchange_", geneofinterest[i], ".svg", sep="")
    svglite(file_name)
    print(plot_list[[i]])
    dev.off()
}
```

#### 5.2 Make volcano plot

```
#subset data to hdlbpa
myTopHits.df <- kmt5b_hdlbpa %>% dplyr::filter(mutation == "hdlbpa")

#add column about whether chr is 6
myTopHits.df <- myTopHits.df %>% dplyr::mutate(chrom = replace_na(chrom, "none"))
myTopHits.df <- myTopHits.df %>% mutate(chromosome = ifelse(chrom == "chr6", "chromosome 6", "other chromosome"))

myTopHits.cellcycle <- myTopHits.df %>% dplyr::filter(LLgeneAbbrev %in% c("thap1", "cdc40", "esco2", "mki67", "nusap1", "rbbp8"))
myTopHits.cellcycle$category <- "cell cycle"

#list of insulin genes
myTopHits.insulin <- myTopHits.df %>% dplyr::filter(LLgeneAbbrev %in% c("pik3r3b", "pik3r6a", "irs4a", "insl5a"))
myTopHits.insulin$category <- "insulin"

myTopHits.collagen <- myTopHits.df %>% dplyr::filter(LLgeneAbbrev %in% c("col4a3", "col9a3", "col11a2", "fbn2b", "matn1", "prelp", "si:dkey-239b22.2"))
myTopHits.collagen$category <- "collagen/ECM"

myTopHits.glycerophospholipid <- myTopHits.df %>% dplyr::filter(LLgeneAbbrev %in% c("gpd1b", "mboat1", "ppap2d", "pla2g15", "plpp1a", "proca1"))
myTopHits.glycerophospholipid$category <- "glycerophospholipid metabolism"

#genes to label
myTopHits.labels <- rbind(myTopHits.insulin, myTopHits.collagen, myTopHits.cellcycle, myTopHits.glycerophospholipid)

genestolabel <- c("hdlbpa", "ctsll", "p4hb", "stx8", "CR407584.1", "ptprnb", "ecrg4b", "BX511311.3", "gpr27", "mybphb", "XLOC_036545", "stat2", "nupr1b", "BACZ01084081.1", "plek2", "zgc:162999", "CABZ01084081.1", "si:dkey-61p9.11", "insl5a")

#subset to only include labeled genes
myTopHits.labels.all <- myTopHits.df %>% dplyr::filter(LLgeneAbbrev %in% genestolabel)

#make all points other
myTopHits.df <- myTopHits.df %>% dplyr::mutate(mutation = "other")

#make the plot
hdlbpa_volcano <- ggplot() +
  annotate("rect", xmin = 1, xmax = 6, ymin = -log10(0.05), ymax = 20, 
           alpha=.15,   fill="grey") +
  annotate("rect", xmin = -1, xmax = -6, ymin = -log10(0.05), ymax = 20, 
           alpha=.15, fill="grey") +
  geom_point(data=myTopHits.df, aes(y=-log10(padjbound), x=log2FoldChange, shape = chromosome, color = mutation, text = paste("Symbol:", LLgeneAbbrev)), size=2) +
  geom_point(data=myTopHits.labels, aes(y=-log10(padjbound), x = log2FoldChange, color=category, shape=chromosome),  size=2, show.legend = T) +
  theme_bw() +
  coord_cartesian(xlim = c(-6.5, 6.5), ylim = c(-0.5, 10.5), expand = FALSE) +
  ylab("-log10(padj)") + 
  xlab("log2 fold change") +
  geom_label_repel(data=myTopHits.labels.all, aes(x=log2FoldChange, y=-log10(padjbound), label=LLgeneAbbrev), force = 2, nudge_y = -1, size = 2.5, max.overlaps = Inf, show.legend = FALSE, color = "black") + #label selected genes
  theme_bw() +
  scale_color_manual(values = c("#AADC32FF", "#27AD81FF", "#472D7BFF","gold", "darkgrey", "deepskyblue", "orange", "#D697FF"), name = "pathway") +
  scale_shape_manual(values=c(4, 16)) 

hdlbpa_volcano
```

```
#export 700x500
```

#### 5.3 DAVID pathway analysis

Get a list of the top 300 genes dysregulated in hdlbpa by padj to put
into DAVID.

```
#select hdlbpa genes that aren't on chromosome 6
hdlbpa_genesfordavid <- kmt5b_hdlbpa %>% dplyr::filter(mutation == "hdlbpa" & chromosomenumber != 6)

#sort by pvalue
hdlbpa_genesfordavid <- hdlbpa_genesfordavid %>% dplyr::arrange(padj)

#get names of top 300 genes
hdlbpa_genesfordavid <- hdlbpa_genesfordavid$LLgeneAbbrev[1:300]

#write.csv(hdlbpa_genesfordavid, "kmt5b_output/hdlbpa_genesfordavid.csv")

# use the 'gost' function from the gprofiler2 package to run GO enrichment analysis
#genes shared between comparisons - both up and downregulated
gost.res.homwtall <- gost(hdlbpa_genesfordavid, organism = "drerio", correction_method = "fdr", significant = TRUE, evcodes = TRUE, ordered_query = TRUE)

#export to csv
#write.csv(apply(gost.res.homwtall$result,2,as.character), file="kmt5b_output/hdlbpa_GO-analysis.csv")

# produce a manhattan plot of enriched GO terms
gostplot(gost.res.homwtall, interactive = T, capped = T)
```

#### 5.4 hdlbpa GSEA with C5

```
# choose a specific msigdb collection/subcollection
dr_gsea_c5 <- msigdbr(species = "Danio rerio", category = "C5") %>% # choose  msigdb collection of interest
  dplyr::select(gs_name, gene_symbol) #just get the columns corresponding to signature name and gene symbols of genes in each signature 

# pull out data for hdlbpa
GSEAgenes <- kmt5b_hdlbpa %>% dplyr::filter(mutation == "hdlbpa" & chromosomenumber != 6)
mydata.df.sub <- dplyr::select(GSEAgenes, LLgeneAbbrev, log2FoldChange)

#get rid of duplicates
mydata.df.sub <- mydata.df.sub %>% dplyr::filter(LLgeneAbbrev %in% mydata.df.sub$LLgeneAbbrev[!duplicated(mydata.df.sub$LLgeneAbbrev)])

# construct a named vector
mydata.gsea <- mydata.df.sub$log2FoldChange
names(mydata.gsea) <- as.character(mydata.df.sub$LLgeneAbbrev)
mydata.gsea <- sort(mydata.gsea, decreasing = TRUE)

# run GSEA with C5 in kmt5b
myGSEA.res <- GSEA(mydata.gsea, TERM2GENE=dr_gsea_c5, verbose=FALSE, seed=TRUE)

#convert to DF
myGSEA.df <- as_tibble(myGSEA.res)

#look at top terms
datatable(myGSEA.df, 
          extensions = c('KeyTable', "FixedHeader"), 
          options = list(keys = TRUE, searchHighlight = TRUE, pageLength = 10, lengthMenu = c("10", "25", "50", "100"))) %>%
  formatRound(columns=c(2:10), digits=2)
```

```
#export
#write.csv(myGSEA.df, "kmt5b_output/hdlbpa-GSEA-c5.csv")
```

#### 5.5 Make network plot from C5 hdlbpa GSEA

```
#import data
myGSEA.df <- read.csv("kmt5b_output/hdlbpa-GSEA-c5.csv")[,2:12]

#list of terms to include in network
networkterms <- c("GOCC_CHROMOSOME_CENTROMERIC_REGION", "GOBP_MITOTIC_NUCLEAR_DIVISION", "GOCC_APICAL_PLASMA_MEMBRANE", "GOBP_MITOTIC_SISTER_CHROMATID_SEGREGATION  ", "GOCC_CONDENSED_CHROMOSOME", "GOCC_CONDENSED_CHROMOSOME_CENTROMERIC_REGION", "GOBP_CELL_DIVISION", "GOMF_CARBOHYDRATE_TRANSMEMBRANE_TRANSPORTER_ACTIVITY", "GOMF_SUGAR_TRANSMEMBRANE_TRANSPORTER_ACTIVITY")

myGSEA.res.filter <- myGSEA.res
myGSEA.res.filter@result <- myGSEA.res.filter@result %>% dplyr::filter(ID %in% networkterms)

#make network plot
myGSEA.res.filter <- pairwise_termsim(myGSEA.res.filter)
hdlbpanetwork <- emapplot(myGSEA.res.filter, color="NES", categorySize="p.adjust", showCategory=length(networkterms))

#get data of out network plot
hdlbpanetwork <- ggplot_build(hdlbpanetwork)
networkdata <- hdlbpanetwork$plot$data

hdlbpa_c5_network <- ggraph(networkdata) + 
  geom_edge_link(alpha=.8, aes_(width=~I(width)), colour='darkgrey') + 
  geom_node_point(aes(colour = color, size=size)) +
  geom_node_text(aes(label=name), repel=TRUE) + 
  theme_void() +
  scale_color_gradientn(colors = myheatcolors3, limit=c(-3.5,3.5), name="NES")

hdlbpa_c5_network
```

```
#export 700x500

#make heatmap
p3 <- heatplot(myGSEA.res, foldChange=mydata.gsea, showCategory=749) + ggplot2::coord_flip()

#get data of out heatmap
p3 <- ggplot_build(p3)
heatmapdata <- p3$plot$data

#filter to only include fold changes > 0.5
genestoinclude <- heatmapdata %>% dplyr::filter(foldChange > 0.5 | foldChange < -0.5)
heatmapdata <- heatmapdata %>% dplyr::filter(Gene %in% genestoinclude$Gene)

#choose which clusters to keep in heatmap
heatmapdata <- heatmapdata %>% dplyr::filter(categoryID %in% c("GOCC_CHROMOSOME_CENTROMERIC_REGION", "GOBP_MITOTIC_NUCLEAR_DIVISION", "GOCC_APICAL_PLASMA_MEMBRANE", "GOBP_MITOTIC_SISTER_CHROMATID_SEGREGATION ", "GOCC_CONDENSED_CHROMOSOME", "GOCC_CONDENSED_CHROMOSOME_CENTROMERIC_REGION", "GOBP_CELL_DIVISION", "GOMF_CARBOHYDRATE_TRANSMEMBRANE_TRANSPORTER_ACTIVITY", "GOMF_SUGAR_TRANSMEMBRANE_TRANSPORTER_ACTIVITY"))

#use pivot wider to make untidy table
heatmapdata <- heatmapdata %>% pivot_wider(names_from = categoryID, values_from  = foldChange, values_fill=NA) 

#make matrix
heatmapmatrix <- as.matrix(heatmapdata[,2:9])
rownames(heatmapmatrix) <- heatmapdata$Gene

#turn matrix into 1s and 0s and cluster
heatmapmatrix<-ifelse(abs(heatmapmatrix) > 0,1,0)
heatmapmatrix[is.na(heatmapmatrix)] = 0
#cluster rows by pearson correlation
hc <- hclust(as.dist(1-cor(heatmapmatrix, method="spearman")), method="average") #cluster columns by spearman correlation

#turn heatmap data back into tidy table
heatmapdata <- heatmapdata %>% pivot_longer(!Gene, names_to = "categoryID", values_to  = "foldChange") 

#change order of data to be plotted
heatmapdata$categoryID <- factor(heatmapdata$categoryID, levels = hc$labels)
heatmapdata$Gene <- factor(heatmapdata$Gene, levels = unique(heatmapdata$Gene))

#rename dataframe columns
colnames(heatmapdata) <- c("Gene", "categoryID", "log2foldchange")

hdlbpa_c5_heatmap <- ggplot(heatmapdata, aes(x = Gene, y = categoryID, fill = log2foldchange)) + 
  geom_tile(color="black") + 
  theme_classic() + 
  scale_fill_continuous_divergingx(palette = 'RdBu', mid = 0, na.value="white") +
  theme(axis.text.x = element_text(angle = 45, vjust = 1, hjust = 1), axis.title.x=element_blank(), axis.title.y=element_blank()) 

hdlbpa_c5_heatmap + ggplot2::coord_flip()
```

```
#export 400x1200
```

#### 5.6 Use ZFA anatomy to perform GSEA on hdlbpa

```
# Pull out just the columns corresponding to gene symbols and LogFC for at least one pairwise comparison for the enrichment analysis
GSEAgenes <- kmt5b_hdlbpa %>% dplyr::filter(mutation == "hdlbpa" & chromosomenumber != 6)
mydata.df.sub <- dplyr::select(GSEAgenes, LLgeneAbbrev, log2FoldChange)

#get rid of duplicates
mydata.df.sub <- mydata.df.sub %>% dplyr::filter(LLgeneAbbrev %in% mydata.df.sub$LLgeneAbbrev[!duplicated(mydata.df.sub$LLgeneAbbrev)])

# construct a named vector
mydata.gsea <- mydata.df.sub$log2FoldChange
names(mydata.gsea) <- as.character(mydata.df.sub$LLgeneAbbrev)
mydata.gsea <- sort(mydata.gsea, decreasing = TRUE)

#options for doing GSEA using ZFA
#CNStermsGSEA
#headtermsGSEA
#fullsetGSEA

# run GSEA with CNS terms 
myGSEA.res <- GSEA(mydata.gsea, TERM2GENE=CNStermsGSEA, verbose=FALSE, seed=TRUE, minGSSize = 80, pvalueCutoff = 1)
myGSEA.df <- as_tibble(myGSEA.res@result)

#export file
#write.csv(myGSEA.df, file="kmt5b_output/hdlbpa_CNS_GSEA.csv")


# view results as an interactive table
datatable(myGSEA.df, 
          extensions = c('KeyTable', "FixedHeader"), 
          options = list(keys = TRUE, searchHighlight = TRUE, pageLength = 10, lengthMenu = c("10", "25", "50", "100"))) %>%
  formatRound(columns=c(2:10), digits=2)
```

Head terms with positive enrichment scores are lens, eye
photoreceptor cell, and ciliary marginal zone. No terms are significant
for CNS terms.

#### 5.7 Use single cell markers from 5 dpf zebrafish brain to perform GSEA on hdlbpa data

```
# Pull out just the columns corresponding to gene symbols and LogFC
GSEAgenes <- kmt5b_hdlbpa %>% dplyr::filter(mutation == "hdlbpa" & chromosomenumber != 6)
mydata.df.sub <- dplyr::select(GSEAgenes, LLgeneAbbrev, log2FoldChange)

#get rid of duplicates
mydata.df.sub <- mydata.df.sub %>% dplyr::filter(LLgeneAbbrev %in% mydata.df.sub$LLgeneAbbrev[!duplicated(mydata.df.sub$LLgeneAbbrev)])

# construct a named vector
mydata.gsea <- mydata.df.sub$log2FoldChange
names(mydata.gsea) <- as.character(mydata.df.sub$LLgeneAbbrev)
mydata.gsea <- sort(mydata.gsea, decreasing = TRUE)

# run GSEA with single cell markers in 6dpf 23bp
myGSEA.res <- GSEA(mydata.gsea, TERM2GENE=singlecellmarkers, verbose=FALSE, seed=TRUE, minGSSize = 80)
myGSEA.df <- as_tibble(myGSEA.res@result)

#export
#write.csv(myGSEA.df, file="kmt5b_output/hdlbpa-singlecellGSEA.csv")

#import
myGSEA.df <- read.csv(file="kmt5b_output/hdlbpa-singlecellGSEA.csv")[,2:12]

datatable(myGSEA.df, 
          extensions = c('KeyTable', "FixedHeader"), 
          options = list(keys = TRUE, searchHighlight = TRUE, pageLength = 10, lengthMenu = c("10", "25", "50", "100"))) %>%
  formatRound(columns=c(2:10), digits=2)
```

```
#make network plot
myGSEA.res <- pairwise_termsim(myGSEA.res)
hdlbpanetwork <- emapplot(myGSEA.res, color="NES", categorySize="p.adjust", showCategory = 11)

#get data of out network plot
hdlbpanetwork <- ggplot_build(hdlbpanetwork)
networkdata <- hdlbpanetwork$plot$data

#make network plot
hdlbpa_singlecell_network <- ggraph(networkdata) + 
  geom_edge_link(alpha=.8, aes_(width=~I(width)), colour='darkgrey') + 
  geom_node_point(aes(colour = color, size=size)) +
  geom_node_text(aes(label=name), repel=TRUE) + 
  theme_void() +
  scale_color_gradientn(colors = myheatcolors3, limit=c(-3,3), name="NES")

hdlbpa_singlecell_network
```

```
#export 700x500

#repeat 6dpf GSEA but with neuronal clusters only
nonneuronalclusters <- c("#N/A", "cartilage #2", "cartilage #1", "cartilage #3", "cornea", "cranial ganglion", "cranial ganglion (vagal) #1", "cranial ganglion (vagal) #2", "dermal bone", "epidermis (progenitors)", "epithelium (cornea, progenitors)", "erythrocytes", "eye (cornea), cartilage", "eye (cornea, epithelium)", "eye (non retina)", "glia (eye)", "lens #1", "lens #2", "muscle", "neural crest derived (pigment/xanthophore)", "neutrophils", "otic vesicle", "otic/lateral line", "pigment cell (iridophore)", "retina (amacrine, gabaergic)", "retina (cone bipolar cells)", "retina (cones)", "retina (muller glia)", "retina (photoreceptor precursor cells)", "retina (RGC)", "retina (rods)", "retina (RPE differentiation)", "retina (RPE)", "retina neurons (horizontal?)", "rostral blood island (myeloid) #1", "rostral blood island (myeloid) #2", "pharangeal arch/pectoral fin (mesoderm)")

#make GSEA set for CNS single cell data
singlecellmarkers_CNS <- singlecellmarkers %>% dplyr::filter(! cluster.description %in% nonneuronalclusters)

# run GSEA with single cell markers in 6dpf 23bp
myGSEA.res <- GSEA(mydata.gsea, TERM2GENE=singlecellmarkers_CNS, verbose=FALSE, seed=TRUE, minGSSize = 80)
myGSEA.df <- as_tibble(myGSEA.res@result)

#export
#write.csv(myGSEA.df, file="kmt5b_output/hdlbpa-singlecellGSEA-CNS.csv")

#import
myGSEA.df <- read.csv(file="kmt5b_output/hdlbpa-singlecellGSEA-CNS.csv")[,2:12]

#view data table
datatable(myGSEA.df, 
          extensions = c('KeyTable', "FixedHeader"), 
          options = list(keys = TRUE, searchHighlight = TRUE, pageLength = 10, lengthMenu = c("10", "25", "50", "100"))) %>%
  formatRound(columns=c(2:10), digits=2)
```

```
#make network plot
myGSEA.res <- pairwise_termsim(myGSEA.res)
hdlbpanetwork <- emapplot(myGSEA.res, color="NES", categorySize="p.adjust", showCategory = 7)

#get data of out network plot
hdlbpanetwork <- ggplot_build(hdlbpanetwork)
networkdata <- hdlbpanetwork$plot$data

#make new network plot
hdlbpa_singlecellCNS_network <- ggraph(networkdata) + 
  geom_edge_link(alpha=.8, aes_(width=~I(width)), colour='darkgrey') + 
  geom_node_point(aes(colour = color, size=size)) +
  geom_node_text(aes(label=name), repel=TRUE) + 
  theme_void() +
  scale_color_gradientn(colors = myheatcolors3, limit=c(-3,3), name="NES")

hdlbpa_singlecellCNS_network
```

```
#export 700x500

#make heatmap
p3 <- heatplot(myGSEA.res, foldChange=mydata.gsea, showCategory = 7) + ggplot2::coord_flip()

#get data of out heatmap
p3 <- ggplot_build(p3)
heatmapdata <- p3$plot$data

#filter to only include fold changes > 0.5
genestoinclude <- heatmapdata %>% dplyr::filter(foldChange > 0.5 | foldChange < -0.5)
heatmapdata <- heatmapdata %>% dplyr::filter(Gene %in% genestoinclude$Gene)

#use pivot wider to make untidy table
heatmapdata <- heatmapdata %>% pivot_wider(names_from = categoryID, values_from  = foldChange, values_fill=NA) 

#make matrix
heatmapmatrix <- as.matrix(heatmapdata[,2:8])
rownames(heatmapmatrix) <- heatmapdata$Gene

#turn matrix into 1s and 0s and cluster
heatmapmatrix<-ifelse(abs(heatmapmatrix) > 0,1,0)
heatmapmatrix[is.na(heatmapmatrix)] = 0

#cluster your selected genes
hr <- hclust(as.dist(1-cor(t(heatmapmatrix), method="pearson")), method="complete") #cluster rows by pearson correlation
hc <- hclust(as.dist(1-cor(heatmapmatrix, method="spearman")), method="average") #cluster columns by spearman correlation

#turn heatmap data back into tidy table
heatmapdata <- heatmapdata %>% pivot_longer(!Gene, names_to = "categoryID", values_to  = "foldChange") 
    
#change order of data to be plotted
heatmapdata$categoryID <- factor(heatmapdata$categoryID, levels = hc$labels)
heatmapdata$Gene <- factor(heatmapdata$Gene, levels = unique(heatmapdata$Gene))

#rename dataframe columns
colnames(heatmapdata) <- c("Gene", "categoryID", "log2foldchange")

singlecellheatmap <- ggplot(heatmapdata, aes(x = Gene, y = categoryID, fill = log2foldchange)) + 
  geom_tile(color="black") + 
  theme_classic() + 
  scale_fill_continuous_divergingx(palette = 'RdBu', mid = 0, na.value="white") +
  theme(axis.text.x = element_text(angle = 45, vjust = 1, hjust = 1), axis.title.x=element_blank(), axis.title.y=element_blank()) 

singlecellheatmap + ggplot2::coord_flip()
```

```
#export 1200x400
```

### 6 Session info

Packages and versions necessary to reproduce the results in this
report.

```
sessionInfo()
```

```
## R version 4.3.1 (2023-06-16 ucrt)
## Platform: x86_64-w64-mingw32/x64 (64-bit)
## Running under: Windows 10 x64 (build 19045)
## 
## Matrix products: default
## 
## 
## locale:
## [1] LC_COLLATE=English_United States.utf8 
## [2] LC_CTYPE=English_United States.utf8   
## [3] LC_MONETARY=English_United States.utf8
## [4] LC_NUMERIC=C                          
## [5] LC_TIME=English_United States.utf8    
## 
## time zone: America/New_York
## tzcode source: internal
## 
## attached base packages:
## [1] stats4    stats     graphics  grDevices utils     datasets  methods  
## [8] base     
## 
## other attached packages:
##  [1] eulerr_7.0.0                            
##  [2] svglite_2.1.2                           
##  [3] ggraph_2.1.0                            
##  [4] colorspace_2.1-0                        
##  [5] ggpubr_0.6.0                            
##  [6] lattice_0.22-5                          
##  [7] org.Dr.eg.db_3.18.0                     
##  [8] ensembldb_2.26.0                        
##  [9] AnnotationFilter_1.26.0                 
## [10] GenomicFeatures_1.54.1                  
## [11] ChIPseeker_1.38.0                       
## [12] orthogene_1.8.0                         
## [13] plotly_4.10.3                           
## [14] BaseSet_0.9.0                           
## [15] ontologyIndex_2.11                      
## [16] enrichplot_1.22.0                       
## [17] msigdbr_7.5.1                           
## [18] clusterProfiler_4.10.0                  
## [19] gprofiler2_0.2.2                        
## [20] GSVA_1.50.0                             
## [21] GSEABase_1.64.0                         
## [22] graph_1.80.0                            
## [23] annotate_1.80.0                         
## [24] XML_3.99-0.14                           
## [25] AnnotationDbi_1.64.0                    
## [26] DT_0.30                                 
## [27] RColorBrewer_1.1-3                      
## [28] ggrepel_0.9.4                           
## [29] scales_1.2.1                            
## [30] viridis_0.6.4                           
## [31] viridisLite_0.4.2                       
## [32] cowplot_1.1.1                           
## [33] DESeq2_1.42.0                           
## [34] SummarizedExperiment_1.32.0             
## [35] Biobase_2.62.0                          
## [36] MatrixGenerics_1.14.0                   
## [37] matrixStats_1.1.0                       
## [38] GenomicRanges_1.54.1                    
## [39] GenomeInfoDb_1.38.0                     
## [40] IRanges_2.36.0                          
## [41] S4Vectors_0.40.1                        
## [42] BiocGenerics_0.48.0                     
## [43] lubridate_1.9.3                         
## [44] forcats_1.0.0                           
## [45] stringr_1.5.0                           
## [46] dplyr_1.1.3                             
## [47] purrr_1.0.2                             
## [48] readr_2.1.4                             
## [49] tidyr_1.3.0                             
## [50] tibble_3.2.1                            
## [51] ggplot2_3.4.4                           
## [52] tidyverse_2.0.0                         
## [53] knitr_1.45                              
## [54] tinytex_0.48                            
## [55] rmarkdown_2.25                          
## [56] TxDb.Hsapiens.UCSC.hg38.knownGene_3.18.0
## [57] EnsDb.Hsapiens.v75_2.99.0               
## 
## loaded via a namespace (and not attached):
##   [1] ProtGenerics_1.34.0                    
##   [2] fs_1.6.3                               
##   [3] bitops_1.0-7                           
##   [4] HDO.db_0.99.1                          
##   [5] httr_1.4.7                             
##   [6] tools_4.3.1                            
##   [7] backports_1.4.1                        
##   [8] utf8_1.2.4                             
##   [9] R6_2.5.1                               
##  [10] HDF5Array_1.30.0                       
##  [11] lazyeval_0.2.2                         
##  [12] rhdf5filters_1.14.0                    
##  [13] withr_2.5.2                            
##  [14] prettyunits_1.2.0                      
##  [15] gridExtra_2.3                          
##  [16] cli_3.6.1                              
##  [17] scatterpie_0.2.1                       
##  [18] labeling_0.4.3                         
##  [19] sass_0.4.7                             
##  [20] systemfonts_1.0.5                      
##  [21] Rsamtools_2.18.0                       
##  [22] yulab.utils_0.1.0                      
##  [23] gson_0.1.0                             
##  [24] DOSE_3.28.0                            
##  [25] plotrix_3.8-2                          
##  [26] rstudioapi_0.15.0                      
##  [27] RSQLite_2.3.2                          
##  [28] TxDb.Hsapiens.UCSC.hg19.knownGene_3.2.2
##  [29] generics_0.1.3                         
##  [30] gridGraphics_0.5-1                     
##  [31] BiocIO_1.12.0                          
##  [32] crosstalk_1.2.0                        
##  [33] vroom_1.6.4                            
##  [34] gtools_3.9.4                           
##  [35] car_3.1-2                              
##  [36] homologene_1.4.68.19.3.27              
##  [37] GO.db_3.18.0                           
##  [38] Matrix_1.6-1.1                         
##  [39] fansi_1.0.5                            
##  [40] abind_1.4-5                            
##  [41] lifecycle_1.0.4                        
##  [42] yaml_2.3.7                             
##  [43] carData_3.0-5                          
##  [44] gplots_3.1.3                           
##  [45] rhdf5_2.46.0                           
##  [46] qvalue_2.34.0                          
##  [47] SparseArray_1.2.0                      
##  [48] BiocFileCache_2.10.1                   
##  [49] grid_4.3.1                             
##  [50] blob_1.2.4                             
##  [51] promises_1.2.1                         
##  [52] crayon_1.5.2                           
##  [53] beachmat_2.18.0                        
##  [54] KEGGREST_1.42.0                        
##  [55] pillar_1.9.0                           
##  [56] fgsea_1.28.0                           
##  [57] boot_1.3-28.1                          
##  [58] rjson_0.2.21                           
##  [59] codetools_0.2-19                       
##  [60] fastmatch_1.1-4                        
##  [61] glue_1.6.2                             
##  [62] ggfun_0.1.3                            
##  [63] data.table_1.14.8                      
##  [64] vctrs_0.6.4                            
##  [65] png_0.1-8                              
##  [66] treeio_1.26.0                          
##  [67] gtable_0.3.4                           
##  [68] cachem_1.0.8                           
##  [69] xfun_0.41                              
##  [70] S4Arrays_1.2.0                         
##  [71] mime_0.12                              
##  [72] tidygraph_1.2.3                        
##  [73] SingleCellExperiment_1.24.0            
##  [74] interactiveDisplayBase_1.40.0          
##  [75] ellipsis_0.3.2                         
##  [76] nlme_3.1-163                           
##  [77] ggtree_3.10.0                          
##  [78] bit64_4.0.5                            
##  [79] progress_1.2.2                         
##  [80] filelock_1.0.2                         
##  [81] bslib_0.5.1                            
##  [82] irlba_2.3.5.1                          
##  [83] KernSmooth_2.23-22                     
##  [84] DBI_1.1.3                              
##  [85] tidyselect_1.2.0                       
##  [86] bit_4.0.5                              
##  [87] compiler_4.3.1                         
##  [88] curl_5.1.0                             
##  [89] xml2_1.3.5                             
##  [90] DelayedArray_0.28.0                    
##  [91] shadowtext_0.1.2                       
##  [92] rtracklayer_1.62.0                     
##  [93] caTools_1.18.2                         
##  [94] rappdirs_0.3.3                         
##  [95] digest_0.6.33                          
##  [96] XVector_0.42.0                         
##  [97] htmltools_0.5.6.1                      
##  [98] pkgconfig_2.0.3                        
##  [99] sparseMatrixStats_1.14.0               
## [100] highr_0.10                             
## [101] dbplyr_2.4.0                           
## [102] fastmap_1.1.1                          
## [103] rlang_1.1.1                            
## [104] htmlwidgets_1.6.2                      
## [105] shiny_1.7.5.1                          
## [106] DelayedMatrixStats_1.24.0              
## [107] farver_2.1.1                           
## [108] jquerylib_0.1.4                        
## [109] jsonlite_1.8.7                         
## [110] BiocParallel_1.36.0                    
## [111] GOSemSim_2.28.0                        
## [112] BiocSingular_1.18.0                    
## [113] RCurl_1.98-1.12                        
## [114] magrittr_2.0.3                         
## [115] GenomeInfoDbData_1.2.11                
## [116] ggplotify_0.1.2                        
## [117] patchwork_1.1.3                        
## [118] Rhdf5lib_1.24.0                        
## [119] munsell_0.5.0                          
## [120] Rcpp_1.0.11                            
## [121] ggnewscale_0.4.9                       
## [122] ape_5.7-1                              
## [123] babelgene_22.9                         
## [124] stringi_1.8.1                          
## [125] zlibbioc_1.48.0                        
## [126] MASS_7.3-60                            
## [127] AnnotationHub_3.10.0                   
## [128] plyr_1.8.9                             
## [129] parallel_4.3.1                         
## [130] HPO.db_0.99.2                          
## [131] Biostrings_2.70.1                      
## [132] graphlayouts_1.0.1                     
## [133] splines_4.3.1                          
## [134] hms_1.1.3                              
## [135] polylabelr_0.2.0                       
## [136] locfit_1.5-9.8                         
## [137] igraph_1.5.1                           
## [138] ggsignif_0.6.4                         
## [139] reshape2_1.4.4                         
## [140] biomaRt_2.58.0                         
## [141] ScaledMatrix_1.10.0                    
## [142] BiocVersion_3.18.0                     
## [143] evaluate_0.23                          
## [144] BiocManager_1.30.22                    
## [145] tzdb_0.4.0                             
## [146] tweenr_2.0.2                           
## [147] httpuv_1.6.12                          
## [148] grr_0.9.5                              
## [149] polyclip_1.10-6                        
## [150] ggforce_0.4.1                          
## [151] rsvd_1.0.5                             
## [152] broom_1.0.5                            
## [153] xtable_1.8-4                           
## [154] restfulr_0.0.15                        
## [155] tidytree_0.4.5                         
## [156] MPO.db_0.99.7                          
## [157] rstatix_0.7.2                          
## [158] later_1.3.1                            
## [159] snow_0.4-4                             
## [160] aplot_0.2.2                            
## [161] GenomicAlignments_1.38.0               
## [162] memoise_2.0.1                          
## [163] timechange_0.2.0
```
